# Multivariate Analyses of Codon Usage of SARS-CoV-2 and other betacoronaviruses

**DOI:** 10.1101/2020.02.15.950568

**Authors:** Haogao Gu, Daniel Chu, Malik Peiris, Leo L.M. Poon

## Abstract

Coronavirus disease 2019 (COVID-19) is a global health concern as it continues to spread within China and beyond. The causative agent of this disease, severe acute respiratory syndrome coronavirus 2 (SARS-CoV-2), belongs to the genus *Betacoronavirus* which also includes severe acute respiratory syndrome related coronavirus (SARSr-CoV) and Middle East respiratory syndrome related coronavirus (MERSr-CoV). Codon usage of viral genes are believed to be subjected to different selection pressures in different host environments. Previous studies on codon usage of influenza A viruses can help identify viral host origins and evolution trends, however, similar studies on coronaviruses are lacking. In this study, global correspondence analysis (CA), within-group correspondence analysis (WCA) and between-group correspondence analysis (BCA) were performed among different genes in coronavirus viral sequences. The amino acid usage pattern of SARS-CoV-2 was generally found similar to bat and human SARSr-CoVs. However, we found greater synonymous codon usage differences between SARS-CoV-2 and its phylogenetic relatives on spike and membrane genes, suggesting these two genes of SARS-CoV-2 are subjected to different evolutionary pressures.

## Introduction

A novel coronavirus outbreak took place in Wuhan, Hubei province, China in December 2019^1^. This novel coronavirus (SARS-CoV-2) causes pneumonia in patients^2^ and it has rapidly spread to other provinces in China and other countries ^3^. This novel coronavirus outbreak had raised global concern but current knowledge on the origin and transmission route of the pathogen is still limited. The SARS-CoV-2 belongs to the genus *Betacoronavirus*, which also includes two highly virulent human coronaviruses, SARS-CoV and MERS-CoV. Apart from human, many animal species, such as bat, rat, camel, swine and hedgehog, can be infected by different types of coronaviruses. Further sequence analyses of this novel and other betacoronaviruses might provide additional information to better understand the evolution of SARS-CoV-2.

Preferential codon usage is commonly seen in different organisms, and it has been evident that the uneven codon usage is not neutral but related to gene expression or other selection pressures ^4–6^. There are two levels of codon usage biases, one is at amino acid level and the other is at synonymous codon level. The amino acid composition of proteins can be an important factor that explaining certain sequence traits. For example integral membrane proteins that are enriched in hydrophobic amino acids can create significant codon usage bias^7^. Amino acid composition sometime can also introduce confounding effects when one only focuses on studying the variations of synonymous codon usage. The use of global correspondence analysis (CA) and its derivatives within-group correspondence analysis (WCA) and between-group correspondence analysis (BCA) to analyze codon usages can overcome the above problem. In fact, WCA becomes “model of choice” for analyzing synonymous codon usage in recent years, as it is more robust than other traditional methods (e.g. CA with relative codon frequency or CA with RSCU values)^7,8^. This analytic approach, however, has not been used in studying viral sequences. As the natural history of the SARS-CoV-2 remains largely unknown, an in-depth codon usage analysis of this newly emerging virus might provide some novel insights.

In this study, we used both CA and WCA to analyses codon usage patterns of a vast number of betacoronavirus sequences. We found SARS-CoV-2 and bat SARSr-CoV have similar amino acid usage. However, our analyses suggested that the spike and member genes of SARS-CoV-2 have rather distinct synonymous codon usage patterns.

## Methods

### Sequence data

To construct a reference sequence dataset, available full-length complete genome sequences of coronavirus were collected through Virus Pathogen Resource database (https://www.viprbrc.org/brc/home.spg?decorator=corona, accessed 13 Jul 2019, ticket 958868915368). The sequences were filtered by the following steps: (1) Remove sequences without protein annotation, (2) Keep only sequences with complete set of desired replicase and structural proteins (sequences coding for orf1ab, spike, membrane and nucleocapsid), (3) Filter out sequences that are unusually long and short (>130% or <70% of the median length for each group of gene sequences), (4) Limit our analysis to genus *Betacoronavirus* and (5) Concatenate orf1a and orf1b sequences to form orf1ab if necessary.

The final dataset comprised 769 individual strains (3076 individual gene sequences) that contain complete sets of coding regions for orf1ab, spike, membrane and nucleocapsid genes (see Supplementary Figure 1). The sequences for envelope gene were not included in the analysis because of the short length and potential bias in codon usage. Corresponding metadata for the sequences were extracted by the sequence name field. 24 complete genome sequences of the newly identified SARS-CoV-2 and its phylogenetically close relatives were retrieved from Genbank and GISAID (accessed 22 Jan 2020). Six genomes in this study were used as special references (BetaCoV/bat/Yunnan/RaTG13/2013|EPI_ISL_402131; BetaCoV/pangolin/Guangxi/P1E/2017|EPI_ISL_410539; MG772934.1_Bat_SARS-like_coronavirus_isolate_bat-SL-CoVZXC21; MG772933.1_Bat_SARS-like_coronavirus_isolate_bat-SL-CoVZC45; KY352407.1_Severe_acute_respiratory_syndrome-related_coronavirus_strain_BtKY72 and GU190215.1_Bat_coronavirus_BM48-31/BGR/2008), as they have previously been reported to have close phylogenetic relationship with SARS-CoV-2^9–11^. Detailed accession ID for the above data are provided in the Supplementary Table S1.

The codon count for every gene sequence input for the correspondence analysis was calculated by the SynMut^12^ package. The implementation of the different correspondence analyses in this study was performed by functions in the package ade4^13^. Three stop codons (TAA, TAG and TGA) were excluded in the correspondence analysis.

### Global correspondence analysis (CA) on codon usage

Correspondence analysis (CA) is a dimension reduction method which is well suited for amino acid and codon usage analysis. The concept in correspondence analysis is similar to Pearson’s χ^2^ test (i.e., the expected counts are calculated under the hypothesis of independence, based on the observed contingency table). With the deduced expected count table, the Euclidean distance or the χ^2^ distance can be used to evaluate the difference between two observations. The χ^2^ distance that we are using in the global correspondence analysis is applied for the row profile (adjusted for the size effect among difference genes) and the column profile (adjusted for the size effect among difference codons) and therefore the raw codon count rather than the Relative Synonymous Codon Usage (RSCU) values are more informative and suitable input for our model. The calculation of the χ^2^ distance is included in the Supplementary Method.

All the correspondence analyses in this study were performed individually for each gene, to achieve better resolution on gene specific codon usage pattern.

### Within-group correspondence analysis and between-group correspondence analysis

In contrast to the ordinary correspondence analysis, the within-block correspondence analysis^14^ (WCA) can segregate the effects of different codon compositions in different amino acids. WCA has been recognized as the most accurate and effective CA method for studying the synonymous codon usage in various genomic profile^8^. WCA focuses on the within-amino acid variability, and it technically excludes the variation of amino acid usage differences. WCA was implemented based on the existing global CA, with additional information for factoring.

Between-group correspondence analysis (BCA) is complementary to WCA; BCA focuses on the between-group variability. BCA can be interpreted as the CA on amino acid usage. We used BCA in this study to investigate the amino acid usage pattern in different coronaviruses.

### Grand Average of Hydropathy (GRAVY) score

Gravy score provides an easy way to estimate the hydropathy character of a protein^15^. It was used in this study as a proxy to identify proteins that are likely to be membrane-bound proteins. The GRAVY score was calculated in a linear form on codon frequencies as:

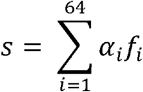

Where *α*_*i*_ is the coefficient for a particular amino acid (provided by data *EXP* in *Seqinr* package^16^) encoded by codon *i*, *f*_*i*_ correspond to the relative frequency of codon *i*.

## Results

### General sequence features in *Betacoronavirus*

A total of 3,076 individual gene sequences passed the filtering criteria and were included in this study. Viral sequences from 3 different species (*Middle East respiratory syndrome related coronavirus* (*MERSr-CoV*), *Betacoronavirus 1*, *SARS related coronavirus* (*SARSr-Cov*)) were the three most dominant species (see Supplementary Figure S1) in the filtered dataset.

Four conserved protein sequence encoding regions of *Betacoronavirus* were analysed separately. The median lengths of the studied sequence regions were 21237 nt for orf1ab gene, 4062 nt for spike gene, 660 nt for membrane gene and 1242 nt for nucleocapsid gene. Spike gene has the lowest average and median G + C contents among these four genes (median: 37.45%, 37.31%, 42.60% and 47.22% for orf1ab, spike, membrane and nucleocapsid respectively). The G +C contents of the orf1ab and spike genes were found distributed in bi-modal patterns, and the G + C contents of SARS-CoV-2 were found located at the lesser half of the data of these two genes. The G + C contents for membrane and nucleocapsid genes of studied viral sequences were distributed in unimodal pattern (see Supplementary Figure S2).

The overall amino acid and codon usage of the dataset are plotted in an ascending order (Figure 1). We observed that leucine and valine were the two most frequently used amino acids in the four studied genes, while tryptophan, histidine and methionine were the three least used ones. We also found that codons ending with cytosine or guanine were generally less frequent than the codons ending with adenine or thymine. This pattern of uneven usage in synonymous codons is in accordance with the G + C content distribution results (codons ending with guanine or cytosine were less frequently observed).

**Figure 1.**
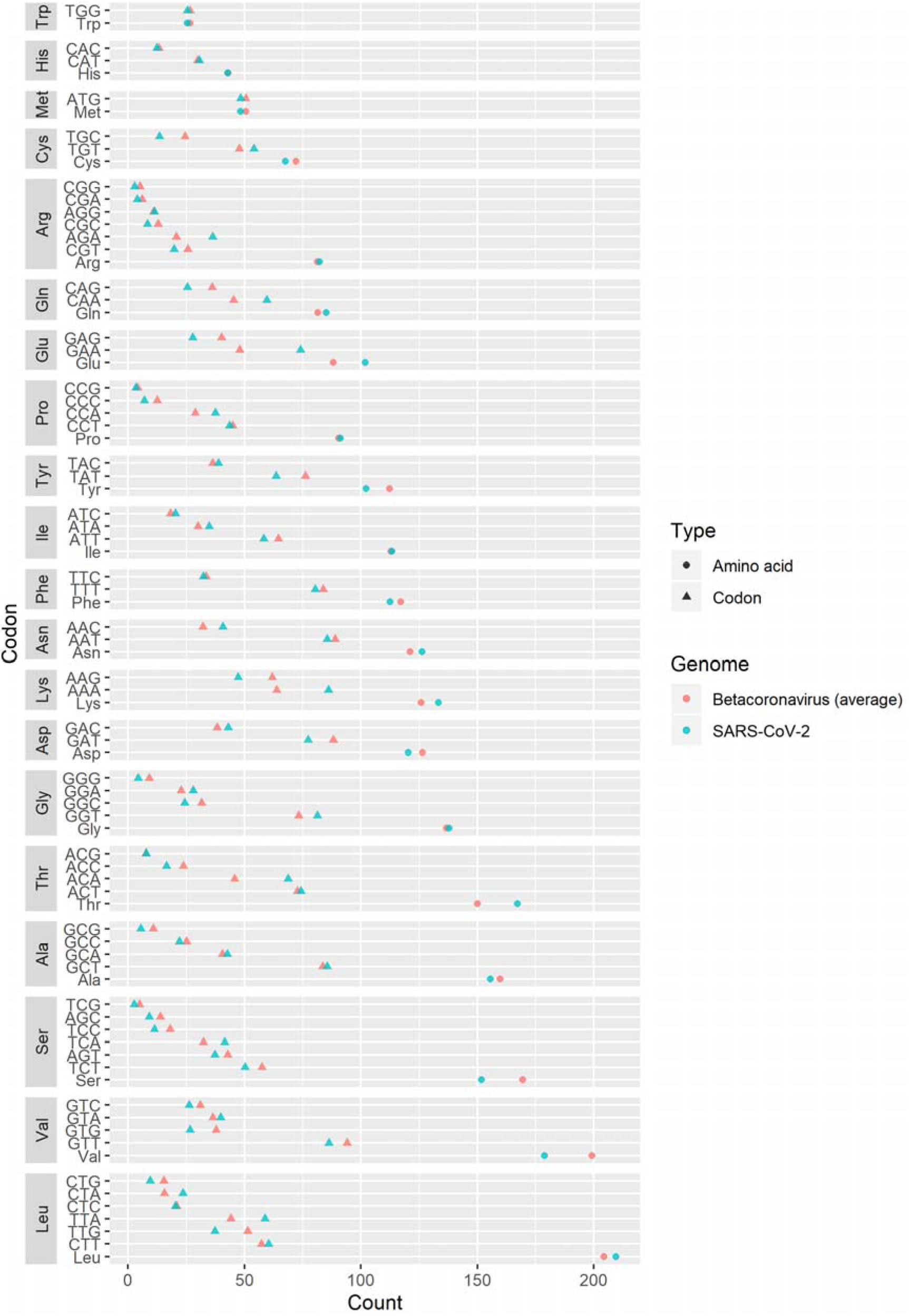
Codon usage in *Betacoronavirus* (Cleveland’s dot plot). Points in green showed the count of codons in a sample SARS-CoV-2 genome (MN908947).

We found a substantial bias in amino acid usage among these four genes, and this bias is well explained by the hydropathy of the encoded proteins (results from global correspondence analysis on all the four genes, collectively, data not shown). The GRAVY scores for every sequence were calculated to represent the degree of hydropathy. We discovered that the nucleocapsid protein sequences had significantly lower GRAVY scores as compared to those from other genes, while the membrane protein sequences had highest GRAVY scores (see Supplementary Figure S3).

### Correspondence analysis

We first conducted a multivariate analysis of codon usage on the dataset by using global correspondence analysis. We also conducted WCA and BCA to study these sequences at synonymous codon usage and amino acid usage levels, respectively. Given that there were different amino acid usage biases among different genes (Supplementary Figure S3), we performed correspondence analyses of these genes separately.

Of all the four correspondence analyses for the four genes, the extracted first factors explained more than 50% of the total variance (see Supplementary Figure S4). The first two factors in orf1ab global CA represented 67.7% and 16.8% of total inertia. Similarly, the first two factors of the spike, membrane and nucleocapsid global CA represented 51.0% and 18.5%, 52.6% and 20.2%, and 54.8% and 14.2%, respectively, of total inertia. With only these two factors, we could extract ~70% of the variability of the overall codon usage for each studied gene. These levels of representations were higher than or similar to those deduced from other codon usage analyses^8,17,18^.

### The overall codon usage of SARS-CoV-2 in orf1ab, spike and membrane genes are similar to those of bat and pangolin CoVs

Based on the above CA analysis, the data points are shown in different colours that represent different features of the sequences (e.g. viral host or viral species). There were no neighbouring human viruses around SARS-CoV-2 in CA results of orf1ab, spike and membrane (Figure 2), suggesting that the overall codon usage of SARS-CoV-2 in the orf1ab, spike or membrane gene was significantly different from those of human betacoronaviruses. By contrast, the nucleocapsid genes of SARS coronavirus and SARS-CoV-2 are found to be relatively similar (Supplementary Figure S5A). Except for the nucleocapsid gene, virus sequences adjacent to the SARS-CoV-2 were all from bat coronaviruses (coloured in purple in Figure 2).

**Figure 2.**
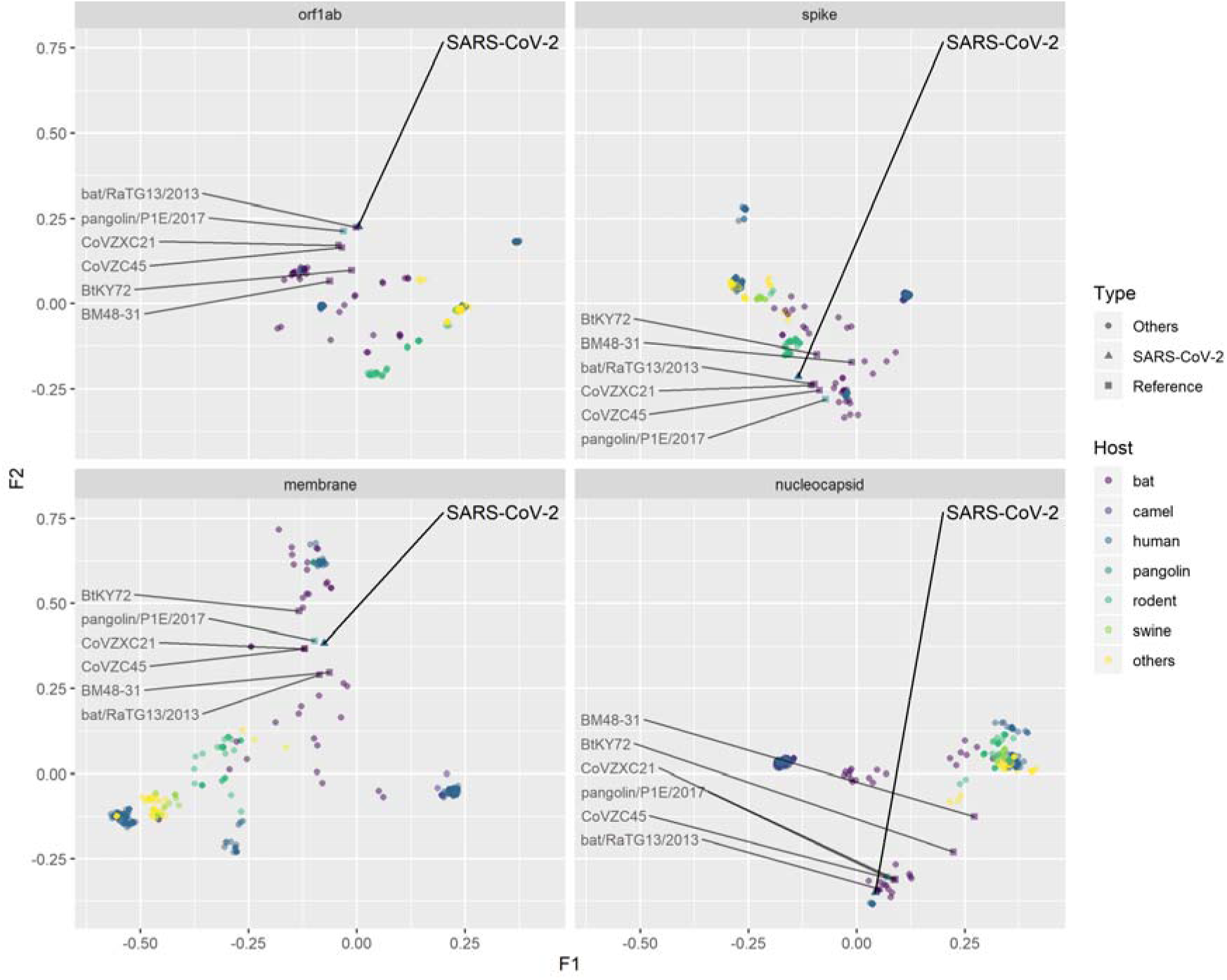
Factorial map of the first and second factors for global CA by different genes, coloured by different viral host. The SARS-CoV-2 and related reference data points were labelled.

There are five groups of viral sequences of human origin in the dataset (SARS-CoV-2, Betacoronavirus 1, human coronavirus HKU 1, MERS-CoV and SARS-CoV). These five groups of viral sequences were well separated from each other in terms of codon usage, except the nucleocapsid gene sequences of SARS-CoV-2 and SARS-CoV as mentioned above. There was no overlap between SARS-CoV-2 and human SARS-CoV in orf1ab, spike and membrane, yet SARS-CoV codon usage processed more similar to SARS-CoV-2 compared to the other three types of human coronaviruses (i.e. yellow point always closest to SARS-CoV-2 in Supplementary Figure S5A).

Compared to human coronavirus sequences, the bat coronavirus sequences have more scattered codon usage, even within the same viral species (Supplementary S5B). Some viral species in bats formed their own clusters in all four genes (e.g. SARSr-CoV). SARSr-CoV is a group of coronavirus that can be found in both humans and bats. We observed that the data points of human SARSr-CoV are clustered with those of bat SARSr-CoV in all the four genes (by comparing the yellow points in Supplementary Figure S5A and S5B). The codon usage of SARS-CoV-2 in orf1ab, spike and membrane were slightly different from the SARS-CoV clusters and these data points are located in between SARSr-CoV and other coronavirus species (e.g. MERSr-CoV and bat coronavirus HKU9 etc.)

The global codon usages of bat RatG13 virus were found most similar to SARS-CoV-2 in orf1ab, spike and nucleocapsid genes, but not in membrane gene (Figure. 2). In the analysis of membrane protein, pangolin P1E virus had a more similar codon usage to SARS-CoV-2 than all the other viruses. We found the similarity in codon usage between pangolin P1E and SARS-CoV-2 were also high in orf1ab, where P1E was the second closest data point to SARS-CoV-2. But this is not the case for spike and nucleocapsid genes.

We also observed that the codon usage pattern in spike gene was more complex than in other genes. For example, data points adjacent to the spike gene of SARS-CoV-2 were coronaviruses from bat, human and rodent hosts (Figure 2). The codon usage of rodent coronaviruses was generally distinct from human or bat coronaviruses in orf1ab, membrane and nucleocapsid gene sequences. By contrast, the spike gene sequences of murine coronaviruses were found located between SARSr-CoV and other coronaviruses, just like SARS-CoV-2 (Figure 2 and Supplementary Figure S6B). The codon usage from camel, swine and other coronaviruses were found to be well clustered and relatively distant to SARS-CoV-2 (see Supplementary Figure S6A, S5C, S5D).

### The codon usage at synonymous level suggested novel patterns of SARS-CoV-2 in spike and membrane genes

WCA and BCA were used to further differentiate codon usage of these betacoronaviruses at synonymous codon usage and amino acid usage levels, respectively. After applying the row-block structure to the original global CA model, we found that most of the variability in codon usage can be explained at synonymous codon usage level (90.36% for orf1ab gene, 85.29% for spike gene, 83.71% for member gene and 84.07% for nucleocapsid gene) (Table 1).

**Table 1.**
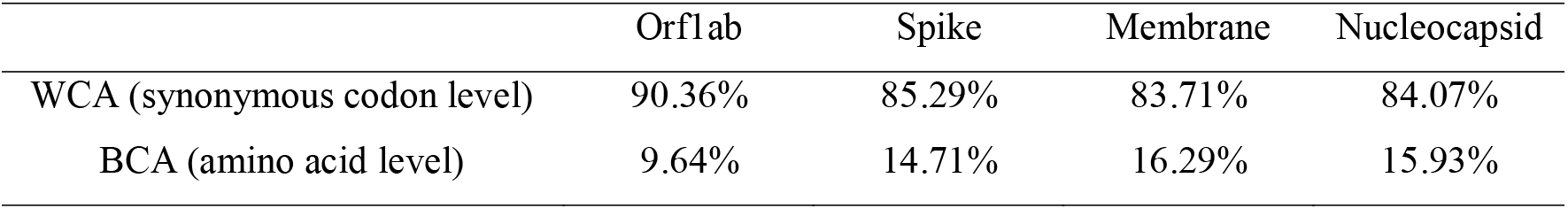
Variability explained by the synonymous codon usage level and the amino acid level.

|  | Orf1ab | Spike | Membrane | Nucleocapsid |
| --- | --- | --- | --- | --- |
| WCA (synonymous codon level) | 90.36% | 85.29% | 83.71% | 84.07% |
| BCA (amino acid level) | 9.64% | 14.71% | 16.29% | 15.93% |

Results from the BCA suggested that the amino acid usage of SARS-CoV-2 is closely related to bat and human SARSr-CoVs in all four genes (Figure 3B and Figure 4B). Specifically, we discovered that the SARS-CoV-2 had amino acid usage pattern most similar to bat RaTG13 virus, followed by pangolin P1E, bat CovVZC45 and bat CoVZXC21. The sequences of BtKY72 and BM48-31 were from a more phylogenetically distant clade, and, accordingly, they had relatively distinct amino acid usage to SARS-CoV-2 as expected in all four studied genes. This result agrees with the result in the full-genome phylogenetic analysis (Supplementary Figure S7).

**Figure 3.**
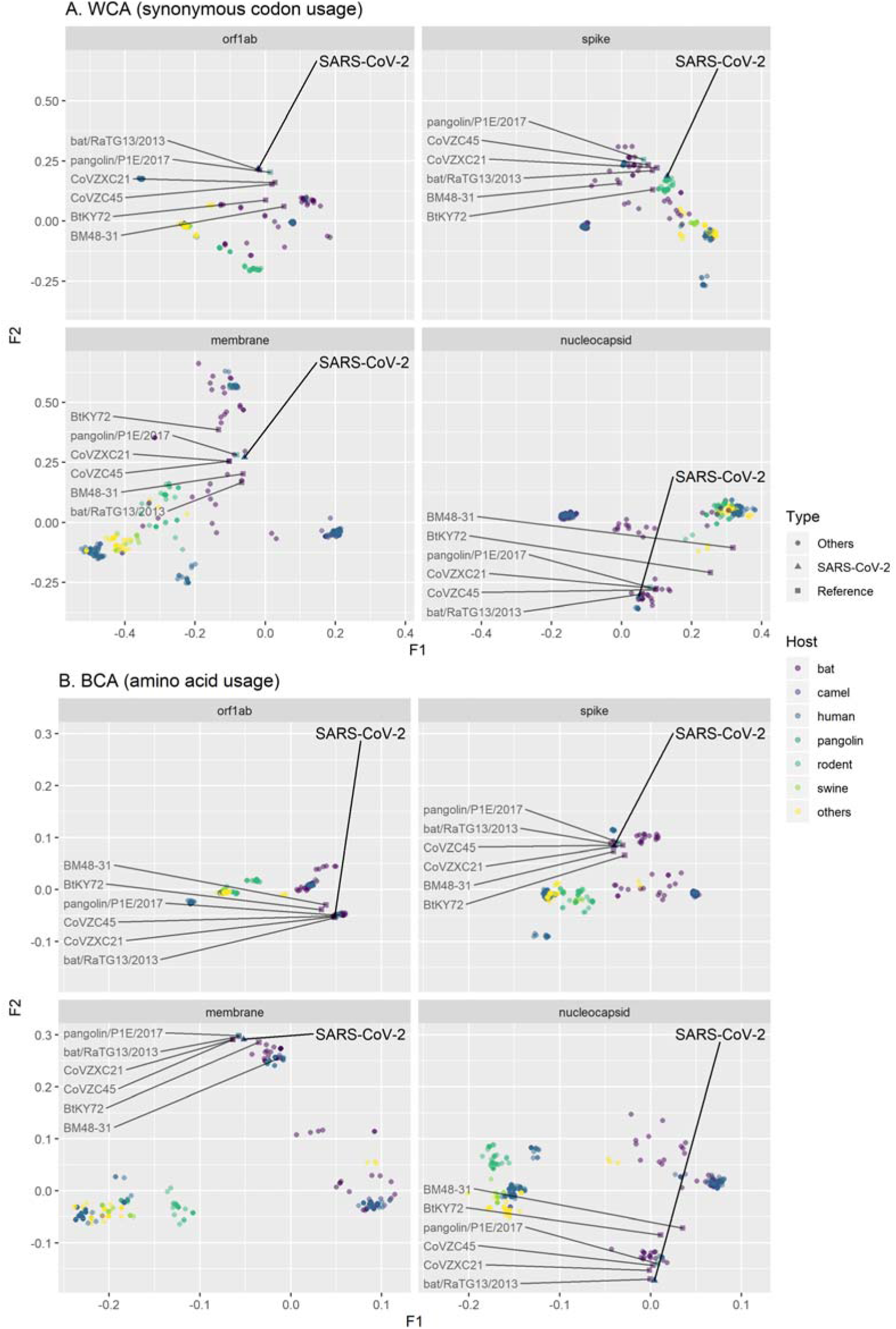
Factorial map of the first and second factors for WCA and BCA by different genes, coloured by different viral host. The SARS-CoV-2 and related reference data points were labelled.

**Figure 4.**
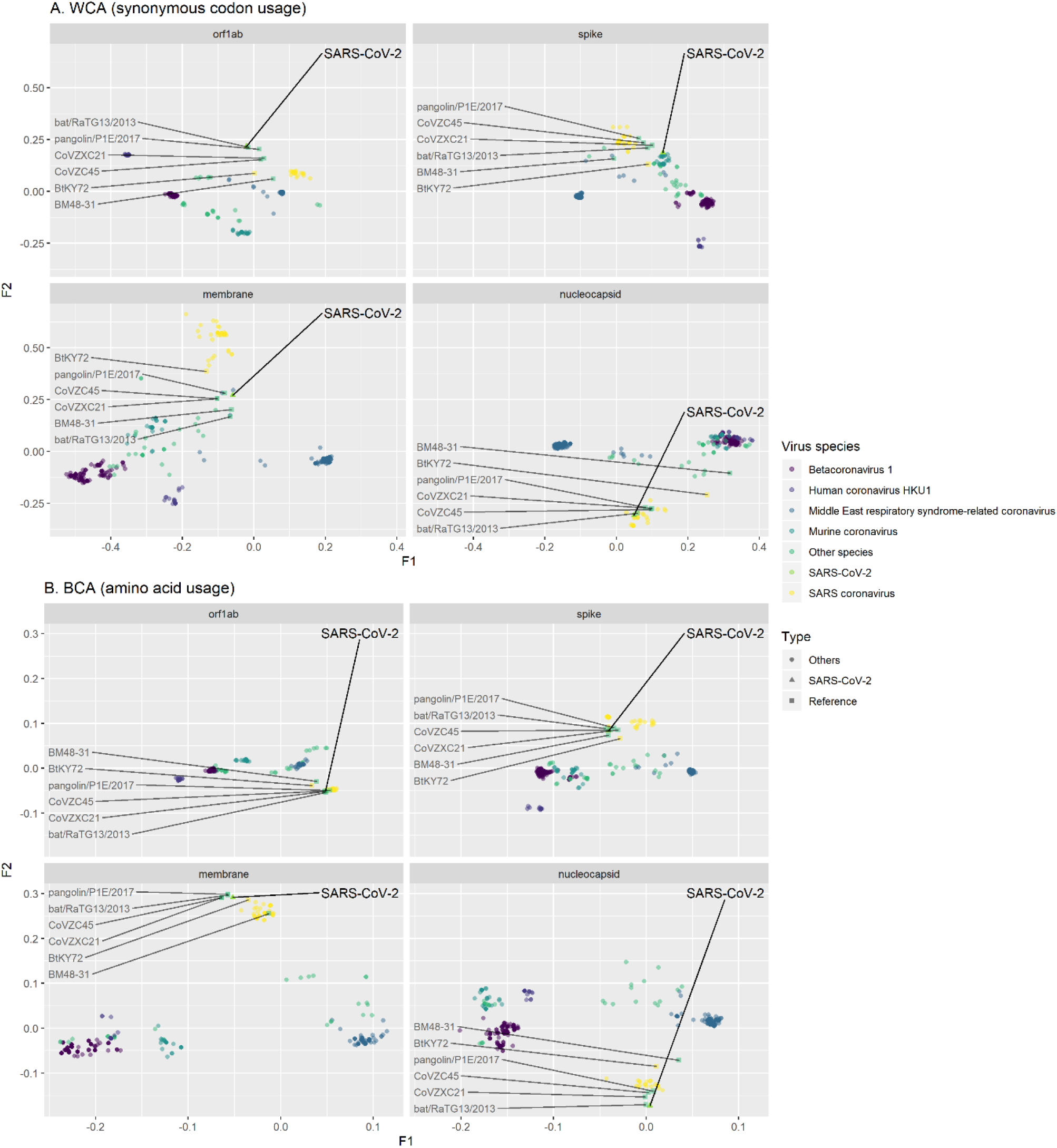
Factorial map of the first and second factors for WCA and BCA by different genes, coloured by different viral species. The SARS-CoV-2 and related reference data points were labelled.

The difference between SARS-CoV-2 and RaTG13 at synonymous codon usage level was marginal in orf1ab and nucleocapsid sequences. Interestingly, there were noticeable differences in the spike and membrane gene analyses. Our results suggest the synonymous codon usage patterns in the spike and membrane gene of SARS-CoV-2 are different from those of its genetically related viruses (i.e. RaTG13 and other reference relatives). For example, the synonymous codon usage pattern of SARS-CoV-2 was found to be closer to a cluster of rodent murine coronaviruses at the first two factorial levels (Figure 3A and Figure 4A).

Further analysis on spike gene, however, suggested that the codon usage of SARS-CoV-2 and rodent murine coronaviruses were distinct at the third factorial level (Supplementary Figure S8A). The results show that although RaTG13 was not the point most adjacent to SARS-CoV-2 at the first and second dimension, it surpassed murine coronaviruses at the third dimension. Our results suggest a complex genomic background in the spike gene of SARS-CoV-2, which made its synonymous codon usage harder to differentiate from other genomic sequences in our WCA analysis. Despite the proximity between RaTG13 and SARS-CoV-2 at three-dimensional level, they were still formed into two separated clusters (Supplementary Figure S8A). It is evident that the synonymous codon usage pattern of SARS-CoV-2 is distinct from other bat origin coronaviruses. The difference in synonymous codon usage is largely explained by the first factor (more than 50%), and our analysis on codon usages suggest that the first factor maybe highly related to the preferential usage of codons ending with cytosine (Supplementary Figure S9). We also had similar observation for the membrane gene. Our three-dimensional analysis revealed that the synonymous codon usage of SARS-CoV-2 in membrane was most similar to P1E and CoVZXC21 (Supplementary Figure S8B). It is worth noting that comparing to RaTG13, P1E and CoVZXC21 had lower synonymous codon usage similarity to SARS-CoV-2 in the other three genes.

Overall, our WCA results support a more complex synonymous codon usage background on spike and membrane genes, though we identified unique codon usage patterns of SARS-CoV-2 on these two genes.

## Discussion

Codon usage can be affected by many sequence features, including nucleotide composition, dinucleotide composition, amino acid preference, host adaption, etc^8,19,20^. The codon usages of viral sequences can vary by genes and host origins^21–23^. The bias in codon usage is a unique and distinctive characteristic that can reflect the “signature” of a genomic sequence. Codon usage analyses are often complementary to ordinary sequence alignment-based analyses which focus on the genetic distance at nucleotide level, whereas codon usage analyses enable capturing signals at different sequence parameters. Therefore, codon usage bias can be another good proxy for identifying unique traits (e.g. virus origin, host origin, or some functions of proteins) of a genome. The goal of this study was to investigate the codon usage bias of betacoronaviruses. By studying the codon usags of these viruses in a systematic manner, we identified viral sequences carrying traits similar to those of SARS-CoV-2, which provided useful information for studying the host origin and evolutionary history of SARS-CoV-2.

The codon usage of different genes in betacoronaviruses are very different. The G+C content, especially the GC3 content is known to be influential to the codon usage of some bacteria and viruses ^7,24,25^. The GC3 content has pronounced effects on our WCA analysis of the orf1ab and spike genes. The GC3 content was found correlated with high WCA values on the first factor of orf1ab (Supplementary Figure S9). By contrast, codons ending with cytosine had lower factorial values in the spike gene analysis (Supplementary Figure S9). The G + C contents in membrane and nucleocapsid genes were less suppressed (Supplementary Figure S2). This can be partly explained by the fact that membrane and nucleocapsid are two genes with shorter lengths which may limit the flexibilities for mutation or codon usage adaptation.

In addition to global CA analysis, the application of WCA and BCA can eliminate the effects caused by amino acid compositions and synonymous codon usage, respectively. These alternative analytical tools were important to our study. It is because the amino acid sequences are expected to be more conserved such that they can preserve biological functions of the translated genes. By contrast, mutations at synonymous level tend to be more frequent, as most of these codon alternatives do not affect the biological function of a protein.

Of all the existing genomes in the dataset, RaTG13 best matched the overall codon usage pattern of the SARS-CoV-2. Although the SARS-CoV-2 had amino acid usage similar to bat and human SARSr-CoVs, the synonymous codon usages between them were relatively different, which indicates similar protein characteristics but maybe different evolutionary histories. The codon usage of bat coronaviruses are more scattered than coronaviruses of other hosts. This result agrees with the fact that bat is a major host reservoir of coronavirus^26^, thus it harbours coronaviruses with more complex genomic backgrounds.

SARS-CoV-2 was first identified in human, but its codon usage pattern is very different from those of other human betacoroanviruses (Supplementary Figure S5A). In fact, the codon usage at both the amino acid level and synonymous level denote that the orf1ab gene in SARS-CoV-2 had closest relationship to SARSr-CoV, especially RaTG13. The CoVZX45 and CoVZXC21 had similar amino acid usage but relatively different synonymous codon usage to SARS-CoV-2 (Figure 3). Besides bat-origin SARSr-CoV, the pangolin P1E also had similar codon usage to SARS-CoV-2 both at amino acid and synonymous codon levels. The result in orf1ab is in accordance with the full-genome phylogenetic analysis (Supplementary Figure S7), showing a close relationship between SARS-CoV-2 and RaTG13 by the overall backbone of the genome.

The S protein is responsible for receptor binding which is important for viral entry. The genetic variability is extreme in spike gene^27^, and this highly mutable gene may possess valuable information about recent evolution history. In our results, the synonymous codon usage of SARS-CoV-2 in spike gene was distinct from those of RaTG13 and other phylogenetic relatives (Figure 3A), which was not observed in orf1ab or nucleocapsid gene. Although the codon usage in spike of SARS-CoV-2, RaTG13 and P1E were similar at amino acid level, the difference at synonymous codon usage level indicates that they are unlikely to share a very recent common ancestor. It is more likely that SARS-CoV-2, RaTG13 and P1E might have undergone different evolution pathways for a certain period of time. The amino acid usage of SARS-CoV-2 in membrane was clustered with bat SARSr-CoV, however the synonymous codon usage of SARS-CoV-2 was still distinct to these bat coronaviruses. Notably, in membrane gene, pangolin P1E had a more similar synonymous codon usage to SARS-CoV-2 than RaTG13. These findings suggest that there may be different selection forces between genes. Our result supports different evolutionary background or currently unknown host adaption history in SARS-CoV-2. The codon usage of SARS-CoV-2 in nucleocapsid gene was similar to bat SARSr-CoV both at amino acid level and synonymous level, suggesting that no highly significant mutation happened in this gene.

Codon usage can be shaped by many different selection forces, including the influence from host factors. Some researchers have hypothesised that the codon usage in SARS-CoV-2 maybe directly correlated to the codon usage of its host^28^. However our recent study on influenza A viruses implied that these may not be the most influential factors shaping the codon usage of a viral genome^19^. Our analysis took advantage of the existing genomes of *Betacoronavirus* to study the complex host effect on codon usage, which warrants more accurate but relatively conserved estimation.

## Acknowledgements

We thank researchers for making viral sequences available for public access. We gratefully acknowledge the Originating and Submitting Laboratories for sharing genetic sequences and other associated data through the GISAID Initiative, on which this research is based. A list of the authors can be found in Supplementary Table S2.

