## Supplementary Information for "Multivariate Analyses of Codon Usage of SARS-CoV-2 and other betacoronaviruses"

**Supplementary Method**

The $\chi^{2}$ distance^1^ was calculated as:

$$d^{2}\left( X_{i},X_{i^{'}} \right)= n_{++}\sum_{j=1}^{J} {(\frac{r_{ij}}{\sqrt{n_{+j}}}- \frac{r_{i^{'}j}}{\sqrt{n_{+j}}})}^{2}$$

$$r_{ij}= \frac{n_{ij}}{n_{i+}}$$

$$r_{i^{'}j}= \frac{n_{i^{'}j}}{n_{i^{'}+}}$$

where $X_{i}$ is the $i^{th}$ row variable in the $i\times j$ contingency table. The count of the $j^{th}$ codon for the $i^{th}$ sequence is noted $n_{ij}$. The row and column marginal totals are noted $n_{i+}$ and $n_{+j}$ respectively. The total number of observations is $n_{++}$.

**Supplementary Figures**

Figure S1. Count of different virus species in the database.

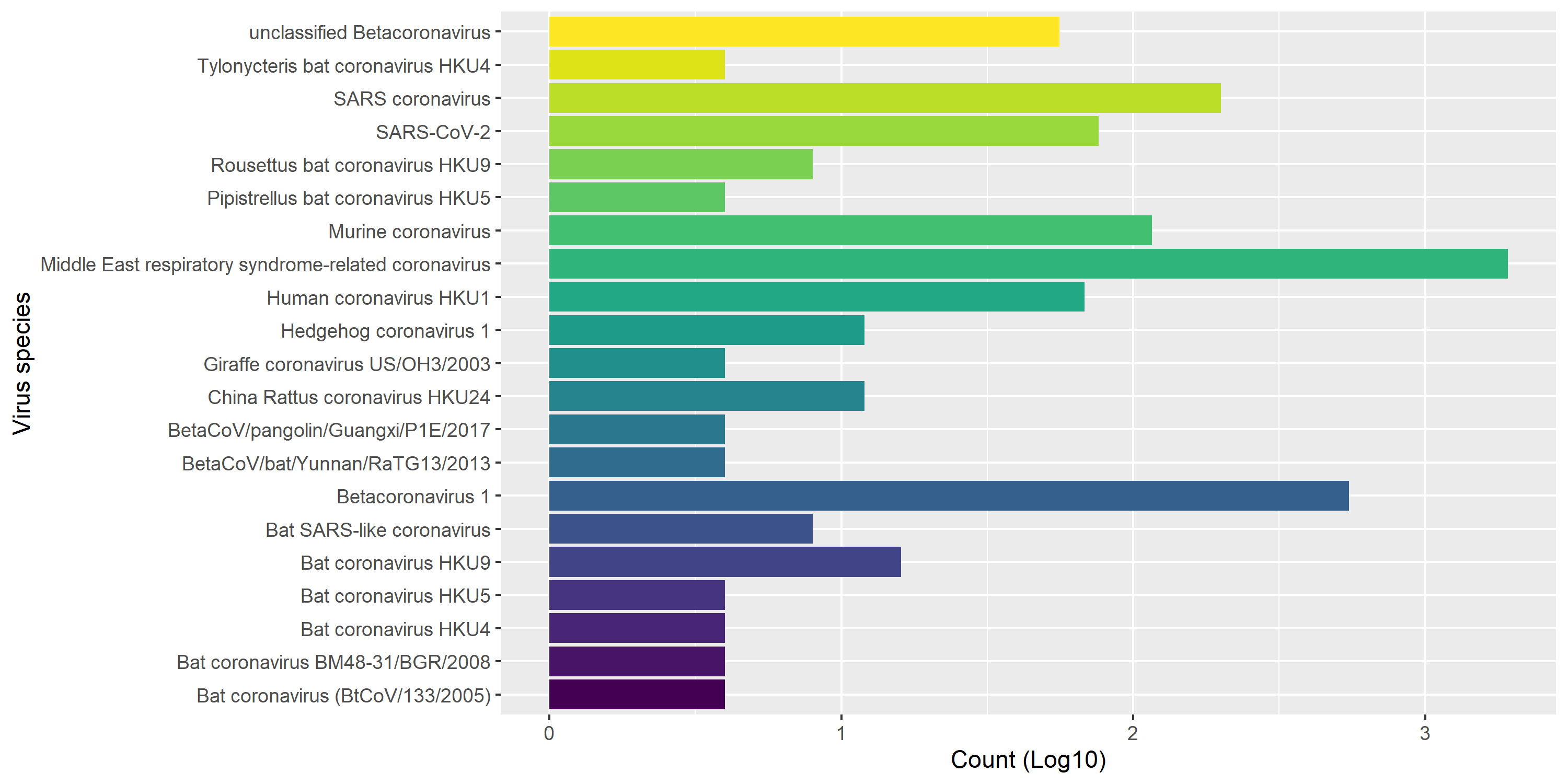

Figure S2. Histogram of G + C content by different gene in *Betacoronavirus.* The G + C content values of SARS-CoV-2 were plotted separately in red.

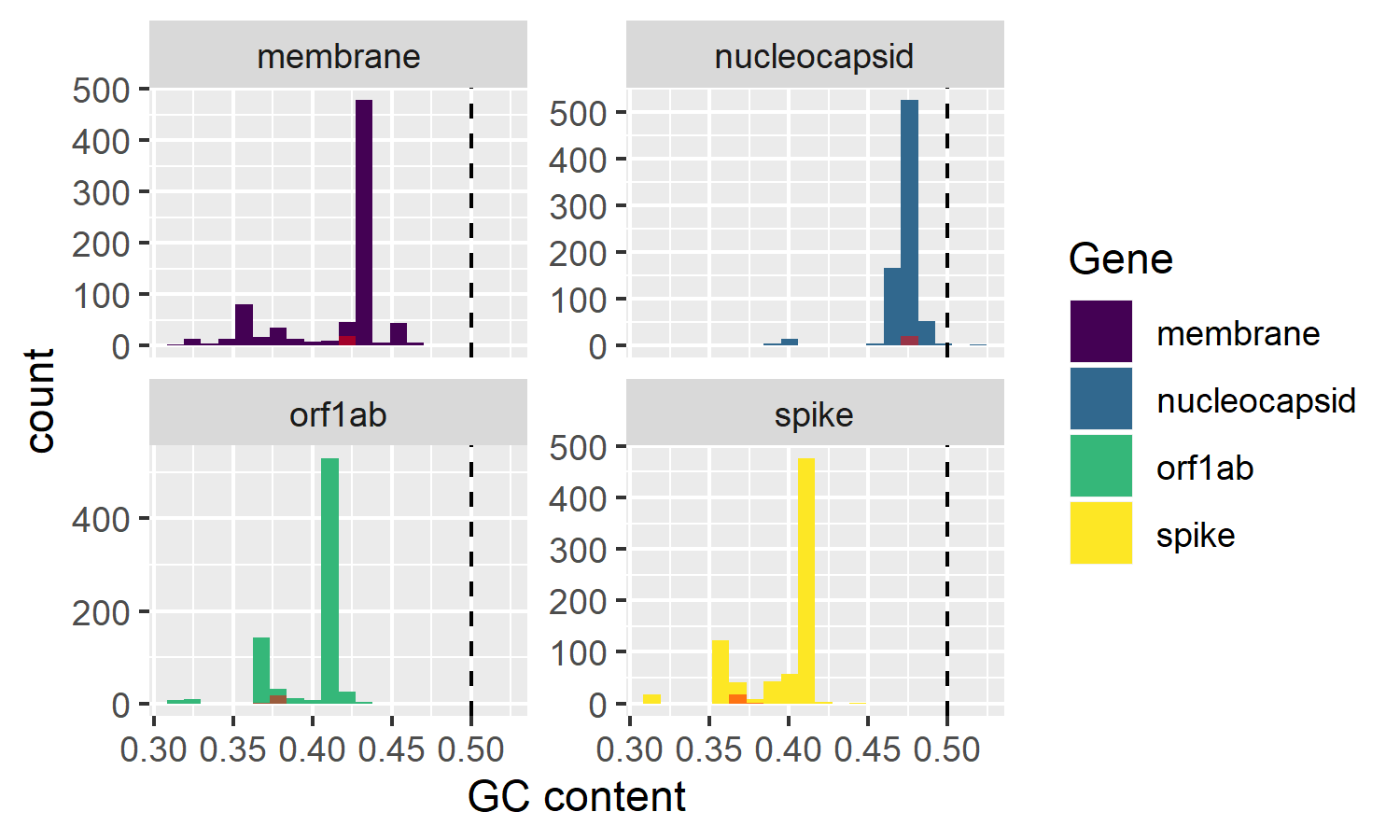

Figure S3. Distribution of the GRAVY (Grand Average of Hydropathy) scores by different genes.

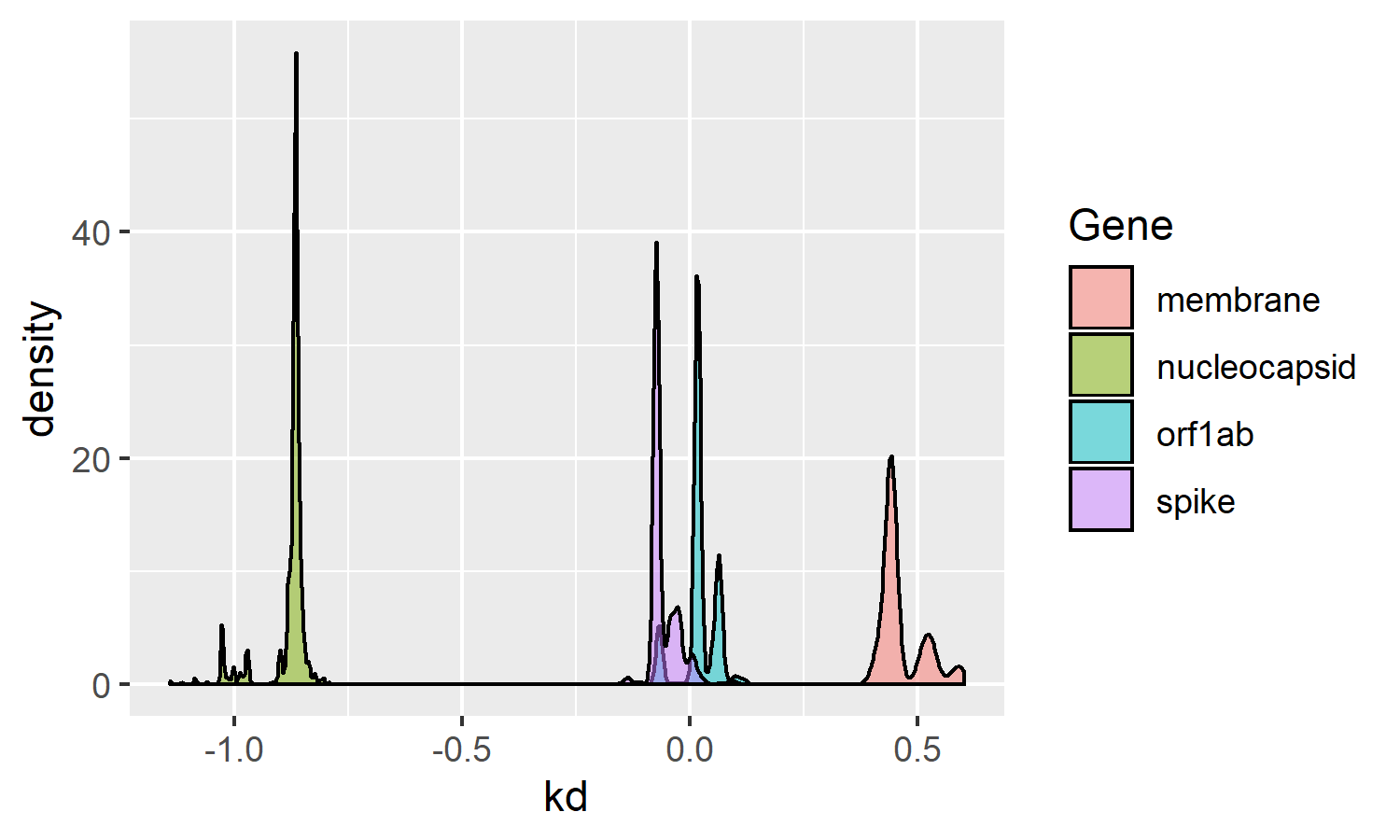

Figure S4. Global correspondence analysis scree plot by different genes

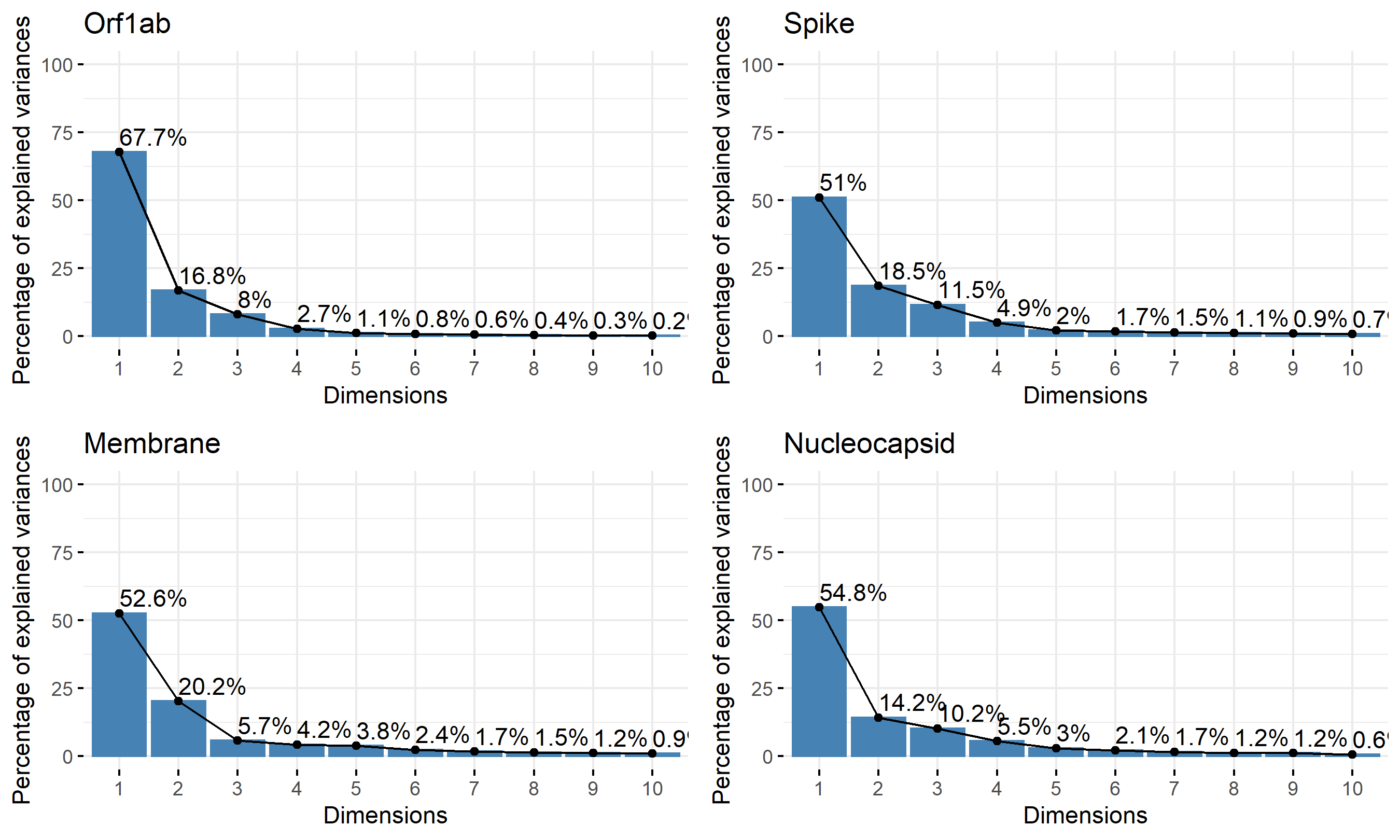

Figure S5. Factorial map of the first and second factors for global CA by different genes, coloured by different viral species, separated by (A) human and pangolin host and (B) bat host.

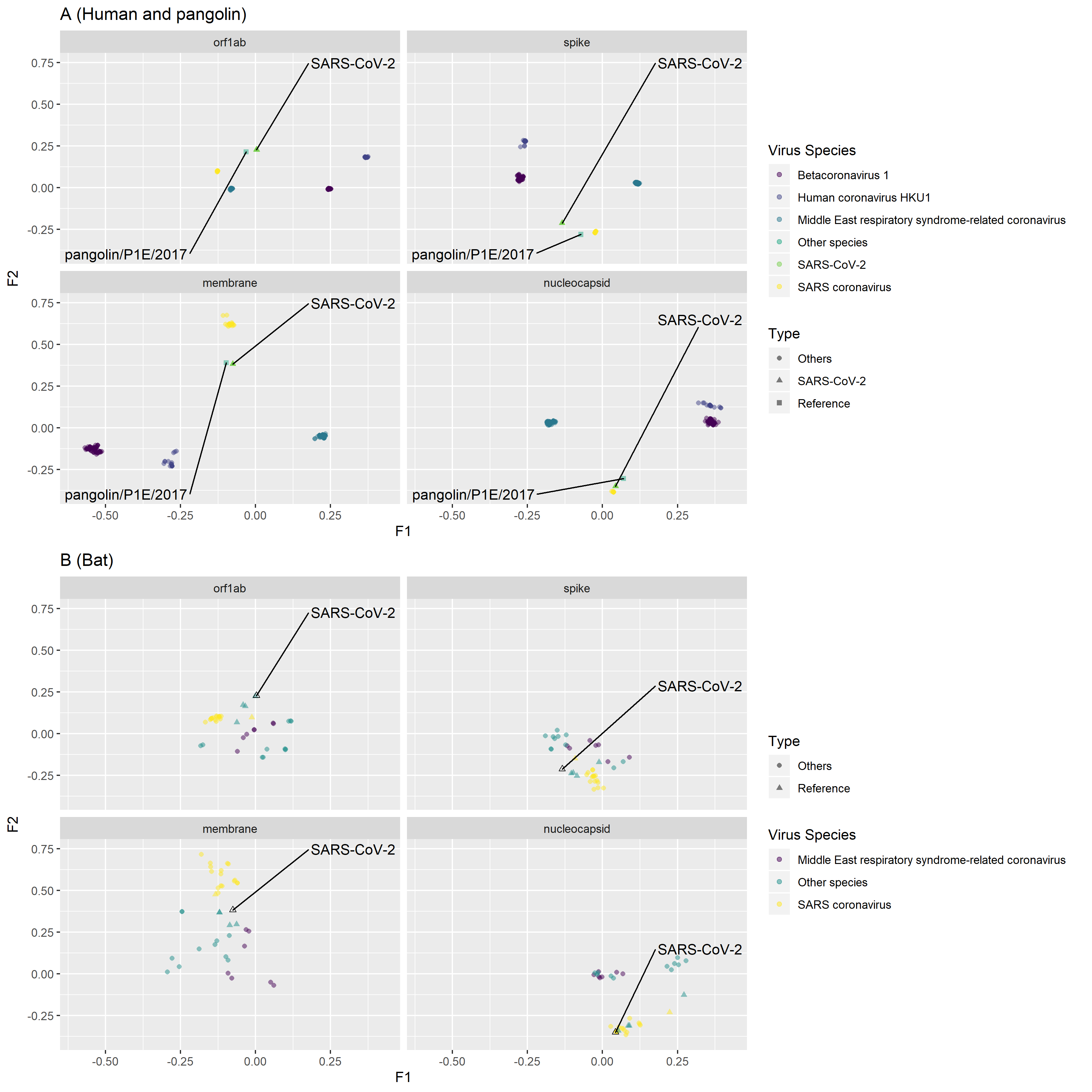

Figure S6. Factorial map of the first and second factors for global CA by different genes, coloured by different virus species, separated by hosts of (A) camel, (B) rodent, (C) swine and (D) others.

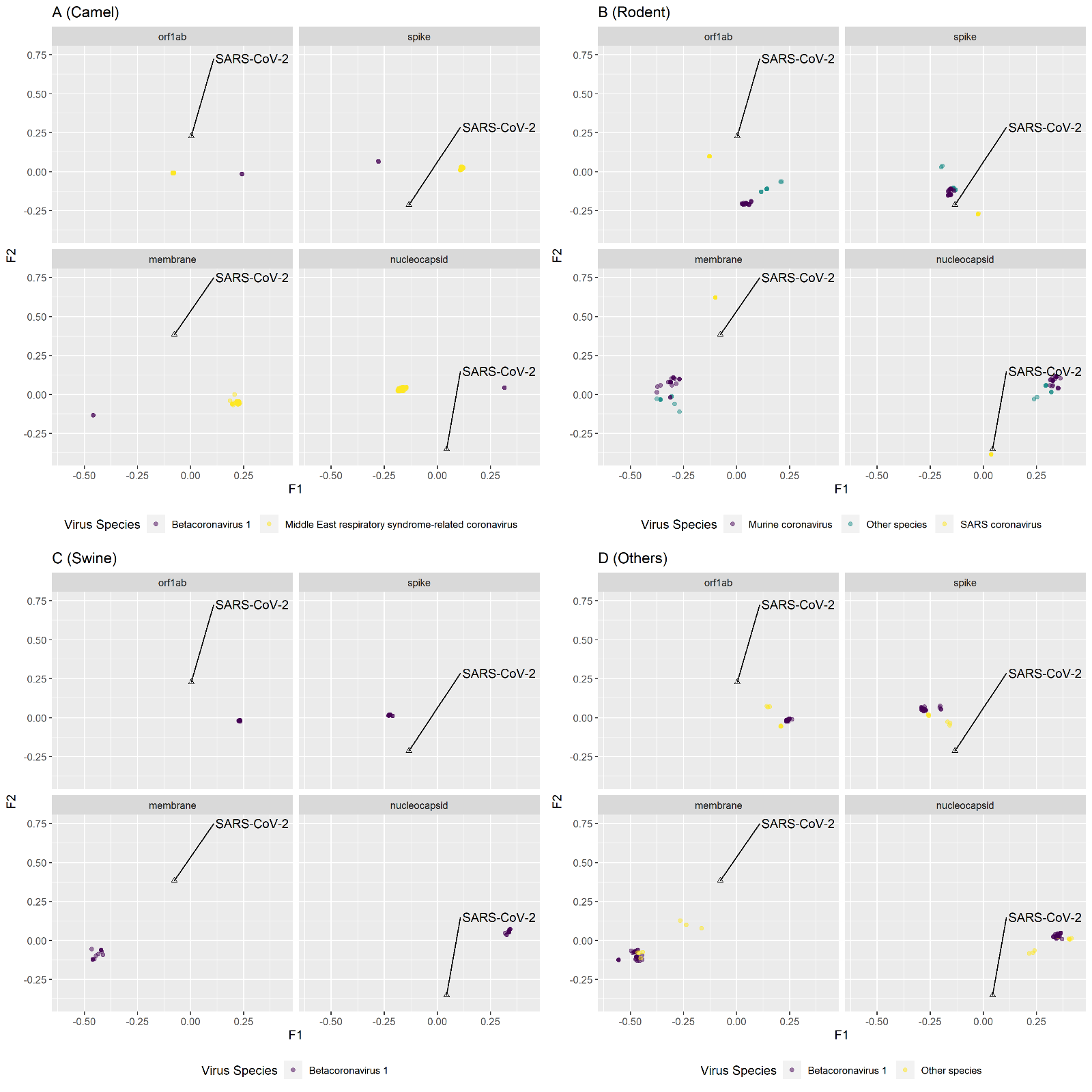

Figure S7. Phylogenetic tree of SARS-CoV-2 and related strains. The tree was built by IQ-TREE ^2^, with Best-fit model: GTR+F+R3 chosen according to AIC. The alignment was performed by MAFFT^3^.

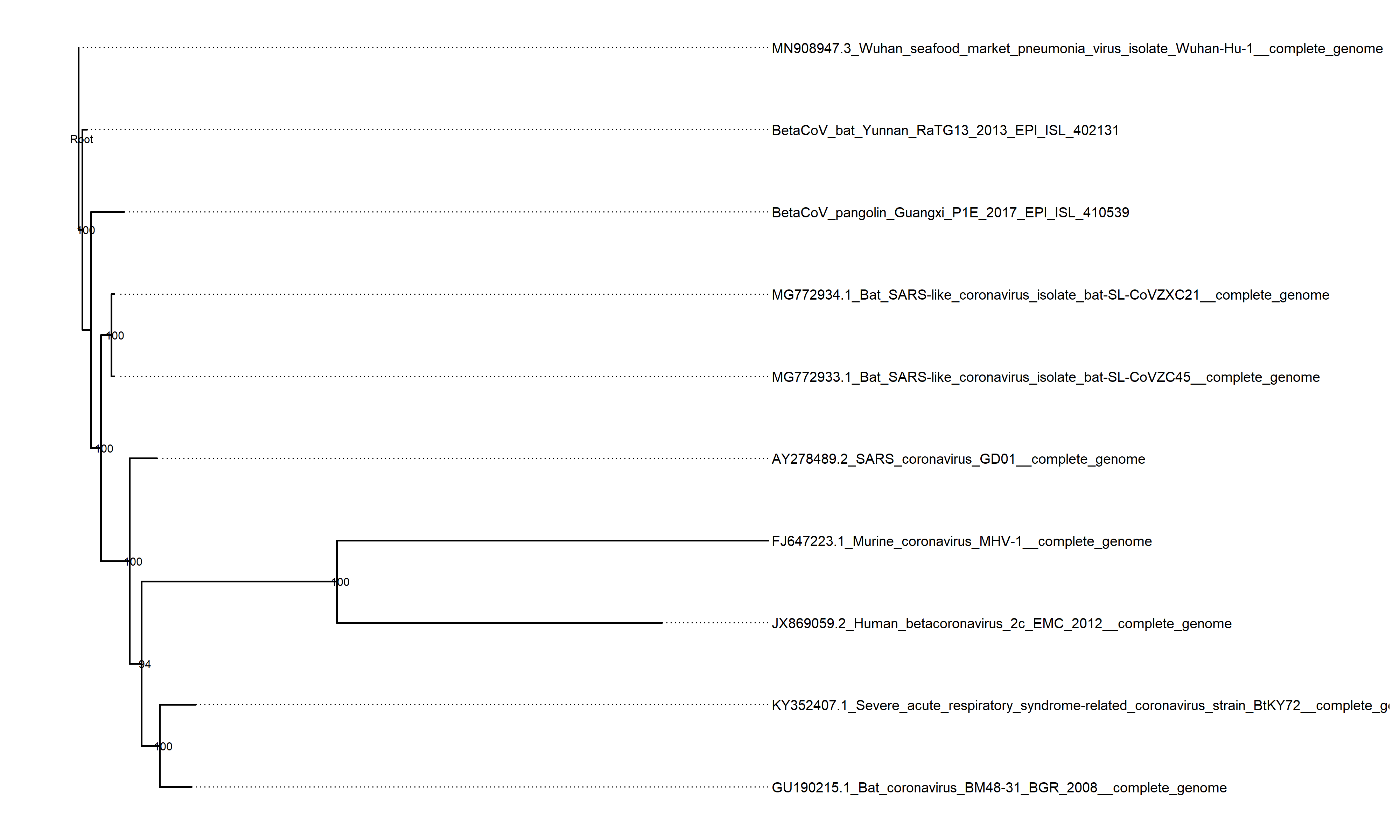

Figure S8. Factorial map of the first, second and third factors for WCA analyses. (The data points of SARS-CoV-2 were pointed by blue arrows)

A. spike gene (interactive online version available at https://koohoko.github.io/CoV_codon_usage/plot_wca_spike.html)

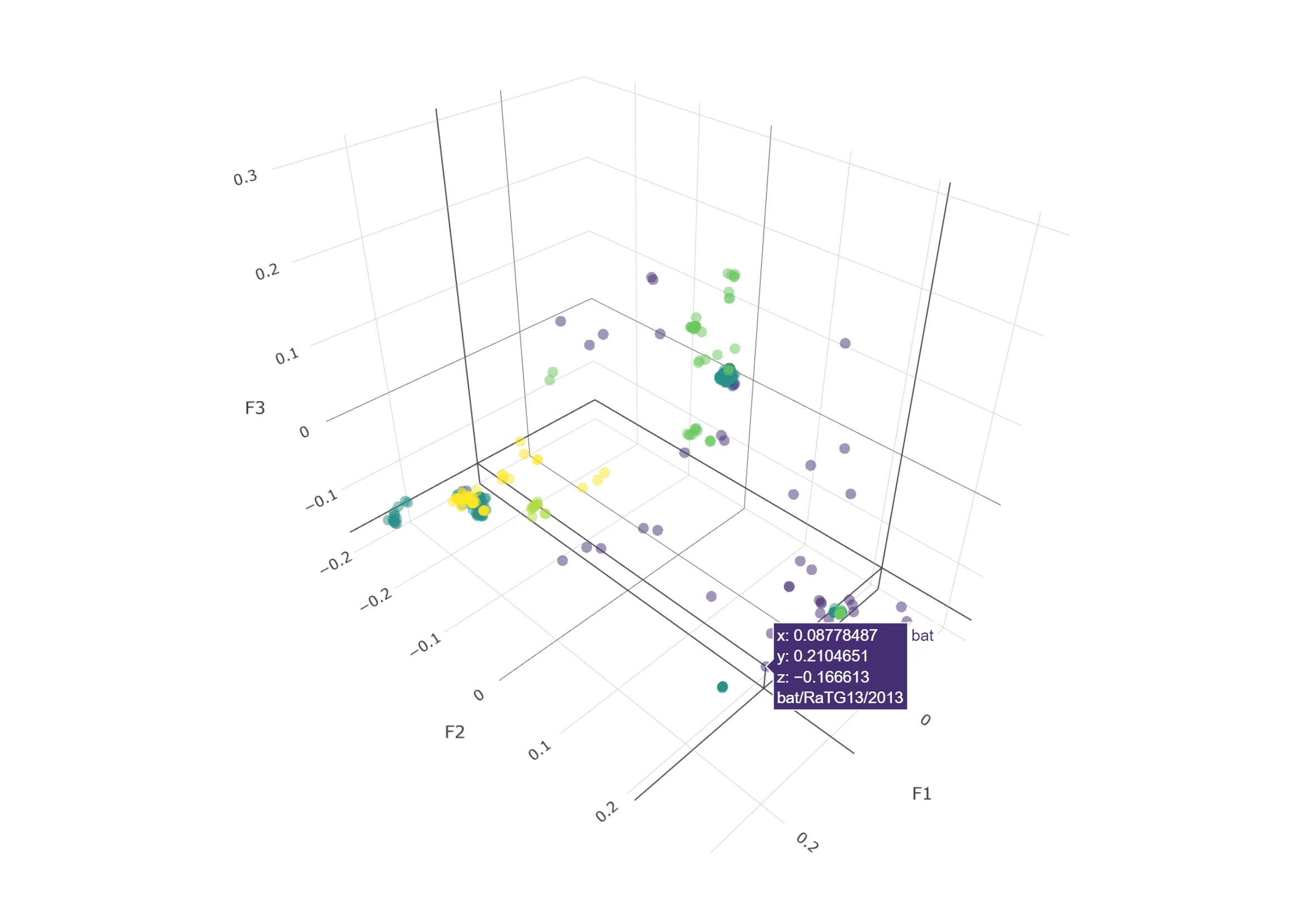

B. membrane gene (interactive online version available at https://koohoko.github.io/CoV_codon_usage/plot_wca_membrane.html)

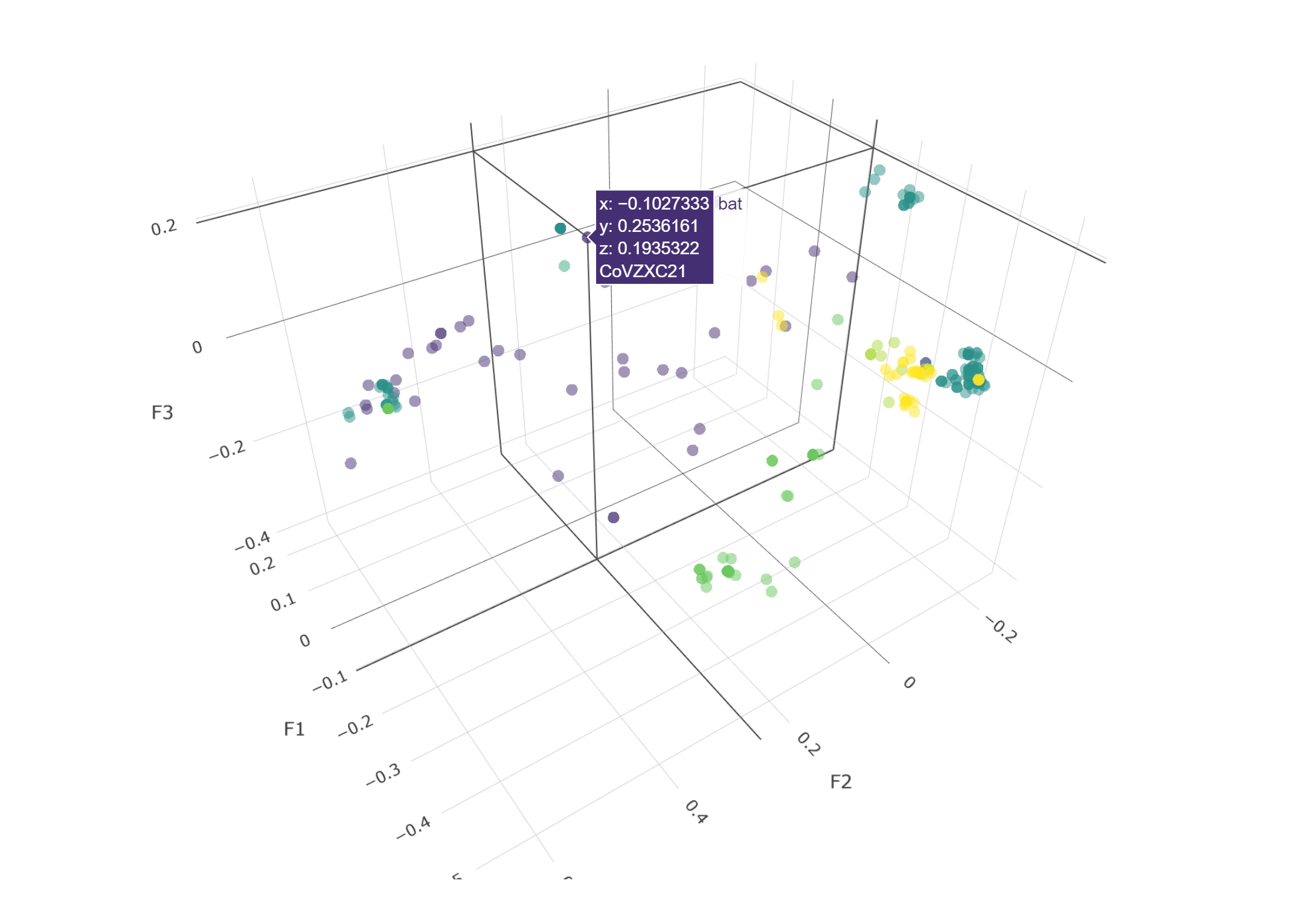

Figure S9. Distribution of codons on the first factors of WCA. Different color indicated different nucleotide on the third base of the codons.

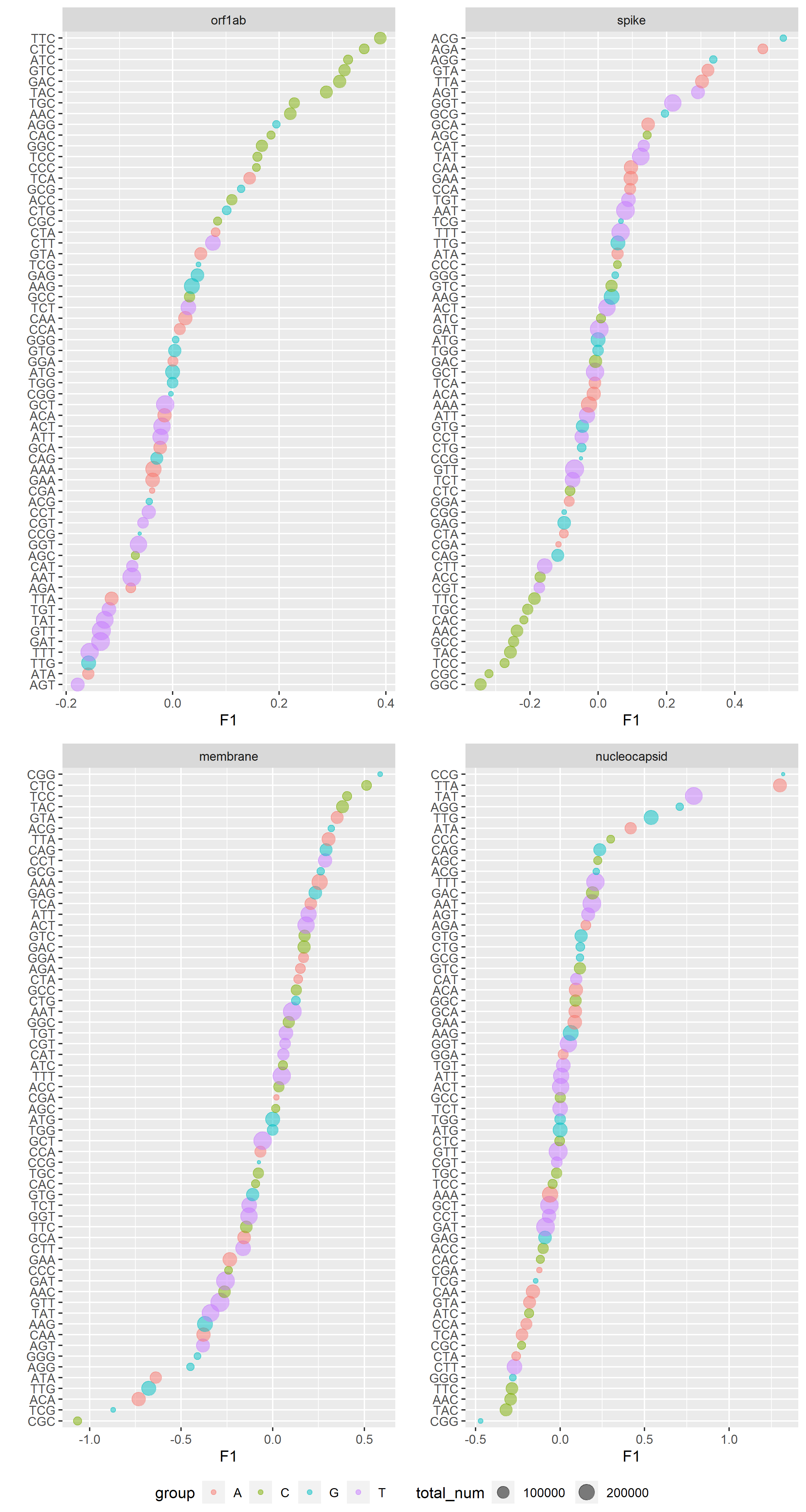

Figure S10. Scree plot of eigen values of all correspondence analyses in this study.

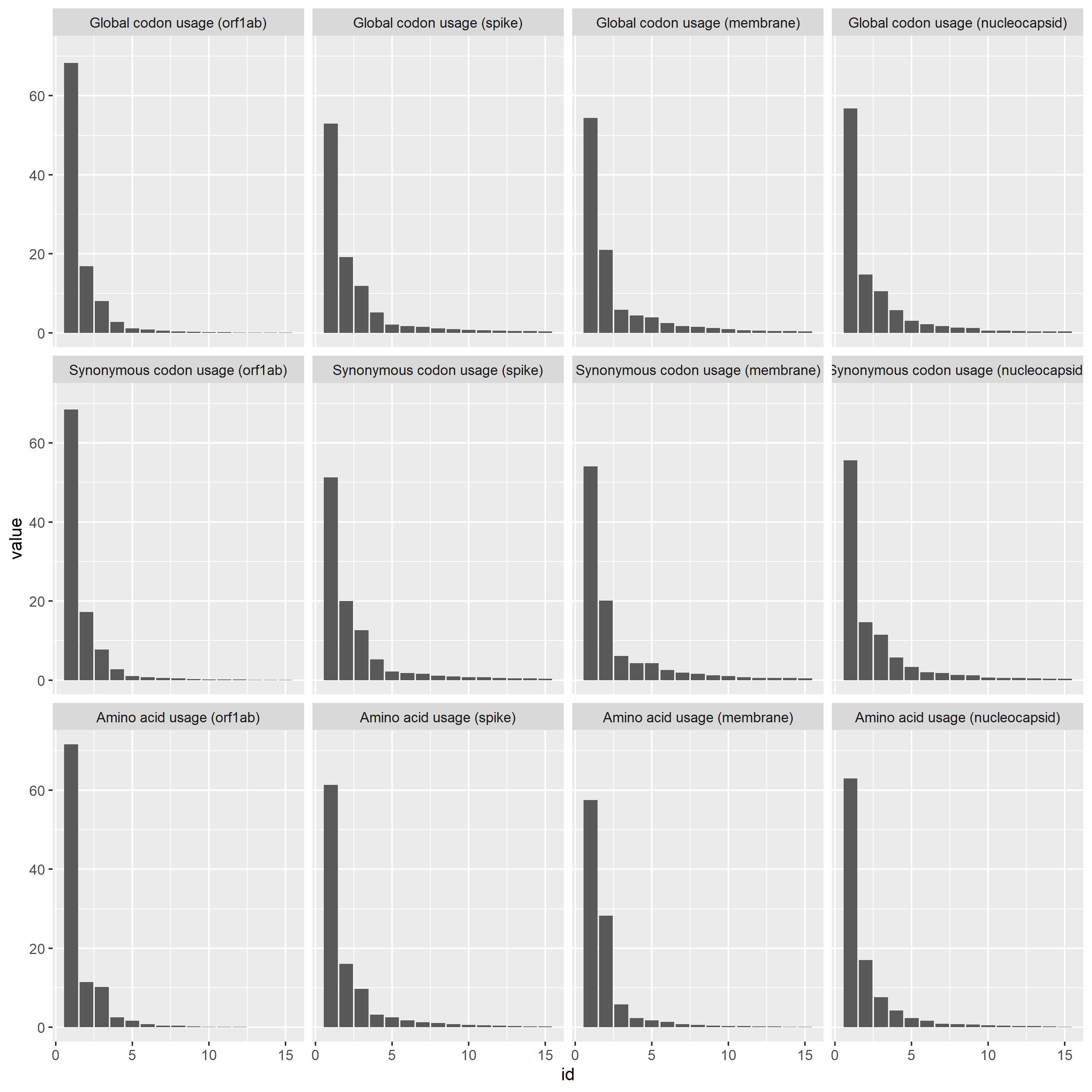

Table S1. Accession ID for the sequences including in this study

id	range	organism	strain_name	Gene

MH687968	NA	Betacoronavirus sp.	VZ_BetaCoV_16715_52	membrane

MH687968	NA	Betacoronavirus sp.	VZ_BetaCoV_16715_52	nucleocapsid

MH687968	NA	Betacoronavirus sp.	VZ_BetaCoV_16715_52	orf1ab

MH687968	NA	Betacoronavirus sp.	VZ_BetaCoV_16715_52	spike

MH687969	NA	Betacoronavirus sp.	VZ_BetaCoV_20724_33	membrane

MH687969	NA	Betacoronavirus sp.	VZ_BetaCoV_20724_33	nucleocapsid

MH687969	NA	Betacoronavirus sp.	VZ_BetaCoV_20724_33	orf1ab

MH687969	NA	Betacoronavirus sp.	VZ_BetaCoV_20724_33	spike

MH687970	NA	Betacoronavirus sp.	VZ_BetaCoV_20724_34_c12	membrane

MH687970	NA	Betacoronavirus sp.	VZ_BetaCoV_20724_34_c12	nucleocapsid

MH687970	NA	Betacoronavirus sp.	VZ_BetaCoV_20724_34_c12	orf1ab

MH687970	NA	Betacoronavirus sp.	VZ_BetaCoV_20724_34_c12	spike

MH687971	NA	Betacoronavirus sp.	VZ_BetaCoV_20724_34_c13	membrane

MH687971	NA	Betacoronavirus sp.	VZ_BetaCoV_20724_34_c13	nucleocapsid

MH687971	NA	Betacoronavirus sp.	VZ_BetaCoV_20724_34_c13	orf1ab

MH687971	NA	Betacoronavirus sp.	VZ_BetaCoV_20724_34_c13	spike

MH687972	NA	Betacoronavirus sp.	VZ_BetaCoV_20724_38	membrane

MH687972	NA	Betacoronavirus sp.	VZ_BetaCoV_20724_38	nucleocapsid

MH687972	NA	Betacoronavirus sp.	VZ_BetaCoV_20724_38	orf1ab

MH687972	NA	Betacoronavirus sp.	VZ_BetaCoV_20724_38	spike

MH687973	NA	Betacoronavirus sp.	VZ_BetaCoV_20724_39	membrane

MH687973	NA	Betacoronavirus sp.	VZ_BetaCoV_20724_39	nucleocapsid

MH687973	NA	Betacoronavirus sp.	VZ_BetaCoV_20724_39	orf1ab

MH687973	NA	Betacoronavirus sp.	VZ_BetaCoV_20724_39	spike

MH687974	NA	Betacoronavirus sp.	VZ_BetaCoV_20724_43	membrane

MH687974	NA	Betacoronavirus sp.	VZ_BetaCoV_20724_43	nucleocapsid

MH687974	NA	Betacoronavirus sp.	VZ_BetaCoV_20724_43	orf1ab

MH687974	NA	Betacoronavirus sp.	VZ_BetaCoV_20724_43	spike

MH687976	NA	Betacoronavirus sp.	VZ_BetaCoV_22084_1	membrane

MH687976	NA	Betacoronavirus sp.	VZ_BetaCoV_22084_1	nucleocapsid

MH687976	NA	Betacoronavirus sp.	VZ_BetaCoV_22084_1	orf1ab

MH687976	NA	Betacoronavirus sp.	VZ_BetaCoV_22084_1	spike

MH687977	NA	Betacoronavirus sp.	VZ_BetaCoV_22084_10	membrane

MH687977	NA	Betacoronavirus sp.	VZ_BetaCoV_22084_10	nucleocapsid

MH687977	NA	Betacoronavirus sp.	VZ_BetaCoV_22084_10	orf1ab

MH687977	NA	Betacoronavirus sp.	VZ_BetaCoV_22084_10	spike

MH687978	NA	Betacoronavirus sp.	VZ_BetaCoV_22084_6	membrane

MH687978	NA	Betacoronavirus sp.	VZ_BetaCoV_22084_6	nucleocapsid

MH687978	NA	Betacoronavirus sp.	VZ_BetaCoV_22084_6	orf1ab

MH687978	NA	Betacoronavirus sp.	VZ_BetaCoV_22084_6	spike

NC_039207	27842-28498	Betacoronavirus Erinaceus/VMC/DEU/2012	ErinaceusCoV/2012-174/GER/2012	membrane

NC_039207	28552-29826	Betacoronavirus Erinaceus/VMC/DEU/2012	ErinaceusCoV/2012-174/GER/2012	nucleocapsid

NC_039207	21610-25602	Betacoronavirus Erinaceus/VMC/DEU/2012	ErinaceusCoV/2012-174/GER/2012	spike

NC_039207	241-21692	Betacoronavirus Erinaceus/VMC/DEU/2012	ErinaceusCoV/2012-174/GER/2012	orf1ab

MK679660	27863-28519	Hedgehog coronavirus 1	UNKNOWN-MK679660	membrane

MK679660	28573-29850	Hedgehog coronavirus 1	UNKNOWN-MK679660	nucleocapsid

MK679660	21628-25614	Hedgehog coronavirus 1	UNKNOWN-MK679660	spike

MK679660	241-21710	Hedgehog coronavirus 1	UNKNOWN-MK679660	orf1ab

NC_038294	27852-28511	Betacoronavirus England 1	England 1	membrane

NC_038294	28565-29800	Betacoronavirus England 1	England 1	nucleocapsid

NC_038294	278-21513	Betacoronavirus England 1	England 1	orf1ab

NC_038294	21455-25516	Betacoronavirus England 1	England 1	spike

KJ473821	NA	BtVs-BetaCoV/SC2013	UNKNOWN-KJ473821	orf1ab

KJ473821	NA	BtVs-BetaCoV/SC2013	UNKNOWN-KJ473821	membrane

KJ473821	NA	BtVs-BetaCoV/SC2013	UNKNOWN-KJ473821	nucleocapsid

KJ473821	NA	BtVs-BetaCoV/SC2013	UNKNOWN-KJ473821	spike

MK357908	NA	Middle East respiratory syndrome-related coronavirus	011/DAB/C8/F<1	membrane

MK357908	NA	Middle East respiratory syndrome-related coronavirus	011/DAB/C8/F<1	nucleocapsid

MK357908	NA	Middle East respiratory syndrome-related coronavirus	011/DAB/C8/F<1	orf1ab

MK357908	NA	Middle East respiratory syndrome-related coronavirus	011/DAB/C8/F<1	spike

MK357909	NA	Middle East respiratory syndrome-related coronavirus	011/LOM/C20/F<1	membrane

MK357909	NA	Middle East respiratory syndrome-related coronavirus	011/LOM/C20/F<1	nucleocapsid

MK357909	NA	Middle East respiratory syndrome-related coronavirus	011/LOM/C20/F<1	orf1ab

MK357909	NA	Middle East respiratory syndrome-related coronavirus	011/LOM/C20/F<1	spike

MH371127	279-21514	Middle East respiratory syndrome-related coronavirus	2362	orf1ab

MH371127	27853-28512	Middle East respiratory syndrome-related coronavirus	2362	membrane

MH371127	28566-29807	Middle East respiratory syndrome-related coronavirus	2362	nucleocapsid

MH371127	21456-25517	Middle East respiratory syndrome-related coronavirus	2362	spike

MH395139	279-21514	Middle East respiratory syndrome-related coronavirus	2363	orf1ab

MH395139	27853-28512	Middle East respiratory syndrome-related coronavirus	2363	membrane

MH395139	28566-29807	Middle East respiratory syndrome-related coronavirus	2363	nucleocapsid

MH395139	21456-25517	Middle East respiratory syndrome-related coronavirus	2363	spike

MH432120	279-21514	Middle East respiratory syndrome-related coronavirus	2366	orf1ab

MH432120	27853-28512	Middle East respiratory syndrome-related coronavirus	2366	membrane

MH432120	28566-29807	Middle East respiratory syndrome-related coronavirus	2366	nucleocapsid

MH432120	21456-25517	Middle East respiratory syndrome-related coronavirus	2366	spike

MG596802	217-21446	Middle East respiratory syndrome-related coronavirus	Bat-CoV/H.savii/Italy/206645-40/2011	orf1ab

MG596802	27756-28412	Middle East respiratory syndrome-related coronavirus	Bat-CoV/H.savii/Italy/206645-40/2011	membrane

MG596802	28460-29749	Middle East respiratory syndrome-related coronavirus	Bat-CoV/H.savii/Italy/206645-40/2011	nucleocapsid

MG596802	21388-25425	Middle East respiratory syndrome-related coronavirus	Bat-CoV/H.savii/Italy/206645-40/2011	spike

MG596803	208-21437	Middle East respiratory syndrome-related coronavirus	Bat-CoV/P.khulii/Italy/206645-63/2011	orf1ab

MG596803	27747-28403	Middle East respiratory syndrome-related coronavirus	Bat-CoV/P.khulii/Italy/206645-63/2011	membrane

MG596803	28451-29740	Middle East respiratory syndrome-related coronavirus	Bat-CoV/P.khulii/Italy/206645-63/2011	nucleocapsid

MG596803	21379-25416	Middle East respiratory syndrome-related coronavirus	Bat-CoV/P.khulii/Italy/206645-63/2011	spike

MK564474	NA	Middle East respiratory syndrome-related coronavirus	camel/MERS/Amibara/118/2017	orf1ab

MK564474	NA	Middle East respiratory syndrome-related coronavirus	camel/MERS/Amibara/118/2017	membrane

MK564474	NA	Middle East respiratory syndrome-related coronavirus	camel/MERS/Amibara/118/2017	nucleocapsid

MK564474	NA	Middle East respiratory syndrome-related coronavirus	camel/MERS/Amibara/118/2017	spike

MK564475	NA	Middle East respiratory syndrome-related coronavirus	camel/MERS/Amibara/126/2017	orf1ab

MK564475	NA	Middle East respiratory syndrome-related coronavirus	camel/MERS/Amibara/126/2017	membrane

MK564475	NA	Middle East respiratory syndrome-related coronavirus	camel/MERS/Amibara/126/2017	nucleocapsid

MK564475	NA	Middle East respiratory syndrome-related coronavirus	camel/MERS/Amibara/126/2017	spike

KY673149	27853-28512	Middle East respiratory syndrome-related coronavirus	Camel/Oman_1_2015	membrane

KY673149	28566-29807	Middle East respiratory syndrome-related coronavirus	Camel/Oman_1_2015	nucleocapsid

KY673149	279-21514	Middle East respiratory syndrome-related coronavirus	Camel/Oman_1_2015	orf1ab

KY673149	21456-25517	Middle East respiratory syndrome-related coronavirus	Camel/Oman_1_2015	spike

KY581695	279-21514	Middle East respiratory syndrome-related coronavirus	Camel/UAE_1B-A_2014	orf1ab

KY581695	27853-28512	Middle East respiratory syndrome-related coronavirus	Camel/UAE_1B-A_2014	membrane

KY581695	28566-29807	Middle East respiratory syndrome-related coronavirus	Camel/UAE_1B-A_2014	nucleocapsid

KY581695	21456-25517	Middle East respiratory syndrome-related coronavirus	Camel/UAE_1B-A_2014	spike

KY581696	279-21514	Middle East respiratory syndrome-related coronavirus	Camel/UAE_1H-B_2014	orf1ab

KY581696	27853-28512	Middle East respiratory syndrome-related coronavirus	Camel/UAE_1H-B_2014	membrane

KY581696	28566-29807	Middle East respiratory syndrome-related coronavirus	Camel/UAE_1H-B_2014	nucleocapsid

KY581696	21456-25517	Middle East respiratory syndrome-related coronavirus	Camel/UAE_1H-B_2014	spike

KY581697	279-21514	Middle East respiratory syndrome-related coronavirus	Camel/UAE_1H-D_2014	orf1ab

KY581697	27853-28512	Middle East respiratory syndrome-related coronavirus	Camel/UAE_1H-D_2014	membrane

KY581697	28566-29807	Middle East respiratory syndrome-related coronavirus	Camel/UAE_1H-D_2014	nucleocapsid

KY581697	21456-25517	Middle East respiratory syndrome-related coronavirus	Camel/UAE_1H-D_2014	spike

MF598702	279-21514	Middle East respiratory syndrome-related coronavirus	camel/UAE_414377_W3_2015	orf1ab

MF598702	27853-28512	Middle East respiratory syndrome-related coronavirus	camel/UAE_414377_W3_2015	membrane

MF598702	28566-29807	Middle East respiratory syndrome-related coronavirus	camel/UAE_414377_W3_2015	nucleocapsid

MF598702	21456-25517	Middle East respiratory syndrome-related coronavirus	camel/UAE_414377_W3_2015	spike

MF598700	279-21514	Middle East respiratory syndrome-related coronavirus	camel/UAE_414379_W3_2015	orf1ab

MF598700	27853-28512	Middle East respiratory syndrome-related coronavirus	camel/UAE_414379_W3_2015	membrane

MF598700	28566-29807	Middle East respiratory syndrome-related coronavirus	camel/UAE_414379_W3_2015	nucleocapsid

MF598700	21456-25517	Middle East respiratory syndrome-related coronavirus	camel/UAE_414379_W3_2015	spike

MF598711	279-21514	Middle East respiratory syndrome-related coronavirus	camel/UAE_414481_W4_2015	orf1ab

MF598711	27853-28512	Middle East respiratory syndrome-related coronavirus	camel/UAE_414481_W4_2015	membrane

MF598711	28566-29807	Middle East respiratory syndrome-related coronavirus	camel/UAE_414481_W4_2015	nucleocapsid

MF598711	21456-25517	Middle East respiratory syndrome-related coronavirus	camel/UAE_414481_W4_2015	spike

MF598699	279-21514	Middle East respiratory syndrome-related coronavirus	camel/UAE_414485_W3_2015	orf1ab

MF598699	27853-28512	Middle East respiratory syndrome-related coronavirus	camel/UAE_414485_W3_2015	membrane

MF598699	28566-29807	Middle East respiratory syndrome-related coronavirus	camel/UAE_414485_W3_2015	nucleocapsid

MF598699	21456-25517	Middle East respiratory syndrome-related coronavirus	camel/UAE_414485_W3_2015	spike

MF598709	279-21514	Middle East respiratory syndrome-related coronavirus	camel/UAE_414485_W4_2015	orf1ab

MF598709	27853-28512	Middle East respiratory syndrome-related coronavirus	camel/UAE_414485_W4_2015	membrane

MF598709	28566-29807	Middle East respiratory syndrome-related coronavirus	camel/UAE_414485_W4_2015	nucleocapsid

MF598709	21456-25517	Middle East respiratory syndrome-related coronavirus	camel/UAE_414485_W4_2015	spike

MF598705	279-21514	Middle East respiratory syndrome-related coronavirus	camel/UAE_414486_W3_2015	orf1ab

MF598705	27853-28512	Middle East respiratory syndrome-related coronavirus	camel/UAE_414486_W3_2015	membrane

MF598705	28566-29807	Middle East respiratory syndrome-related coronavirus	camel/UAE_414486_W3_2015	nucleocapsid

MF598705	21456-25517	Middle East respiratory syndrome-related coronavirus	camel/UAE_414486_W3_2015	spike

MF598716	279-21514	Middle East respiratory syndrome-related coronavirus	camel/UAE_414486_W4_2015	orf1ab

MF598716	27853-28512	Middle East respiratory syndrome-related coronavirus	camel/UAE_414486_W4_2015	membrane

MF598716	28566-29807	Middle East respiratory syndrome-related coronavirus	camel/UAE_414486_W4_2015	nucleocapsid

MF598716	21456-25517	Middle East respiratory syndrome-related coronavirus	camel/UAE_414486_W4_2015	spike

MF598712	279-21514	Middle East respiratory syndrome-related coronavirus	camel/UAE_414492_W4_2015	orf1ab

MF598712	27853-28512	Middle East respiratory syndrome-related coronavirus	camel/UAE_414492_W4_2015	membrane

MF598712	28566-29807	Middle East respiratory syndrome-related coronavirus	camel/UAE_414492_W4_2015	nucleocapsid

MF598712	21456-25517	Middle East respiratory syndrome-related coronavirus	camel/UAE_414492_W4_2015	spike

MF598701	279-21514	Middle East respiratory syndrome-related coronavirus	camel/UAE_414498_W3_2015	orf1ab

MF598701	27853-28512	Middle East respiratory syndrome-related coronavirus	camel/UAE_414498_W3_2015	membrane

MF598701	28566-29807	Middle East respiratory syndrome-related coronavirus	camel/UAE_414498_W3_2015	nucleocapsid

MF598701	21456-25517	Middle East respiratory syndrome-related coronavirus	camel/UAE_414498_W3_2015	spike

MF598714	279-21514	Middle East respiratory syndrome-related coronavirus	camel/UAE_414500_W4_2015	orf1ab

MF598714	27853-28512	Middle East respiratory syndrome-related coronavirus	camel/UAE_414500_W4_2015	membrane

MF598714	28566-29807	Middle East respiratory syndrome-related coronavirus	camel/UAE_414500_W4_2015	nucleocapsid

MF598714	21456-25517	Middle East respiratory syndrome-related coronavirus	camel/UAE_414500_W4_2015	spike

MF598706	279-21514	Middle East respiratory syndrome-related coronavirus	camel/UAE_415911_W3_2015	orf1ab

MF598706	27853-28512	Middle East respiratory syndrome-related coronavirus	camel/UAE_415911_W3_2015	membrane

MF598706	28566-29807	Middle East respiratory syndrome-related coronavirus	camel/UAE_415911_W3_2015	nucleocapsid

MF598706	21456-25517	Middle East respiratory syndrome-related coronavirus	camel/UAE_415911_W3_2015	spike

MF598719	279-21514	Middle East respiratory syndrome-related coronavirus	camel/UAE_415911_W4_2015	orf1ab

MF598719	27853-28512	Middle East respiratory syndrome-related coronavirus	camel/UAE_415911_W4_2015	membrane

MF598719	28566-29807	Middle East respiratory syndrome-related coronavirus	camel/UAE_415911_W4_2015	nucleocapsid

MF598719	21456-25517	Middle East respiratory syndrome-related coronavirus	camel/UAE_415911_W4_2015	spike

MF598707	279-21514	Middle East respiratory syndrome-related coronavirus	camel/UAE_415915_W3_2015	orf1ab

MF598707	27853-28512	Middle East respiratory syndrome-related coronavirus	camel/UAE_415915_W3_2015	membrane

MF598707	28566-29807	Middle East respiratory syndrome-related coronavirus	camel/UAE_415915_W3_2015	nucleocapsid

MF598707	21456-25517	Middle East respiratory syndrome-related coronavirus	camel/UAE_415915_W3_2015	spike

MF598722	279-21514	Middle East respiratory syndrome-related coronavirus	camel/UAE_415915_W6_2015	orf1ab

MF598722	27853-28512	Middle East respiratory syndrome-related coronavirus	camel/UAE_415915_W6_2015	membrane

MF598722	28566-29807	Middle East respiratory syndrome-related coronavirus	camel/UAE_415915_W6_2015	nucleocapsid

MF598722	21456-25517	Middle East respiratory syndrome-related coronavirus	camel/UAE_415915_W6_2015	spike

MF598715	279-21514	Middle East respiratory syndrome-related coronavirus	camel/UAE_416452_W4_2015	orf1ab

MF598715	27853-28512	Middle East respiratory syndrome-related coronavirus	camel/UAE_416452_W4_2015	membrane

MF598715	28566-29807	Middle East respiratory syndrome-related coronavirus	camel/UAE_416452_W4_2015	nucleocapsid

MF598715	21456-25517	Middle East respiratory syndrome-related coronavirus	camel/UAE_416452_W4_2015	spike

MF598721	279-21514	Middle East respiratory syndrome-related coronavirus	camel/UAE_416452_W6_2015	orf1ab

MF598721	27853-28512	Middle East respiratory syndrome-related coronavirus	camel/UAE_416452_W6_2015	membrane

MF598721	28566-29807	Middle East respiratory syndrome-related coronavirus	camel/UAE_416452_W6_2015	nucleocapsid

MF598721	21456-25517	Middle East respiratory syndrome-related coronavirus	camel/UAE_416452_W6_2015	spike

MF598710	279-21514	Middle East respiratory syndrome-related coronavirus	camel/UAE_417162_W4_2015	orf1ab

MF598710	27853-28512	Middle East respiratory syndrome-related coronavirus	camel/UAE_417162_W4_2015	membrane

MF598710	28566-29807	Middle East respiratory syndrome-related coronavirus	camel/UAE_417162_W4_2015	nucleocapsid

MF598710	21456-25517	Middle East respiratory syndrome-related coronavirus	camel/UAE_417162_W4_2015	spike

MF598717	279-21514	Middle East respiratory syndrome-related coronavirus	camel/UAE_417163_W4_2015	orf1ab

MF598717	27853-28512	Middle East respiratory syndrome-related coronavirus	camel/UAE_417163_W4_2015	membrane

MF598717	28566-29807	Middle East respiratory syndrome-related coronavirus	camel/UAE_417163_W4_2015	nucleocapsid

MF598717	21456-25517	Middle East respiratory syndrome-related coronavirus	camel/UAE_417163_W4_2015	spike

MF598687	279-21514	Middle East respiratory syndrome-related coronavirus	camel/UAE_B100_2015	orf1ab

MF598687	27853-28512	Middle East respiratory syndrome-related coronavirus	camel/UAE_B100_2015	membrane

MF598687	28566-29807	Middle East respiratory syndrome-related coronavirus	camel/UAE_B100_2015	nucleocapsid

MF598687	21456-25517	Middle East respiratory syndrome-related coronavirus	camel/UAE_B100_2015	spike

MF598688	279-21514	Middle East respiratory syndrome-related coronavirus	camel/UAE_B101_2015	orf1ab

MF598688	27853-28512	Middle East respiratory syndrome-related coronavirus	camel/UAE_B101_2015	membrane

MF598688	28566-29807	Middle East respiratory syndrome-related coronavirus	camel/UAE_B101_2015	nucleocapsid

MF598688	21456-25517	Middle East respiratory syndrome-related coronavirus	camel/UAE_B101_2015	spike

MF598689	277-21512	Middle East respiratory syndrome-related coronavirus	camel/UAE_B102_2015	orf1ab

MF598689	27851-28510	Middle East respiratory syndrome-related coronavirus	camel/UAE_B102_2015	membrane

MF598689	28564-29805	Middle East respiratory syndrome-related coronavirus	camel/UAE_B102_2015	nucleocapsid

MF598689	21454-25515	Middle East respiratory syndrome-related coronavirus	camel/UAE_B102_2015	spike

MF598690	279-21514	Middle East respiratory syndrome-related coronavirus	camel/UAE_B103_2015	orf1ab

MF598690	27853-28512	Middle East respiratory syndrome-related coronavirus	camel/UAE_B103_2015	membrane

MF598690	28566-29807	Middle East respiratory syndrome-related coronavirus	camel/UAE_B103_2015	nucleocapsid

MF598690	21456-25517	Middle East respiratory syndrome-related coronavirus	camel/UAE_B103_2015	spike

MF598691	279-21514	Middle East respiratory syndrome-related coronavirus	camel/UAE_B104_2015	orf1ab

MF598691	27853-28512	Middle East respiratory syndrome-related coronavirus	camel/UAE_B104_2015	membrane

MF598691	28566-29807	Middle East respiratory syndrome-related coronavirus	camel/UAE_B104_2015	nucleocapsid

MF598691	21456-25517	Middle East respiratory syndrome-related coronavirus	camel/UAE_B104_2015	spike

MF598693	279-21514	Middle East respiratory syndrome-related coronavirus	camel/UAE_B106_2015	orf1ab

MF598693	27853-28512	Middle East respiratory syndrome-related coronavirus	camel/UAE_B106_2015	membrane

MF598693	28566-29807	Middle East respiratory syndrome-related coronavirus	camel/UAE_B106_2015	nucleocapsid

MF598693	21456-25517	Middle East respiratory syndrome-related coronavirus	camel/UAE_B106_2015	spike

MF598694	279-21514	Middle East respiratory syndrome-related coronavirus	camel/UAE_B107_2015	orf1ab

MF598694	27853-28512	Middle East respiratory syndrome-related coronavirus	camel/UAE_B107_2015	membrane

MF598694	28566-29807	Middle East respiratory syndrome-related coronavirus	camel/UAE_B107_2015	nucleocapsid

MF598694	21456-25517	Middle East respiratory syndrome-related coronavirus	camel/UAE_B107_2015	spike

MF598695	279-21514	Middle East respiratory syndrome-related coronavirus	camel/UAE_B108_2015	orf1ab

MF598695	27853-28512	Middle East respiratory syndrome-related coronavirus	camel/UAE_B108_2015	membrane

MF598695	28566-29807	Middle East respiratory syndrome-related coronavirus	camel/UAE_B108_2015	nucleocapsid

MF598695	21456-25517	Middle East respiratory syndrome-related coronavirus	camel/UAE_B108_2015	spike

MF598696	279-21514	Middle East respiratory syndrome-related coronavirus	camel/UAE_B109_2015	orf1ab

MF598696	27853-28512	Middle East respiratory syndrome-related coronavirus	camel/UAE_B109_2015	membrane

MF598696	28566-29807	Middle East respiratory syndrome-related coronavirus	camel/UAE_B109_2015	nucleocapsid

MF598696	21456-25517	Middle East respiratory syndrome-related coronavirus	camel/UAE_B109_2015	spike

MF598602	279-21514	Middle East respiratory syndrome-related coronavirus	camel/UAE_B10_2015	orf1ab

MF598602	27853-28512	Middle East respiratory syndrome-related coronavirus	camel/UAE_B10_2015	membrane

MF598602	28566-29807	Middle East respiratory syndrome-related coronavirus	camel/UAE_B10_2015	nucleocapsid

MF598602	21456-25517	Middle East respiratory syndrome-related coronavirus	camel/UAE_B10_2015	spike

MF598697	279-21514	Middle East respiratory syndrome-related coronavirus	camel/UAE_B110_2015	orf1ab

MF598697	27853-28512	Middle East respiratory syndrome-related coronavirus	camel/UAE_B110_2015	membrane

MF598697	28566-29807	Middle East respiratory syndrome-related coronavirus	camel/UAE_B110_2015	nucleocapsid

MF598697	21456-25517	Middle East respiratory syndrome-related coronavirus	camel/UAE_B110_2015	spike

MF598698	279-21514	Middle East respiratory syndrome-related coronavirus	camel/UAE_B111_2015	orf1ab

MF598698	27853-28512	Middle East respiratory syndrome-related coronavirus	camel/UAE_B111_2015	membrane

MF598698	28566-29807	Middle East respiratory syndrome-related coronavirus	camel/UAE_B111_2015	nucleocapsid

MF598698	21456-25517	Middle East respiratory syndrome-related coronavirus	camel/UAE_B111_2015	spike

MF598603	279-21514	Middle East respiratory syndrome-related coronavirus	camel/UAE_B11_2015	orf1ab

MF598603	27853-28512	Middle East respiratory syndrome-related coronavirus	camel/UAE_B11_2015	membrane

MF598603	28566-29807	Middle East respiratory syndrome-related coronavirus	camel/UAE_B11_2015	nucleocapsid

MF598603	21456-25517	Middle East respiratory syndrome-related coronavirus	camel/UAE_B11_2015	spike

MF598604	279-21514	Middle East respiratory syndrome-related coronavirus	camel/UAE_B12_2015	orf1ab

MF598604	27853-28512	Middle East respiratory syndrome-related coronavirus	camel/UAE_B12_2015	membrane

MF598604	28566-29807	Middle East respiratory syndrome-related coronavirus	camel/UAE_B12_2015	nucleocapsid

MF598604	21456-25517	Middle East respiratory syndrome-related coronavirus	camel/UAE_B12_2015	spike

MF598606	279-21514	Middle East respiratory syndrome-related coronavirus	camel/UAE_B14_2015	orf1ab

MF598606	27853-28512	Middle East respiratory syndrome-related coronavirus	camel/UAE_B14_2015	membrane

MF598606	28566-29807	Middle East respiratory syndrome-related coronavirus	camel/UAE_B14_2015	nucleocapsid

MF598606	21456-25517	Middle East respiratory syndrome-related coronavirus	camel/UAE_B14_2015	spike

MF598607	279-21514	Middle East respiratory syndrome-related coronavirus	camel/UAE_B15_2015	orf1ab

MF598607	27853-28512	Middle East respiratory syndrome-related coronavirus	camel/UAE_B15_2015	membrane

MF598607	28566-29807	Middle East respiratory syndrome-related coronavirus	camel/UAE_B15_2015	nucleocapsid

MF598607	21456-25517	Middle East respiratory syndrome-related coronavirus	camel/UAE_B15_2015	spike

MF598608	279-21514	Middle East respiratory syndrome-related coronavirus	camel/UAE_B16_2015	orf1ab

MF598608	27853-28512	Middle East respiratory syndrome-related coronavirus	camel/UAE_B16_2015	membrane

MF598608	28566-29807	Middle East respiratory syndrome-related coronavirus	camel/UAE_B16_2015	nucleocapsid

MF598608	21456-25517	Middle East respiratory syndrome-related coronavirus	camel/UAE_B16_2015	spike

MF598609	279-21514	Middle East respiratory syndrome-related coronavirus	camel/UAE_B17_2015	orf1ab

MF598609	27853-28512	Middle East respiratory syndrome-related coronavirus	camel/UAE_B17_2015	membrane

MF598609	28566-29807	Middle East respiratory syndrome-related coronavirus	camel/UAE_B17_2015	nucleocapsid

MF598609	21456-25517	Middle East respiratory syndrome-related coronavirus	camel/UAE_B17_2015	spike

MF598610	279-21514	Middle East respiratory syndrome-related coronavirus	camel/UAE_B18_2015	orf1ab

MF598610	27853-28512	Middle East respiratory syndrome-related coronavirus	camel/UAE_B18_2015	membrane

MF598610	28566-29807	Middle East respiratory syndrome-related coronavirus	camel/UAE_B18_2015	nucleocapsid

MF598610	21456-25517	Middle East respiratory syndrome-related coronavirus	camel/UAE_B18_2015	spike

MF598611	279-21514	Middle East respiratory syndrome-related coronavirus	camel/UAE_B19_2015	orf1ab

MF598611	27853-28512	Middle East respiratory syndrome-related coronavirus	camel/UAE_B19_2015	membrane

MF598611	28566-29807	Middle East respiratory syndrome-related coronavirus	camel/UAE_B19_2015	nucleocapsid

MF598611	21456-25517	Middle East respiratory syndrome-related coronavirus	camel/UAE_B19_2015	spike

MF598594	279-21514	Middle East respiratory syndrome-related coronavirus	camel/UAE_B1_2015	orf1ab

MF598594	27853-28512	Middle East respiratory syndrome-related coronavirus	camel/UAE_B1_2015	membrane

MF598594	28566-29807	Middle East respiratory syndrome-related coronavirus	camel/UAE_B1_2015	nucleocapsid

MF598594	21456-25517	Middle East respiratory syndrome-related coronavirus	camel/UAE_B1_2015	spike

MF598612	279-21514	Middle East respiratory syndrome-related coronavirus	camel/UAE_B20_2015	orf1ab

MF598612	27853-28512	Middle East respiratory syndrome-related coronavirus	camel/UAE_B20_2015	membrane

MF598612	28566-29807	Middle East respiratory syndrome-related coronavirus	camel/UAE_B20_2015	nucleocapsid

MF598612	21456-25517	Middle East respiratory syndrome-related coronavirus	camel/UAE_B20_2015	spike

MF598613	279-21514	Middle East respiratory syndrome-related coronavirus	camel/UAE_B21_2015	orf1ab

MF598613	27853-28512	Middle East respiratory syndrome-related coronavirus	camel/UAE_B21_2015	membrane

MF598613	28566-29807	Middle East respiratory syndrome-related coronavirus	camel/UAE_B21_2015	nucleocapsid

MF598613	21456-25517	Middle East respiratory syndrome-related coronavirus	camel/UAE_B21_2015	spike

MF598614	279-21514	Middle East respiratory syndrome-related coronavirus	camel/UAE_B22_2015	orf1ab

MF598614	27853-28512	Middle East respiratory syndrome-related coronavirus	camel/UAE_B22_2015	membrane

MF598614	28566-29807	Middle East respiratory syndrome-related coronavirus	camel/UAE_B22_2015	nucleocapsid

MF598614	21456-25517	Middle East respiratory syndrome-related coronavirus	camel/UAE_B22_2015	spike

MF598615	279-21514	Middle East respiratory syndrome-related coronavirus	camel/UAE_B23_2015	orf1ab

MF598615	27853-28512	Middle East respiratory syndrome-related coronavirus	camel/UAE_B23_2015	membrane

MF598615	28566-29807	Middle East respiratory syndrome-related coronavirus	camel/UAE_B23_2015	nucleocapsid

MF598615	21456-25517	Middle East respiratory syndrome-related coronavirus	camel/UAE_B23_2015	spike

MF598616	279-21514	Middle East respiratory syndrome-related coronavirus	camel/UAE_B24_2015	orf1ab

MF598616	27853-28512	Middle East respiratory syndrome-related coronavirus	camel/UAE_B24_2015	membrane

MF598616	28566-29807	Middle East respiratory syndrome-related coronavirus	camel/UAE_B24_2015	nucleocapsid

MF598616	21456-25517	Middle East respiratory syndrome-related coronavirus	camel/UAE_B24_2015	spike

MF598617	279-21514	Middle East respiratory syndrome-related coronavirus	camel/UAE_B25_2015	orf1ab

MF598617	27853-28512	Middle East respiratory syndrome-related coronavirus	camel/UAE_B25_2015	membrane

MF598617	28566-29807	Middle East respiratory syndrome-related coronavirus	camel/UAE_B25_2015	nucleocapsid

MF598617	21456-25517	Middle East respiratory syndrome-related coronavirus	camel/UAE_B25_2015	spike

MF598618	279-21514	Middle East respiratory syndrome-related coronavirus	camel/UAE_B26_2015	orf1ab

MF598618	27853-28512	Middle East respiratory syndrome-related coronavirus	camel/UAE_B26_2015	membrane

MF598618	28566-29807	Middle East respiratory syndrome-related coronavirus	camel/UAE_B26_2015	nucleocapsid

MF598618	21456-25517	Middle East respiratory syndrome-related coronavirus	camel/UAE_B26_2015	spike

MF598619	279-21514	Middle East respiratory syndrome-related coronavirus	camel/UAE_B27_2015	orf1ab

MF598619	27853-28512	Middle East respiratory syndrome-related coronavirus	camel/UAE_B27_2015	membrane

MF598619	28566-29807	Middle East respiratory syndrome-related coronavirus	camel/UAE_B27_2015	nucleocapsid

MF598619	21456-25517	Middle East respiratory syndrome-related coronavirus	camel/UAE_B27_2015	spike

MF598620	279-21514	Middle East respiratory syndrome-related coronavirus	camel/UAE_B28_2015	orf1ab

MF598620	27853-28512	Middle East respiratory syndrome-related coronavirus	camel/UAE_B28_2015	membrane

MF598620	28566-29807	Middle East respiratory syndrome-related coronavirus	camel/UAE_B28_2015	nucleocapsid

MF598620	21456-25517	Middle East respiratory syndrome-related coronavirus	camel/UAE_B28_2015	spike

MF598621	279-21514	Middle East respiratory syndrome-related coronavirus	camel/UAE_B29_2015	orf1ab

MF598621	27853-28512	Middle East respiratory syndrome-related coronavirus	camel/UAE_B29_2015	membrane

MF598621	28566-29807	Middle East respiratory syndrome-related coronavirus	camel/UAE_B29_2015	nucleocapsid

MF598621	21456-25517	Middle East respiratory syndrome-related coronavirus	camel/UAE_B29_2015	spike

MF598595	279-21514	Middle East respiratory syndrome-related coronavirus	camel/UAE_B2_2015	orf1ab

MF598595	27853-28512	Middle East respiratory syndrome-related coronavirus	camel/UAE_B2_2015	membrane

MF598595	28566-29807	Middle East respiratory syndrome-related coronavirus	camel/UAE_B2_2015	nucleocapsid

MF598595	21456-25517	Middle East respiratory syndrome-related coronavirus	camel/UAE_B2_2015	spike

MF598622	279-21514	Middle East respiratory syndrome-related coronavirus	camel/UAE_B30_2015	orf1ab

MF598622	27853-28512	Middle East respiratory syndrome-related coronavirus	camel/UAE_B30_2015	membrane

MF598622	28566-29807	Middle East respiratory syndrome-related coronavirus	camel/UAE_B30_2015	nucleocapsid

MF598622	21456-25517	Middle East respiratory syndrome-related coronavirus	camel/UAE_B30_2015	spike

MF598623	279-21514	Middle East respiratory syndrome-related coronavirus	camel/UAE_B31_2015	orf1ab

MF598623	27853-28512	Middle East respiratory syndrome-related coronavirus	camel/UAE_B31_2015	membrane

MF598623	28566-29807	Middle East respiratory syndrome-related coronavirus	camel/UAE_B31_2015	nucleocapsid

MF598623	21456-25517	Middle East respiratory syndrome-related coronavirus	camel/UAE_B31_2015	spike

MF598624	279-21514	Middle East respiratory syndrome-related coronavirus	camel/UAE_B32_2015	orf1ab

MF598624	27853-28512	Middle East respiratory syndrome-related coronavirus	camel/UAE_B32_2015	membrane

MF598624	28566-29807	Middle East respiratory syndrome-related coronavirus	camel/UAE_B32_2015	nucleocapsid

MF598624	21456-25517	Middle East respiratory syndrome-related coronavirus	camel/UAE_B32_2015	spike

MF598625	279-21514	Middle East respiratory syndrome-related coronavirus	camel/UAE_B33_2015	orf1ab

MF598625	27853-28512	Middle East respiratory syndrome-related coronavirus	camel/UAE_B33_2015	membrane

MF598625	28566-29807	Middle East respiratory syndrome-related coronavirus	camel/UAE_B33_2015	nucleocapsid

MF598625	21456-25517	Middle East respiratory syndrome-related coronavirus	camel/UAE_B33_2015	spike

MF598626	279-21514	Middle East respiratory syndrome-related coronavirus	camel/UAE_B34_2015	orf1ab

MF598626	27853-28512	Middle East respiratory syndrome-related coronavirus	camel/UAE_B34_2015	membrane

MF598626	28566-29807	Middle East respiratory syndrome-related coronavirus	camel/UAE_B34_2015	nucleocapsid

MF598626	21456-25517	Middle East respiratory syndrome-related coronavirus	camel/UAE_B34_2015	spike

MF598627	279-21514	Middle East respiratory syndrome-related coronavirus	camel/UAE_B35_2015	orf1ab

MF598627	27853-28512	Middle East respiratory syndrome-related coronavirus	camel/UAE_B35_2015	membrane

MF598627	28566-29807	Middle East respiratory syndrome-related coronavirus	camel/UAE_B35_2015	nucleocapsid

MF598627	21456-25517	Middle East respiratory syndrome-related coronavirus	camel/UAE_B35_2015	spike

MF598629	279-21514	Middle East respiratory syndrome-related coronavirus	camel/UAE_B37_2015	orf1ab

MF598629	27853-28512	Middle East respiratory syndrome-related coronavirus	camel/UAE_B37_2015	membrane

MF598629	28566-29807	Middle East respiratory syndrome-related coronavirus	camel/UAE_B37_2015	nucleocapsid

MF598629	21456-25517	Middle East respiratory syndrome-related coronavirus	camel/UAE_B37_2015	spike

MF598630	279-21514	Middle East respiratory syndrome-related coronavirus	camel/UAE_B38_2015	orf1ab

MF598630	27853-28512	Middle East respiratory syndrome-related coronavirus	camel/UAE_B38_2015	membrane

MF598630	28566-29807	Middle East respiratory syndrome-related coronavirus	camel/UAE_B38_2015	nucleocapsid

MF598630	21456-25517	Middle East respiratory syndrome-related coronavirus	camel/UAE_B38_2015	spike

MF598631	279-21514	Middle East respiratory syndrome-related coronavirus	camel/UAE_B39_2015	orf1ab

MF598631	27853-28512	Middle East respiratory syndrome-related coronavirus	camel/UAE_B39_2015	membrane

MF598631	28566-29807	Middle East respiratory syndrome-related coronavirus	camel/UAE_B39_2015	nucleocapsid

MF598631	21456-25517	Middle East respiratory syndrome-related coronavirus	camel/UAE_B39_2015	spike

MF598632	279-21514	Middle East respiratory syndrome-related coronavirus	camel/UAE_B40_2015	orf1ab

MF598632	27853-28512	Middle East respiratory syndrome-related coronavirus	camel/UAE_B40_2015	membrane

MF598632	28566-29807	Middle East respiratory syndrome-related coronavirus	camel/UAE_B40_2015	nucleocapsid

MF598632	21456-25517	Middle East respiratory syndrome-related coronavirus	camel/UAE_B40_2015	spike

MF598634	279-21514	Middle East respiratory syndrome-related coronavirus	camel/UAE_B42_2015	orf1ab

MF598634	27853-28512	Middle East respiratory syndrome-related coronavirus	camel/UAE_B42_2015	membrane

MF598634	28566-29807	Middle East respiratory syndrome-related coronavirus	camel/UAE_B42_2015	nucleocapsid

MF598634	21456-25517	Middle East respiratory syndrome-related coronavirus	camel/UAE_B42_2015	spike

MF598635	279-21514	Middle East respiratory syndrome-related coronavirus	camel/UAE_B44_2015	orf1ab

MF598635	27853-28512	Middle East respiratory syndrome-related coronavirus	camel/UAE_B44_2015	membrane

MF598635	28566-29807	Middle East respiratory syndrome-related coronavirus	camel/UAE_B44_2015	nucleocapsid

MF598635	21456-25517	Middle East respiratory syndrome-related coronavirus	camel/UAE_B44_2015	spike

MF598636	279-21514	Middle East respiratory syndrome-related coronavirus	camel/UAE_B45_2015	orf1ab

MF598636	27853-28512	Middle East respiratory syndrome-related coronavirus	camel/UAE_B45_2015	membrane

MF598636	28566-29807	Middle East respiratory syndrome-related coronavirus	camel/UAE_B45_2015	nucleocapsid

MF598636	21456-25517	Middle East respiratory syndrome-related coronavirus	camel/UAE_B45_2015	spike

MF598637	279-21514	Middle East respiratory syndrome-related coronavirus	camel/UAE_B46_2015	orf1ab

MF598637	27853-28512	Middle East respiratory syndrome-related coronavirus	camel/UAE_B46_2015	membrane

MF598637	28566-29807	Middle East respiratory syndrome-related coronavirus	camel/UAE_B46_2015	nucleocapsid

MF598637	21456-25517	Middle East respiratory syndrome-related coronavirus	camel/UAE_B46_2015	spike

MF598638	279-21514	Middle East respiratory syndrome-related coronavirus	camel/UAE_B47_2015	orf1ab

MF598638	27853-28512	Middle East respiratory syndrome-related coronavirus	camel/UAE_B47_2015	membrane

MF598638	28566-29807	Middle East respiratory syndrome-related coronavirus	camel/UAE_B47_2015	nucleocapsid

MF598638	21456-25517	Middle East respiratory syndrome-related coronavirus	camel/UAE_B47_2015	spike

MF598639	279-21514	Middle East respiratory syndrome-related coronavirus	camel/UAE_B48_2015	orf1ab

MF598639	27853-28512	Middle East respiratory syndrome-related coronavirus	camel/UAE_B48_2015	membrane

MF598639	28566-29807	Middle East respiratory syndrome-related coronavirus	camel/UAE_B48_2015	nucleocapsid

MF598639	21456-25517	Middle East respiratory syndrome-related coronavirus	camel/UAE_B48_2015	spike

MF598640	279-21514	Middle East respiratory syndrome-related coronavirus	camel/UAE_B49_2015	orf1ab

MF598640	27853-28512	Middle East respiratory syndrome-related coronavirus	camel/UAE_B49_2015	membrane

MF598640	28566-29807	Middle East respiratory syndrome-related coronavirus	camel/UAE_B49_2015	nucleocapsid

MF598640	21456-25517	Middle East respiratory syndrome-related coronavirus	camel/UAE_B49_2015	spike

MF598596	279-21514	Middle East respiratory syndrome-related coronavirus	camel/UAE_B4_2015	orf1ab

MF598596	27853-28512	Middle East respiratory syndrome-related coronavirus	camel/UAE_B4_2015	membrane

MF598596	28566-29807	Middle East respiratory syndrome-related coronavirus	camel/UAE_B4_2015	nucleocapsid

MF598596	21456-25517	Middle East respiratory syndrome-related coronavirus	camel/UAE_B4_2015	spike

MF598641	279-21514	Middle East respiratory syndrome-related coronavirus	camel/UAE_B50_2015	orf1ab

MF598641	27853-28512	Middle East respiratory syndrome-related coronavirus	camel/UAE_B50_2015	membrane

MF598641	28566-29807	Middle East respiratory syndrome-related coronavirus	camel/UAE_B50_2015	nucleocapsid

MF598641	21456-25517	Middle East respiratory syndrome-related coronavirus	camel/UAE_B50_2015	spike

MF598643	279-21514	Middle East respiratory syndrome-related coronavirus	camel/UAE_B52_2015	orf1ab

MF598643	27853-28512	Middle East respiratory syndrome-related coronavirus	camel/UAE_B52_2015	membrane

MF598643	28566-29807	Middle East respiratory syndrome-related coronavirus	camel/UAE_B52_2015	nucleocapsid

MF598643	21456-25517	Middle East respiratory syndrome-related coronavirus	camel/UAE_B52_2015	spike

MF598644	279-21514	Middle East respiratory syndrome-related coronavirus	camel/UAE_B53_2015	orf1ab

MF598644	27853-28512	Middle East respiratory syndrome-related coronavirus	camel/UAE_B53_2015	membrane

MF598644	28566-29807	Middle East respiratory syndrome-related coronavirus	camel/UAE_B53_2015	nucleocapsid

MF598644	21456-25517	Middle East respiratory syndrome-related coronavirus	camel/UAE_B53_2015	spike

MF598645	279-21514	Middle East respiratory syndrome-related coronavirus	camel/UAE_B54_2015	orf1ab

MF598645	27853-28512	Middle East respiratory syndrome-related coronavirus	camel/UAE_B54_2015	membrane

MF598645	28566-29807	Middle East respiratory syndrome-related coronavirus	camel/UAE_B54_2015	nucleocapsid

MF598645	21456-25517	Middle East respiratory syndrome-related coronavirus	camel/UAE_B54_2015	spike

MF598646	279-21514	Middle East respiratory syndrome-related coronavirus	camel/UAE_B55_2015	orf1ab

MF598646	27853-28512	Middle East respiratory syndrome-related coronavirus	camel/UAE_B55_2015	membrane

MF598646	28566-29807	Middle East respiratory syndrome-related coronavirus	camel/UAE_B55_2015	nucleocapsid

MF598646	21456-25517	Middle East respiratory syndrome-related coronavirus	camel/UAE_B55_2015	spike

MF598647	279-21514	Middle East respiratory syndrome-related coronavirus	camel/UAE_B56_2015	orf1ab

MF598647	27853-28512	Middle East respiratory syndrome-related coronavirus	camel/UAE_B56_2015	membrane

MF598647	28566-29807	Middle East respiratory syndrome-related coronavirus	camel/UAE_B56_2015	nucleocapsid

MF598647	21456-25517	Middle East respiratory syndrome-related coronavirus	camel/UAE_B56_2015	spike

MF598648	279-21514	Middle East respiratory syndrome-related coronavirus	camel/UAE_B58_2015	orf1ab

MF598648	27853-28512	Middle East respiratory syndrome-related coronavirus	camel/UAE_B58_2015	membrane

MF598648	28566-29807	Middle East respiratory syndrome-related coronavirus	camel/UAE_B58_2015	nucleocapsid

MF598648	21456-25517	Middle East respiratory syndrome-related coronavirus	camel/UAE_B58_2015	spike

MF598649	279-21514	Middle East respiratory syndrome-related coronavirus	camel/UAE_B59_2015	orf1ab

MF598649	27853-28512	Middle East respiratory syndrome-related coronavirus	camel/UAE_B59_2015	membrane

MF598649	28566-29807	Middle East respiratory syndrome-related coronavirus	camel/UAE_B59_2015	nucleocapsid

MF598649	21456-25517	Middle East respiratory syndrome-related coronavirus	camel/UAE_B59_2015	spike

MF598597	279-21514	Middle East respiratory syndrome-related coronavirus	camel/UAE_B5_2015	orf1ab

MF598597	27853-28512	Middle East respiratory syndrome-related coronavirus	camel/UAE_B5_2015	membrane

MF598597	28566-29807	Middle East respiratory syndrome-related coronavirus	camel/UAE_B5_2015	nucleocapsid

MF598597	21456-25517	Middle East respiratory syndrome-related coronavirus	camel/UAE_B5_2015	spike

MF598650	279-21514	Middle East respiratory syndrome-related coronavirus	camel/UAE_B60_2015	orf1ab

MF598650	27853-28512	Middle East respiratory syndrome-related coronavirus	camel/UAE_B60_2015	membrane

MF598650	28566-29807	Middle East respiratory syndrome-related coronavirus	camel/UAE_B60_2015	nucleocapsid

MF598650	21456-25517	Middle East respiratory syndrome-related coronavirus	camel/UAE_B60_2015	spike

MF598651	279-21514	Middle East respiratory syndrome-related coronavirus	camel/UAE_B61_2015	orf1ab

MF598651	27853-28512	Middle East respiratory syndrome-related coronavirus	camel/UAE_B61_2015	membrane

MF598651	28566-29807	Middle East respiratory syndrome-related coronavirus	camel/UAE_B61_2015	nucleocapsid

MF598651	21456-25517	Middle East respiratory syndrome-related coronavirus	camel/UAE_B61_2015	spike

MF598652	279-21514	Middle East respiratory syndrome-related coronavirus	camel/UAE_B62_2015	orf1ab

MF598652	27853-28512	Middle East respiratory syndrome-related coronavirus	camel/UAE_B62_2015	membrane

MF598652	28566-29807	Middle East respiratory syndrome-related coronavirus	camel/UAE_B62_2015	nucleocapsid

MF598652	21456-25517	Middle East respiratory syndrome-related coronavirus	camel/UAE_B62_2015	spike

MF598653	279-21514	Middle East respiratory syndrome-related coronavirus	camel/UAE_B63_2015	orf1ab

MF598653	27853-28512	Middle East respiratory syndrome-related coronavirus	camel/UAE_B63_2015	membrane

MF598653	28566-29807	Middle East respiratory syndrome-related coronavirus	camel/UAE_B63_2015	nucleocapsid

MF598653	21456-25517	Middle East respiratory syndrome-related coronavirus	camel/UAE_B63_2015	spike

MF598654	279-21514	Middle East respiratory syndrome-related coronavirus	camel/UAE_B64_2015	orf1ab

MF598654	27853-28512	Middle East respiratory syndrome-related coronavirus	camel/UAE_B64_2015	membrane

MF598654	28566-29807	Middle East respiratory syndrome-related coronavirus	camel/UAE_B64_2015	nucleocapsid

MF598654	21456-25517	Middle East respiratory syndrome-related coronavirus	camel/UAE_B64_2015	spike

MF598655	279-21514	Middle East respiratory syndrome-related coronavirus	camel/UAE_B65_2015	orf1ab

MF598655	27853-28512	Middle East respiratory syndrome-related coronavirus	camel/UAE_B65_2015	membrane

MF598655	28566-29807	Middle East respiratory syndrome-related coronavirus	camel/UAE_B65_2015	nucleocapsid

MF598655	21456-25517	Middle East respiratory syndrome-related coronavirus	camel/UAE_B65_2015	spike

MF598656	279-21514	Middle East respiratory syndrome-related coronavirus	camel/UAE_B66_2015	orf1ab

MF598656	27853-28512	Middle East respiratory syndrome-related coronavirus	camel/UAE_B66_2015	membrane

MF598656	28566-29807	Middle East respiratory syndrome-related coronavirus	camel/UAE_B66_2015	nucleocapsid

MF598656	21456-25517	Middle East respiratory syndrome-related coronavirus	camel/UAE_B66_2015	spike

MF598657	279-21514	Middle East respiratory syndrome-related coronavirus	camel/UAE_B67_2015	orf1ab

MF598657	27853-28512	Middle East respiratory syndrome-related coronavirus	camel/UAE_B67_2015	membrane

MF598657	28566-29807	Middle East respiratory syndrome-related coronavirus	camel/UAE_B67_2015	nucleocapsid

MF598657	21456-25517	Middle East respiratory syndrome-related coronavirus	camel/UAE_B67_2015	spike

MF598658	279-21514	Middle East respiratory syndrome-related coronavirus	camel/UAE_B68_2015	orf1ab

MF598658	27853-28512	Middle East respiratory syndrome-related coronavirus	camel/UAE_B68_2015	membrane

MF598658	28566-29807	Middle East respiratory syndrome-related coronavirus	camel/UAE_B68_2015	nucleocapsid

MF598658	21456-25517	Middle East respiratory syndrome-related coronavirus	camel/UAE_B68_2015	spike

MF598659	279-21514	Middle East respiratory syndrome-related coronavirus	camel/UAE_B69_2015	orf1ab

MF598659	27853-28512	Middle East respiratory syndrome-related coronavirus	camel/UAE_B69_2015	membrane

MF598659	28566-29807	Middle East respiratory syndrome-related coronavirus	camel/UAE_B69_2015	nucleocapsid

MF598659	21456-25517	Middle East respiratory syndrome-related coronavirus	camel/UAE_B69_2015	spike

MF598598	279-21514	Middle East respiratory syndrome-related coronavirus	camel/UAE_B6_2015	orf1ab

MF598598	27853-28512	Middle East respiratory syndrome-related coronavirus	camel/UAE_B6_2015	membrane

MF598598	28566-29807	Middle East respiratory syndrome-related coronavirus	camel/UAE_B6_2015	nucleocapsid

MF598598	21456-25517	Middle East respiratory syndrome-related coronavirus	camel/UAE_B6_2015	spike

MF598660	279-21514	Middle East respiratory syndrome-related coronavirus	camel/UAE_B70_2015	orf1ab

MF598660	27853-28512	Middle East respiratory syndrome-related coronavirus	camel/UAE_B70_2015	membrane

MF598660	28566-29807	Middle East respiratory syndrome-related coronavirus	camel/UAE_B70_2015	nucleocapsid

MF598660	21456-25517	Middle East respiratory syndrome-related coronavirus	camel/UAE_B70_2015	spike

MF598661	279-21514	Middle East respiratory syndrome-related coronavirus	camel/UAE_B71_2015	orf1ab

MF598661	27853-28512	Middle East respiratory syndrome-related coronavirus	camel/UAE_B71_2015	membrane

MF598661	28566-29807	Middle East respiratory syndrome-related coronavirus	camel/UAE_B71_2015	nucleocapsid

MF598661	21456-25517	Middle East respiratory syndrome-related coronavirus	camel/UAE_B71_2015	spike

MF598662	279-21514	Middle East respiratory syndrome-related coronavirus	camel/UAE_B72_2015	orf1ab

MF598662	27853-28512	Middle East respiratory syndrome-related coronavirus	camel/UAE_B72_2015	membrane

MF598662	28566-29807	Middle East respiratory syndrome-related coronavirus	camel/UAE_B72_2015	nucleocapsid

MF598662	21456-25517	Middle East respiratory syndrome-related coronavirus	camel/UAE_B72_2015	spike

MF598663	279-21514	Middle East respiratory syndrome-related coronavirus	camel/UAE_B73_2015	orf1ab

MF598663	27853-28512	Middle East respiratory syndrome-related coronavirus	camel/UAE_B73_2015	membrane

MF598663	28566-29807	Middle East respiratory syndrome-related coronavirus	camel/UAE_B73_2015	nucleocapsid

MF598663	21456-25517	Middle East respiratory syndrome-related coronavirus	camel/UAE_B73_2015	spike

MF598664	279-21514	Middle East respiratory syndrome-related coronavirus	camel/UAE_B74_2015	orf1ab

MF598664	27853-28512	Middle East respiratory syndrome-related coronavirus	camel/UAE_B74_2015	membrane

MF598664	28566-29807	Middle East respiratory syndrome-related coronavirus	camel/UAE_B74_2015	nucleocapsid

MF598664	21456-25517	Middle East respiratory syndrome-related coronavirus	camel/UAE_B74_2015	spike

MF598665	279-21514	Middle East respiratory syndrome-related coronavirus	camel/UAE_B75_2015	orf1ab

MF598665	27853-28512	Middle East respiratory syndrome-related coronavirus	camel/UAE_B75_2015	membrane

MF598665	28566-29807	Middle East respiratory syndrome-related coronavirus	camel/UAE_B75_2015	nucleocapsid

MF598665	21456-25517	Middle East respiratory syndrome-related coronavirus	camel/UAE_B75_2015	spike

MF598666	279-21514	Middle East respiratory syndrome-related coronavirus	camel/UAE_B76_2015	orf1ab

MF598666	27853-28512	Middle East respiratory syndrome-related coronavirus	camel/UAE_B76_2015	membrane

MF598666	28566-29807	Middle East respiratory syndrome-related coronavirus	camel/UAE_B76_2015	nucleocapsid

MF598666	21456-25517	Middle East respiratory syndrome-related coronavirus	camel/UAE_B76_2015	spike

MF598667	279-21514	Middle East respiratory syndrome-related coronavirus	camel/UAE_B77_2015	orf1ab

MF598667	27853-28512	Middle East respiratory syndrome-related coronavirus	camel/UAE_B77_2015	membrane

MF598667	28566-29807	Middle East respiratory syndrome-related coronavirus	camel/UAE_B77_2015	nucleocapsid

MF598667	21456-25517	Middle East respiratory syndrome-related coronavirus	camel/UAE_B77_2015	spike

MF598668	279-21514	Middle East respiratory syndrome-related coronavirus	camel/UAE_B78_2015	orf1ab

MF598668	27853-28512	Middle East respiratory syndrome-related coronavirus	camel/UAE_B78_2015	membrane

MF598668	28566-29807	Middle East respiratory syndrome-related coronavirus	camel/UAE_B78_2015	nucleocapsid

MF598668	21456-25517	Middle East respiratory syndrome-related coronavirus	camel/UAE_B78_2015	spike

MF598669	279-21514	Middle East respiratory syndrome-related coronavirus	camel/UAE_B79_2015	orf1ab

MF598669	27853-28512	Middle East respiratory syndrome-related coronavirus	camel/UAE_B79_2015	membrane

MF598669	28566-29807	Middle East respiratory syndrome-related coronavirus	camel/UAE_B79_2015	nucleocapsid

MF598669	21456-25517	Middle East respiratory syndrome-related coronavirus	camel/UAE_B79_2015	spike

MF598599	279-21514	Middle East respiratory syndrome-related coronavirus	camel/UAE_B7_2015	orf1ab

MF598599	27853-28512	Middle East respiratory syndrome-related coronavirus	camel/UAE_B7_2015	membrane

MF598599	28566-29807	Middle East respiratory syndrome-related coronavirus	camel/UAE_B7_2015	nucleocapsid

MF598599	21456-25517	Middle East respiratory syndrome-related coronavirus	camel/UAE_B7_2015	spike

MF598670	279-21514	Middle East respiratory syndrome-related coronavirus	camel/UAE_B80_2015	orf1ab

MF598670	27853-28512	Middle East respiratory syndrome-related coronavirus	camel/UAE_B80_2015	membrane

MF598670	28566-29807	Middle East respiratory syndrome-related coronavirus	camel/UAE_B80_2015	nucleocapsid

MF598670	21456-25517	Middle East respiratory syndrome-related coronavirus	camel/UAE_B80_2015	spike

MF598671	279-21514	Middle East respiratory syndrome-related coronavirus	camel/UAE_B81_2015	orf1ab

MF598671	27853-28512	Middle East respiratory syndrome-related coronavirus	camel/UAE_B81_2015	membrane

MF598671	28566-29807	Middle East respiratory syndrome-related coronavirus	camel/UAE_B81_2015	nucleocapsid

MF598671	21456-25517	Middle East respiratory syndrome-related coronavirus	camel/UAE_B81_2015	spike

MF598672	279-21514	Middle East respiratory syndrome-related coronavirus	camel/UAE_B82_2015	orf1ab

MF598672	27853-28512	Middle East respiratory syndrome-related coronavirus	camel/UAE_B82_2015	membrane

MF598672	28566-29807	Middle East respiratory syndrome-related coronavirus	camel/UAE_B82_2015	nucleocapsid

MF598672	21456-25517	Middle East respiratory syndrome-related coronavirus	camel/UAE_B82_2015	spike

MF598673	279-21514	Middle East respiratory syndrome-related coronavirus	camel/UAE_B83_2015	orf1ab

MF598673	27853-28512	Middle East respiratory syndrome-related coronavirus	camel/UAE_B83_2015	membrane

MF598673	28566-29807	Middle East respiratory syndrome-related coronavirus	camel/UAE_B83_2015	nucleocapsid

MF598673	21456-25517	Middle East respiratory syndrome-related coronavirus	camel/UAE_B83_2015	spike

MF598674	279-21514	Middle East respiratory syndrome-related coronavirus	camel/UAE_B84_2015	orf1ab

MF598674	27853-28512	Middle East respiratory syndrome-related coronavirus	camel/UAE_B84_2015	membrane

MF598674	28566-29807	Middle East respiratory syndrome-related coronavirus	camel/UAE_B84_2015	nucleocapsid

MF598674	21456-25517	Middle East respiratory syndrome-related coronavirus	camel/UAE_B84_2015	spike

MF598675	279-21514	Middle East respiratory syndrome-related coronavirus	camel/UAE_B85_2015	orf1ab

MF598675	27853-28512	Middle East respiratory syndrome-related coronavirus	camel/UAE_B85_2015	membrane

MF598675	28566-29807	Middle East respiratory syndrome-related coronavirus	camel/UAE_B85_2015	nucleocapsid

MF598675	21456-25517	Middle East respiratory syndrome-related coronavirus	camel/UAE_B85_2015	spike

MF598676	279-21514	Middle East respiratory syndrome-related coronavirus	camel/UAE_B86_2015	orf1ab

MF598676	27853-28512	Middle East respiratory syndrome-related coronavirus	camel/UAE_B86_2015	membrane

MF598676	28566-29807	Middle East respiratory syndrome-related coronavirus	camel/UAE_B86_2015	nucleocapsid

MF598676	21456-25517	Middle East respiratory syndrome-related coronavirus	camel/UAE_B86_2015	spike

MF598677	279-21514	Middle East respiratory syndrome-related coronavirus	camel/UAE_B87_2015	orf1ab

MF598677	27853-28512	Middle East respiratory syndrome-related coronavirus	camel/UAE_B87_2015	membrane

MF598677	28566-29807	Middle East respiratory syndrome-related coronavirus	camel/UAE_B87_2015	nucleocapsid

MF598677	21456-25517	Middle East respiratory syndrome-related coronavirus	camel/UAE_B87_2015	spike

MF598678	279-21514	Middle East respiratory syndrome-related coronavirus	camel/UAE_B88_2015	orf1ab

MF598678	27853-28512	Middle East respiratory syndrome-related coronavirus	camel/UAE_B88_2015	membrane

MF598678	28566-29807	Middle East respiratory syndrome-related coronavirus	camel/UAE_B88_2015	nucleocapsid

MF598678	21456-25517	Middle East respiratory syndrome-related coronavirus	camel/UAE_B88_2015	spike

MF598600	279-21514	Middle East respiratory syndrome-related coronavirus	camel/UAE_B8_2015	orf1ab

MF598600	27853-28512	Middle East respiratory syndrome-related coronavirus	camel/UAE_B8_2015	membrane

MF598600	28566-29807	Middle East respiratory syndrome-related coronavirus	camel/UAE_B8_2015	nucleocapsid

MF598600	21456-25517	Middle East respiratory syndrome-related coronavirus	camel/UAE_B8_2015	spike

MF598679	279-21514	Middle East respiratory syndrome-related coronavirus	camel/UAE_B90_2015	orf1ab

MF598679	27853-28512	Middle East respiratory syndrome-related coronavirus	camel/UAE_B90_2015	membrane

MF598679	28566-29807	Middle East respiratory syndrome-related coronavirus	camel/UAE_B90_2015	nucleocapsid

MF598679	21456-25517	Middle East respiratory syndrome-related coronavirus	camel/UAE_B90_2015	spike

MF598680	279-21514	Middle East respiratory syndrome-related coronavirus	camel/UAE_B91_2015	orf1ab

MF598680	27853-28512	Middle East respiratory syndrome-related coronavirus	camel/UAE_B91_2015	membrane

MF598680	28566-29807	Middle East respiratory syndrome-related coronavirus	camel/UAE_B91_2015	nucleocapsid

MF598680	21456-25517	Middle East respiratory syndrome-related coronavirus	camel/UAE_B91_2015	spike

MF598681	279-21514	Middle East respiratory syndrome-related coronavirus	camel/UAE_B92_2015	orf1ab

MF598681	27853-28512	Middle East respiratory syndrome-related coronavirus	camel/UAE_B92_2015	membrane

MF598681	28566-29807	Middle East respiratory syndrome-related coronavirus	camel/UAE_B92_2015	nucleocapsid

MF598681	21456-25517	Middle East respiratory syndrome-related coronavirus	camel/UAE_B92_2015	spike

MF598682	279-21514	Middle East respiratory syndrome-related coronavirus	camel/UAE_B93_2015	orf1ab

MF598682	27853-28512	Middle East respiratory syndrome-related coronavirus	camel/UAE_B93_2015	membrane

MF598682	28566-29807	Middle East respiratory syndrome-related coronavirus	camel/UAE_B93_2015	nucleocapsid

MF598682	21456-25517	Middle East respiratory syndrome-related coronavirus	camel/UAE_B93_2015	spike

MF598683	279-21514	Middle East respiratory syndrome-related coronavirus	camel/UAE_B95_2015	orf1ab

MF598683	27853-28512	Middle East respiratory syndrome-related coronavirus	camel/UAE_B95_2015	membrane

MF598683	28566-29807	Middle East respiratory syndrome-related coronavirus	camel/UAE_B95_2015	nucleocapsid

MF598683	21456-25517	Middle East respiratory syndrome-related coronavirus	camel/UAE_B95_2015	spike

MF598684	279-21514	Middle East respiratory syndrome-related coronavirus	camel/UAE_B96_2015	orf1ab

MF598684	27853-28512	Middle East respiratory syndrome-related coronavirus	camel/UAE_B96_2015	membrane

MF598684	28566-29807	Middle East respiratory syndrome-related coronavirus	camel/UAE_B96_2015	nucleocapsid

MF598684	21456-25517	Middle East respiratory syndrome-related coronavirus	camel/UAE_B96_2015	spike

MF598686	279-21514	Middle East respiratory syndrome-related coronavirus	camel/UAE_B99_2015	orf1ab

MF598686	27853-28512	Middle East respiratory syndrome-related coronavirus	camel/UAE_B99_2015	membrane

MF598686	28566-29807	Middle East respiratory syndrome-related coronavirus	camel/UAE_B99_2015	nucleocapsid

MF598686	21456-25517	Middle East respiratory syndrome-related coronavirus	camel/UAE_B99_2015	spike

MF598601	279-21514	Middle East respiratory syndrome-related coronavirus	camel/UAE_B9_2015	orf1ab

MF598601	27853-28512	Middle East respiratory syndrome-related coronavirus	camel/UAE_B9_2015	membrane

MF598601	28566-29807	Middle East respiratory syndrome-related coronavirus	camel/UAE_B9_2015	nucleocapsid

MF598601	21456-25517	Middle East respiratory syndrome-related coronavirus	camel/UAE_B9_2015	spike

MH013216	279-21514	Middle East respiratory syndrome-related coronavirus	HCoV-EMC	orf1ab

MH013216	27853-28512	Middle East respiratory syndrome-related coronavirus	HCoV-EMC	membrane

MH013216	28566-29807	Middle East respiratory syndrome-related coronavirus	HCoV-EMC	nucleocapsid

MH013216	21456-25517	Middle East respiratory syndrome-related coronavirus	HCoV-EMC	spike

MH306207	279-21514	Middle East respiratory syndrome-related coronavirus	HCoV-EMC	orf1ab

MH306207	27853-28512	Middle East respiratory syndrome-related coronavirus	HCoV-EMC	membrane

MH306207	28566-29807	Middle East respiratory syndrome-related coronavirus	HCoV-EMC	nucleocapsid

MH306207	21456-25517	Middle East respiratory syndrome-related coronavirus	HCoV-EMC	spike

MH454272	279-21514	Middle East respiratory syndrome-related coronavirus	HCoV-EMC	orf1ab

MH454272	27853-28512	Middle East respiratory syndrome-related coronavirus	HCoV-EMC	membrane

MH454272	28566-29807	Middle East respiratory syndrome-related coronavirus	HCoV-EMC	nucleocapsid

MH454272	21456-25517	Middle East respiratory syndrome-related coronavirus	HCoV-EMC	spike

NC_019843	279-21514	Middle East respiratory syndrome-related coronavirus	HCoV-EMC	orf1ab

NC_019843	27853-28512	Middle East respiratory syndrome-related coronavirus	HCoV-EMC	membrane

NC_019843	28566-29807	Middle East respiratory syndrome-related coronavirus	HCoV-EMC	nucleocapsid

NC_019843	21456-25517	Middle East respiratory syndrome-related coronavirus	HCoV-EMC	spike

MK483839	27853-28512	Middle East respiratory syndrome-related coronavirus	Hu/Albaha-KSA-0800H/2018	membrane

MK483839	28566-29807	Middle East respiratory syndrome-related coronavirus	Hu/Albaha-KSA-0800H/2018	nucleocapsid

MK483839	279-21514	Middle East respiratory syndrome-related coronavirus	Hu/Albaha-KSA-0800H/2018	orf1ab

MK483839	21456-25517	Middle East respiratory syndrome-related coronavirus	Hu/Albaha-KSA-0800H/2018	spike

MF000457	231-21466	Middle East respiratory syndrome-related coronavirus	Hu/Amman-Jordan-12641/2015	orf1ab

MF000457	27757-28416	Middle East respiratory syndrome-related coronavirus	Hu/Amman-Jordan-12641/2015	membrane

MF000457	28470-29711	Middle East respiratory syndrome-related coronavirus	Hu/Amman-Jordan-12641/2015	nucleocapsid

MF000457	21408-25469	Middle East respiratory syndrome-related coronavirus	Hu/Amman-Jordan-12641/2015	spike

MF000460	258-21493	Middle East respiratory syndrome-related coronavirus	Hu/Amman-Jordan-12716/2015	orf1ab

MF000460	27775-28434	Middle East respiratory syndrome-related coronavirus	Hu/Amman-Jordan-12716/2015	membrane

MF000460	28488-29729	Middle East respiratory syndrome-related coronavirus	Hu/Amman-Jordan-12716/2015	nucleocapsid

MF000460	21435-25496	Middle East respiratory syndrome-related coronavirus	Hu/Amman-Jordan-12716/2015	spike

MF000459	231-21466	Middle East respiratory syndrome-related coronavirus	Hu/Amman-Jordan-12918/2015	orf1ab

MF000459	27748-28407	Middle East respiratory syndrome-related coronavirus	Hu/Amman-Jordan-12918/2015	membrane

MF000459	28461-29702	Middle East respiratory syndrome-related coronavirus	Hu/Amman-Jordan-12918/2015	nucleocapsid

MF000459	21408-25469	Middle East respiratory syndrome-related coronavirus	Hu/Amman-Jordan-12918/2015	spike

MF000458	258-21493	Middle East respiratory syndrome-related coronavirus	Hu/Amman-Jordan-13030/2015	orf1ab

MF000458	27775-28434	Middle East respiratory syndrome-related coronavirus	Hu/Amman-Jordan-13030/2015	membrane

MF000458	28488-29729	Middle East respiratory syndrome-related coronavirus	Hu/Amman-Jordan-13030/2015	nucleocapsid

MF000458	21435-25496	Middle East respiratory syndrome-related coronavirus	Hu/Amman-Jordan-13030/2015	spike

KX154694	27831-28490	Middle East respiratory syndrome-related coronavirus	Hu/Artawiyah-KSA-13328/2016	membrane

KX154694	28544-29785	Middle East respiratory syndrome-related coronavirus	Hu/Artawiyah-KSA-13328/2016	nucleocapsid

KX154694	257-21492	Middle East respiratory syndrome-related coronavirus	Hu/Artawiyah-KSA-13328/2016	orf1ab

KX154694	21434-25495	Middle East respiratory syndrome-related coronavirus	Hu/Artawiyah-KSA-13328/2016	spike

MK462244	27853-28512	Middle East respiratory syndrome-related coronavirus	Hu/Aseer-KSA-173RS1288/2017	membrane

MK462244	28566-29807	Middle East respiratory syndrome-related coronavirus	Hu/Aseer-KSA-173RS1288/2017	nucleocapsid

MK462244	279-21514	Middle East respiratory syndrome-related coronavirus	Hu/Aseer-KSA-173RS1288/2017	orf1ab

MK462244	21456-25517	Middle East respiratory syndrome-related coronavirus	Hu/Aseer-KSA-173RS1288/2017	spike

KY688119	258-21493	Middle East respiratory syndrome-related coronavirus	Hu/Aseer-KSA-Rs924/2015	orf1ab

KY688119	27791-28450	Middle East respiratory syndrome-related coronavirus	Hu/Aseer-KSA-Rs924/2015	membrane

KY688119	28504-29745	Middle East respiratory syndrome-related coronavirus	Hu/Aseer-KSA-Rs924/2015	nucleocapsid

KY688119	21435-25496	Middle East respiratory syndrome-related coronavirus	Hu/Aseer-KSA-Rs924/2015	spike

KY688120	258-21493	Middle East respiratory syndrome-related coronavirus	Hu/Hufuf-KSA-11002/2015	orf1ab

KY688120	27832-28491	Middle East respiratory syndrome-related coronavirus	Hu/Hufuf-KSA-11002/2015	membrane

KY688120	28545-29786	Middle East respiratory syndrome-related coronavirus	Hu/Hufuf-KSA-11002/2015	nucleocapsid

KY688120	21435-25496	Middle East respiratory syndrome-related coronavirus	Hu/Hufuf-KSA-11002/2015	spike

KY688123	258-21493	Middle East respiratory syndrome-related coronavirus	Hu/Hufuf-KSA-11150/2015	orf1ab

KY688123	27832-28491	Middle East respiratory syndrome-related coronavirus	Hu/Hufuf-KSA-11150/2015	membrane

KY688123	28545-29786	Middle East respiratory syndrome-related coronavirus	Hu/Hufuf-KSA-11150/2015	nucleocapsid

KY688123	21435-25496	Middle East respiratory syndrome-related coronavirus	Hu/Hufuf-KSA-11150/2015	spike

KY688121	258-21493	Middle East respiratory syndrome-related coronavirus	Hu/Hufuf-KSA-11442/2015	orf1ab

KY688121	27832-28491	Middle East respiratory syndrome-related coronavirus	Hu/Hufuf-KSA-11442/2015	membrane

KY688121	28545-29786	Middle East respiratory syndrome-related coronavirus	Hu/Hufuf-KSA-11442/2015	nucleocapsid

KY688121	21435-25496	Middle East respiratory syndrome-related coronavirus	Hu/Hufuf-KSA-11442/2015	spike

KY688122	258-21493	Middle East respiratory syndrome-related coronavirus	Hu/Hufuf-KSA-11767/2015	orf1ab

KY688122	27832-28491	Middle East respiratory syndrome-related coronavirus	Hu/Hufuf-KSA-11767/2015	membrane

KY688122	28545-29786	Middle East respiratory syndrome-related coronavirus	Hu/Hufuf-KSA-11767/2015	nucleocapsid

KY688122	21435-25496	Middle East respiratory syndrome-related coronavirus	Hu/Hufuf-KSA-11767/2015	spike

KX154690	27831-28490	Middle East respiratory syndrome-related coronavirus	Hu/Jeddah-KSA-161RS1146/2016	membrane

KX154690	28544-29785	Middle East respiratory syndrome-related coronavirus	Hu/Jeddah-KSA-161RS1146/2016	nucleocapsid

KX154690	257-21492	Middle East respiratory syndrome-related coronavirus	Hu/Jeddah-KSA-161RS1146/2016	orf1ab

KX154690	21434-25495	Middle East respiratory syndrome-related coronavirus	Hu/Jeddah-KSA-161RS1146/2016	spike

MK462243	27850-28509	Middle East respiratory syndrome-related coronavirus	Hu/Jeddah-KSA-173RS1101/2017	membrane

MK462243	28563-29804	Middle East respiratory syndrome-related coronavirus	Hu/Jeddah-KSA-173RS1101/2017	nucleocapsid

MK462243	279-21514	Middle East respiratory syndrome-related coronavirus	Hu/Jeddah-KSA-173RS1101/2017	orf1ab

MK462243	21456-25517	Middle East respiratory syndrome-related coronavirus	Hu/Jeddah-KSA-173RS1101/2017	spike

MK462245	27850-28509	Middle East respiratory syndrome-related coronavirus	Hu/Jeddah-KSA-173RS1512/2017	membrane

MK462245	28563-29804	Middle East respiratory syndrome-related coronavirus	Hu/Jeddah-KSA-173RS1512/2017	nucleocapsid

MK462245	279-21514	Middle East respiratory syndrome-related coronavirus	Hu/Jeddah-KSA-173RS1512/2017	orf1ab

MK462245	21456-25517	Middle East respiratory syndrome-related coronavirus	Hu/Jeddah-KSA-173RS1512/2017	spike

MK462246	27850-28509	Middle East respiratory syndrome-related coronavirus	Hu/Jeddah-KSA-173RS1570/2017	membrane

MK462246	28563-29804	Middle East respiratory syndrome-related coronavirus	Hu/Jeddah-KSA-173RS1570/2017	nucleocapsid

MK462246	279-21514	Middle East respiratory syndrome-related coronavirus	Hu/Jeddah-KSA-173RS1570/2017	orf1ab

MK462246	21456-25517	Middle East respiratory syndrome-related coronavirus	Hu/Jeddah-KSA-173RS1570/2017	spike

MK462247	27853-28512	Middle East respiratory syndrome-related coronavirus	Hu/Jeddah-KSA-182RS2449/2018	membrane

MK462247	28566-29807	Middle East respiratory syndrome-related coronavirus	Hu/Jeddah-KSA-182RS2449/2018	nucleocapsid

MK462247	279-21514	Middle East respiratory syndrome-related coronavirus	Hu/Jeddah-KSA-182RS2449/2018	orf1ab

MK462247	21456-25517	Middle East respiratory syndrome-related coronavirus	Hu/Jeddah-KSA-182RS2449/2018	spike

MK039552	279-21514	Middle East respiratory syndrome-related coronavirus	Hu/Jordan-201440011123/2014	orf1ab

MK039552	27853-28512	Middle East respiratory syndrome-related coronavirus	Hu/Jordan-201440011123/2014	membrane

MK039552	28566-29807	Middle East respiratory syndrome-related coronavirus	Hu/Jordan-201440011123/2014	nucleocapsid

MK039552	21456-25517	Middle East respiratory syndrome-related coronavirus	Hu/Jordan-201440011123/2014	spike

MK039553	279-21514	Middle East respiratory syndrome-related coronavirus	Hu/Jordan-201440011858/2014	orf1ab

MK039553	27853-28512	Middle East respiratory syndrome-related coronavirus	Hu/Jordan-201440011858/2014	membrane

MK039553	28566-29807	Middle East respiratory syndrome-related coronavirus	Hu/Jordan-201440011858/2014	nucleocapsid

MK039553	21456-25517	Middle East respiratory syndrome-related coronavirus	Hu/Jordan-201440011858/2014	spike

KY688118	231-21466	Middle East respiratory syndrome-related coronavirus	Hu/Khobar-KSA-6736/2015	orf1ab

KY688118	27805-28464	Middle East respiratory syndrome-related coronavirus	Hu/Khobar-KSA-6736/2015	membrane

KY688118	28518-29759	Middle East respiratory syndrome-related coronavirus	Hu/Khobar-KSA-6736/2015	nucleocapsid

KY688118	21408-25469	Middle East respiratory syndrome-related coronavirus	Hu/Khobar-KSA-6736/2015	spike

MG011351	27831-28490	Middle East respiratory syndrome-related coronavirus	Hu/Madinah-KSA-390/2016	membrane

MG011351	28544-29785	Middle East respiratory syndrome-related coronavirus	Hu/Madinah-KSA-390/2016	nucleocapsid

MG011351	257-21492	Middle East respiratory syndrome-related coronavirus	Hu/Madinah-KSA-390/2016	orf1ab

MG011351	21434-25495	Middle East respiratory syndrome-related coronavirus	Hu/Madinah-KSA-390/2016	spike

MK462248	27853-28512	Middle East respiratory syndrome-related coronavirus	Hu/Najran-KSA-182RS2567/2018	membrane

MK462248	28566-29807	Middle East respiratory syndrome-related coronavirus	Hu/Najran-KSA-182RS2567/2018	nucleocapsid

MK462248	279-21514	Middle East respiratory syndrome-related coronavirus	Hu/Najran-KSA-182RS2567/2018	orf1ab

MK462248	21456-25517	Middle East respiratory syndrome-related coronavirus	Hu/Najran-KSA-182RS2567/2018	spike

MK462250	27853-28512	Middle East respiratory syndrome-related coronavirus	Hu/Najran-KSA-183RS279/2018	membrane

MK462250	28566-29807	Middle East respiratory syndrome-related coronavirus	Hu/Najran-KSA-183RS279/2018	nucleocapsid

MK462250	279-21514	Middle East respiratory syndrome-related coronavirus	Hu/Najran-KSA-183RS279/2018	orf1ab

MK462250	21456-25517	Middle East respiratory syndrome-related coronavirus	Hu/Najran-KSA-183RS279/2018	spike

MK462251	27853-28512	Middle East respiratory syndrome-related coronavirus	Hu/Northern-KSA-1847784/2018	membrane

MK462251	28566-29807	Middle East respiratory syndrome-related coronavirus	Hu/Northern-KSA-1847784/2018	nucleocapsid

MK462251	279-21514	Middle East respiratory syndrome-related coronavirus	Hu/Northern-KSA-1847784/2018	orf1ab

MK462251	21456-25517	Middle East respiratory syndrome-related coronavirus	Hu/Northern-KSA-1847784/2018	spike

KY673148	279-21514	Middle East respiratory syndrome-related coronavirus	Hu/Oman_50_2015	orf1ab

KY673148	27853-28512	Middle East respiratory syndrome-related coronavirus	Hu/Oman_50_2015	membrane

KY673148	28566-29807	Middle East respiratory syndrome-related coronavirus	Hu/Oman_50_2015	nucleocapsid

KY673148	21456-25517	Middle East respiratory syndrome-related coronavirus	Hu/Oman_50_2015	spike

MK462254	27853-28512	Middle East respiratory syndrome-related coronavirus	Hu/Qaseem-KSA-18013897/2018	membrane

MK462254	28566-29807	Middle East respiratory syndrome-related coronavirus	Hu/Qaseem-KSA-18013897/2018	nucleocapsid

MK462254	279-21514	Middle East respiratory syndrome-related coronavirus	Hu/Qaseem-KSA-18013897/2018	orf1ab

MK462254	21456-25517	Middle East respiratory syndrome-related coronavirus	Hu/Qaseem-KSA-18013897/2018	spike

MG011341	27831-28490	Middle East respiratory syndrome-related coronavirus	Hu/Qasim-KSA-13922/2016	membrane

MG011341	28544-29785	Middle East respiratory syndrome-related coronavirus	Hu/Qasim-KSA-13922/2016	nucleocapsid

MG011341	257-21492	Middle East respiratory syndrome-related coronavirus	Hu/Qasim-KSA-13922/2016	orf1ab

MG011341	21434-25495	Middle East respiratory syndrome-related coronavirus	Hu/Qasim-KSA-13922/2016	spike

MG011352	27831-28490	Middle East respiratory syndrome-related coronavirus	Hu/Qasim-KSA-893/2016	membrane

MG011352	28544-29785	Middle East respiratory syndrome-related coronavirus	Hu/Qasim-KSA-893/2016	nucleocapsid

MG011352	257-21492	Middle East respiratory syndrome-related coronavirus	Hu/Qasim-KSA-893/2016	orf1ab

MG011352	21434-25495	Middle East respiratory syndrome-related coronavirus	Hu/Qasim-KSA-893/2016	spike

MG757593	27831-28490	Middle East respiratory syndrome-related coronavirus	Hu/Riyadh-KSA-002D1N/2015	membrane

MG757593	28544-29785	Middle East respiratory syndrome-related coronavirus	Hu/Riyadh-KSA-002D1N/2015	nucleocapsid

MG757593	257-21492	Middle East respiratory syndrome-related coronavirus	Hu/Riyadh-KSA-002D1N/2015	orf1ab

MG757593	21434-25495	Middle East respiratory syndrome-related coronavirus	Hu/Riyadh-KSA-002D1N/2015	spike

MG757594	27831-28490	Middle East respiratory syndrome-related coronavirus	Hu/Riyadh-KSA-004D1S/2015	membrane

MG757594	28544-29785	Middle East respiratory syndrome-related coronavirus	Hu/Riyadh-KSA-004D1S/2015	nucleocapsid

MG757594	257-21492	Middle East respiratory syndrome-related coronavirus	Hu/Riyadh-KSA-004D1S/2015	orf1ab

MG757594	21434-25495	Middle East respiratory syndrome-related coronavirus	Hu/Riyadh-KSA-004D1S/2015	spike

MG757595	27831-28490	Middle East respiratory syndrome-related coronavirus	Hu/Riyadh-KSA-005D1S/2015	membrane

MG757595	28544-29785	Middle East respiratory syndrome-related coronavirus	Hu/Riyadh-KSA-005D1S/2015	nucleocapsid

MG757595	257-21492	Middle East respiratory syndrome-related coronavirus	Hu/Riyadh-KSA-005D1S/2015	orf1ab

MG757595	21434-25495	Middle East respiratory syndrome-related coronavirus	Hu/Riyadh-KSA-005D1S/2015	spike

MG757596	27831-28490	Middle East respiratory syndrome-related coronavirus	Hu/Riyadh-KSA-006D1S/2015	membrane

MG757596	28544-29785	Middle East respiratory syndrome-related coronavirus	Hu/Riyadh-KSA-006D1S/2015	nucleocapsid

MG757596	257-21492	Middle East respiratory syndrome-related coronavirus	Hu/Riyadh-KSA-006D1S/2015	orf1ab

MG757596	21434-25495	Middle East respiratory syndrome-related coronavirus	Hu/Riyadh-KSA-006D1S/2015	spike

MG757597	27831-28490	Middle East respiratory syndrome-related coronavirus	Hu/Riyadh-KSA-007D1N/2015	membrane

MG757597	28544-29785	Middle East respiratory syndrome-related coronavirus	Hu/Riyadh-KSA-007D1N/2015	nucleocapsid

MG757597	257-21492	Middle East respiratory syndrome-related coronavirus	Hu/Riyadh-KSA-007D1N/2015	orf1ab

MG757597	21434-25495	Middle East respiratory syndrome-related coronavirus	Hu/Riyadh-KSA-007D1N/2015	spike

MG757598	27831-28490	Middle East respiratory syndrome-related coronavirus	Hu/Riyadh-KSA-008D3N/2015	membrane

MG757598	28544-29785	Middle East respiratory syndrome-related coronavirus	Hu/Riyadh-KSA-008D3N/2015	nucleocapsid

MG757598	257-21492	Middle East respiratory syndrome-related coronavirus	Hu/Riyadh-KSA-008D3N/2015	orf1ab

MG757598	21434-25495	Middle East respiratory syndrome-related coronavirus	Hu/Riyadh-KSA-008D3N/2015	spike

MH029552	279-21514	Middle East respiratory syndrome-related coronavirus	Hu/Riyadh-KSA-008D3N/2015	orf1ab

MH029552	27853-28512	Middle East respiratory syndrome-related coronavirus	Hu/Riyadh-KSA-008D3N/2015	membrane

MH029552	28566-29807	Middle East respiratory syndrome-related coronavirus	Hu/Riyadh-KSA-008D3N/2015	nucleocapsid

MH029552	21456-25517	Middle East respiratory syndrome-related coronavirus	Hu/Riyadh-KSA-008D3N/2015	spike

MG757599	27831-28490	Middle East respiratory syndrome-related coronavirus	Hu/Riyadh-KSA-009D1N/2015	membrane

MG757599	28544-29785	Middle East respiratory syndrome-related coronavirus	Hu/Riyadh-KSA-009D1N/2015	nucleocapsid

MG757599	257-21492	Middle East respiratory syndrome-related coronavirus	Hu/Riyadh-KSA-009D1N/2015	orf1ab

MG757599	21434-25495	Middle East respiratory syndrome-related coronavirus	Hu/Riyadh-KSA-009D1N/2015	spike

MG757600	27831-28490	Middle East respiratory syndrome-related coronavirus	Hu/Riyadh-KSA-012D1S/2015	membrane

MG757600	28544-29785	Middle East respiratory syndrome-related coronavirus	Hu/Riyadh-KSA-012D1S/2015	nucleocapsid

MG757600	257-21492	Middle East respiratory syndrome-related coronavirus	Hu/Riyadh-KSA-012D1S/2015	orf1ab

MG757600	21434-25495	Middle East respiratory syndrome-related coronavirus	Hu/Riyadh-KSA-012D1S/2015	spike

MG757601	27831-28490	Middle East respiratory syndrome-related coronavirus	Hu/Riyadh-KSA-014D1N/2015	membrane

MG757601	28544-29785	Middle East respiratory syndrome-related coronavirus	Hu/Riyadh-KSA-014D1N/2015	nucleocapsid

MG757601	257-21492	Middle East respiratory syndrome-related coronavirus	Hu/Riyadh-KSA-014D1N/2015	orf1ab

MG757601	21434-25495	Middle East respiratory syndrome-related coronavirus	Hu/Riyadh-KSA-014D1N/2015	spike

MG757602	27831-28490	Middle East respiratory syndrome-related coronavirus	Hu/Riyadh-KSA-015D1P/2015	membrane

MG757602	28544-29785	Middle East respiratory syndrome-related coronavirus	Hu/Riyadh-KSA-015D1P/2015	nucleocapsid

MG757602	257-21492	Middle East respiratory syndrome-related coronavirus	Hu/Riyadh-KSA-015D1P/2015	orf1ab

MG757602	21434-25495	Middle East respiratory syndrome-related coronavirus	Hu/Riyadh-KSA-015D1P/2015	spike

MG520075	27831-28490	Middle East respiratory syndrome-related coronavirus	Hu/Riyadh-KSA-023D1N/2015	membrane

MG520075	28544-29785	Middle East respiratory syndrome-related coronavirus	Hu/Riyadh-KSA-023D1N/2015	nucleocapsid

MG520075	257-21492	Middle East respiratory syndrome-related coronavirus	Hu/Riyadh-KSA-023D1N/2015	orf1ab

MG520075	21434-25495	Middle East respiratory syndrome-related coronavirus	Hu/Riyadh-KSA-023D1N/2015	spike

MG546330	279-21514	Middle East respiratory syndrome-related coronavirus	Hu/Riyadh-KSA-023D1N/2015	orf1ab

MG546330	27853-28512	Middle East respiratory syndrome-related coronavirus	Hu/Riyadh-KSA-023D1N/2015	membrane

MG546330	28566-29807	Middle East respiratory syndrome-related coronavirus	Hu/Riyadh-KSA-023D1N/2015	nucleocapsid

MG546330	21456-25517	Middle East respiratory syndrome-related coronavirus	Hu/Riyadh-KSA-023D1N/2015	spike

MG546331	279-21514	Middle East respiratory syndrome-related coronavirus	Hu/Riyadh-KSA-023D3N/2015	orf1ab

MG546331	27853-28512	Middle East respiratory syndrome-related coronavirus	Hu/Riyadh-KSA-023D3N/2015	membrane

MG546331	28566-29807	Middle East respiratory syndrome-related coronavirus	Hu/Riyadh-KSA-023D3N/2015	nucleocapsid

MG546331	21456-25517	Middle East respiratory syndrome-related coronavirus	Hu/Riyadh-KSA-023D3N/2015	spike

MG757603	27831-28490	Middle East respiratory syndrome-related coronavirus	Hu/Riyadh-KSA-031D1N/2015	membrane

MG757603	28544-29785	Middle East respiratory syndrome-related coronavirus	Hu/Riyadh-KSA-031D1N/2015	nucleocapsid

MG757603	257-21492	Middle East respiratory syndrome-related coronavirus	Hu/Riyadh-KSA-031D1N/2015	orf1ab

MG757603	21434-25495	Middle East respiratory syndrome-related coronavirus	Hu/Riyadh-KSA-031D1N/2015	spike

MG757605	27831-28490	Middle East respiratory syndrome-related coronavirus	Hu/Riyadh-KSA-036D1N/2016	membrane

MG757605	28544-29785	Middle East respiratory syndrome-related coronavirus	Hu/Riyadh-KSA-036D1N/2016	nucleocapsid

MG757605	257-21492	Middle East respiratory syndrome-related coronavirus	Hu/Riyadh-KSA-036D1N/2016	orf1ab

MG757605	21434-25495	Middle East respiratory syndrome-related coronavirus	Hu/Riyadh-KSA-036D1N/2016	spike

MG757604	27831-28490	Middle East respiratory syndrome-related coronavirus	Hu/Riyadh-KSA-039D3T/2016	membrane

MG757604	28544-29785	Middle East respiratory syndrome-related coronavirus	Hu/Riyadh-KSA-039D3T/2016	nucleocapsid

MG757604	257-21492	Middle East respiratory syndrome-related coronavirus	Hu/Riyadh-KSA-039D3T/2016	orf1ab

MG757604	21434-25495	Middle East respiratory syndrome-related coronavirus	Hu/Riyadh-KSA-039D3T/2016	spike

MG366883	279-21514	Middle East respiratory syndrome-related coronavirus	Hu/Riyadh-KSA-10024/2017	orf1ab

MG366883	27853-28512	Middle East respiratory syndrome-related coronavirus	Hu/Riyadh-KSA-10024/2017	membrane

MG366883	28566-29807	Middle East respiratory syndrome-related coronavirus	Hu/Riyadh-KSA-10024/2017	nucleocapsid

MG366883	21456-25517	Middle East respiratory syndrome-related coronavirus	Hu/Riyadh-KSA-10024/2017	spike

KX154689	27831-28490	Middle East respiratory syndrome-related coronavirus	Hu/Riyadh-KSA-10208/2016	membrane

KX154689	28544-29785	Middle East respiratory syndrome-related coronavirus	Hu/Riyadh-KSA-10208/2016	nucleocapsid

KX154689	257-21492	Middle East respiratory syndrome-related coronavirus	Hu/Riyadh-KSA-10208/2016	orf1ab

KX154689	21434-25495	Middle East respiratory syndrome-related coronavirus	Hu/Riyadh-KSA-10208/2016	spike

MG912601	279-21514	Middle East respiratory syndrome-related coronavirus	Hu/Riyadh-KSA-10308/2017	orf1ab

MG912601	27853-28512	Middle East respiratory syndrome-related coronavirus	Hu/Riyadh-KSA-10308/2017	membrane

MG912601	28566-29807	Middle East respiratory syndrome-related coronavirus	Hu/Riyadh-KSA-10308/2017	nucleocapsid

MG912601	21456-25517	Middle East respiratory syndrome-related coronavirus	Hu/Riyadh-KSA-10308/2017	spike

MH310911	27831-28490	Middle East respiratory syndrome-related coronavirus	Hu/Riyadh-KSA-10717/2017	membrane

MH310911	28544-29785	Middle East respiratory syndrome-related coronavirus	Hu/Riyadh-KSA-10717/2017	nucleocapsid

MH310911	257-21492	Middle East respiratory syndrome-related coronavirus	Hu/Riyadh-KSA-10717/2017	orf1ab

MH310911	21434-25495	Middle East respiratory syndrome-related coronavirus	Hu/Riyadh-KSA-10717/2017	spike

KX154684	27831-28490	Middle East respiratory syndrome-related coronavirus	Hu/Riyadh-KSA-11739/2016	membrane

KX154684	28544-29785	Middle East respiratory syndrome-related coronavirus	Hu/Riyadh-KSA-11739/2016	nucleocapsid

KX154684	257-21492	Middle East respiratory syndrome-related coronavirus	Hu/Riyadh-KSA-11739/2016	orf1ab

KX154684	21434-25495	Middle East respiratory syndrome-related coronavirus	Hu/Riyadh-KSA-11739/2016	spike

KX154685	27831-28490	Middle East respiratory syndrome-related coronavirus	Hu/Riyadh-KSA-11740/2016	membrane

KX154685	28544-29785	Middle East respiratory syndrome-related coronavirus	Hu/Riyadh-KSA-11740/2016	nucleocapsid

KX154685	257-21492	Middle East respiratory syndrome-related coronavirus	Hu/Riyadh-KSA-11740/2016	orf1ab

KX154685	21434-25495	Middle East respiratory syndrome-related coronavirus	Hu/Riyadh-KSA-11740/2016	spike

KX154686	27831-28490	Middle East respiratory syndrome-related coronavirus	Hu/Riyadh-KSA-11898/2016	membrane

KX154686	28544-29785	Middle East respiratory syndrome-related coronavirus	Hu/Riyadh-KSA-11898/2016	nucleocapsid

KX154686	257-21492	Middle East respiratory syndrome-related coronavirus	Hu/Riyadh-KSA-11898/2016	orf1ab

KX154686	21434-25495	Middle East respiratory syndrome-related coronavirus	Hu/Riyadh-KSA-11898/2016	spike

KX154687	27831-28490	Middle East respiratory syndrome-related coronavirus	Hu/Riyadh-KSA-11958/2016	membrane

KX154687	28544-29785	Middle East respiratory syndrome-related coronavirus	Hu/Riyadh-KSA-11958/2016	nucleocapsid

KX154687	257-21492	Middle East respiratory syndrome-related coronavirus	Hu/Riyadh-KSA-11958/2016	orf1ab

KX154687	21434-25495	Middle East respiratory syndrome-related coronavirus	Hu/Riyadh-KSA-11958/2016	spike

KX154688	27831-28490	Middle East respiratory syndrome-related coronavirus	Hu/Riyadh-KSA-12160/2016	membrane

KX154688	28544-29785	Middle East respiratory syndrome-related coronavirus	Hu/Riyadh-KSA-12160/2016	nucleocapsid

KX154688	257-21492	Middle East respiratory syndrome-related coronavirus	Hu/Riyadh-KSA-12160/2016	orf1ab

KX154688	21434-25495	Middle East respiratory syndrome-related coronavirus	Hu/Riyadh-KSA-12160/2016	spike

KX154691	27831-28490	Middle East respiratory syndrome-related coronavirus	Hu/Riyadh-KSA-12832/2016	membrane

KX154691	28544-29785	Middle East respiratory syndrome-related coronavirus	Hu/Riyadh-KSA-12832/2016	nucleocapsid

KX154691	257-21492	Middle East respiratory syndrome-related coronavirus	Hu/Riyadh-KSA-12832/2016	orf1ab

KX154691	21434-25495	Middle East respiratory syndrome-related coronavirus	Hu/Riyadh-KSA-12832/2016	spike

KX154692	27831-28490	Middle East respiratory syndrome-related coronavirus	Hu/Riyadh-KSA-12969/2016	membrane

KX154692	28544-29785	Middle East respiratory syndrome-related coronavirus	Hu/Riyadh-KSA-12969/2016	nucleocapsid

KX154692	257-21492	Middle East respiratory syndrome-related coronavirus	Hu/Riyadh-KSA-12969/2016	orf1ab

KX154692	21434-25495	Middle East respiratory syndrome-related coronavirus	Hu/Riyadh-KSA-12969/2016	spike

KX154693	27831-28490	Middle East respiratory syndrome-related coronavirus	Hu/Riyadh-KSA-13127/2016	membrane

KX154693	28544-29785	Middle East respiratory syndrome-related coronavirus	Hu/Riyadh-KSA-13127/2016	nucleocapsid

KX154693	257-21492	Middle East respiratory syndrome-related coronavirus	Hu/Riyadh-KSA-13127/2016	orf1ab

KX154693	21434-25495	Middle East respiratory syndrome-related coronavirus	Hu/Riyadh-KSA-13127/2016	spike

MG011340	27831-28490	Middle East respiratory syndrome-related coronavirus	Hu/Riyadh-KSA-13798/2016	membrane

MG011340	28544-29785	Middle East respiratory syndrome-related coronavirus	Hu/Riyadh-KSA-13798/2016	nucleocapsid

MG011340	257-21492	Middle East respiratory syndrome-related coronavirus	Hu/Riyadh-KSA-13798/2016	orf1ab

MG011340	21434-25495	Middle East respiratory syndrome-related coronavirus	Hu/Riyadh-KSA-13798/2016	spike

MG011342	27831-28490	Middle East respiratory syndrome-related coronavirus	Hu/Riyadh-KSA-13984/2016	membrane

MG011342	28544-29785	Middle East respiratory syndrome-related coronavirus	Hu/Riyadh-KSA-13984/2016	nucleocapsid

MG011342	257-21492	Middle East respiratory syndrome-related coronavirus	Hu/Riyadh-KSA-13984/2016	orf1ab

MG011342	21434-25495	Middle East respiratory syndrome-related coronavirus	Hu/Riyadh-KSA-13984/2016	spike

MG011343	27831-28490	Middle East respiratory syndrome-related coronavirus	Hu/Riyadh-KSA-14607/2016	membrane

MG011343	28544-29785	Middle East respiratory syndrome-related coronavirus	Hu/Riyadh-KSA-14607/2016	nucleocapsid

MG011343	257-21492	Middle East respiratory syndrome-related coronavirus	Hu/Riyadh-KSA-14607/2016	orf1ab

MG011343	21434-25495	Middle East respiratory syndrome-related coronavirus	Hu/Riyadh-KSA-14607/2016	spike

MG011347	27831-28490	Middle East respiratory syndrome-related coronavirus	Hu/Riyadh-KSA-14670/2016	membrane

MG011347	28544-29785	Middle East respiratory syndrome-related coronavirus	Hu/Riyadh-KSA-14670/2016	nucleocapsid

MG011347	257-21492	Middle East respiratory syndrome-related coronavirus	Hu/Riyadh-KSA-14670/2016	orf1ab

MG011347	21434-25495	Middle East respiratory syndrome-related coronavirus	Hu/Riyadh-KSA-14670/2016	spike

MG011344	27831-28490	Middle East respiratory syndrome-related coronavirus	Hu/Riyadh-KSA-14675/2016	membrane

MG011344	28544-29785	Middle East respiratory syndrome-related coronavirus	Hu/Riyadh-KSA-14675/2016	nucleocapsid

MG011344	257-21492	Middle East respiratory syndrome-related coronavirus	Hu/Riyadh-KSA-14675/2016	orf1ab

MG011344	21434-25495	Middle East respiratory syndrome-related coronavirus	Hu/Riyadh-KSA-14675/2016	spike

MG011346	27831-28490	Middle East respiratory syndrome-related coronavirus	Hu/Riyadh-KSA-14949/2016	membrane

MG011346	28544-29785	Middle East respiratory syndrome-related coronavirus	Hu/Riyadh-KSA-14949/2016	nucleocapsid

MG011346	257-21492	Middle East respiratory syndrome-related coronavirus	Hu/Riyadh-KSA-14949/2016	orf1ab

MG011346	21434-25495	Middle East respiratory syndrome-related coronavirus	Hu/Riyadh-KSA-14949/2016	spike

MG011345	27831-28490	Middle East respiratory syndrome-related coronavirus	Hu/Riyadh-KSA-15385/2016	membrane

MG011345	28544-29785	Middle East respiratory syndrome-related coronavirus	Hu/Riyadh-KSA-15385/2016	nucleocapsid

MG011345	257-21492	Middle East respiratory syndrome-related coronavirus	Hu/Riyadh-KSA-15385/2016	orf1ab

MG011345	21434-25495	Middle East respiratory syndrome-related coronavirus	Hu/Riyadh-KSA-15385/2016	spike

MG011359	27831-28490	Middle East respiratory syndrome-related coronavirus	Hu/Riyadh-KSA-16559/2016	membrane

MG011359	28544-29785	Middle East respiratory syndrome-related coronavirus	Hu/Riyadh-KSA-16559/2016	nucleocapsid

MG011359	257-21492	Middle East respiratory syndrome-related coronavirus	Hu/Riyadh-KSA-16559/2016	orf1ab

MG011359	21434-25495	Middle East respiratory syndrome-related coronavirus	Hu/Riyadh-KSA-16559/2016	spike

MG011350	27831-28490	Middle East respiratory syndrome-related coronavirus	Hu/Riyadh-KSA-16849/2016	membrane

MG011350	28544-29785	Middle East respiratory syndrome-related coronavirus	Hu/Riyadh-KSA-16849/2016	nucleocapsid

MG011350	257-21492	Middle East respiratory syndrome-related coronavirus	Hu/Riyadh-KSA-16849/2016	orf1ab

MG011350	21434-25495	Middle East respiratory syndrome-related coronavirus	Hu/Riyadh-KSA-16849/2016	spike

MG011349	27831-28490	Middle East respiratory syndrome-related coronavirus	Hu/Riyadh-KSA-17382/2016	membrane

MG011349	28544-29785	Middle East respiratory syndrome-related coronavirus	Hu/Riyadh-KSA-17382/2016	nucleocapsid

MG011349	257-21492	Middle East respiratory syndrome-related coronavirus	Hu/Riyadh-KSA-17382/2016	orf1ab

MG011349	21434-25495	Middle East respiratory syndrome-related coronavirus	Hu/Riyadh-KSA-17382/2016	spike

MG011348	27831-28490	Middle East respiratory syndrome-related coronavirus	Hu/Riyadh-KSA-17756/2016	membrane

MG011348	28544-29785	Middle East respiratory syndrome-related coronavirus	Hu/Riyadh-KSA-17756/2016	nucleocapsid

MG011348	257-21492	Middle East respiratory syndrome-related coronavirus	Hu/Riyadh-KSA-17756/2016	orf1ab

MG011348	21434-25495	Middle East respiratory syndrome-related coronavirus	Hu/Riyadh-KSA-17756/2016	spike

MK462249	27853-28512	Middle East respiratory syndrome-related coronavirus	Hu/Riyadh-KSA-18012493/2018	membrane

MK462249	28566-29807	Middle East respiratory syndrome-related coronavirus	Hu/Riyadh-KSA-18012493/2018	nucleocapsid

MK462249	279-21514	Middle East respiratory syndrome-related coronavirus	Hu/Riyadh-KSA-18012493/2018	orf1ab

MK462249	21456-25517	Middle East respiratory syndrome-related coronavirus	Hu/Riyadh-KSA-18012493/2018	spike

MK462253	27853-28512	Middle East respiratory syndrome-related coronavirus	Hu/Riyadh-KSA-18013832/2018	membrane

MK462253	28566-29807	Middle East respiratory syndrome-related coronavirus	Hu/Riyadh-KSA-18013832/2018	nucleocapsid

MK462253	279-21514	Middle East respiratory syndrome-related coronavirus	Hu/Riyadh-KSA-18013832/2018	orf1ab

MK462253	21456-25517	Middle East respiratory syndrome-related coronavirus	Hu/Riyadh-KSA-18013832/2018	spike

MK462255	27787-28446	Middle East respiratory syndrome-related coronavirus	Hu/Riyadh-KSA-18014504/2018	membrane

MK462255	28500-29741	Middle East respiratory syndrome-related coronavirus	Hu/Riyadh-KSA-18014504/2018	nucleocapsid

MK462255	279-21488	Middle East respiratory syndrome-related coronavirus	Hu/Riyadh-KSA-18014504/2018	orf1ab

MK462255	21390-25451	Middle East respiratory syndrome-related coronavirus	Hu/Riyadh-KSA-18014504/2018	spike

MK462256	27853-28512	Middle East respiratory syndrome-related coronavirus	Hu/Riyadh-KSA-18014506/2018	membrane

MK462256	28566-29807	Middle East respiratory syndrome-related coronavirus	Hu/Riyadh-KSA-18014506/2018	nucleocapsid

MK462256	279-21514	Middle East respiratory syndrome-related coronavirus	Hu/Riyadh-KSA-18014506/2018	orf1ab

MK462256	21456-25517	Middle East respiratory syndrome-related coronavirus	Hu/Riyadh-KSA-18014506/2018	spike

MG011353	27831-28490	Middle East respiratory syndrome-related coronavirus	Hu/Riyadh-KSA-19949/2016	membrane

MG011353	28544-29785	Middle East respiratory syndrome-related coronavirus	Hu/Riyadh-KSA-19949/2016	nucleocapsid

MG011353	257-21492	Middle East respiratory syndrome-related coronavirus	Hu/Riyadh-KSA-19949/2016	orf1ab

MG011353	21434-25495	Middle East respiratory syndrome-related coronavirus	Hu/Riyadh-KSA-19949/2016	spike

MG011354	27831-28490	Middle East respiratory syndrome-related coronavirus	Hu/Riyadh-KSA-20570/2016	membrane

MG011354	28544-29785	Middle East respiratory syndrome-related coronavirus	Hu/Riyadh-KSA-20570/2016	nucleocapsid

MG011354	257-21492	Middle East respiratory syndrome-related coronavirus	Hu/Riyadh-KSA-20570/2016	orf1ab

MG011354	21434-25495	Middle East respiratory syndrome-related coronavirus	Hu/Riyadh-KSA-20570/2016	spike

MG011355	27831-28490	Middle East respiratory syndrome-related coronavirus	Hu/Riyadh-KSA-21155/2016	membrane

MG011355	28544-29785	Middle East respiratory syndrome-related coronavirus	Hu/Riyadh-KSA-21155/2016	nucleocapsid

MG011355	257-21492	Middle East respiratory syndrome-related coronavirus	Hu/Riyadh-KSA-21155/2016	orf1ab

MG011355	21434-25495	Middle East respiratory syndrome-related coronavirus	Hu/Riyadh-KSA-21155/2016	spike

MG011356	27831-28490	Middle East respiratory syndrome-related coronavirus	Hu/Riyadh-KSA-21891/2016	membrane

MG011356	28544-29785	Middle East respiratory syndrome-related coronavirus	Hu/Riyadh-KSA-21891/2016	nucleocapsid

MG011356	257-21492	Middle East respiratory syndrome-related coronavirus	Hu/Riyadh-KSA-21891/2016	orf1ab

MG011356	21434-25495	Middle East respiratory syndrome-related coronavirus	Hu/Riyadh-KSA-21891/2016	spike

MG011358	27831-28490	Middle East respiratory syndrome-related coronavirus	Hu/Riyadh-KSA-24241/2016	membrane

MG011358	28544-29785	Middle East respiratory syndrome-related coronavirus	Hu/Riyadh-KSA-24241/2016	nucleocapsid

MG011358	257-21492	Middle East respiratory syndrome-related coronavirus	Hu/Riyadh-KSA-24241/2016	orf1ab

MG011358	21434-25495	Middle East respiratory syndrome-related coronavirus	Hu/Riyadh-KSA-24241/2016	spike

MG011357	27831-28490	Middle East respiratory syndrome-related coronavirus	Hu/Riyadh-KSA-24244/2016	membrane

MG011357	28544-29785	Middle East respiratory syndrome-related coronavirus	Hu/Riyadh-KSA-24244/2016	nucleocapsid

MG011357	257-21492	Middle East respiratory syndrome-related coronavirus	Hu/Riyadh-KSA-24244/2016	orf1ab

MG011357	21434-25495	Middle East respiratory syndrome-related coronavirus	Hu/Riyadh-KSA-24244/2016	spike

MH310909	27800-28459	Middle East respiratory syndrome-related coronavirus	Hu/Riyadh-KSA-5767/2017	membrane

MH310909	28513-29754	Middle East respiratory syndrome-related coronavirus	Hu/Riyadh-KSA-5767/2017	nucleocapsid

MH310909	257-21492	Middle East respiratory syndrome-related coronavirus	Hu/Riyadh-KSA-5767/2017	orf1ab

MH310909	21434-25495	Middle East respiratory syndrome-related coronavirus	Hu/Riyadh-KSA-5767/2017	spike

MG912595	279-21514	Middle East respiratory syndrome-related coronavirus	Hu/Riyadh-KSA-7178/2017	orf1ab

MG912595	27853-28512	Middle East respiratory syndrome-related coronavirus	Hu/Riyadh-KSA-7178/2017	membrane

MG912595	28566-29807	Middle East respiratory syndrome-related coronavirus	Hu/Riyadh-KSA-7178/2017	nucleocapsid

MG912595	21456-25517	Middle East respiratory syndrome-related coronavirus	Hu/Riyadh-KSA-7178/2017	spike

MH310912	27820-28479	Middle East respiratory syndrome-related coronavirus	Hu/Riyadh-KSA-7344/2017	membrane

MH310912	28533-29774	Middle East respiratory syndrome-related coronavirus	Hu/Riyadh-KSA-7344/2017	nucleocapsid

MH310912	246-21481	Middle East respiratory syndrome-related coronavirus	Hu/Riyadh-KSA-7344/2017	orf1ab

MH310912	21423-25484	Middle East respiratory syndrome-related coronavirus	Hu/Riyadh-KSA-7344/2017	spike

MG912596	279-21514	Middle East respiratory syndrome-related coronavirus	Hu/Riyadh-KSA-7373/2017	orf1ab

MG912596	27853-28512	Middle East respiratory syndrome-related coronavirus	Hu/Riyadh-KSA-7373/2017	membrane

MG912596	28566-29807	Middle East respiratory syndrome-related coronavirus	Hu/Riyadh-KSA-7373/2017	nucleocapsid

MG912596	21456-25517	Middle East respiratory syndrome-related coronavirus	Hu/Riyadh-KSA-7373/2017	spike

MG366483	27831-28490	Middle East respiratory syndrome-related coronavirus	Hu/Riyadh-KSA-7413/2017	membrane

MG366483	28544-29785	Middle East respiratory syndrome-related coronavirus	Hu/Riyadh-KSA-7413/2017	nucleocapsid

MG366483	257-21492	Middle East respiratory syndrome-related coronavirus	Hu/Riyadh-KSA-7413/2017	orf1ab

MG366483	21434-25495	Middle East respiratory syndrome-related coronavirus	Hu/Riyadh-KSA-7413/2017	spike

MG912597	279-21514	Middle East respiratory syndrome-related coronavirus	Hu/Riyadh-KSA-7423/2017	orf1ab

MG912597	27853-28512	Middle East respiratory syndrome-related coronavirus	Hu/Riyadh-KSA-7423/2017	membrane

MG912597	28566-29807	Middle East respiratory syndrome-related coronavirus	Hu/Riyadh-KSA-7423/2017	nucleocapsid

MG912597	21456-25517	Middle East respiratory syndrome-related coronavirus	Hu/Riyadh-KSA-7423/2017	spike

MG366880	279-21514	Middle East respiratory syndrome-related coronavirus	Hu/Riyadh-KSA-7436/2017	orf1ab

MG366880	27853-28512	Middle East respiratory syndrome-related coronavirus	Hu/Riyadh-KSA-7436/2017	membrane

MG366880	28566-29807	Middle East respiratory syndrome-related coronavirus	Hu/Riyadh-KSA-7436/2017	nucleocapsid

MG366880	21456-25517	Middle East respiratory syndrome-related coronavirus	Hu/Riyadh-KSA-7436/2017	spike

MG912598	279-21514	Middle East respiratory syndrome-related coronavirus	Hu/Riyadh-KSA-7680/2017	orf1ab

MG912598	27853-28512	Middle East respiratory syndrome-related coronavirus	Hu/Riyadh-KSA-7680/2017	membrane

MG912598	28566-29807	Middle East respiratory syndrome-related coronavirus	Hu/Riyadh-KSA-7680/2017	nucleocapsid

MG912598	21456-25517	Middle East respiratory syndrome-related coronavirus	Hu/Riyadh-KSA-7680/2017	spike

MH310910	27831-28490	Middle East respiratory syndrome-related coronavirus	Hu/Riyadh-KSA-7696/2017	membrane

MH310910	28544-29785	Middle East respiratory syndrome-related coronavirus	Hu/Riyadh-KSA-7696/2017	nucleocapsid

MH310910	257-21492	Middle East respiratory syndrome-related coronavirus	Hu/Riyadh-KSA-7696/2017	orf1ab

MH310910	21434-25495	Middle East respiratory syndrome-related coronavirus	Hu/Riyadh-KSA-7696/2017	spike

MG912599	279-21514	Middle East respiratory syndrome-related coronavirus	Hu/Riyadh-KSA-8667/2017	orf1ab

MG912599	27853-28512	Middle East respiratory syndrome-related coronavirus	Hu/Riyadh-KSA-8667/2017	membrane

MG912599	28566-29807	Middle East respiratory syndrome-related coronavirus	Hu/Riyadh-KSA-8667/2017	nucleocapsid

MG912599	21456-25517	Middle East respiratory syndrome-related coronavirus	Hu/Riyadh-KSA-8667/2017	spike

MG366881	279-21514	Middle East respiratory syndrome-related coronavirus	Hu/Riyadh-KSA-8677/2017	orf1ab

MG366881	27853-28512	Middle East respiratory syndrome-related coronavirus	Hu/Riyadh-KSA-8677/2017	membrane

MG366881	28566-29807	Middle East respiratory syndrome-related coronavirus	Hu/Riyadh-KSA-8677/2017	nucleocapsid

MG366881	21456-25517	Middle East respiratory syndrome-related coronavirus	Hu/Riyadh-KSA-8677/2017	spike

MG366882	279-21514	Middle East respiratory syndrome-related coronavirus	Hu/Riyadh-KSA-8683/2017	orf1ab

MG366882	27853-28512	Middle East respiratory syndrome-related coronavirus	Hu/Riyadh-KSA-8683/2017	membrane

MG366882	28566-29807	Middle East respiratory syndrome-related coronavirus	Hu/Riyadh-KSA-8683/2017	nucleocapsid

MG366882	21456-25517	Middle East respiratory syndrome-related coronavirus	Hu/Riyadh-KSA-8683/2017	spike

MG912600	279-21514	Middle East respiratory syndrome-related coronavirus	Hu/Riyadh-KSA-8882/2017	orf1ab

MG912600	27853-28512	Middle East respiratory syndrome-related coronavirus	Hu/Riyadh-KSA-8882/2017	membrane

MG912600	28566-29807	Middle East respiratory syndrome-related coronavirus	Hu/Riyadh-KSA-8882/2017	nucleocapsid

MG912600	21456-25517	Middle East respiratory syndrome-related coronavirus	Hu/Riyadh-KSA-8882/2017	spike

MG912607	279-21514	Middle East respiratory syndrome-related coronavirus	Hu/Riyadh-KSA-9522/2017	orf1ab

MG912607	27853-28512	Middle East respiratory syndrome-related coronavirus	Hu/Riyadh-KSA-9522/2017	membrane

MG912607	28566-29807	Middle East respiratory syndrome-related coronavirus	Hu/Riyadh-KSA-9522/2017	nucleocapsid

MG912607	21456-25517	Middle East respiratory syndrome-related coronavirus	Hu/Riyadh-KSA-9522/2017	spike

MG912602	279-21514	Middle East respiratory syndrome-related coronavirus	Hu/Riyadh-KSA-9614/2017	orf1ab

MG912602	27853-28512	Middle East respiratory syndrome-related coronavirus	Hu/Riyadh-KSA-9614/2017	membrane

MG912602	28566-29807	Middle East respiratory syndrome-related coronavirus	Hu/Riyadh-KSA-9614/2017	nucleocapsid

MG912602	21456-25517	Middle East respiratory syndrome-related coronavirus	Hu/Riyadh-KSA-9614/2017	spike

MG912604	279-21514	Middle East respiratory syndrome-related coronavirus	Hu/Riyadh-KSA-9689/2017	orf1ab

MG912604	27853-28512	Middle East respiratory syndrome-related coronavirus	Hu/Riyadh-KSA-9689/2017	membrane

MG912604	28566-29807	Middle East respiratory syndrome-related coronavirus	Hu/Riyadh-KSA-9689/2017	nucleocapsid

MG912604	21456-25517	Middle East respiratory syndrome-related coronavirus	Hu/Riyadh-KSA-9689/2017	spike

MG912603	279-21514	Middle East respiratory syndrome-related coronavirus	Hu/Riyadh-KSA-9693/2017	orf1ab

MG912603	27853-28512	Middle East respiratory syndrome-related coronavirus	Hu/Riyadh-KSA-9693/2017	membrane

MG912603	28566-29807	Middle East respiratory syndrome-related coronavirus	Hu/Riyadh-KSA-9693/2017	nucleocapsid

MG912603	21456-25517	Middle East respiratory syndrome-related coronavirus	Hu/Riyadh-KSA-9693/2017	spike

MG912608	279-21514	Middle East respiratory syndrome-related coronavirus	Hu/Riyadh-KSA-9730/2017	orf1ab

MG912608	27853-28512	Middle East respiratory syndrome-related coronavirus	Hu/Riyadh-KSA-9730/2017	membrane

MG912608	28566-29807	Middle East respiratory syndrome-related coronavirus	Hu/Riyadh-KSA-9730/2017	nucleocapsid

MG912608	21456-25517	Middle East respiratory syndrome-related coronavirus	Hu/Riyadh-KSA-9730/2017	spike

MG912605	279-21514	Middle East respiratory syndrome-related coronavirus	Hu/Riyadh-KSA-9835/2017	orf1ab

MG912605	27853-28512	Middle East respiratory syndrome-related coronavirus	Hu/Riyadh-KSA-9835/2017	membrane

MG912605	28566-29807	Middle East respiratory syndrome-related coronavirus	Hu/Riyadh-KSA-9835/2017	nucleocapsid

MG912605	21456-25517	Middle East respiratory syndrome-related coronavirus	Hu/Riyadh-KSA-9835/2017	spike

MG912606	279-21514	Middle East respiratory syndrome-related coronavirus	Hu/Riyadh-KSA-9852/2017	orf1ab

MG912606	27853-28512	Middle East respiratory syndrome-related coronavirus	Hu/Riyadh-KSA-9852/2017	membrane

MG912606	28566-29807	Middle East respiratory syndrome-related coronavirus	Hu/Riyadh-KSA-9852/2017	nucleocapsid

MG912606	21456-25517	Middle East respiratory syndrome-related coronavirus	Hu/Riyadh-KSA-9852/2017	spike

MG011361	27831-28490	Middle East respiratory syndrome-related coronavirus	Hu/Riyadh-KSA-K17000405/2017	membrane

MG011361	28544-29785	Middle East respiratory syndrome-related coronavirus	Hu/Riyadh-KSA-K17000405/2017	nucleocapsid

MG011361	257-21492	Middle East respiratory syndrome-related coronavirus	Hu/Riyadh-KSA-K17000405/2017	orf1ab

MG011361	21434-25495	Middle East respiratory syndrome-related coronavirus	Hu/Riyadh-KSA-K17000405/2017	spike

MG011360	27831-28490	Middle East respiratory syndrome-related coronavirus	Hu/Riyadh-KSA-K17000887/2017	membrane

MG011360	28544-29785	Middle East respiratory syndrome-related coronavirus	Hu/Riyadh-KSA-K17000887/2017	nucleocapsid

MG011360	257-21492	Middle East respiratory syndrome-related coronavirus	Hu/Riyadh-KSA-K17000887/2017	orf1ab

MG011360	21434-25495	Middle East respiratory syndrome-related coronavirus	Hu/Riyadh-KSA-K17000887/2017	spike

MG011362	27831-28490	Middle East respiratory syndrome-related coronavirus	Hu/Riyadh-KSA-K37029157/2016	membrane

MG011362	28544-29785	Middle East respiratory syndrome-related coronavirus	Hu/Riyadh-KSA-K37029157/2016	nucleocapsid

MG011362	257-21492	Middle East respiratory syndrome-related coronavirus	Hu/Riyadh-KSA-K37029157/2016	orf1ab

MG011362	21434-25495	Middle East respiratory syndrome-related coronavirus	Hu/Riyadh-KSA-K37029157/2016	spike

MK462252	27853-28512	Middle East respiratory syndrome-related coronavirus	Hu/Tabuk-KSA-153/2018	membrane

MK462252	28566-29807	Middle East respiratory syndrome-related coronavirus	Hu/Tabuk-KSA-153/2018	nucleocapsid

MK462252	279-21514	Middle East respiratory syndrome-related coronavirus	Hu/Tabuk-KSA-153/2018	orf1ab

MK462252	21456-25517	Middle East respiratory syndrome-related coronavirus	Hu/Tabuk-KSA-153/2018	spike

KY581684	279-21514	Middle East respiratory syndrome-related coronavirus	Hu/UAE_002_2013	orf1ab

KY581684	27853-28512	Middle East respiratory syndrome-related coronavirus	Hu/UAE_002_2013	membrane

KY581684	28566-29807	Middle East respiratory syndrome-related coronavirus	Hu/UAE_002_2013	nucleocapsid

KY581684	21456-25517	Middle East respiratory syndrome-related coronavirus	Hu/UAE_002_2013	spike

KY581685	279-21514	Middle East respiratory syndrome-related coronavirus	Hu/UAE_004_2013	orf1ab

KY581685	27853-28512	Middle East respiratory syndrome-related coronavirus	Hu/UAE_004_2013	membrane

KY581685	28566-29807	Middle East respiratory syndrome-related coronavirus	Hu/UAE_004_2013	nucleocapsid

KY581685	21456-25517	Middle East respiratory syndrome-related coronavirus	Hu/UAE_004_2013	spike

KY581686	279-21514	Middle East respiratory syndrome-related coronavirus	Hu/UAE_009_2014	orf1ab

KY581686	27853-28512	Middle East respiratory syndrome-related coronavirus	Hu/UAE_009_2014	membrane

KY581686	28566-29807	Middle East respiratory syndrome-related coronavirus	Hu/UAE_009_2014	nucleocapsid

KY581686	21456-25517	Middle East respiratory syndrome-related coronavirus	Hu/UAE_009_2014	spike

KY581687	279-21514	Middle East respiratory syndrome-related coronavirus	Hu/UAE_011_2013	orf1ab

KY581687	27853-28512	Middle East respiratory syndrome-related coronavirus	Hu/UAE_011_2013	membrane

KY581687	28566-29807	Middle East respiratory syndrome-related coronavirus	Hu/UAE_011_2013	nucleocapsid

KY581687	21456-25517	Middle East respiratory syndrome-related coronavirus	Hu/UAE_011_2013	spike

KY581688	279-21514	Middle East respiratory syndrome-related coronavirus	Hu/UAE_011_2014	orf1ab

KY581688	27853-28512	Middle East respiratory syndrome-related coronavirus	Hu/UAE_011_2014	membrane

KY581688	28566-29807	Middle East respiratory syndrome-related coronavirus	Hu/UAE_011_2014	nucleocapsid

KY581688	21456-25517	Middle East respiratory syndrome-related coronavirus	Hu/UAE_011_2014	spike

KY581689	279-21514	Middle East respiratory syndrome-related coronavirus	Hu/UAE_015_2014	orf1ab

KY581689	27853-28512	Middle East respiratory syndrome-related coronavirus	Hu/UAE_015_2014	membrane

KY581689	28566-29807	Middle East respiratory syndrome-related coronavirus	Hu/UAE_015_2014	nucleocapsid

KY581689	21456-25517	Middle East respiratory syndrome-related coronavirus	Hu/UAE_015_2014	spike

KY581690	279-21514	Middle East respiratory syndrome-related coronavirus	Hu/UAE_017_2014	orf1ab

KY581690	27853-28512	Middle East respiratory syndrome-related coronavirus	Hu/UAE_017_2014	membrane

KY581690	28566-29807	Middle East respiratory syndrome-related coronavirus	Hu/UAE_017_2014	nucleocapsid

KY581690	21456-25517	Middle East respiratory syndrome-related coronavirus	Hu/UAE_017_2014	spike

KY581691	279-21514	Middle East respiratory syndrome-related coronavirus	Hu/UAE_023_2014	orf1ab

KY581691	27853-28512	Middle East respiratory syndrome-related coronavirus	Hu/UAE_023_2014	membrane

KY581691	28566-29807	Middle East respiratory syndrome-related coronavirus	Hu/UAE_023_2014	nucleocapsid

KY581691	21456-25517	Middle East respiratory syndrome-related coronavirus	Hu/UAE_023_2014	spike

KY581692	279-21514	Middle East respiratory syndrome-related coronavirus	Hu/UAE_025_2014	orf1ab

KY581692	27853-28512	Middle East respiratory syndrome-related coronavirus	Hu/UAE_025_2014	membrane

KY581692	28566-29807	Middle East respiratory syndrome-related coronavirus	Hu/UAE_025_2014	nucleocapsid

KY581692	21456-25517	Middle East respiratory syndrome-related coronavirus	Hu/UAE_025_2014	spike

KY581693	279-21514	Middle East respiratory syndrome-related coronavirus	Hu/UAE_032_2014	orf1ab

KY581693	27853-28512	Middle East respiratory syndrome-related coronavirus	Hu/UAE_032_2014	membrane

KY581693	28566-29807	Middle East respiratory syndrome-related coronavirus	Hu/UAE_032_2014	nucleocapsid

KY581693	21456-25517	Middle East respiratory syndrome-related coronavirus	Hu/UAE_032_2014	spike

KY581694	279-21514	Middle East respiratory syndrome-related coronavirus	Hu/UAE_X_2014	orf1ab

KY581694	27853-28512	Middle East respiratory syndrome-related coronavirus	Hu/UAE_X_2014	membrane

KY581694	28566-29807	Middle East respiratory syndrome-related coronavirus	Hu/UAE_X_2014	nucleocapsid

KY581694	21456-25517	Middle East respiratory syndrome-related coronavirus	Hu/UAE_X_2014	spike

MK796425	NA	Middle East respiratory syndrome-related coronavirus	KNIH/002_05_2015	membrane

MK796425	NA	Middle East respiratory syndrome-related coronavirus	KNIH/002_05_2015	nucleocapsid

MK796425	NA	Middle East respiratory syndrome-related coronavirus	KNIH/002_05_2015	orf1ab

MK796425	NA	Middle East respiratory syndrome-related coronavirus	KNIH/002_05_2015	spike

MH259485	NA	Middle East respiratory syndrome-related coronavirus	KSA_1722	orf1ab

MH259485	NA	Middle East respiratory syndrome-related coronavirus	KSA_1722	membrane

MH259485	NA	Middle East respiratory syndrome-related coronavirus	KSA_1722	nucleocapsid

MH259485	NA	Middle East respiratory syndrome-related coronavirus	KSA_1722	spike

MH259486	NA	Middle East respiratory syndrome-related coronavirus	KSA_1723	orf1ab

MH259486	NA	Middle East respiratory syndrome-related coronavirus	KSA_1723	membrane

MH259486	NA	Middle East respiratory syndrome-related coronavirus	KSA_1723	nucleocapsid

MH259486	NA	Middle East respiratory syndrome-related coronavirus	KSA_1723	spike

MG923470	27469-28128	Middle East respiratory syndrome-related coronavirus	MERS-CoV camel/Burkina Faso/CIRAD-HKU434/2015	membrane

MG923470	28182-29423	Middle East respiratory syndrome-related coronavirus	MERS-CoV camel/Burkina Faso/CIRAD-HKU434/2015	nucleocapsid

MG923470	261-21496	Middle East respiratory syndrome-related coronavirus	MERS-CoV camel/Burkina Faso/CIRAD-HKU434/2015	orf1ab

MG923470	21438-25499	Middle East respiratory syndrome-related coronavirus	MERS-CoV camel/Burkina Faso/CIRAD-HKU434/2015	spike

MG923473	27420-28079	Middle East respiratory syndrome-related coronavirus	MERS-CoV camel/Burkina Faso/CIRAD-HKU697/2015	membrane

MG923473	28133-29374	Middle East respiratory syndrome-related coronavirus	MERS-CoV camel/Burkina Faso/CIRAD-HKU697/2015	nucleocapsid

MG923473	261-21496	Middle East respiratory syndrome-related coronavirus	MERS-CoV camel/Burkina Faso/CIRAD-HKU697/2015	orf1ab

MG923473	21438-25499	Middle East respiratory syndrome-related coronavirus	MERS-CoV camel/Burkina Faso/CIRAD-HKU697/2015	spike

MG923471	27470-28129	Middle East respiratory syndrome-related coronavirus	MERS-CoV camel/Burkina Faso/CIRAD-HKU785/2015	membrane

MG923471	28183-29424	Middle East respiratory syndrome-related coronavirus	MERS-CoV camel/Burkina Faso/CIRAD-HKU785/2015	nucleocapsid

MG923471	262-21497	Middle East respiratory syndrome-related coronavirus	MERS-CoV camel/Burkina Faso/CIRAD-HKU785/2015	orf1ab

MG923471	21439-25500	Middle East respiratory syndrome-related coronavirus	MERS-CoV camel/Burkina Faso/CIRAD-HKU785/2015	spike

MG923466	27842-28501	Middle East respiratory syndrome-related coronavirus	MERS-CoV camel/Ethiopia/AAU-EPHI-HKU4412/2017	membrane

MG923466	28555-29796	Middle East respiratory syndrome-related coronavirus	MERS-CoV camel/Ethiopia/AAU-EPHI-HKU4412/2017	nucleocapsid

MG923466	268-21503	Middle East respiratory syndrome-related coronavirus	MERS-CoV camel/Ethiopia/AAU-EPHI-HKU4412/2017	orf1ab

MG923466	21445-25506	Middle East respiratory syndrome-related coronavirus	MERS-CoV camel/Ethiopia/AAU-EPHI-HKU4412/2017	spike

MG923467	27839-28498	Middle East respiratory syndrome-related coronavirus	MERS-CoV camel/Ethiopia/AAU-EPHI-HKU4448/2017	membrane

MG923467	28552-29793	Middle East respiratory syndrome-related coronavirus	MERS-CoV camel/Ethiopia/AAU-EPHI-HKU4448/2017	nucleocapsid

MG923467	268-21500	Middle East respiratory syndrome-related coronavirus	MERS-CoV camel/Ethiopia/AAU-EPHI-HKU4448/2017	orf1ab

MG923467	21442-25503	Middle East respiratory syndrome-related coronavirus	MERS-CoV camel/Ethiopia/AAU-EPHI-HKU4448/2017	spike

MG923468	27839-28498	Middle East respiratory syndrome-related coronavirus	MERS-CoV camel/Ethiopia/AAU-EPHI-HKU4458/2017	membrane

MG923468	28552-29793	Middle East respiratory syndrome-related coronavirus	MERS-CoV camel/Ethiopia/AAU-EPHI-HKU4458/2017	nucleocapsid

MG923468	268-21500	Middle East respiratory syndrome-related coronavirus	MERS-CoV camel/Ethiopia/AAU-EPHI-HKU4458/2017	orf1ab

MG923468	21442-25503	Middle East respiratory syndrome-related coronavirus	MERS-CoV camel/Ethiopia/AAU-EPHI-HKU4458/2017	spike

MH734114	NA	Middle East respiratory syndrome-related coronavirus	MERS-CoV camel/Kenya/C1215/2018	membrane

MH734114	NA	Middle East respiratory syndrome-related coronavirus	MERS-CoV camel/Kenya/C1215/2018	nucleocapsid

MH734114	NA	Middle East respiratory syndrome-related coronavirus	MERS-CoV camel/Kenya/C1215/2018	orf1ab

MH734114	NA	Middle East respiratory syndrome-related coronavirus	MERS-CoV camel/Kenya/C1215/2018	spike

MG923469	27467-28126	Middle East respiratory syndrome-related coronavirus	MERS-CoV camel/Morocco/CIRAD-HKU213/2015	membrane

MG923469	28180-29421	Middle East respiratory syndrome-related coronavirus	MERS-CoV camel/Morocco/CIRAD-HKU213/2015	nucleocapsid

MG923469	264-21499	Middle East respiratory syndrome-related coronavirus	MERS-CoV camel/Morocco/CIRAD-HKU213/2015	orf1ab

MG923469	21441-25502	Middle East respiratory syndrome-related coronavirus	MERS-CoV camel/Morocco/CIRAD-HKU213/2015	spike

MG923472	27434-28093	Middle East respiratory syndrome-related coronavirus	MERS-CoV camel/Nigeria/NS004/2015	membrane

MG923472	28147-29382	Middle East respiratory syndrome-related coronavirus	MERS-CoV camel/Nigeria/NS004/2015	nucleocapsid

MG923472	250-21482	Middle East respiratory syndrome-related coronavirus	MERS-CoV camel/Nigeria/NS004/2015	orf1ab

MG923472	21424-25485	Middle East respiratory syndrome-related coronavirus	MERS-CoV camel/Nigeria/NS004/2015	spike

MG923474	27413-28072	Middle East respiratory syndrome-related coronavirus	MERS-CoV camel/Nigeria/NV1405/2016	membrane

MG923474	28126-29367	Middle East respiratory syndrome-related coronavirus	MERS-CoV camel/Nigeria/NV1405/2016	nucleocapsid

MG923474	267-21499	Middle East respiratory syndrome-related coronavirus	MERS-CoV camel/Nigeria/NV1405/2016	orf1ab

MG923474	21441-25502	Middle East respiratory syndrome-related coronavirus	MERS-CoV camel/Nigeria/NV1405/2016	spike

MG923475	27421-28080	Middle East respiratory syndrome-related coronavirus	MERS-CoV camel/Nigeria/NV1657/2016	membrane

MG923475	28134-29375	Middle East respiratory syndrome-related coronavirus	MERS-CoV camel/Nigeria/NV1657/2016	nucleocapsid

MG923475	267-21499	Middle East respiratory syndrome-related coronavirus	MERS-CoV camel/Nigeria/NV1657/2016	orf1ab

MG923475	21441-25502	Middle East respiratory syndrome-related coronavirus	MERS-CoV camel/Nigeria/NV1657/2016	spike

MG923478	27421-28080	Middle East respiratory syndrome-related coronavirus	MERS-CoV camel/Nigeria/NV1673/2016	membrane

MG923478	28134-29375	Middle East respiratory syndrome-related coronavirus	MERS-CoV camel/Nigeria/NV1673/2016	nucleocapsid

MG923478	267-21499	Middle East respiratory syndrome-related coronavirus	MERS-CoV camel/Nigeria/NV1673/2016	orf1ab

MG923478	21441-25502	Middle East respiratory syndrome-related coronavirus	MERS-CoV camel/Nigeria/NV1673/2016	spike

MG923479	27421-28080	Middle East respiratory syndrome-related coronavirus	MERS-CoV camel/Nigeria/NV1712/2016	membrane

MG923479	28134-29375	Middle East respiratory syndrome-related coronavirus	MERS-CoV camel/Nigeria/NV1712/2016	nucleocapsid

MG923479	267-21499	Middle East respiratory syndrome-related coronavirus	MERS-CoV camel/Nigeria/NV1712/2016	orf1ab

MG923479	21441-25502	Middle East respiratory syndrome-related coronavirus	MERS-CoV camel/Nigeria/NV1712/2016	spike

MG923480	27421-28080	Middle East respiratory syndrome-related coronavirus	MERS-CoV camel/Nigeria/NV1787/2016	membrane

MG923480	28134-29375	Middle East respiratory syndrome-related coronavirus	MERS-CoV camel/Nigeria/NV1787/2016	nucleocapsid

MG923480	267-21499	Middle East respiratory syndrome-related coronavirus	MERS-CoV camel/Nigeria/NV1787/2016	orf1ab

MG923480	21441-25502	Middle East respiratory syndrome-related coronavirus	MERS-CoV camel/Nigeria/NV1787/2016	spike

MG923476	27421-28080	Middle East respiratory syndrome-related coronavirus	MERS-CoV camel/Nigeria/NV1989/2016	membrane

MG923476	28134-29375	Middle East respiratory syndrome-related coronavirus	MERS-CoV camel/Nigeria/NV1989/2016	nucleocapsid

MG923476	267-21499	Middle East respiratory syndrome-related coronavirus	MERS-CoV camel/Nigeria/NV1989/2016	orf1ab

MG923476	21441-25502	Middle East respiratory syndrome-related coronavirus	MERS-CoV camel/Nigeria/NV1989/2016	spike

MG923481	27421-28080	Middle East respiratory syndrome-related coronavirus	MERS-CoV camel/Nigeria/NV2020/2016	membrane

MG923481	28134-29375	Middle East respiratory syndrome-related coronavirus	MERS-CoV camel/Nigeria/NV2020/2016	nucleocapsid

MG923481	267-21499	Middle East respiratory syndrome-related coronavirus	MERS-CoV camel/Nigeria/NV2020/2016	orf1ab

MG923481	21441-25502	Middle East respiratory syndrome-related coronavirus	MERS-CoV camel/Nigeria/NV2020/2016	spike

MG923477	27417-28076	Middle East respiratory syndrome-related coronavirus	MERS-CoV camel/Nigeria/NV2040/2016	membrane

MG923477	28130-29371	Middle East respiratory syndrome-related coronavirus	MERS-CoV camel/Nigeria/NV2040/2016	nucleocapsid

MG923477	263-21495	Middle East respiratory syndrome-related coronavirus	MERS-CoV camel/Nigeria/NV2040/2016	orf1ab

MG923477	21437-25498	Middle East respiratory syndrome-related coronavirus	MERS-CoV camel/Nigeria/NV2040/2016	spike

MK129253	279-21514	Middle East respiratory syndrome-related coronavirus	MERS-CoV/KOR/KCDC/001_2018-TSVi	orf1ab

MK129253	27853-28512	Middle East respiratory syndrome-related coronavirus	MERS-CoV/KOR/KCDC/001_2018-TSVi	membrane

MK129253	28566-29807	Middle East respiratory syndrome-related coronavirus	MERS-CoV/KOR/KCDC/001_2018-TSVi	nucleocapsid

MK129253	21456-25517	Middle East respiratory syndrome-related coronavirus	MERS-CoV/KOR/KCDC/001_2018-TSVi	spike

KT326819	NA	Middle East respiratory syndrome-related coronavirus	MERS-CoV/KOR/KNIH/001_05_2015	membrane

KT326819	NA	Middle East respiratory syndrome-related coronavirus	MERS-CoV/KOR/KNIH/001_05_2015	nucleocapsid

KT326819	NA	Middle East respiratory syndrome-related coronavirus	MERS-CoV/KOR/KNIH/001_05_2015	orf1ab

KT326819	NA	Middle East respiratory syndrome-related coronavirus	MERS-CoV/KOR/KNIH/001_05_2015	spike

KT225476	NA	Middle East respiratory syndrome-related coronavirus	MERS-CoV/THA/CU/17_06_2015	membrane

KT225476	NA	Middle East respiratory syndrome-related coronavirus	MERS-CoV/THA/CU/17_06_2015	nucleocapsid

KT225476	NA	Middle East respiratory syndrome-related coronavirus	MERS-CoV/THA/CU/17_06_2015	orf1ab

KT225476	NA	Middle East respiratory syndrome-related coronavirus	MERS-CoV/THA/CU/17_06_2015	spike

MH822886	267-21502	Middle East respiratory syndrome-related coronavirus	MERS-CoV_England-KSA/1/2018	orf1ab

MH822886	27841-28500	Middle East respiratory syndrome-related coronavirus	MERS-CoV_England-KSA/1/2018	membrane

MH822886	28554-29795	Middle East respiratory syndrome-related coronavirus	MERS-CoV_England-KSA/1/2018	nucleocapsid

MH822886	21444-25505	Middle East respiratory syndrome-related coronavirus	MERS-CoV_England-KSA/1/2018	spike

MF593268	NA	Middle East respiratory syndrome-related coronavirus	Neoromicia/5038	orf1ab

MF593268	NA	Middle East respiratory syndrome-related coronavirus	Neoromicia/5038	membrane

MF593268	NA	Middle East respiratory syndrome-related coronavirus	Neoromicia/5038	nucleocapsid

MF593268	NA	Middle East respiratory syndrome-related coronavirus	Neoromicia/5038	spike

MG021451	27814-28470	Middle East respiratory syndrome-related coronavirus	NL13845	membrane

MG021451	28526-29824	Middle East respiratory syndrome-related coronavirus	NL13845	nucleocapsid

MG021451	237-21517	Middle East respiratory syndrome-related coronavirus	NL13845	orf1ab

MG021451	21459-25529	Middle East respiratory syndrome-related coronavirus	NL13845	spike

MG021452	27810-28466	Middle East respiratory syndrome-related coronavirus	NL140422	membrane

MG021452	28521-29822	Middle East respiratory syndrome-related coronavirus	NL140422	nucleocapsid

MG021452	238-21560	Middle East respiratory syndrome-related coronavirus	NL140422	orf1ab

MG021452	21502-25551	Middle East respiratory syndrome-related coronavirus	NL140422	spike

MK280984	27826-28485	Middle East respiratory syndrome-related coronavirus	Qatar15	membrane

MK280984	28539-29780	Middle East respiratory syndrome-related coronavirus	Qatar15	nucleocapsid

MK280984	252-21487	Middle East respiratory syndrome-related coronavirus	Qatar15	orf1ab

MK280984	21429-25490	Middle East respiratory syndrome-related coronavirus	Qatar15	spike

JX993988	NA	Bat coronavirus Cp/Yunnan2011	Cp/Yunnan2011	membrane

JX993988	NA	Bat coronavirus Cp/Yunnan2011	Cp/Yunnan2011	nucleocapsid

JX993988	NA	Bat coronavirus Cp/Yunnan2011	Cp/Yunnan2011	spike

JX993987	NA	Bat coronavirus Rp/Shaanxi2011	Rp/Shaanxi2011	membrane

JX993987	NA	Bat coronavirus Rp/Shaanxi2011	Rp/Shaanxi2011	nucleocapsid

JX993987	NA	Bat coronavirus Rp/Shaanxi2011	Rp/Shaanxi2011	spike

KJ473812	NA	BtRf-BetaCoV/HeB2013	BtRf-HeB2013	orf1ab

KJ473812	NA	BtRf-BetaCoV/HeB2013	BtRf-HeB2013	membrane

KJ473812	NA	BtRf-BetaCoV/HeB2013	BtRf-HeB2013	nucleocapsid

KJ473812	NA	BtRf-BetaCoV/HeB2013	BtRf-HeB2013	spike

KJ473811	NA	BtRf-BetaCoV/JL2012	BtRf-JL2012	orf1ab

KJ473811	NA	BtRf-BetaCoV/JL2012	BtRf-JL2012	membrane

KJ473811	NA	BtRf-BetaCoV/JL2012	BtRf-JL2012	nucleocapsid

KJ473811	NA	BtRf-BetaCoV/JL2012	BtRf-JL2012	spike

KJ473813	NA	BtRf-BetaCoV/SX2013	BtRf-SX2013	orf1ab

KJ473813	NA	BtRf-BetaCoV/SX2013	BtRf-SX2013	membrane

KJ473813	NA	BtRf-BetaCoV/SX2013	BtRf-SX2013	nucleocapsid

KJ473813	NA	BtRf-BetaCoV/SX2013	BtRf-SX2013	spike

KJ473815	NA	BtRs-BetaCoV/GX2013	BtRs-GX2013	orf1ab

KJ473815	NA	BtRs-BetaCoV/GX2013	BtRs-GX2013	membrane

KJ473815	NA	BtRs-BetaCoV/GX2013	BtRs-GX2013	nucleocapsid

KJ473815	NA	BtRs-BetaCoV/GX2013	BtRs-GX2013	spike

KJ473814	NA	BtRs-BetaCoV/HuB2013	BtRs-HuB2013	orf1ab

KJ473814	NA	BtRs-BetaCoV/HuB2013	BtRs-HuB2013	membrane

KJ473814	NA	BtRs-BetaCoV/HuB2013	BtRs-HuB2013	nucleocapsid

KJ473814	NA	BtRs-BetaCoV/HuB2013	BtRs-HuB2013	spike

KJ473816	NA	BtRs-BetaCoV/YN2013	BtRs-YN2013	orf1ab

KJ473816	NA	BtRs-BetaCoV/YN2013	BtRs-YN2013	membrane

KJ473816	NA	BtRs-BetaCoV/YN2013	BtRs-YN2013	nucleocapsid

KJ473816	NA	BtRs-BetaCoV/YN2013	BtRs-YN2013	spike

DQ648794	NA	Bat coronavirus (BtCoV/133/2005)	BtCoV/133/2005	orf1ab

DQ648794	NA	Bat coronavirus (BtCoV/133/2005)	BtCoV/133/2005	membrane

DQ648794	NA	Bat coronavirus (BtCoV/133/2005)	BtCoV/133/2005	nucleocapsid

DQ648794	NA	Bat coronavirus (BtCoV/133/2005)	BtCoV/133/2005	spike

KU558922	28684-29376	Betacoronavirus 1	Buffalo coronavirus B1-24F	membrane

KU558922	29386-30732	Betacoronavirus 1	Buffalo coronavirus B1-24F	nucleocapsid

KU558922	211-21494	Betacoronavirus 1	Buffalo coronavirus B1-24F	orf1ab

KU558922	23641-27732	Betacoronavirus 1	Buffalo coronavirus B1-24F	spike

KU558923	28639-29331	Betacoronavirus 1	Buffalo coronavirus B1-28F	membrane

KU558923	29341-30687	Betacoronavirus 1	Buffalo coronavirus B1-28F	nucleocapsid

KU558923	165-21448	Betacoronavirus 1	Buffalo coronavirus B1-28F	orf1ab

KU558923	23595-27686	Betacoronavirus 1	Buffalo coronavirus B1-28F	spike

NC_003045	28691-29383	Bovine coronavirus	BCoV-ENT	membrane

NC_003045	29393-30739	Bovine coronavirus	BCoV-ENT	nucleocapsid

NC_003045	211-21494	Bovine coronavirus	BCoV-ENT	orf1ab

NC_003045	23641-27732	Bovine coronavirus	BCoV-ENT	spike

KU886219	NA	Bovine coronavirus	BCV-AKS-01	membrane

KU886219	NA	Bovine coronavirus	BCV-AKS-01	nucleocapsid

KU886219	NA	Bovine coronavirus	BCV-AKS-01	spike

DQ811784	NA	Bovine coronavirus DB2	DB2	membrane

DQ811784	NA	Bovine coronavirus DB2	DB2	nucleocapsid

DQ811784	NA	Bovine coronavirus DB2	DB2	orf1ab

DQ811784	NA	Bovine coronavirus DB2	DB2	spike

EF424619	28675-29367	Bovine coronavirus E-AH187	E-AH187	membrane

EF424619	29377-30723	Bovine coronavirus E-AH187	E-AH187	nucleocapsid

EF424619	23625-27716	Bovine coronavirus E-AH187	E-AH187	spike

EF424615	28691-29383	Bovine coronavirus E-AH65	E-AH65	membrane

EF424615	29393-30739	Bovine coronavirus E-AH65	E-AH65	nucleocapsid

EF424615	23641-27732	Bovine coronavirus E-AH65	E-AH65	spike

EF424616	28675-29367	Bovine coronavirus E-AH65-TC	E-AH65-TC	membrane

EF424616	29377-30723	Bovine coronavirus E-AH65-TC	E-AH65-TC	nucleocapsid

EF424616	23625-27716	Bovine coronavirus E-AH65-TC	E-AH65-TC	spike

EF424620	28675-29367	Bovine coronavirus R-AH187	R-AH187	membrane

EF424620	29377-30723	Bovine coronavirus R-AH187	R-AH187	nucleocapsid

EF424620	23625-27716	Bovine coronavirus R-AH187	R-AH187	spike

EF424617	28675-29367	Bovine coronavirus R-AH65	R-AH65	membrane

EF424617	29377-30723	Bovine coronavirus R-AH65	R-AH65	nucleocapsid

EF424617	23625-27716	Bovine coronavirus R-AH65	R-AH65	spike

EF424618	28675-29367	Bovine coronavirus R-AH65-TC	R-AH65-TC	membrane

EF424618	29377-30723	Bovine coronavirus R-AH65-TC	R-AH65-TC	nucleocapsid

EF424618	23625-27716	Bovine coronavirus R-AH65-TC	R-AH65-TC	spike

EF424624	28659-29351	Calf-giraffe coronavirus US/OH3/2006	US/OH3/2006	membrane

EF424624	29361-30707	Calf-giraffe coronavirus US/OH3/2006	US/OH3/2006	nucleocapsid

EF424624	23625-27701	Calf-giraffe coronavirus US/OH3/2006	US/OH3/2006	spike

KX432213	28624-29316	Canine respiratory coronavirus	BJ232	membrane

KX432213	29326-30672	Canine respiratory coronavirus	BJ232	nucleocapsid

KX432213	144-21427	Canine respiratory coronavirus	BJ232	orf1ab

KX432213	23574-27665	Canine respiratory coronavirus	BJ232	spike

KF906249	28699-29391	Dromedary camel coronavirus HKU23	HKU23-265F	membrane

KF906249	29401-30747	Dromedary camel coronavirus HKU23	HKU23-265F	nucleocapsid

KF906249	215-21498	Dromedary camel coronavirus HKU23	HKU23-265F	orf1ab

KF906249	23645-27745	Dromedary camel coronavirus HKU23	HKU23-265F	spike

KF906250	28699-29391	Dromedary camel coronavirus HKU23	HKU23-362F	membrane

KF906250	29401-30747	Dromedary camel coronavirus HKU23	HKU23-362F	nucleocapsid

KF906250	215-21498	Dromedary camel coronavirus HKU23	HKU23-362F	orf1ab

KF906250	23645-27745	Dromedary camel coronavirus HKU23	HKU23-362F	spike

LC061273	209-21561	Equine coronavirus	Obihiro12-1	orf1ab

LC061273	28585-29277	Equine coronavirus	Obihiro12-1	membrane

LC061273	29287-30627	Equine coronavirus	Obihiro12-1	nucleocapsid

LC061273	23708-27799	Equine coronavirus	Obihiro12-1	spike

LC061274	209-21561	Equine coronavirus	Obihiro12-2	orf1ab

LC061274	28585-29277	Equine coronavirus	Obihiro12-2	membrane

LC061274	29287-30627	Equine coronavirus	Obihiro12-2	nucleocapsid

LC061274	23708-27799	Equine coronavirus	Obihiro12-2	spike

LC061272	209-21570	Equine coronavirus	Tokachi09	orf1ab

LC061272	28451-29143	Equine coronavirus	Tokachi09	membrane

LC061272	29153-30493	Equine coronavirus	Tokachi09	nucleocapsid

LC061272	23719-27810	Equine coronavirus	Tokachi09	spike

EF424622	28659-29351	Giraffe coronavirus US/OH3-TC/2006	US/OH3-TC/2006	membrane

EF424622	29361-30707	Giraffe coronavirus US/OH3-TC/2006	US/OH3-TC/2006	nucleocapsid

EF424622	23625-27701	Giraffe coronavirus US/OH3-TC/2006	US/OH3-TC/2006	spike

EF424623	28659-29351	Giraffe coronavirus US/OH3/2003	US/OH3/2003	membrane

EF424623	29361-30707	Giraffe coronavirus US/OH3/2003	US/OH3/2003	nucleocapsid

EF424623	23625-27701	Giraffe coronavirus US/OH3/2003	US/OH3/2003	spike

KP198611	28400-29092	Human coronavirus OC43	1783A/10	membrane

KP198611	29102-30448	Human coronavirus OC43	1783A/10	nucleocapsid

KP198611	NA	Human coronavirus OC43	1783A/10	orf1ab

KP198611	23640-27728	Human coronavirus OC43	1783A/10	spike

KY014281	125-21411	Human coronavirus OC43	2002-04	orf1ab

KY014281	28300-28992	Human coronavirus OC43	2002-04	membrane

KY014281	29002-30348	Human coronavirus OC43	2002-04	nucleocapsid

KY014281	23558-27628	Human coronavirus OC43	2002-04	spike

KY014282	181-21467	Human coronavirus OC43	2007-09	orf1ab

KY014282	28371-29063	Human coronavirus OC43	2007-09	membrane

KY014282	29073-30419	Human coronavirus OC43	2007-09	nucleocapsid

KY014282	23611-27699	Human coronavirus OC43	2007-09	spike

KP198610	28400-29092	Human coronavirus OC43	2058A/10	membrane

KP198610	29102-30448	Human coronavirus OC43	2058A/10	nucleocapsid

KP198610	NA	Human coronavirus OC43	2058A/10	orf1ab

KP198610	23640-27728	Human coronavirus OC43	2058A/10	spike

KU131570	28393-29085	Human coronavirus OC43	HCoV-OC43/UK/London/2011	membrane

KU131570	29095-30441	Human coronavirus OC43	HCoV-OC43/UK/London/2011	nucleocapsid

KU131570	23642-27721	Human coronavirus OC43	HCoV-OC43/UK/London/2011	spike

KU131570	212-21948	Human coronavirus OC43	HCoV-OC43/UK/London/2011	orf1ab

MF314143	28401-29093	Human coronavirus OC43	HCoV-OC43/USA/ACRI_0052/2016	membrane

MF314143	29103-30449	Human coronavirus OC43	HCoV-OC43/USA/ACRI_0052/2016	nucleocapsid

MF314143	211-21497	Human coronavirus OC43	HCoV-OC43/USA/ACRI_0052/2016	orf1ab

MF314143	23641-27729	Human coronavirus OC43	HCoV-OC43/USA/ACRI_0052/2016	spike

MH121121	28386-29078	Human coronavirus OC43	HCoV-OC43/USA/ACRI_0213/2016	membrane

MH121121	29088-30434	Human coronavirus OC43	HCoV-OC43/USA/ACRI_0213/2016	nucleocapsid

MH121121	207-21493	Human coronavirus OC43	HCoV-OC43/USA/ACRI_0213/2016	orf1ab

MH121121	23649-27725	Human coronavirus OC43	HCoV-OC43/USA/ACRI_0213/2016	spike

MF374984	28393-29085	Human coronavirus OC43	HCoV-OC43/USA/TCNP_00204/2017	membrane

MF374984	29095-30441	Human coronavirus OC43	HCoV-OC43/USA/TCNP_00204/2017	nucleocapsid

MF374984	202-21497	Human coronavirus OC43	HCoV-OC43/USA/TCNP_00204/2017	orf1ab

MF374984	23632-27720	Human coronavirus OC43	HCoV-OC43/USA/TCNP_00204/2017	spike

MF374985	28385-29077	Human coronavirus OC43	HCoV-OC43/USA/TCNP_00212/2017	membrane

MF374985	29087-30433	Human coronavirus OC43	HCoV-OC43/USA/TCNP_00212/2017	nucleocapsid

MF374985	200-21486	Human coronavirus OC43	HCoV-OC43/USA/TCNP_00212/2017	orf1ab

MF374985	23642-27724	Human coronavirus OC43	HCoV-OC43/USA/TCNP_00212/2017	spike

MF374983	28384-29076	Human coronavirus OC43	HCoV-OC43/USA/TCNP_0070/2016	membrane

MF374983	29086-30432	Human coronavirus OC43	HCoV-OC43/USA/TCNP_0070/2016	nucleocapsid

MF374983	205-21491	Human coronavirus OC43	HCoV-OC43/USA/TCNP_0070/2016	orf1ab

MF374983	23647-27723	Human coronavirus OC43	HCoV-OC43/USA/TCNP_0070/2016	spike

JN129835	28386-29078	Human coronavirus OC43	HK04-02	membrane

JN129835	29088-30434	Human coronavirus OC43	HK04-02	nucleocapsid

JN129835	NA	Human coronavirus OC43	HK04-02	orf1ab

JN129835	23640-27725	Human coronavirus OC43	HK04-02	spike

KJ958218	28378-29070	Human coronavirus OC43	LY341	membrane

KJ958218	29080-30426	Human coronavirus OC43	LY341	nucleocapsid

KJ958218	211-21497	Human coronavirus OC43	LY341	orf1ab

KJ958218	23641-27717	Human coronavirus OC43	LY341	spike

KJ958219	28378-29070	Human coronavirus OC43	LY342	membrane

KJ958219	29080-30426	Human coronavirus OC43	LY342	nucleocapsid

KJ958219	211-21497	Human coronavirus OC43	LY342	orf1ab

KJ958219	23641-27717	Human coronavirus OC43	LY342	spike

KX538964	28377-29069	Human coronavirus OC43	MY-U002/12	membrane

KX538964	29079-30425	Human coronavirus OC43	MY-U002/12	nucleocapsid

KX538964	210-21496	Human coronavirus OC43	MY-U002/12	orf1ab

KX538964	23640-27716	Human coronavirus OC43	MY-U002/12	spike

KX538975	28380-29072	Human coronavirus OC43	MY-U1024/12	membrane

KX538975	29082-30428	Human coronavirus OC43	MY-U1024/12	nucleocapsid

KX538975	210-21496	Human coronavirus OC43	MY-U1024/12	orf1ab

KX538975	23640-27719	Human coronavirus OC43	MY-U1024/12	spike

KX538976	28377-29069	Human coronavirus OC43	MY-U1057/12	membrane

KX538976	29079-30425	Human coronavirus OC43	MY-U1057/12	nucleocapsid

KX538976	210-21496	Human coronavirus OC43	MY-U1057/12	orf1ab

KX538976	23640-27716	Human coronavirus OC43	MY-U1057/12	spike

KX538977	28380-29072	Human coronavirus OC43	MY-U1140/12	membrane

KX538977	29082-30428	Human coronavirus OC43	MY-U1140/12	nucleocapsid

KX538977	210-21496	Human coronavirus OC43	MY-U1140/12	orf1ab

KX538977	23640-27719	Human coronavirus OC43	MY-U1140/12	spike

KX538978	28377-29069	Human coronavirus OC43	MY-U1758/13	membrane

KX538978	29079-30425	Human coronavirus OC43	MY-U1758/13	nucleocapsid

KX538978	210-21496	Human coronavirus OC43	MY-U1758/13	orf1ab

KX538978	23640-27716	Human coronavirus OC43	MY-U1758/13	spike

KX538979	28377-29069	Human coronavirus OC43	MY-U1975/13	membrane

KX538979	29079-30425	Human coronavirus OC43	MY-U1975/13	nucleocapsid

KX538979	210-21496	Human coronavirus OC43	MY-U1975/13	orf1ab

KX538979	23640-27716	Human coronavirus OC43	MY-U1975/13	spike

KX538965	28380-29072	Human coronavirus OC43	MY-U208/12	membrane

KX538965	29082-30428	Human coronavirus OC43	MY-U208/12	nucleocapsid

KX538965	210-21496	Human coronavirus OC43	MY-U208/12	orf1ab

KX538965	23640-27719	Human coronavirus OC43	MY-U208/12	spike

KX538966	28377-29069	Human coronavirus OC43	MY-U236/12	membrane

KX538966	29079-30425	Human coronavirus OC43	MY-U236/12	nucleocapsid

KX538966	210-21496	Human coronavirus OC43	MY-U236/12	orf1ab

KX538966	23640-27716	Human coronavirus OC43	MY-U236/12	spike

KX538967	28380-29072	Human coronavirus OC43	MY-U413/12	membrane

KX538967	29082-30428	Human coronavirus OC43	MY-U413/12	nucleocapsid

KX538967	210-21496	Human coronavirus OC43	MY-U413/12	orf1ab

KX538967	23640-27719	Human coronavirus OC43	MY-U413/12	spike

KX538968	28380-29072	Human coronavirus OC43	MY-U464/12	membrane

KX538968	29082-30428	Human coronavirus OC43	MY-U464/12	nucleocapsid

KX538968	210-21496	Human coronavirus OC43	MY-U464/12	orf1ab

KX538968	23640-27719	Human coronavirus OC43	MY-U464/12	spike

KX538969	28380-29072	Human coronavirus OC43	MY-U523/12	membrane

KX538969	29082-30428	Human coronavirus OC43	MY-U523/12	nucleocapsid

KX538969	210-21496	Human coronavirus OC43	MY-U523/12	orf1ab

KX538969	23640-27719	Human coronavirus OC43	MY-U523/12	spike

KX538970	28377-29069	Human coronavirus OC43	MY-U710/12	membrane

KX538970	29079-30425	Human coronavirus OC43	MY-U710/12	nucleocapsid

KX538970	210-21496	Human coronavirus OC43	MY-U710/12	orf1ab

KX538970	23640-27716	Human coronavirus OC43	MY-U710/12	spike

KX538971	28380-29072	Human coronavirus OC43	MY-U732/12	membrane

KX538971	29082-30428	Human coronavirus OC43	MY-U732/12	nucleocapsid

KX538971	210-21496	Human coronavirus OC43	MY-U732/12	orf1ab

KX538971	23640-27719	Human coronavirus OC43	MY-U732/12	spike

KX538972	28377-29069	Human coronavirus OC43	MY-U774/12	membrane

KX538972	29079-30425	Human coronavirus OC43	MY-U774/12	nucleocapsid

KX538972	210-21496	Human coronavirus OC43	MY-U774/12	orf1ab

KX538972	23640-27716	Human coronavirus OC43	MY-U774/12	spike

KX538973	28380-29072	Human coronavirus OC43	MY-U868/12	membrane

KX538973	29082-30428	Human coronavirus OC43	MY-U868/12	nucleocapsid

KX538973	210-21496	Human coronavirus OC43	MY-U868/12	orf1ab

KX538973	23640-27719	Human coronavirus OC43	MY-U868/12	spike

KX538974	28380-29072	Human coronavirus OC43	MY-U945/12	membrane

KX538974	29082-30428	Human coronavirus OC43	MY-U945/12	nucleocapsid

KX538974	210-21496	Human coronavirus OC43	MY-U945/12	orf1ab

KX538974	23640-27719	Human coronavirus OC43	MY-U945/12	spike

KY554972	NA	Human coronavirus OC43	N07-1541B_433X	membrane

KY554972	NA	Human coronavirus OC43	N07-1541B_433X	nucleocapsid

KY554972	NA	Human coronavirus OC43	N07-1541B_433X	orf1ab

KY554972	NA	Human coronavirus OC43	N07-1541B_433X	spike

KY554973	NA	Human coronavirus OC43	N07-1689B_116X	membrane

KY554973	NA	Human coronavirus OC43	N07-1689B_116X	nucleocapsid

KY554973	NA	Human coronavirus OC43	N07-1689B_116X	orf1ab

KY554973	NA	Human coronavirus OC43	N07-1689B_116X	spike

KY554974	NA	Human coronavirus OC43	N08-33B_360X	membrane

KY554974	NA	Human coronavirus OC43	N08-33B_360X	nucleocapsid

KY554974	NA	Human coronavirus OC43	N08-33B_360X	orf1ab

KY554974	NA	Human coronavirus OC43	N08-33B_360X	spike

KY554975	NA	Human coronavirus OC43	N09-382B	membrane

KY554975	NA	Human coronavirus OC43	N09-382B	nucleocapsid

KY554975	NA	Human coronavirus OC43	N09-382B	orf1ab

KY554975	NA	Human coronavirus OC43	N09-382B	spike

KX344031	28377-29069	Human coronavirus OC43	OC43/human/Mex/LRTI_238/2011	membrane

KX344031	29079-30425	Human coronavirus OC43	OC43/human/Mex/LRTI_238/2011	nucleocapsid

KX344031	210-21496	Human coronavirus OC43	OC43/human/Mex/LRTI_238/2011	orf1ab

KX344031	23640-27716	Human coronavirus OC43	OC43/human/Mex/LRTI_238/2011	spike

KF530068	28361-29053	Human coronavirus OC43	OC43/human/USA/007-11/2000	membrane

KF530068	29063-30409	Human coronavirus OC43	OC43/human/USA/007-11/2000	nucleocapsid

KF530068	191-21477	Human coronavirus OC43	OC43/human/USA/007-11/2000	orf1ab

KF530068	23621-27706	Human coronavirus OC43	OC43/human/USA/007-11/2000	spike

KF530092	28366-29058	Human coronavirus OC43	OC43/human/USA/008-5/2000	membrane

KF530092	29068-30414	Human coronavirus OC43	OC43/human/USA/008-5/2000	nucleocapsid

KF530092	191-21477	Human coronavirus OC43	OC43/human/USA/008-5/2000	orf1ab

KF530092	23624-27694	Human coronavirus OC43	OC43/human/USA/008-5/2000	spike

KF530060	28366-29058	Human coronavirus OC43	OC43/human/USA/851-15/1985	membrane

KF530060	29068-30414	Human coronavirus OC43	OC43/human/USA/851-15/1985	nucleocapsid

KF530060	191-21477	Human coronavirus OC43	OC43/human/USA/851-15/1985	orf1ab

KF530060	23624-27694	Human coronavirus OC43	OC43/human/USA/851-15/1985	spike

KF530085	28358-29050	Human coronavirus OC43	OC43/human/USA/871-25/1987	membrane

KF530085	29060-30406	Human coronavirus OC43	OC43/human/USA/871-25/1987	nucleocapsid

KF530085	183-21469	Human coronavirus OC43	OC43/human/USA/871-25/1987	orf1ab

KF530085	23616-27686	Human coronavirus OC43	OC43/human/USA/871-25/1987	spike

KF530086	28366-29058	Human coronavirus OC43	OC43/human/USA/872-5/1987	membrane

KF530086	29068-30414	Human coronavirus OC43	OC43/human/USA/872-5/1987	nucleocapsid

KF530086	191-21477	Human coronavirus OC43	OC43/human/USA/872-5/1987	orf1ab

KF530086	23624-27694	Human coronavirus OC43	OC43/human/USA/872-5/1987	spike

KF530077	28366-29058	Human coronavirus OC43	OC43/human/USA/873-16/1987	membrane

KF530077	29068-30414	Human coronavirus OC43	OC43/human/USA/873-16/1987	nucleocapsid

KF530077	191-21477	Human coronavirus OC43	OC43/human/USA/873-16/1987	orf1ab

KF530077	23624-27694	Human coronavirus OC43	OC43/human/USA/873-16/1987	spike

KF530083	28366-29058	Human coronavirus OC43	OC43/human/USA/873-19/1987	membrane

KF530083	29068-30414	Human coronavirus OC43	OC43/human/USA/873-19/1987	nucleocapsid

KF530083	191-21477	Human coronavirus OC43	OC43/human/USA/873-19/1987	orf1ab

KF530083	23624-27694	Human coronavirus OC43	OC43/human/USA/873-19/1987	spike

KF530087	28366-29058	Human coronavirus OC43	OC43/human/USA/873-6/1987	membrane

KF530087	29068-30414	Human coronavirus OC43	OC43/human/USA/873-6/1987	nucleocapsid

KF530087	191-21477	Human coronavirus OC43	OC43/human/USA/873-6/1987	orf1ab

KF530087	23624-27694	Human coronavirus OC43	OC43/human/USA/873-6/1987	spike

KF530073	28366-29058	Human coronavirus OC43	OC43/human/USA/8912-37/1989	membrane

KF530073	29068-30414	Human coronavirus OC43	OC43/human/USA/8912-37/1989	nucleocapsid

KF530073	191-21477	Human coronavirus OC43	OC43/human/USA/8912-37/1989	orf1ab

KF530073	23624-27694	Human coronavirus OC43	OC43/human/USA/8912-37/1989	spike

KF530066	28353-29045	Human coronavirus OC43	OC43/human/USA/901-33/1990	membrane

KF530066	29055-30401	Human coronavirus OC43	OC43/human/USA/901-33/1990	nucleocapsid

KF530066	178-21464	Human coronavirus OC43	OC43/human/USA/901-33/1990	orf1ab

KF530066	23611-27681	Human coronavirus OC43	OC43/human/USA/901-33/1990	spike

KF530065	28366-29058	Human coronavirus OC43	OC43/human/USA/901-41/1990	membrane

KF530065	29068-30414	Human coronavirus OC43	OC43/human/USA/901-41/1990	nucleocapsid

KF530065	191-21477	Human coronavirus OC43	OC43/human/USA/901-41/1990	orf1ab

KF530065	23624-27694	Human coronavirus OC43	OC43/human/USA/901-41/1990	spike

KF530061	28366-29058	Human coronavirus OC43	OC43/human/USA/901-43/1990	membrane

KF530061	29068-30414	Human coronavirus OC43	OC43/human/USA/901-43/1990	nucleocapsid

KF530061	191-21477	Human coronavirus OC43	OC43/human/USA/901-43/1990	orf1ab

KF530061	23624-27694	Human coronavirus OC43	OC43/human/USA/901-43/1990	spike

KF530088	28367-29059	Human coronavirus OC43	OC43/human/USA/901-54/1990	membrane

KF530088	29069-30415	Human coronavirus OC43	OC43/human/USA/901-54/1990	nucleocapsid

KF530088	191-21477	Human coronavirus OC43	OC43/human/USA/901-54/1990	orf1ab

KF530088	23624-27706	Human coronavirus OC43	OC43/human/USA/901-54/1990	spike

KF530076	28381-29073	Human coronavirus OC43	OC43/human/USA/911-11/1991	membrane

KF530076	29083-30429	Human coronavirus OC43	OC43/human/USA/911-11/1991	nucleocapsid

KF530076	205-21491	Human coronavirus OC43	OC43/human/USA/911-11/1991	orf1ab

KF530076	23638-27720	Human coronavirus OC43	OC43/human/USA/911-11/1991	spike

KF530096	28364-29056	Human coronavirus OC43	OC43/human/USA/911-38/1991	membrane

KF530096	29066-30412	Human coronavirus OC43	OC43/human/USA/911-38/1991	nucleocapsid

KF530096	188-21474	Human coronavirus OC43	OC43/human/USA/911-38/1991	orf1ab

KF530096	23621-27703	Human coronavirus OC43	OC43/human/USA/911-38/1991	spike

KF530091	28381-29073	Human coronavirus OC43	OC43/human/USA/911-58/1991	membrane

KF530091	29083-30429	Human coronavirus OC43	OC43/human/USA/911-58/1991	nucleocapsid

KF530091	205-21491	Human coronavirus OC43	OC43/human/USA/911-58/1991	orf1ab

KF530091	23638-27720	Human coronavirus OC43	OC43/human/USA/911-58/1991	spike

KF530089	28384-29076	Human coronavirus OC43	OC43/human/USA/911-66/1991	membrane

KF530089	29086-30432	Human coronavirus OC43	OC43/human/USA/911-66/1991	nucleocapsid

KF530089	205-21491	Human coronavirus OC43	OC43/human/USA/911-66/1991	orf1ab

KF530089	23638-27723	Human coronavirus OC43	OC43/human/USA/911-66/1991	spike

KF530067	28383-29075	Human coronavirus OC43	OC43/human/USA/912-10/1991	membrane

KF530067	29085-30431	Human coronavirus OC43	OC43/human/USA/912-10/1991	nucleocapsid

KF530067	204-21490	Human coronavirus OC43	OC43/human/USA/912-10/1991	orf1ab

KF530067	23637-27722	Human coronavirus OC43	OC43/human/USA/912-10/1991	spike

KF530082	28370-29062	Human coronavirus OC43	OC43/human/USA/912-11/1991	membrane

KF530082	29072-30418	Human coronavirus OC43	OC43/human/USA/912-11/1991	nucleocapsid

KF530082	191-21477	Human coronavirus OC43	OC43/human/USA/912-11/1991	orf1ab

KF530082	23624-27709	Human coronavirus OC43	OC43/human/USA/912-11/1991	spike

KF530094	28370-29062	Human coronavirus OC43	OC43/human/USA/912-36/1991	membrane

KF530094	29072-30418	Human coronavirus OC43	OC43/human/USA/912-36/1991	nucleocapsid

KF530094	191-21477	Human coronavirus OC43	OC43/human/USA/912-36/1991	orf1ab

KF530094	23624-27709	Human coronavirus OC43	OC43/human/USA/912-36/1991	spike

KF530095	28377-29069	Human coronavirus OC43	OC43/human/USA/912-6/1991	membrane

KF530095	29079-30425	Human coronavirus OC43	OC43/human/USA/912-6/1991	nucleocapsid

KF530095	201-21487	Human coronavirus OC43	OC43/human/USA/912-6/1991	orf1ab

KF530095	23634-27716	Human coronavirus OC43	OC43/human/USA/912-6/1991	spike

KF530079	28370-29062	Human coronavirus OC43	OC43/human/USA/913-29/1991	membrane

KF530079	29072-30418	Human coronavirus OC43	OC43/human/USA/913-29/1991	nucleocapsid

KF530079	191-21477	Human coronavirus OC43	OC43/human/USA/913-29/1991	orf1ab

KF530079	23624-27709	Human coronavirus OC43	OC43/human/USA/913-29/1991	spike

KF530097	28366-29058	Human coronavirus OC43	OC43/human/USA/9211-43/1992	membrane

KF530097	29068-30414	Human coronavirus OC43	OC43/human/USA/9211-43/1992	nucleocapsid

KF530097	191-21477	Human coronavirus OC43	OC43/human/USA/9211-43/1992	orf1ab

KF530097	23624-27694	Human coronavirus OC43	OC43/human/USA/9211-43/1992	spike

KF530074	28363-29055	Human coronavirus OC43	OC43/human/USA/9212-33/1992	membrane

KF530074	29065-30411	Human coronavirus OC43	OC43/human/USA/9212-33/1992	nucleocapsid

KF530074	188-21474	Human coronavirus OC43	OC43/human/USA/9212-33/1992	orf1ab

KF530074	23621-27691	Human coronavirus OC43	OC43/human/USA/9212-33/1992	spike

KF530071	28367-29059	Human coronavirus OC43	OC43/human/USA/925-1/1992	membrane

KF530071	29069-30415	Human coronavirus OC43	OC43/human/USA/925-1/1992	nucleocapsid

KF530071	191-21477	Human coronavirus OC43	OC43/human/USA/925-1/1992	orf1ab

KF530071	23624-27706	Human coronavirus OC43	OC43/human/USA/925-1/1992	spike

KF530090	28366-29058	Human coronavirus OC43	OC43/human/USA/931-85/1993	membrane

KF530090	29068-30414	Human coronavirus OC43	OC43/human/USA/931-85/1993	nucleocapsid

KF530090	191-21477	Human coronavirus OC43	OC43/human/USA/931-85/1993	orf1ab

KF530090	23624-27694	Human coronavirus OC43	OC43/human/USA/931-85/1993	spike

KF530084	28367-29059	Human coronavirus OC43	OC43/human/USA/951-18/1995	membrane

KF530084	29069-30415	Human coronavirus OC43	OC43/human/USA/951-18/1995	nucleocapsid

KF530084	191-21477	Human coronavirus OC43	OC43/human/USA/951-18/1995	orf1ab

KF530084	23621-27706	Human coronavirus OC43	OC43/human/USA/951-18/1995	spike

KF530075	28366-29058	Human coronavirus OC43	OC43/human/USA/953-23/1995	membrane

KF530075	29068-30414	Human coronavirus OC43	OC43/human/USA/953-23/1995	nucleocapsid

KF530075	191-21477	Human coronavirus OC43	OC43/human/USA/953-23/1995	orf1ab

KF530075	23624-27694	Human coronavirus OC43	OC43/human/USA/953-23/1995	spike

KF530078	28353-29045	Human coronavirus OC43	OC43/human/USA/9612-29/1996	membrane

KF530078	29055-30401	Human coronavirus OC43	OC43/human/USA/9612-29/1996	nucleocapsid

KF530078	178-21464	Human coronavirus OC43	OC43/human/USA/9612-29/1996	orf1ab

KF530078	23611-27681	Human coronavirus OC43	OC43/human/USA/9612-29/1996	spike

KF530063	28366-29058	Human coronavirus OC43	OC43/human/USA/9612-48/1996	membrane

KF530063	29068-30414	Human coronavirus OC43	OC43/human/USA/9612-48/1996	nucleocapsid

KF530063	191-21477	Human coronavirus OC43	OC43/human/USA/9612-48/1996	orf1ab

KF530063	23621-27706	Human coronavirus OC43	OC43/human/USA/9612-48/1996	spike

KF530064	28366-29058	Human coronavirus OC43	OC43/human/USA/9612-9/1996	membrane

KF530064	29068-30414	Human coronavirus OC43	OC43/human/USA/9612-9/1996	nucleocapsid

KF530064	191-21477	Human coronavirus OC43	OC43/human/USA/9612-9/1996	orf1ab

KF530064	23624-27694	Human coronavirus OC43	OC43/human/USA/9612-9/1996	spike

KF530098	28367-29059	Human coronavirus OC43	OC43/human/USA/965-6/1996	membrane

KF530098	29069-30415	Human coronavirus OC43	OC43/human/USA/965-6/1996	nucleocapsid

KF530098	191-21477	Human coronavirus OC43	OC43/human/USA/965-6/1996	orf1ab

KF530098	23621-27706	Human coronavirus OC43	OC43/human/USA/965-6/1996	spike

KF530099	28367-29059	Human coronavirus OC43	OC43/human/USA/971-5/1997	membrane

KF530099	29069-30415	Human coronavirus OC43	OC43/human/USA/971-5/1997	nucleocapsid

KF530099	191-21477	Human coronavirus OC43	OC43/human/USA/971-5/1997	orf1ab

KF530099	23621-27706	Human coronavirus OC43	OC43/human/USA/971-5/1997	spike

KF530072	28366-29058	Human coronavirus OC43	OC43/human/USA/9712-13/1997	membrane

KF530072	29068-30414	Human coronavirus OC43	OC43/human/USA/9712-13/1997	nucleocapsid

KF530072	191-21477	Human coronavirus OC43	OC43/human/USA/9712-13/1997	orf1ab

KF530072	23624-27694	Human coronavirus OC43	OC43/human/USA/9712-13/1997	spike

KF530080	28366-29058	Human coronavirus OC43	OC43/human/USA/9712-31/1997	membrane

KF530080	29068-30414	Human coronavirus OC43	OC43/human/USA/9712-31/1997	nucleocapsid

KF530080	191-21477	Human coronavirus OC43	OC43/human/USA/9712-31/1997	orf1ab

KF530080	23624-27694	Human coronavirus OC43	OC43/human/USA/9712-31/1997	spike

KF530069	28367-29059	Human coronavirus OC43	OC43/human/USA/982-4/1998	membrane

KF530069	29069-30415	Human coronavirus OC43	OC43/human/USA/982-4/1998	nucleocapsid

KF530069	191-21477	Human coronavirus OC43	OC43/human/USA/982-4/1998	orf1ab

KF530069	23621-27706	Human coronavirus OC43	OC43/human/USA/982-4/1998	spike

KF530070	28362-29054	Human coronavirus OC43	OC43/human/USA/991-19/1999	membrane

KF530070	29064-30410	Human coronavirus OC43	OC43/human/USA/991-19/1999	nucleocapsid

KF530070	192-21478	Human coronavirus OC43	OC43/human/USA/991-19/1999	orf1ab

KF530070	23622-27707	Human coronavirus OC43	OC43/human/USA/991-19/1999	spike

KF530081	28360-29052	Human coronavirus OC43	OC43/human/USA/991-5/1999	membrane

KF530081	29062-30408	Human coronavirus OC43	OC43/human/USA/991-5/1999	nucleocapsid

KF530081	190-21476	Human coronavirus OC43	OC43/human/USA/991-5/1999	orf1ab

KF530081	23620-27705	Human coronavirus OC43	OC43/human/USA/991-5/1999	spike

KY684759	NA	Human coronavirus OC43	SC2269	membrane

KY684759	NA	Human coronavirus OC43	SC2269	nucleocapsid

KY684759	NA	Human coronavirus OC43	SC2269	orf1ab

KY684759	NA	Human coronavirus OC43	SC2269	spike

KY967361	NA	Human coronavirus OC43	SC2345	membrane

KY967361	NA	Human coronavirus OC43	SC2345	nucleocapsid

KY967361	NA	Human coronavirus OC43	SC2345	orf1ab

KY967361	NA	Human coronavirus OC43	SC2345	spike

KY967360	NA	Human coronavirus OC43	SC2476	membrane

KY967360	NA	Human coronavirus OC43	SC2476	nucleocapsid

KY967360	NA	Human coronavirus OC43	SC2476	orf1ab

KY967360	NA	Human coronavirus OC43	SC2476	spike

KY983583	NA	Human coronavirus OC43	SC2481	membrane

KY983583	NA	Human coronavirus OC43	SC2481	nucleocapsid

KY983583	NA	Human coronavirus OC43	SC2481	orf1ab

KY983583	NA	Human coronavirus OC43	SC2481	spike

KY967359	NA	Human coronavirus OC43	SC2730	membrane

KY967359	NA	Human coronavirus OC43	SC2730	nucleocapsid

KY967359	NA	Human coronavirus OC43	SC2730	orf1ab

KY967359	NA	Human coronavirus OC43	SC2730	spike

KY967358	NA	Human coronavirus OC43	SC2770	membrane

KY967358	NA	Human coronavirus OC43	SC2770	nucleocapsid

KY967358	NA	Human coronavirus OC43	SC2770	orf1ab

KY967358	NA	Human coronavirus OC43	SC2770	spike

KY983585	NA	Human coronavirus OC43	SC2854	membrane

KY983585	NA	Human coronavirus OC43	SC2854	nucleocapsid

KY983585	NA	Human coronavirus OC43	SC2854	orf1ab

KY983585	NA	Human coronavirus OC43	SC2854	spike

KY967356	NA	Human coronavirus OC43	SC2924	membrane

KY967356	NA	Human coronavirus OC43	SC2924	nucleocapsid

KY967356	NA	Human coronavirus OC43	SC2924	orf1ab

KY967356	NA	Human coronavirus OC43	SC2924	spike

KY983588	NA	Human coronavirus OC43	SC3118	membrane

KY983588	NA	Human coronavirus OC43	SC3118	nucleocapsid

KY983588	NA	Human coronavirus OC43	SC3118	orf1ab

KY983588	NA	Human coronavirus OC43	SC3118	spike

KY369906	NA	Human coronavirus OC43	SC622	membrane

KY369906	NA	Human coronavirus OC43	SC622	nucleocapsid

KY369906	NA	Human coronavirus OC43	SC622	orf1ab

KY369906	NA	Human coronavirus OC43	SC622	spike

KY369905	NA	Human coronavirus OC43	SC831	membrane

KY369905	NA	Human coronavirus OC43	SC831	nucleocapsid

KY369905	NA	Human coronavirus OC43	SC831	orf1ab

KY369905	NA	Human coronavirus OC43	SC831	spike

KY369907	NA	Human coronavirus OC43	SC9741	membrane

KY369907	NA	Human coronavirus OC43	SC9741	nucleocapsid

KY369907	NA	Human coronavirus OC43	SC9741	orf1ab

KY369907	NA	Human coronavirus OC43	SC9741	spike

MG977451	28386-29078	Human coronavirus OC43	TNP 12636	membrane

MG977451	29088-30434	Human coronavirus OC43	TNP 12636	nucleocapsid

MG977451	204-21490	Human coronavirus OC43	TNP 12636	orf1ab

MG977451	23646-27725	Human coronavirus OC43	TNP 12636	spike

MG977452	28368-29060	Human coronavirus OC43	TNP 12643	membrane

MG977452	29070-30416	Human coronavirus OC43	TNP 12643	nucleocapsid

MG977452	186-21472	Human coronavirus OC43	TNP 12643	orf1ab

MG977452	23628-27707	Human coronavirus OC43	TNP 12643	spike

MG977444	28248-28940	Human coronavirus OC43	TNP F1778_2	membrane

MG977444	28950-30296	Human coronavirus OC43	TNP F1778_2	nucleocapsid

MG977444	66-21352	Human coronavirus OC43	TNP F1778_2	orf1ab

MG977444	23508-27587	Human coronavirus OC43	TNP F1778_2	spike

MG977445	28343-29035	Human coronavirus OC43	TNP F1790_2	membrane

MG977445	29045-30391	Human coronavirus OC43	TNP F1790_2	nucleocapsid

MG977445	161-21447	Human coronavirus OC43	TNP F1790_2	orf1ab

MG977445	23603-27682	Human coronavirus OC43	TNP F1790_2	spike

MG977447	28369-29061	Human coronavirus OC43	TNP F1832_2	membrane

MG977447	29071-30417	Human coronavirus OC43	TNP F1832_2	nucleocapsid

MG977447	187-21473	Human coronavirus OC43	TNP F1832_2	orf1ab

MG977447	23629-27708	Human coronavirus OC43	TNP F1832_2	spike

MG977449	28385-29077	Human coronavirus OC43	TNP F1834_2	membrane

MG977449	29087-30433	Human coronavirus OC43	TNP F1834_2	nucleocapsid

MG977449	203-21489	Human coronavirus OC43	TNP F1834_2	orf1ab

MG977449	23645-27724	Human coronavirus OC43	TNP F1834_2	spike

MF083115	28345-29037	Porcine hemagglutinating encephalomyelitis virus	CC14	membrane

MF083115	29047-30396	Porcine hemagglutinating encephalomyelitis virus	CC14	nucleocapsid

MF083115	211-21494	Porcine hemagglutinating encephalomyelitis virus	CC14	orf1ab

MF083115	23643-27692	Porcine hemagglutinating encephalomyelitis virus	CC14	spike

KY994645	28345-29037	Porcine hemagglutinating encephalomyelitis virus	JL/2008	membrane

KY994645	29047-30396	Porcine hemagglutinating encephalomyelitis virus	JL/2008	nucleocapsid

KY994645	211-21494	Porcine hemagglutinating encephalomyelitis virus	JL/2008	orf1ab

KY994645	23643-27692	Porcine hemagglutinating encephalomyelitis virus	JL/2008	spike

KY419104	NA	Porcine hemagglutinating encephalomyelitis virus	PHEV CoV USA-15TOSU0331	membrane

KY419104	NA	Porcine hemagglutinating encephalomyelitis virus	PHEV CoV USA-15TOSU0331	nucleocapsid

KY419104	NA	Porcine hemagglutinating encephalomyelitis virus	PHEV CoV USA-15TOSU0331	orf1ab

KY419104	NA	Porcine hemagglutinating encephalomyelitis virus	PHEV CoV USA-15TOSU0331	spike

KY419105	NA	Porcine hemagglutinating encephalomyelitis virus	PHEV CoV USA-15TOSU0582	membrane

KY419105	NA	Porcine hemagglutinating encephalomyelitis virus	PHEV CoV USA-15TOSU0582	nucleocapsid

KY419105	NA	Porcine hemagglutinating encephalomyelitis virus	PHEV CoV USA-15TOSU0582	orf1ab

KY419105	NA	Porcine hemagglutinating encephalomyelitis virus	PHEV CoV USA-15TOSU0582	spike

KY419110	NA	Porcine hemagglutinating encephalomyelitis virus	PHEV CoV USA-15TOSU1362	membrane

KY419110	NA	Porcine hemagglutinating encephalomyelitis virus	PHEV CoV USA-15TOSU1362	nucleocapsid

KY419110	NA	Porcine hemagglutinating encephalomyelitis virus	PHEV CoV USA-15TOSU1362	orf1ab

KY419110	NA	Porcine hemagglutinating encephalomyelitis virus	PHEV CoV USA-15TOSU1362	spike

KY419113	NA	Porcine hemagglutinating encephalomyelitis virus	PHEV CoV USA-15TOSU1582	membrane

KY419113	NA	Porcine hemagglutinating encephalomyelitis virus	PHEV CoV USA-15TOSU1582	nucleocapsid

KY419113	NA	Porcine hemagglutinating encephalomyelitis virus	PHEV CoV USA-15TOSU1582	orf1ab

KY419113	NA	Porcine hemagglutinating encephalomyelitis virus	PHEV CoV USA-15TOSU1582	spike

KY419109	NA	Porcine hemagglutinating encephalomyelitis virus	PHEV CoV USA-15TOSU1655	membrane

KY419109	NA	Porcine hemagglutinating encephalomyelitis virus	PHEV CoV USA-15TOSU1655	nucleocapsid

KY419109	NA	Porcine hemagglutinating encephalomyelitis virus	PHEV CoV USA-15TOSU1655	orf1ab

KY419109	NA	Porcine hemagglutinating encephalomyelitis virus	PHEV CoV USA-15TOSU1655	spike

KY419111	NA	Porcine hemagglutinating encephalomyelitis virus	PHEV CoV USA-15TOSU1727	membrane

KY419111	NA	Porcine hemagglutinating encephalomyelitis virus	PHEV CoV USA-15TOSU1727	nucleocapsid

KY419111	NA	Porcine hemagglutinating encephalomyelitis virus	PHEV CoV USA-15TOSU1727	orf1ab

KY419111	NA	Porcine hemagglutinating encephalomyelitis virus	PHEV CoV USA-15TOSU1727	spike

KY419112	NA	Porcine hemagglutinating encephalomyelitis virus	PHEV CoV USA-15TOSU1765	membrane

KY419112	NA	Porcine hemagglutinating encephalomyelitis virus	PHEV CoV USA-15TOSU1765	nucleocapsid

KY419112	NA	Porcine hemagglutinating encephalomyelitis virus	PHEV CoV USA-15TOSU1765	orf1ab

KY419112	NA	Porcine hemagglutinating encephalomyelitis virus	PHEV CoV USA-15TOSU1765	spike

KY419106	NA	Porcine hemagglutinating encephalomyelitis virus	PHEV CoV USA-15TOSU1785	membrane

KY419106	NA	Porcine hemagglutinating encephalomyelitis virus	PHEV CoV USA-15TOSU1785	nucleocapsid

KY419106	NA	Porcine hemagglutinating encephalomyelitis virus	PHEV CoV USA-15TOSU1785	orf1ab

KY419106	NA	Porcine hemagglutinating encephalomyelitis virus	PHEV CoV USA-15TOSU1785	spike

KY419103	NA	Porcine hemagglutinating encephalomyelitis virus	PHEV CoV USA-15TOSU25049	membrane

KY419103	NA	Porcine hemagglutinating encephalomyelitis virus	PHEV CoV USA-15TOSU25049	nucleocapsid

KY419103	NA	Porcine hemagglutinating encephalomyelitis virus	PHEV CoV USA-15TOSU25049	orf1ab

KY419103	NA	Porcine hemagglutinating encephalomyelitis virus	PHEV CoV USA-15TOSU25049	spike

EF424621	28675-29367	Sable antelope coronavirus US/OH1/2003	US/OH1/2003	membrane

EF424621	29377-30723	Sable antelope coronavirus US/OH1/2003	US/OH1/2003	nucleocapsid

EF424621	23626-27717	Sable antelope coronavirus US/OH1/2003	US/OH1/2003	spike

FJ425188	NA	Sambar deer coronavirus US/OH-WD388-TC/1994	UNKNOWN-FJ425188	membrane

FJ425188	NA	Sambar deer coronavirus US/OH-WD388-TC/1994	UNKNOWN-FJ425188	nucleocapsid

FJ425188	NA	Sambar deer coronavirus US/OH-WD388-TC/1994	UNKNOWN-FJ425188	spike

FJ425190	NA	Sambar deer coronavirus US/OH-WD388-TC/1994	UNKNOWN-FJ425190	membrane

FJ425190	NA	Sambar deer coronavirus US/OH-WD388-TC/1994	UNKNOWN-FJ425190	nucleocapsid

FJ425190	NA	Sambar deer coronavirus US/OH-WD388-TC/1994	UNKNOWN-FJ425190	spike

FJ425189	NA	Sambar deer coronavirus US/OH-WD388/1994	UNKNOWN-FJ425189	membrane

FJ425189	NA	Sambar deer coronavirus US/OH-WD388/1994	UNKNOWN-FJ425189	nucleocapsid

FJ425189	NA	Sambar deer coronavirus US/OH-WD388/1994	UNKNOWN-FJ425189	spike

MG518518	NA	Water deer coronavirus	W17-18	orf1ab

MG518518	NA	Water deer coronavirus	W17-18	membrane

MG518518	NA	Water deer coronavirus	W17-18	nucleocapsid

MG518518	NA	Water deer coronavirus	W17-18	spike

FJ425185	NA	Waterbuck coronavirus US/OH-WD358-GnC/1994	UNKNOWN-FJ425185	membrane

FJ425185	NA	Waterbuck coronavirus US/OH-WD358-GnC/1994	UNKNOWN-FJ425185	nucleocapsid

FJ425185	NA	Waterbuck coronavirus US/OH-WD358-GnC/1994	UNKNOWN-FJ425185	spike

FJ425184	NA	Waterbuck coronavirus US/OH-WD358-TC/1994	UNKNOWN-FJ425184	membrane

FJ425184	NA	Waterbuck coronavirus US/OH-WD358-TC/1994	UNKNOWN-FJ425184	nucleocapsid

FJ425184	NA	Waterbuck coronavirus US/OH-WD358-TC/1994	UNKNOWN-FJ425184	spike

FJ425186	NA	Waterbuck coronavirus US/OH-WD358/1994	UNKNOWN-FJ425186	membrane

FJ425186	NA	Waterbuck coronavirus US/OH-WD358/1994	UNKNOWN-FJ425186	nucleocapsid

FJ425186	NA	Waterbuck coronavirus US/OH-WD358/1994	UNKNOWN-FJ425186	spike

FJ425187	NA	White-tailed deer coronavirus US/OH-WD470/1994	UNKNOWN-FJ425187	membrane

FJ425187	NA	White-tailed deer coronavirus US/OH-WD470/1994	UNKNOWN-FJ425187	nucleocapsid

FJ425187	NA	White-tailed deer coronavirus US/OH-WD470/1994	UNKNOWN-FJ425187	spike

MH810163	NA	Yak coronavirus	YAK/HY24/CH/2017	membrane

MH810163	NA	Yak coronavirus	YAK/HY24/CH/2017	nucleocapsid

MH810163	NA	Yak coronavirus	YAK/HY24/CH/2017	spike

AF220295	28707-29399	Bovine coronavirus	Quebec	membrane

AF220295	29409-30755	Bovine coronavirus	Quebec	nucleocapsid

AF220295	23655-27746	Bovine coronavirus	Quebec	spike

FJ938065	NA	Bovine respiratory coronavirus AH187	AH187	membrane

FJ938065	NA	Bovine respiratory coronavirus AH187	AH187	nucleocapsid

FJ938065	NA	Bovine respiratory coronavirus AH187	AH187	orf1ab

FJ938065	NA	Bovine respiratory coronavirus AH187	AH187	spike

FJ938066	NA	Bovine respiratory coronavirus bovine/US/OH-440-TC/1996	bovine/US/OH-440-TC/1996	membrane

FJ938066	NA	Bovine respiratory coronavirus bovine/US/OH-440-TC/1996	bovine/US/OH-440-TC/1996	nucleocapsid

FJ938066	NA	Bovine respiratory coronavirus bovine/US/OH-440-TC/1996	bovine/US/OH-440-TC/1996	orf1ab

FJ938066	NA	Bovine respiratory coronavirus bovine/US/OH-440-TC/1996	bovine/US/OH-440-TC/1996	spike

EF446615	210-21595	Equine coronavirus	NC99	orf1ab

EF446615	28661-29353	Equine coronavirus	NC99	membrane

EF446615	29363-30703	Equine coronavirus	NC99	nucleocapsid

EF446615	23744-27835	Equine coronavirus	NC99	spike

AY391777	28402-29094	Human coronavirus OC43	ATCC VR-759	membrane

AY391777	29104-30450	Human coronavirus OC43	ATCC VR-759	nucleocapsid

AY391777	23644-27729	Human coronavirus OC43	ATCC VR-759	spike

AY391777	NA	Human coronavirus OC43	ATCC VR-759	orf1ab

JN129834	28374-29066	Human coronavirus OC43	HK04-01	membrane

JN129834	29076-30422	Human coronavirus OC43	HK04-01	nucleocapsid

JN129834	NA	Human coronavirus OC43	HK04-01	orf1ab

JN129834	23628-27713	Human coronavirus OC43	HK04-01	spike

DQ011855	28141-28833	Porcine hemagglutinating encephalomyelitis virus	VW572	membrane

DQ011855	28843-30192	Porcine hemagglutinating encephalomyelitis virus	VW572	nucleocapsid

DQ011855	NA	Porcine hemagglutinating encephalomyelitis virus	VW572	orf1ab

DQ011855	23427-27476	Porcine hemagglutinating encephalomyelitis virus	VW572	spike

NC_022643	27869-28525	Betacoronavirus Erinaceus/VMC/DEU/2012	ErinaceusCoV/2012-216/GER/2012	membrane

NC_022643	28580-29854	Betacoronavirus Erinaceus/VMC/DEU/2012	ErinaceusCoV/2012-216/GER/2012	nucleocapsid

NC_022643	21637-25629	Betacoronavirus Erinaceus/VMC/DEU/2012	ErinaceusCoV/2012-216/GER/2012	spike

NC_022643	241-21719	Betacoronavirus Erinaceus/VMC/DEU/2012	ErinaceusCoV/2012-216/GER/2012	orf1ab

NC_026011	28908-29603	Betacoronavirus HKU24	HKU24-R05005I	membrane

NC_026011	29613-30944	Betacoronavirus HKU24	HKU24-R05005I	nucleocapsid

NC_026011	213-21637	Betacoronavirus HKU24	HKU24-R05005I	orf1ab

NC_026011	23777-27853	Betacoronavirus HKU24	HKU24-R05005I	spike

KM349743	28908-29603	Betacoronavirus HKU24	HKU24-R05009I	membrane

KM349743	29613-30944	Betacoronavirus HKU24	HKU24-R05009I	nucleocapsid

KM349743	213-21637	Betacoronavirus HKU24	HKU24-R05009I	orf1ab

KM349743	23777-27853	Betacoronavirus HKU24	HKU24-R05009I	spike

KM349744	28908-29603	Betacoronavirus HKU24	HKU24-R05010I	membrane

KM349744	29613-30944	Betacoronavirus HKU24	HKU24-R05010I	nucleocapsid

KM349744	213-21637	Betacoronavirus HKU24	HKU24-R05010I	orf1ab

KM349744	23777-27853	Betacoronavirus HKU24	HKU24-R05010I	spike

KT779555	27594-28265	Human coronavirus HKU1	BJ01-p3	membrane

KT779555	28281-29606	Human coronavirus HKU1	BJ01-p3	nucleocapsid

KT779555	200-21714	Human coronavirus HKU1	BJ01-p3	orf1ab

KT779555	22903-26973	Human coronavirus HKU1	BJ01-p3	spike

KT779556	27594-28265	Human coronavirus HKU1	BJ01-p9	membrane

KT779556	28281-29606	Human coronavirus HKU1	BJ01-p9	nucleocapsid

KT779556	200-21714	Human coronavirus HKU1	BJ01-p9	orf1ab

KT779556	22903-26973	Human coronavirus HKU1	BJ01-p9	spike

HM034837	27597-28268	Human coronavirus HKU1	Caen1	membrane

HM034837	28284-29609	Human coronavirus HKU1	Caen1	nucleocapsid

HM034837	170-21717	Human coronavirus HKU1	Caen1	orf1ab

HM034837	22906-26976	Human coronavirus HKU1	Caen1	spike

NC_006577	27633-28304	Human coronavirus HKU1	HKU1	membrane

NC_006577	28320-29645	Human coronavirus HKU1	HKU1	nucleocapsid

NC_006577	206-21753	Human coronavirus HKU1	HKU1	orf1ab

NC_006577	22942-27012	Human coronavirus HKU1	HKU1	spike

KF686341	27563-28234	Human coronavirus HKU1	HKU1/human/USA/HKU1-10/2010	membrane

KF686341	28250-29575	Human coronavirus HKU1	HKU1/human/USA/HKU1-10/2010	nucleocapsid

KF686341	181-21683	Human coronavirus HKU1	HKU1/human/USA/HKU1-10/2010	orf1ab

KF686341	22872-26942	Human coronavirus HKU1	HKU1/human/USA/HKU1-10/2010	spike

KF686342	27473-28144	Human coronavirus HKU1	HKU1/human/USA/HKU1-11/2009	membrane

KF686342	28160-29485	Human coronavirus HKU1	HKU1/human/USA/HKU1-11/2009	nucleocapsid

KF686342	181-21593	Human coronavirus HKU1	HKU1/human/USA/HKU1-11/2009	orf1ab

KF686342	22782-26852	Human coronavirus HKU1	HKU1/human/USA/HKU1-11/2009	spike

KF686346	27713-28384	Human coronavirus HKU1	HKU1/human/USA/HKU1-12/2010	membrane

KF686346	28400-29725	Human coronavirus HKU1	HKU1/human/USA/HKU1-12/2010	nucleocapsid

KF686346	181-21833	Human coronavirus HKU1	HKU1/human/USA/HKU1-12/2010	orf1ab

KF686346	23022-27092	Human coronavirus HKU1	HKU1/human/USA/HKU1-12/2010	spike

KF686343	27713-28384	Human coronavirus HKU1	HKU1/human/USA/HKU1-13/2010	membrane

KF686343	28400-29725	Human coronavirus HKU1	HKU1/human/USA/HKU1-13/2010	nucleocapsid

KF686343	181-21833	Human coronavirus HKU1	HKU1/human/USA/HKU1-13/2010	orf1ab

KF686343	23022-27092	Human coronavirus HKU1	HKU1/human/USA/HKU1-13/2010	spike

KF686344	27425-28096	Human coronavirus HKU1	HKU1/human/USA/HKU1-15/2009	membrane

KF686344	28112-29437	Human coronavirus HKU1	HKU1/human/USA/HKU1-15/2009	nucleocapsid

KF686344	181-21545	Human coronavirus HKU1	HKU1/human/USA/HKU1-15/2009	orf1ab

KF686344	22734-26804	Human coronavirus HKU1	HKU1/human/USA/HKU1-15/2009	spike

KF430201	27665-28336	Human coronavirus HKU1	HKU1/human/USA/HKU1-18/2010	membrane

KF430201	28352-29677	Human coronavirus HKU1	HKU1/human/USA/HKU1-18/2010	nucleocapsid

KF430201	196-21788	Human coronavirus HKU1	HKU1/human/USA/HKU1-18/2010	orf1ab

KF430201	22977-27044	Human coronavirus HKU1	HKU1/human/USA/HKU1-18/2010	spike

KF686340	27653-28324	Human coronavirus HKU1	HKU1/human/USA/HKU1-5/2009	membrane

KF686340	28340-29665	Human coronavirus HKU1	HKU1/human/USA/HKU1-5/2009	nucleocapsid

KF686340	181-21773	Human coronavirus HKU1	HKU1/human/USA/HKU1-5/2009	orf1ab

KF686340	22962-27032	Human coronavirus HKU1	HKU1/human/USA/HKU1-5/2009	spike

KY674921	NA	Human coronavirus HKU1	N08-87	membrane

KY674921	NA	Human coronavirus HKU1	N08-87	nucleocapsid

KY674921	NA	Human coronavirus HKU1	N08-87	orf1ab

KY674921	NA	Human coronavirus HKU1	N08-87	spike

KY674943	NA	Human coronavirus HKU1	N09-1605B	membrane

KY674943	NA	Human coronavirus HKU1	N09-1605B	nucleocapsid

KY674943	NA	Human coronavirus HKU1	N09-1605B	orf1ab

KY674943	NA	Human coronavirus HKU1	N09-1605B	spike

KY674942	NA	Human coronavirus HKU1	N09-1627B	membrane

KY674942	NA	Human coronavirus HKU1	N09-1627B	nucleocapsid

KY674942	NA	Human coronavirus HKU1	N09-1627B	orf1ab

KY674942	NA	Human coronavirus HKU1	N09-1627B	spike

KY674941	NA	Human coronavirus HKU1	N09-1663B	membrane

KY674941	NA	Human coronavirus HKU1	N09-1663B	nucleocapsid

KY674941	NA	Human coronavirus HKU1	N09-1663B	orf1ab

KY674941	NA	Human coronavirus HKU1	N09-1663B	spike

MK167038	NA	Human coronavirus HKU1	SC2521	membrane

MK167038	NA	Human coronavirus HKU1	SC2521	nucleocapsid

MK167038	NA	Human coronavirus HKU1	SC2521	orf1ab

MK167038	NA	Human coronavirus HKU1	SC2521	spike

MH940245	27518-28189	Human coronavirus HKU1	SI17244	membrane

MH940245	28205-29530	Human coronavirus HKU1	SI17244	nucleocapsid

MH940245	196-21653	Human coronavirus HKU1	SI17244	orf1ab

MH940245	22858-26913	Human coronavirus HKU1	SI17244	spike

KC776174	27807-28466	Human betacoronavirus 2c Jordan-N3/2012	Jordan-N3/2012	membrane

KC776174	28520-29761	Human betacoronavirus 2c Jordan-N3/2012	Jordan-N3/2012	nucleocapsid

KC776174	233-21468	Human betacoronavirus 2c Jordan-N3/2012	Jordan-N3/2012	orf1ab

KC776174	21410-25471	Human betacoronavirus 2c Jordan-N3/2012	Jordan-N3/2012	spike

KC869678	27840-28499	Coronavirus Neoromicia/PML-PHE1/RSA/2011	Neoromicia/PML-PHE1/RSA/2011	membrane

KC869678	28557-29801	Coronavirus Neoromicia/PML-PHE1/RSA/2011	Neoromicia/PML-PHE1/RSA/2011	nucleocapsid

KC869678	281-21528	Coronavirus Neoromicia/PML-PHE1/RSA/2011	Neoromicia/PML-PHE1/RSA/2011	orf1ab

KC869678	21470-25504	Coronavirus Neoromicia/PML-PHE1/RSA/2011	Neoromicia/PML-PHE1/RSA/2011	spike

KC667074	NA	Human betacoronavirus 2c England-Qatar/2012	England/Qatar/2012	membrane

KC667074	NA	Human betacoronavirus 2c England-Qatar/2012	England/Qatar/2012	nucleocapsid

KC667074	NA	Human betacoronavirus 2c England-Qatar/2012	England/Qatar/2012	orf1ab

KC667074	NA	Human betacoronavirus 2c England-Qatar/2012	England/Qatar/2012	spike

KJ614529	27838-28497	Human betacoronavirus 2c Jordan-N3/2012	Jordan-N3/2012	membrane

KJ614529	28551-29792	Human betacoronavirus 2c Jordan-N3/2012	Jordan-N3/2012	nucleocapsid

KJ614529	264-21499	Human betacoronavirus 2c Jordan-N3/2012	Jordan-N3/2012	orf1ab

KJ614529	21441-25502	Human betacoronavirus 2c Jordan-N3/2012	Jordan-N3/2012	spike

KP209310	27853-28512	Middle East respiratory syndrome coronavirus	Abu Dhabi/Gayathi_UAE_2_2014	membrane

KP209310	28566-29807	Middle East respiratory syndrome coronavirus	Abu Dhabi/Gayathi_UAE_2_2014	nucleocapsid

KP209310	279-21514	Middle East respiratory syndrome coronavirus	Abu Dhabi/Gayathi_UAE_2_2014	orf1ab

KP209310	21456-25517	Middle East respiratory syndrome coronavirus	Abu Dhabi/Gayathi_UAE_2_2014	spike

KP209308	27853-28512	Middle East respiratory syndrome coronavirus	Abu Dhabi_UAE_16_2014	membrane

KP209308	28566-29807	Middle East respiratory syndrome coronavirus	Abu Dhabi_UAE_16_2014	nucleocapsid

KP209308	279-21514	Middle East respiratory syndrome coronavirus	Abu Dhabi_UAE_16_2014	orf1ab

KP209308	21456-25517	Middle East respiratory syndrome coronavirus	Abu Dhabi_UAE_16_2014	spike

KP209307	27853-28512	Middle East respiratory syndrome coronavirus	Abu Dhabi_UAE_18_2014	membrane

KP209307	28566-29807	Middle East respiratory syndrome coronavirus	Abu Dhabi_UAE_18_2014	nucleocapsid

KP209307	279-21514	Middle East respiratory syndrome coronavirus	Abu Dhabi_UAE_18_2014	orf1ab

KP209307	21456-25517	Middle East respiratory syndrome coronavirus	Abu Dhabi_UAE_18_2014	spike

KP209313	27853-28512	Middle East respiratory syndrome coronavirus	Abu Dhabi_UAE_26_2014	membrane

KP209313	28566-29807	Middle East respiratory syndrome coronavirus	Abu Dhabi_UAE_26_2014	nucleocapsid

KP209313	279-21514	Middle East respiratory syndrome coronavirus	Abu Dhabi_UAE_26_2014	orf1ab

KP209313	21456-25517	Middle East respiratory syndrome coronavirus	Abu Dhabi_UAE_26_2014	spike

KP209306	27853-28512	Middle East respiratory syndrome coronavirus	Abu Dhabi_UAE_8_2014	membrane

KP209306	28566-29807	Middle East respiratory syndrome coronavirus	Abu Dhabi_UAE_8_2014	nucleocapsid

KP209306	279-21514	Middle East respiratory syndrome coronavirus	Abu Dhabi_UAE_8_2014	orf1ab

KP209306	21456-25517	Middle East respiratory syndrome coronavirus	Abu Dhabi_UAE_8_2014	spike

KP209312	27853-28512	Middle East respiratory syndrome coronavirus	Abu Dhabi_UAE_9_2013	membrane

KP209312	28566-29807	Middle East respiratory syndrome coronavirus	Abu Dhabi_UAE_9_2013	nucleocapsid

KP209312	279-21514	Middle East respiratory syndrome coronavirus	Abu Dhabi_UAE_9_2013	orf1ab

KP209312	21456-25517	Middle East respiratory syndrome coronavirus	Abu Dhabi_UAE_9_2013	spike

KF600627	NA	Middle East respiratory syndrome coronavirus	Al-Hasa_12_2013	membrane

KF600627	NA	Middle East respiratory syndrome coronavirus	Al-Hasa_12_2013	nucleocapsid

KF600627	NA	Middle East respiratory syndrome coronavirus	Al-Hasa_12_2013	orf1ab

KF600627	NA	Middle East respiratory syndrome coronavirus	Al-Hasa_12_2013	spike

KF600645	NA	Middle East respiratory syndrome coronavirus	Al-Hasa_15_2013	membrane

KF600645	NA	Middle East respiratory syndrome coronavirus	Al-Hasa_15_2013	nucleocapsid

KF600645	NA	Middle East respiratory syndrome coronavirus	Al-Hasa_15_2013	orf1ab

KF600645	NA	Middle East respiratory syndrome coronavirus	Al-Hasa_15_2013	spike

KF600644	NA	Middle East respiratory syndrome coronavirus	Al-Hasa_16_2013	membrane

KF600644	NA	Middle East respiratory syndrome coronavirus	Al-Hasa_16_2013	nucleocapsid

KF600644	NA	Middle East respiratory syndrome coronavirus	Al-Hasa_16_2013	orf1ab

KF600644	NA	Middle East respiratory syndrome coronavirus	Al-Hasa_16_2013	spike

KF600647	NA	Middle East respiratory syndrome coronavirus	Al-Hasa_17_2013	membrane

KF600647	NA	Middle East respiratory syndrome coronavirus	Al-Hasa_17_2013	nucleocapsid

KF600647	NA	Middle East respiratory syndrome coronavirus	Al-Hasa_17_2013	orf1ab

KF600647	NA	Middle East respiratory syndrome coronavirus	Al-Hasa_17_2013	spike

KF600651	NA	Middle East respiratory syndrome coronavirus	Al-Hasa_18_2013	membrane

KF600651	NA	Middle East respiratory syndrome coronavirus	Al-Hasa_18_2013	nucleocapsid

KF600651	NA	Middle East respiratory syndrome coronavirus	Al-Hasa_18_2013	orf1ab

KF600651	NA	Middle East respiratory syndrome coronavirus	Al-Hasa_18_2013	spike

KF600632	NA	Middle East respiratory syndrome coronavirus	Al-Hasa_19_2013	membrane

KF600632	NA	Middle East respiratory syndrome coronavirus	Al-Hasa_19_2013	nucleocapsid

KF600632	NA	Middle East respiratory syndrome coronavirus	Al-Hasa_19_2013	orf1ab

KF600632	NA	Middle East respiratory syndrome coronavirus	Al-Hasa_19_2013	spike

KF186567	NA	Middle East respiratory syndrome coronavirus	Al-Hasa_1_2013	membrane

KF186567	NA	Middle East respiratory syndrome coronavirus	Al-Hasa_1_2013	nucleocapsid

KF186567	NA	Middle East respiratory syndrome coronavirus	Al-Hasa_1_2013	orf1ab

KF186567	NA	Middle East respiratory syndrome coronavirus	Al-Hasa_1_2013	spike

KF600634	NA	Middle East respiratory syndrome coronavirus	Al-Hasa_21_2013	membrane

KF600634	NA	Middle East respiratory syndrome coronavirus	Al-Hasa_21_2013	nucleocapsid

KF600634	NA	Middle East respiratory syndrome coronavirus	Al-Hasa_21_2013	orf1ab

KF600634	NA	Middle East respiratory syndrome coronavirus	Al-Hasa_21_2013	spike

KJ156866	NA	Middle East respiratory syndrome coronavirus	Al-Hasa_25_2013	membrane

KJ156866	NA	Middle East respiratory syndrome coronavirus	Al-Hasa_25_2013	nucleocapsid

KJ156866	NA	Middle East respiratory syndrome coronavirus	Al-Hasa_25_2013	orf1ab

KJ156866	NA	Middle East respiratory syndrome coronavirus	Al-Hasa_25_2013	spike

KF186566	NA	Middle East respiratory syndrome coronavirus	Al-Hasa_2_2013	membrane

KF186566	NA	Middle East respiratory syndrome coronavirus	Al-Hasa_2_2013	nucleocapsid

KF186566	NA	Middle East respiratory syndrome coronavirus	Al-Hasa_2_2013	orf1ab

KF186566	NA	Middle East respiratory syndrome coronavirus	Al-Hasa_2_2013	spike

KF186565	NA	Middle East respiratory syndrome coronavirus	Al-Hasa_3_2013	membrane

KF186565	NA	Middle East respiratory syndrome coronavirus	Al-Hasa_3_2013	nucleocapsid

KF186565	NA	Middle East respiratory syndrome coronavirus	Al-Hasa_3_2013	orf1ab

KF186565	NA	Middle East respiratory syndrome coronavirus	Al-Hasa_3_2013	spike

KF186564	NA	Middle East respiratory syndrome coronavirus	Al-Hasa_4_2013	membrane

KF186564	NA	Middle East respiratory syndrome coronavirus	Al-Hasa_4_2013	nucleocapsid

KF186564	NA	Middle East respiratory syndrome coronavirus	Al-Hasa_4_2013	orf1ab

KF186564	NA	Middle East respiratory syndrome coronavirus	Al-Hasa_4_2013	spike

KF600620	NA	Middle East respiratory syndrome coronavirus	Bisha_1_2012	membrane

KF600620	NA	Middle East respiratory syndrome coronavirus	Bisha_1_2012	nucleocapsid

KF600620	NA	Middle East respiratory syndrome coronavirus	Bisha_1_2012	orf1ab

KF600620	NA	Middle East respiratory syndrome coronavirus	Bisha_1_2012	spike

KF600630	NA	Middle East respiratory syndrome coronavirus	Buraidah_1_2013	membrane

KF600630	NA	Middle East respiratory syndrome coronavirus	Buraidah_1_2013	nucleocapsid

KF600630	NA	Middle East respiratory syndrome coronavirus	Buraidah_1_2013	orf1ab

KF600630	NA	Middle East respiratory syndrome coronavirus	Buraidah_1_2013	spike

KT368827	27853-28512	Middle East respiratory syndrome coronavirus	camel/Jeddah/401/2014	membrane

KT368827	28566-29807	Middle East respiratory syndrome coronavirus	camel/Jeddah/401/2014	nucleocapsid

KT368827	279-21514	Middle East respiratory syndrome coronavirus	camel/Jeddah/401/2014	orf1ab

KT368827	21456-25517	Middle East respiratory syndrome coronavirus	camel/Jeddah/401/2014	spike

KT368828	27853-28512	Middle East respiratory syndrome coronavirus	camel/Jeddah/D100/2014	membrane

KT368828	28566-29807	Middle East respiratory syndrome coronavirus	camel/Jeddah/D100/2014	nucleocapsid

KT368828	279-21514	Middle East respiratory syndrome coronavirus	camel/Jeddah/D100/2014	orf1ab

KT368828	21456-25517	Middle East respiratory syndrome coronavirus	camel/Jeddah/D100/2014	spike

KT368829	27853-28512	Middle East respiratory syndrome coronavirus	camel/Jeddah/D33(b)/2014	membrane

KT368829	28566-29807	Middle East respiratory syndrome coronavirus	camel/Jeddah/D33(b)/2014	nucleocapsid

KT368829	279-21514	Middle East respiratory syndrome coronavirus	camel/Jeddah/D33(b)/2014	orf1ab

KT368829	21456-25517	Middle East respiratory syndrome coronavirus	camel/Jeddah/D33(b)/2014	spike

KT368830	27853-28512	Middle East respiratory syndrome coronavirus	camel/Jeddah/D34/2014	membrane

KT368830	28566-29807	Middle East respiratory syndrome coronavirus	camel/Jeddah/D34/2014	nucleocapsid

KT368830	279-21514	Middle East respiratory syndrome coronavirus	camel/Jeddah/D34/2014	orf1ab

KT368830	21456-25517	Middle East respiratory syndrome coronavirus	camel/Jeddah/D34/2014	spike

KT368831	27853-28512	Middle East respiratory syndrome coronavirus	camel/Jeddah/D35/2014	membrane

KT368831	28566-29807	Middle East respiratory syndrome coronavirus	camel/Jeddah/D35/2014	nucleocapsid

KT368831	279-21514	Middle East respiratory syndrome coronavirus	camel/Jeddah/D35/2014	orf1ab

KT368831	21456-25517	Middle East respiratory syndrome coronavirus	camel/Jeddah/D35/2014	spike

KT368833	27853-28512	Middle East respiratory syndrome coronavirus	camel/Jeddah/D38(b)/2014	membrane

KT368833	28566-29807	Middle East respiratory syndrome coronavirus	camel/Jeddah/D38(b)/2014	nucleocapsid

KT368833	279-21514	Middle East respiratory syndrome coronavirus	camel/Jeddah/D38(b)/2014	orf1ab

KT368833	21456-25517	Middle East respiratory syndrome coronavirus	camel/Jeddah/D38(b)/2014	spike

KT368834	27853-28512	Middle East respiratory syndrome coronavirus	camel/Jeddah/D40/2014	membrane

KT368834	28566-29807	Middle East respiratory syndrome coronavirus	camel/Jeddah/D40/2014	nucleocapsid

KT368834	279-21514	Middle East respiratory syndrome coronavirus	camel/Jeddah/D40/2014	orf1ab

KT368834	21456-25517	Middle East respiratory syndrome coronavirus	camel/Jeddah/D40/2014	spike

KT368835	27853-28512	Middle East respiratory syndrome coronavirus	camel/Jeddah/D42/2014	membrane

KT368835	28566-29807	Middle East respiratory syndrome coronavirus	camel/Jeddah/D42/2014	nucleocapsid

KT368835	279-21514	Middle East respiratory syndrome coronavirus	camel/Jeddah/D42/2014	orf1ab

KT368835	21456-25517	Middle East respiratory syndrome coronavirus	camel/Jeddah/D42/2014	spike

KT368836	27853-28512	Middle East respiratory syndrome coronavirus	camel/Jeddah/D43(b)/2014	membrane

KT368836	28566-29807	Middle East respiratory syndrome coronavirus	camel/Jeddah/D43(b)/2014	nucleocapsid

KT368836	279-21514	Middle East respiratory syndrome coronavirus	camel/Jeddah/D43(b)/2014	orf1ab

KT368836	21456-25517	Middle East respiratory syndrome coronavirus	camel/Jeddah/D43(b)/2014	spike

KT368838	27853-28512	Middle East respiratory syndrome coronavirus	camel/Jeddah/D46(b)/2014	membrane

KT368838	28566-29807	Middle East respiratory syndrome coronavirus	camel/Jeddah/D46(b)/2014	nucleocapsid

KT368838	279-21514	Middle East respiratory syndrome coronavirus	camel/Jeddah/D46(b)/2014	orf1ab

KT368838	21456-25517	Middle East respiratory syndrome coronavirus	camel/Jeddah/D46(b)/2014	spike

KT368839	27853-28512	Middle East respiratory syndrome coronavirus	camel/Jeddah/D47/2014	membrane

KT368839	28566-29807	Middle East respiratory syndrome coronavirus	camel/Jeddah/D47/2014	nucleocapsid

KT368839	279-21514	Middle East respiratory syndrome coronavirus	camel/Jeddah/D47/2014	orf1ab

KT368839	21456-25517	Middle East respiratory syndrome coronavirus	camel/Jeddah/D47/2014	spike

KT368840	27853-28512	Middle East respiratory syndrome coronavirus	camel/Jeddah/D48/2014	membrane

KT368840	28566-29807	Middle East respiratory syndrome coronavirus	camel/Jeddah/D48/2014	nucleocapsid

KT368840	279-21514	Middle East respiratory syndrome coronavirus	camel/Jeddah/D48/2014	orf1ab

KT368840	21456-25517	Middle East respiratory syndrome coronavirus	camel/Jeddah/D48/2014	spike

KT368841	27853-28512	Middle East respiratory syndrome coronavirus	camel/Jeddah/D49/2014	membrane

KT368841	28566-29807	Middle East respiratory syndrome coronavirus	camel/Jeddah/D49/2014	nucleocapsid

KT368841	279-21514	Middle East respiratory syndrome coronavirus	camel/Jeddah/D49/2014	orf1ab

KT368841	21456-25517	Middle East respiratory syndrome coronavirus	camel/Jeddah/D49/2014	spike

KT368842	27853-28512	Middle East respiratory syndrome coronavirus	camel/Jeddah/D50(b)/2014	membrane

KT368842	28566-29807	Middle East respiratory syndrome coronavirus	camel/Jeddah/D50(b)/2014	nucleocapsid

KT368842	279-21514	Middle East respiratory syndrome coronavirus	camel/Jeddah/D50(b)/2014	orf1ab

KT368842	21456-25517	Middle East respiratory syndrome coronavirus	camel/Jeddah/D50(b)/2014	spike

KT368843	27853-28512	Middle East respiratory syndrome coronavirus	camel/Jeddah/D88/2014	membrane

KT368843	28566-29807	Middle East respiratory syndrome coronavirus	camel/Jeddah/D88/2014	nucleocapsid

KT368843	279-21514	Middle East respiratory syndrome coronavirus	camel/Jeddah/D88/2014	orf1ab

KT368843	21456-25517	Middle East respiratory syndrome coronavirus	camel/Jeddah/D88/2014	spike

KT368844	27853-28512	Middle East respiratory syndrome coronavirus	camel/Jeddah/D90/2014	membrane

KT368844	28566-29807	Middle East respiratory syndrome coronavirus	camel/Jeddah/D90/2014	nucleocapsid

KT368844	279-21514	Middle East respiratory syndrome coronavirus	camel/Jeddah/D90/2014	orf1ab

KT368844	21456-25517	Middle East respiratory syndrome coronavirus	camel/Jeddah/D90/2014	spike

KT368845	27853-28512	Middle East respiratory syndrome coronavirus	camel/Jeddah/D92/2014	membrane

KT368845	28566-29807	Middle East respiratory syndrome coronavirus	camel/Jeddah/D92/2014	nucleocapsid

KT368845	279-21514	Middle East respiratory syndrome coronavirus	camel/Jeddah/D92/2014	orf1ab

KT368845	21456-25517	Middle East respiratory syndrome coronavirus	camel/Jeddah/D92/2014	spike

KT368824	27853-28512	Middle East respiratory syndrome coronavirus	camel/Jeddah/F13A/2014	membrane

KT368824	28566-29807	Middle East respiratory syndrome coronavirus	camel/Jeddah/F13A/2014	nucleocapsid

KT368824	279-21514	Middle East respiratory syndrome coronavirus	camel/Jeddah/F13A/2014	orf1ab

KT368824	21456-25517	Middle East respiratory syndrome coronavirus	camel/Jeddah/F13A/2014	spike

KT368858	27853-28512	Middle East respiratory syndrome coronavirus	camel/Jeddah/Jd1(b)/2015	membrane

KT368858	28566-29807	Middle East respiratory syndrome coronavirus	camel/Jeddah/Jd1(b)/2015	nucleocapsid

KT368858	279-21514	Middle East respiratory syndrome coronavirus	camel/Jeddah/Jd1(b)/2015	orf1ab

KT368858	21456-25517	Middle East respiratory syndrome coronavirus	camel/Jeddah/Jd1(b)/2015	spike

KT368866	27853-28512	Middle East respiratory syndrome coronavirus	camel/Jeddah/Jd175/2015	membrane

KT368866	28566-29807	Middle East respiratory syndrome coronavirus	camel/Jeddah/Jd175/2015	nucleocapsid

KT368866	279-21514	Middle East respiratory syndrome coronavirus	camel/Jeddah/Jd175/2015	orf1ab

KT368866	21456-25517	Middle East respiratory syndrome coronavirus	camel/Jeddah/Jd175/2015	spike

KT368867	27853-28512	Middle East respiratory syndrome coronavirus	camel/Jeddah/Jd199/2015	membrane

KT368867	28566-29807	Middle East respiratory syndrome coronavirus	camel/Jeddah/Jd199/2015	nucleocapsid

KT368867	279-21514	Middle East respiratory syndrome coronavirus	camel/Jeddah/Jd199/2015	orf1ab

KT368867	21456-25517	Middle East respiratory syndrome coronavirus	camel/Jeddah/Jd199/2015	spike

KT368859	27853-28512	Middle East respiratory syndrome coronavirus	camel/Jeddah/Jd4/2015	membrane

KT368859	28566-29807	Middle East respiratory syndrome coronavirus	camel/Jeddah/Jd4/2015	nucleocapsid

KT368859	279-21514	Middle East respiratory syndrome coronavirus	camel/Jeddah/Jd4/2015	orf1ab

KT368859	21456-25517	Middle East respiratory syndrome coronavirus	camel/Jeddah/Jd4/2015	spike

KT368860	27853-28512	Middle East respiratory syndrome coronavirus	camel/Jeddah/Jd6(b)/2015	membrane

KT368860	28566-29807	Middle East respiratory syndrome coronavirus	camel/Jeddah/Jd6(b)/2015	nucleocapsid

KT368860	279-21514	Middle East respiratory syndrome coronavirus	camel/Jeddah/Jd6(b)/2015	orf1ab

KT368860	21456-25517	Middle East respiratory syndrome coronavirus	camel/Jeddah/Jd6(b)/2015	spike

KT368861	27853-28512	Middle East respiratory syndrome coronavirus	camel/Jeddah/Jd7/2015	membrane

KT368861	28566-29807	Middle East respiratory syndrome coronavirus	camel/Jeddah/Jd7/2015	nucleocapsid

KT368861	279-21514	Middle East respiratory syndrome coronavirus	camel/Jeddah/Jd7/2015	orf1ab

KT368861	21456-25517	Middle East respiratory syndrome coronavirus	camel/Jeddah/Jd7/2015	spike

KT368862	27853-28512	Middle East respiratory syndrome coronavirus	camel/Jeddah/Jd85/2015	membrane

KT368862	28566-29807	Middle East respiratory syndrome coronavirus	camel/Jeddah/Jd85/2015	nucleocapsid

KT368862	279-21514	Middle East respiratory syndrome coronavirus	camel/Jeddah/Jd85/2015	orf1ab

KT368862	21456-25517	Middle East respiratory syndrome coronavirus	camel/Jeddah/Jd85/2015	spike

KT368864	27853-28512	Middle East respiratory syndrome coronavirus	camel/Jeddah/Jd87/2015	membrane

KT368864	28566-29807	Middle East respiratory syndrome coronavirus	camel/Jeddah/Jd87/2015	nucleocapsid

KT368864	279-21514	Middle East respiratory syndrome coronavirus	camel/Jeddah/Jd87/2015	orf1ab

KT368864	21456-25517	Middle East respiratory syndrome coronavirus	camel/Jeddah/Jd87/2015	spike

KT368865	27853-28512	Middle East respiratory syndrome coronavirus	camel/Jeddah/Jd90/2015	membrane

KT368865	28566-29807	Middle East respiratory syndrome coronavirus	camel/Jeddah/Jd90/2015	nucleocapsid

KT368865	279-21514	Middle East respiratory syndrome coronavirus	camel/Jeddah/Jd90/2015	orf1ab

KT368865	21456-25517	Middle East respiratory syndrome coronavirus	camel/Jeddah/Jd90/2015	spike

KT368846	27853-28512	Middle East respiratory syndrome coronavirus	camel/Jeddah/N51/2014	membrane

KT368846	28566-29807	Middle East respiratory syndrome coronavirus	camel/Jeddah/N51/2014	nucleocapsid

KT368846	279-21514	Middle East respiratory syndrome coronavirus	camel/Jeddah/N51/2014	orf1ab

KT368846	21456-25517	Middle East respiratory syndrome coronavirus	camel/Jeddah/N51/2014	spike

KT368847	27853-28512	Middle East respiratory syndrome coronavirus	camel/Jeddah/N62(b)/2014	membrane

KT368847	28566-29807	Middle East respiratory syndrome coronavirus	camel/Jeddah/N62(b)/2014	nucleocapsid

KT368847	279-21514	Middle East respiratory syndrome coronavirus	camel/Jeddah/N62(b)/2014	orf1ab

KT368847	21456-25517	Middle East respiratory syndrome coronavirus	camel/Jeddah/N62(b)/2014	spike

KT368848	27853-28512	Middle East respiratory syndrome coronavirus	camel/Jeddah/N68(b)/2014	membrane

KT368848	28566-29807	Middle East respiratory syndrome coronavirus	camel/Jeddah/N68(b)/2014	nucleocapsid

KT368848	279-21514	Middle East respiratory syndrome coronavirus	camel/Jeddah/N68(b)/2014	orf1ab

KT368848	21456-25517	Middle East respiratory syndrome coronavirus	camel/Jeddah/N68(b)/2014	spike

KT368849	27853-28512	Middle East respiratory syndrome coronavirus	camel/Jeddah/O23(b)/2014	membrane

KT368849	28566-29807	Middle East respiratory syndrome coronavirus	camel/Jeddah/O23(b)/2014	nucleocapsid

KT368849	279-21514	Middle East respiratory syndrome coronavirus	camel/Jeddah/O23(b)/2014	orf1ab

KT368849	21456-25517	Middle East respiratory syndrome coronavirus	camel/Jeddah/O23(b)/2014	spike

KT368850	27853-28512	Middle East respiratory syndrome coronavirus	camel/Jeddah/O24/2014	membrane

KT368850	28566-29807	Middle East respiratory syndrome coronavirus	camel/Jeddah/O24/2014	nucleocapsid

KT368850	279-21514	Middle East respiratory syndrome coronavirus	camel/Jeddah/O24/2014	orf1ab

KT368850	21456-25517	Middle East respiratory syndrome coronavirus	camel/Jeddah/O24/2014	spike

KT368851	27853-28512	Middle East respiratory syndrome coronavirus	camel/Jeddah/O30/2014	membrane

KT368851	28566-29807	Middle East respiratory syndrome coronavirus	camel/Jeddah/O30/2014	nucleocapsid

KT368851	279-21514	Middle East respiratory syndrome coronavirus	camel/Jeddah/O30/2014	orf1ab

KT368851	21456-25517	Middle East respiratory syndrome coronavirus	camel/Jeddah/O30/2014	spike

KT368852	27853-28512	Middle East respiratory syndrome coronavirus	camel/Jeddah/O47(b)/2014	membrane

KT368852	28566-29807	Middle East respiratory syndrome coronavirus	camel/Jeddah/O47(b)/2014	nucleocapsid

KT368852	279-21514	Middle East respiratory syndrome coronavirus	camel/Jeddah/O47(b)/2014	orf1ab

KT368852	21456-25517	Middle East respiratory syndrome coronavirus	camel/Jeddah/O47(b)/2014	spike

KT368853	27853-28512	Middle East respiratory syndrome coronavirus	camel/Jeddah/S100/2014	membrane

KT368853	28566-29807	Middle East respiratory syndrome coronavirus	camel/Jeddah/S100/2014	nucleocapsid

KT368853	279-21514	Middle East respiratory syndrome coronavirus	camel/Jeddah/S100/2014	orf1ab

KT368853	21456-25517	Middle East respiratory syndrome coronavirus	camel/Jeddah/S100/2014	spike

KT368854	27853-28512	Middle East respiratory syndrome coronavirus	camel/Jeddah/S73/2014	membrane

KT368854	28566-29807	Middle East respiratory syndrome coronavirus	camel/Jeddah/S73/2014	nucleocapsid

KT368854	279-21514	Middle East respiratory syndrome coronavirus	camel/Jeddah/S73/2014	orf1ab

KT368854	21456-25517	Middle East respiratory syndrome coronavirus	camel/Jeddah/S73/2014	spike

KT368855	27853-28512	Middle East respiratory syndrome coronavirus	camel/Jeddah/S93/2014	membrane

KT368855	28566-29807	Middle East respiratory syndrome coronavirus	camel/Jeddah/S93/2014	nucleocapsid

KT368855	279-21514	Middle East respiratory syndrome coronavirus	camel/Jeddah/S93/2014	orf1ab

KT368855	21456-25517	Middle East respiratory syndrome coronavirus	camel/Jeddah/S93/2014	spike

KT368856	27853-28512	Middle East respiratory syndrome coronavirus	camel/Jeddah/S94/2014	membrane

KT368856	28566-29807	Middle East respiratory syndrome coronavirus	camel/Jeddah/S94/2014	nucleocapsid

KT368856	279-21514	Middle East respiratory syndrome coronavirus	camel/Jeddah/S94/2014	orf1ab

KT368856	21456-25517	Middle East respiratory syndrome coronavirus	camel/Jeddah/S94/2014	spike

KT368857	27853-28512	Middle East respiratory syndrome coronavirus	camel/Jeddah/S99/2014	membrane

KT368857	28566-29807	Middle East respiratory syndrome coronavirus	camel/Jeddah/S99/2014	nucleocapsid

KT368857	279-21514	Middle East respiratory syndrome coronavirus	camel/Jeddah/S99/2014	orf1ab

KT368857	21456-25517	Middle East respiratory syndrome coronavirus	camel/Jeddah/S99/2014	spike

KJ650098	NA	Middle East respiratory syndrome coronavirus	Camel/Qatar_2_2014	orf1ab

KJ650098	NA	Middle East respiratory syndrome coronavirus	Camel/Qatar_2_2014	membrane

KJ650098	NA	Middle East respiratory syndrome coronavirus	Camel/Qatar_2_2014	nucleocapsid

KJ650098	NA	Middle East respiratory syndrome coronavirus	Camel/Qatar_2_2014	spike

KT368868	27853-28512	Middle East respiratory syndrome coronavirus	camel/Riyadh/Ry136/2015	membrane

KT368868	28566-29807	Middle East respiratory syndrome coronavirus	camel/Riyadh/Ry136/2015	nucleocapsid

KT368868	279-21514	Middle East respiratory syndrome coronavirus	camel/Riyadh/Ry136/2015	orf1ab

KT368868	21456-25517	Middle East respiratory syndrome coronavirus	camel/Riyadh/Ry136/2015	spike

KT368869	27842-28501	Middle East respiratory syndrome coronavirus	camel/Riyadh/Ry137/2015	membrane

KT368869	28555-29796	Middle East respiratory syndrome coronavirus	camel/Riyadh/Ry137/2015	nucleocapsid

KT368869	268-21503	Middle East respiratory syndrome coronavirus	camel/Riyadh/Ry137/2015	orf1ab

KT368869	21445-25506	Middle East respiratory syndrome coronavirus	camel/Riyadh/Ry137/2015	spike

KT368870	27853-28512	Middle East respiratory syndrome coronavirus	camel/Riyadh/Ry159(b)/2015	membrane

KT368870	28566-29807	Middle East respiratory syndrome coronavirus	camel/Riyadh/Ry159(b)/2015	nucleocapsid

KT368870	279-21514	Middle East respiratory syndrome coronavirus	camel/Riyadh/Ry159(b)/2015	orf1ab

KT368870	21456-25517	Middle East respiratory syndrome coronavirus	camel/Riyadh/Ry159(b)/2015	spike

KT368871	27853-28512	Middle East respiratory syndrome coronavirus	camel/Riyadh/Ry162/2015	membrane

KT368871	28566-29807	Middle East respiratory syndrome coronavirus	camel/Riyadh/Ry162/2015	nucleocapsid

KT368871	279-21514	Middle East respiratory syndrome coronavirus	camel/Riyadh/Ry162/2015	orf1ab

KT368871	21456-25517	Middle East respiratory syndrome coronavirus	camel/Riyadh/Ry162/2015	spike

KT368872	27853-28512	Middle East respiratory syndrome coronavirus	camel/Riyadh/Ry173/2015	membrane

KT368872	28566-29807	Middle East respiratory syndrome coronavirus	camel/Riyadh/Ry173/2015	nucleocapsid

KT368872	279-21514	Middle East respiratory syndrome coronavirus	camel/Riyadh/Ry173/2015	orf1ab

KT368872	21456-25517	Middle East respiratory syndrome coronavirus	camel/Riyadh/Ry173/2015	spike

KT368873	27853-28512	Middle East respiratory syndrome coronavirus	camel/Riyadh/Ry177/2015	membrane

KT368873	28566-29807	Middle East respiratory syndrome coronavirus	camel/Riyadh/Ry177/2015	nucleocapsid

KT368873	279-21514	Middle East respiratory syndrome coronavirus	camel/Riyadh/Ry177/2015	orf1ab

KT368873	21456-25517	Middle East respiratory syndrome coronavirus	camel/Riyadh/Ry177/2015	spike

KT368874	27853-28512	Middle East respiratory syndrome coronavirus	camel/Riyadh/Ry178/2015	membrane

KT368874	28566-29807	Middle East respiratory syndrome coronavirus	camel/Riyadh/Ry178/2015	nucleocapsid

KT368874	279-21514	Middle East respiratory syndrome coronavirus	camel/Riyadh/Ry178/2015	orf1ab

KT368874	21456-25517	Middle East respiratory syndrome coronavirus	camel/Riyadh/Ry178/2015	spike

KT368875	27848-28507	Middle East respiratory syndrome coronavirus	camel/Riyadh/Ry179/2015	membrane

KT368875	28561-29802	Middle East respiratory syndrome coronavirus	camel/Riyadh/Ry179/2015	nucleocapsid

KT368875	274-21509	Middle East respiratory syndrome coronavirus	camel/Riyadh/Ry179/2015	orf1ab

KT368875	21451-25512	Middle East respiratory syndrome coronavirus	camel/Riyadh/Ry179/2015	spike

KT368825	27853-28512	Middle East respiratory syndrome coronavirus	camel/Riyadh/Ry23N/2014	membrane

KT368825	28566-29807	Middle East respiratory syndrome coronavirus	camel/Riyadh/Ry23N/2014	nucleocapsid

KT368825	279-21514	Middle East respiratory syndrome coronavirus	camel/Riyadh/Ry23N/2014	orf1ab

KT368825	21456-25517	Middle East respiratory syndrome coronavirus	camel/Riyadh/Ry23N/2014	spike

KT368876	27853-28512	Middle East respiratory syndrome coronavirus	camel/Riyadh/Ry63/2015	membrane

KT368876	28566-29807	Middle East respiratory syndrome coronavirus	camel/Riyadh/Ry63/2015	nucleocapsid

KT368876	279-21514	Middle East respiratory syndrome coronavirus	camel/Riyadh/Ry63/2015	orf1ab

KT368876	21456-25517	Middle East respiratory syndrome coronavirus	camel/Riyadh/Ry63/2015	spike

KT368877	27853-28512	Middle East respiratory syndrome coronavirus	camel/Riyadh/Ry64/2015	membrane

KT368877	28566-29807	Middle East respiratory syndrome coronavirus	camel/Riyadh/Ry64/2015	nucleocapsid

KT368877	279-21514	Middle East respiratory syndrome coronavirus	camel/Riyadh/Ry64/2015	orf1ab

KT368877	21456-25517	Middle East respiratory syndrome coronavirus	camel/Riyadh/Ry64/2015	spike

KT368878	27853-28512	Middle East respiratory syndrome coronavirus	camel/Riyadh/Ry79/2015	membrane

KT368878	28566-29807	Middle East respiratory syndrome coronavirus	camel/Riyadh/Ry79/2015	nucleocapsid

KT368878	279-21514	Middle East respiratory syndrome coronavirus	camel/Riyadh/Ry79/2015	orf1ab

KT368878	21456-25517	Middle East respiratory syndrome coronavirus	camel/Riyadh/Ry79/2015	spike

KT368826	27853-28512	Middle East respiratory syndrome coronavirus	camel/Riyadh/Ry84N/2014	membrane

KT368826	28566-29807	Middle East respiratory syndrome coronavirus	camel/Riyadh/Ry84N/2014	nucleocapsid

KT368826	279-21514	Middle East respiratory syndrome coronavirus	camel/Riyadh/Ry84N/2014	orf1ab

KT368826	21456-25517	Middle East respiratory syndrome coronavirus	camel/Riyadh/Ry84N/2014	spike

KT368879	27842-28501	Middle East respiratory syndrome coronavirus	camel/Riyadh/Ry86/2015	membrane

KT368879	28555-29796	Middle East respiratory syndrome coronavirus	camel/Riyadh/Ry86/2015	nucleocapsid

KT368879	268-21503	Middle East respiratory syndrome coronavirus	camel/Riyadh/Ry86/2015	orf1ab

KT368879	21445-25506	Middle East respiratory syndrome coronavirus	camel/Riyadh/Ry86/2015	spike

KT368889	27853-28512	Middle East respiratory syndrome coronavirus	camel/Taif/T150/2015	membrane

KT368889	28566-29807	Middle East respiratory syndrome coronavirus	camel/Taif/T150/2015	nucleocapsid

KT368889	279-21514	Middle East respiratory syndrome coronavirus	camel/Taif/T150/2015	orf1ab

KT368889	21456-25517	Middle East respiratory syndrome coronavirus	camel/Taif/T150/2015	spike

KT368890	27853-28512	Middle East respiratory syndrome coronavirus	camel/Taif/T157(b)/2015	membrane

KT368890	28566-29807	Middle East respiratory syndrome coronavirus	camel/Taif/T157(b)/2015	nucleocapsid

KT368890	279-21514	Middle East respiratory syndrome coronavirus	camel/Taif/T157(b)/2015	orf1ab

KT368890	21456-25517	Middle East respiratory syndrome coronavirus	camel/Taif/T157(b)/2015	spike

KT368882	27853-28512	Middle East respiratory syndrome coronavirus	camel/Taif/T16/2015	membrane

KT368882	28566-29807	Middle East respiratory syndrome coronavirus	camel/Taif/T16/2015	nucleocapsid

KT368882	279-21514	Middle East respiratory syndrome coronavirus	camel/Taif/T16/2015	orf1ab

KT368882	21456-25517	Middle East respiratory syndrome coronavirus	camel/Taif/T16/2015	spike

KT368883	27853-28512	Middle East respiratory syndrome coronavirus	camel/Taif/T22/2015	membrane

KT368883	28566-29807	Middle East respiratory syndrome coronavirus	camel/Taif/T22/2015	nucleocapsid

KT368883	279-21514	Middle East respiratory syndrome coronavirus	camel/Taif/T22/2015	orf1ab

KT368883	21456-25517	Middle East respiratory syndrome coronavirus	camel/Taif/T22/2015	spike

KT368880	27853-28512	Middle East respiratory syndrome coronavirus	camel/Taif/T3/2015	membrane

KT368880	28566-29807	Middle East respiratory syndrome coronavirus	camel/Taif/T3/2015	nucleocapsid

KT368880	279-21514	Middle East respiratory syndrome coronavirus	camel/Taif/T3/2015	orf1ab

KT368880	21456-25517	Middle East respiratory syndrome coronavirus	camel/Taif/T3/2015	spike

KT368884	27853-28512	Middle East respiratory syndrome coronavirus	camel/Taif/T68/2015	membrane

KT368884	28566-29807	Middle East respiratory syndrome coronavirus	camel/Taif/T68/2015	nucleocapsid

KT368884	279-21514	Middle East respiratory syndrome coronavirus	camel/Taif/T68/2015	orf1ab

KT368884	21456-25517	Middle East respiratory syndrome coronavirus	camel/Taif/T68/2015	spike

KT368881	27853-28512	Middle East respiratory syndrome coronavirus	camel/Taif/T7/2015	membrane

KT368881	28566-29807	Middle East respiratory syndrome coronavirus	camel/Taif/T7/2015	nucleocapsid

KT368881	279-21514	Middle East respiratory syndrome coronavirus	camel/Taif/T7/2015	orf1ab

KT368881	21456-25517	Middle East respiratory syndrome coronavirus	camel/Taif/T7/2015	spike

KT368885	27853-28512	Middle East respiratory syndrome coronavirus	camel/Taif/T89/2015	membrane

KT368885	28566-29807	Middle East respiratory syndrome coronavirus	camel/Taif/T89/2015	nucleocapsid

KT368885	279-21514	Middle East respiratory syndrome coronavirus	camel/Taif/T89/2015	orf1ab

KT368885	21456-25517	Middle East respiratory syndrome coronavirus	camel/Taif/T89/2015	spike

KT368887	27847-28506	Middle East respiratory syndrome coronavirus	camel/Taif/T92/2015	membrane

KT368887	28560-29801	Middle East respiratory syndrome coronavirus	camel/Taif/T92/2015	nucleocapsid

KT368887	279-21508	Middle East respiratory syndrome coronavirus	camel/Taif/T92/2015	orf1ab

KT368887	21450-25511	Middle East respiratory syndrome coronavirus	camel/Taif/T92/2015	spike

KT368888	27853-28512	Middle East respiratory syndrome coronavirus	camel/Taif/T98/2015	membrane

KT368888	28566-29807	Middle East respiratory syndrome coronavirus	camel/Taif/T98/2015	nucleocapsid

KT368888	279-21514	Middle East respiratory syndrome coronavirus	camel/Taif/T98/2015	orf1ab

KT368888	21456-25517	Middle East respiratory syndrome coronavirus	camel/Taif/T98/2015	spike

KP719928	NA	Middle East respiratory syndrome coronavirus	Camel/UAE/D1164.10/2014	membrane

KP719928	NA	Middle East respiratory syndrome coronavirus	Camel/UAE/D1164.10/2014	nucleocapsid

KP719928	NA	Middle East respiratory syndrome coronavirus	Camel/UAE/D1164.10/2014	orf1ab

KP719928	NA	Middle East respiratory syndrome coronavirus	Camel/UAE/D1164.10/2014	spike

KP719929	NA	Middle East respiratory syndrome coronavirus	Camel/UAE/D1164.11/2014	membrane

KP719929	NA	Middle East respiratory syndrome coronavirus	Camel/UAE/D1164.11/2014	nucleocapsid

KP719929	NA	Middle East respiratory syndrome coronavirus	Camel/UAE/D1164.11/2014	orf1ab

KP719929	NA	Middle East respiratory syndrome coronavirus	Camel/UAE/D1164.11/2014	spike

KP719930	NA	Middle East respiratory syndrome coronavirus	Camel/UAE/D1164.14/2014	membrane

KP719930	NA	Middle East respiratory syndrome coronavirus	Camel/UAE/D1164.14/2014	nucleocapsid

KP719930	NA	Middle East respiratory syndrome coronavirus	Camel/UAE/D1164.14/2014	orf1ab

KP719930	NA	Middle East respiratory syndrome coronavirus	Camel/UAE/D1164.14/2014	spike

KP719927	NA	Middle East respiratory syndrome coronavirus	Camel/UAE/D1164.9/2014	membrane

KP719927	NA	Middle East respiratory syndrome coronavirus	Camel/UAE/D1164.9/2014	nucleocapsid

KP719927	NA	Middle East respiratory syndrome coronavirus	Camel/UAE/D1164.9/2014	orf1ab

KP719927	NA	Middle East respiratory syndrome coronavirus	Camel/UAE/D1164.9/2014	spike

KP719933	NA	Middle East respiratory syndrome coronavirus	Camel/UAE/D1209/2014	membrane

KP719933	NA	Middle East respiratory syndrome coronavirus	Camel/UAE/D1209/2014	nucleocapsid

KP719933	NA	Middle East respiratory syndrome coronavirus	Camel/UAE/D1209/2014	orf1ab

KP719933	NA	Middle East respiratory syndrome coronavirus	Camel/UAE/D1209/2014	spike

KP719932	NA	Middle East respiratory syndrome coronavirus	Camel/UAE/D1243.12/2014	membrane

KP719932	NA	Middle East respiratory syndrome coronavirus	Camel/UAE/D1243.12/2014	nucleocapsid

KP719932	NA	Middle East respiratory syndrome coronavirus	Camel/UAE/D1243.12/2014	orf1ab

KP719932	NA	Middle East respiratory syndrome coronavirus	Camel/UAE/D1243.12/2014	spike

KP719931	NA	Middle East respiratory syndrome coronavirus	Camel/UAE/D1339.2/2014	membrane

KP719931	NA	Middle East respiratory syndrome coronavirus	Camel/UAE/D1339.2/2014	nucleocapsid

KP719931	NA	Middle East respiratory syndrome coronavirus	Camel/UAE/D1339.2/2014	orf1ab

KP719931	NA	Middle East respiratory syndrome coronavirus	Camel/UAE/D1339.2/2014	spike

KU242424	27847-28506	Middle East respiratory syndrome coronavirus	Camel/UAE/D469-14	membrane

KU242424	28560-29801	Middle East respiratory syndrome coronavirus	Camel/UAE/D469-14	nucleocapsid

KU242424	273-21508	Middle East respiratory syndrome coronavirus	Camel/UAE/D469-14	orf1ab

KU242424	21450-25511	Middle East respiratory syndrome coronavirus	Camel/UAE/D469-14	spike

KT006149	NA	Middle East respiratory syndrome coronavirus	ChinaGD01	membrane

KT006149	NA	Middle East respiratory syndrome coronavirus	ChinaGD01	nucleocapsid

KT006149	NA	Middle East respiratory syndrome coronavirus	ChinaGD01	orf1ab

KT006149	NA	Middle East respiratory syndrome coronavirus	ChinaGD01	spike

KX108944	27834-28493	Middle East respiratory syndrome coronavirus	D1157/15	membrane

KX108944	28547-29788	Middle East respiratory syndrome coronavirus	D1157/15	nucleocapsid

KX108944	260-21495	Middle East respiratory syndrome coronavirus	D1157/15	orf1ab

KX108944	21437-25498	Middle East respiratory syndrome coronavirus	D1157/15	spike

KX108937	27832-28491	Middle East respiratory syndrome coronavirus	D1164.1/14	membrane

KX108937	28545-29786	Middle East respiratory syndrome coronavirus	D1164.1/14	nucleocapsid

KX108937	258-21493	Middle East respiratory syndrome coronavirus	D1164.1/14	orf1ab

KX108937	21435-25496	Middle East respiratory syndrome coronavirus	D1164.1/14	spike

KX108946	27834-28493	Middle East respiratory syndrome coronavirus	D1189.1/15	membrane

KX108946	28547-29788	Middle East respiratory syndrome coronavirus	D1189.1/15	nucleocapsid

KX108946	260-21495	Middle East respiratory syndrome coronavirus	D1189.1/15	orf1ab

KX108946	21437-25498	Middle East respiratory syndrome coronavirus	D1189.1/15	spike

KX108945	27836-28495	Middle East respiratory syndrome coronavirus	D1271/15	membrane

KX108945	28549-29790	Middle East respiratory syndrome coronavirus	D1271/15	nucleocapsid

KX108945	262-21497	Middle East respiratory syndrome coronavirus	D1271/15	orf1ab

KX108945	21439-25500	Middle East respiratory syndrome coronavirus	D1271/15	spike

KX108939	27834-28493	Middle East respiratory syndrome coronavirus	D252/15	membrane

KX108939	28547-29788	Middle East respiratory syndrome coronavirus	D252/15	nucleocapsid

KX108939	260-21495	Middle East respiratory syndrome coronavirus	D252/15	orf1ab

KX108939	21437-25498	Middle East respiratory syndrome coronavirus	D252/15	spike

KX108938	27836-28495	Middle East respiratory syndrome coronavirus	D2597.2/14	membrane

KX108938	28549-29790	Middle East respiratory syndrome coronavirus	D2597.2/14	nucleocapsid

KX108938	262-21497	Middle East respiratory syndrome coronavirus	D2597.2/14	orf1ab

KX108938	21439-25500	Middle East respiratory syndrome coronavirus	D2597.2/14	spike

KT751244	27838-28497	Middle East respiratory syndrome coronavirus	D2731.3/14	membrane

KT751244	28551-29792	Middle East respiratory syndrome coronavirus	D2731.3/14	nucleocapsid

KT751244	264-21499	Middle East respiratory syndrome coronavirus	D2731.3/14	orf1ab

KT751244	21441-25502	Middle East respiratory syndrome coronavirus	D2731.3/14	spike

KX108940	27834-28493	Middle East respiratory syndrome coronavirus	D374/15	membrane

KX108940	28547-29788	Middle East respiratory syndrome coronavirus	D374/15	nucleocapsid

KX108940	260-21495	Middle East respiratory syndrome coronavirus	D374/15	orf1ab

KX108940	21437-25498	Middle East respiratory syndrome coronavirus	D374/15	spike

KX108941	27834-28493	Middle East respiratory syndrome coronavirus	D383/15	membrane

KX108941	28547-29788	Middle East respiratory syndrome coronavirus	D383/15	nucleocapsid

KX108941	262-21497	Middle East respiratory syndrome coronavirus	D383/15	orf1ab

KX108941	21439-25500	Middle East respiratory syndrome coronavirus	D383/15	spike

KX108942	27834-28493	Middle East respiratory syndrome coronavirus	D389/15	membrane

KX108942	28547-29788	Middle East respiratory syndrome coronavirus	D389/15	nucleocapsid

KX108942	260-21495	Middle East respiratory syndrome coronavirus	D389/15	orf1ab

KX108942	21437-25498	Middle East respiratory syndrome coronavirus	D389/15	spike

KX108943	27823-28482	Middle East respiratory syndrome coronavirus	D998/15	membrane

KX108943	28536-29777	Middle East respiratory syndrome coronavirus	D998/15	nucleocapsid

KX108943	258-21493	Middle East respiratory syndrome coronavirus	D998/15	orf1ab

KX108943	21435-25496	Middle East respiratory syndrome coronavirus	D998/15	spike

KM015348	NA	Middle East respiratory syndrome coronavirus	England/2/2013	membrane

KM015348	NA	Middle East respiratory syndrome coronavirus	England/2/2013	nucleocapsid

KM015348	NA	Middle East respiratory syndrome coronavirus	England/2/2013	orf1ab

KM015348	NA	Middle East respiratory syndrome coronavirus	England/2/2013	spike

KM210278	NA	Middle East respiratory syndrome coronavirus	England/3/2013	membrane

KM210278	NA	Middle East respiratory syndrome coronavirus	England/3/2013	nucleocapsid

KM210278	NA	Middle East respiratory syndrome coronavirus	England/3/2013	orf1ab

KM210278	NA	Middle East respiratory syndrome coronavirus	England/3/2013	spike

KM210277	NA	Middle East respiratory syndrome coronavirus	England/4/2013	membrane

KM210277	NA	Middle East respiratory syndrome coronavirus	England/4/2013	nucleocapsid

KM210277	NA	Middle East respiratory syndrome coronavirus	England/4/2013	orf1ab

KM210277	NA	Middle East respiratory syndrome coronavirus	England/4/2013	spike

KJ829365	27853-28512	Middle East respiratory syndrome coronavirus	Florida/USA-2_Saudi Arabia_2014	membrane

KJ829365	28566-29807	Middle East respiratory syndrome coronavirus	Florida/USA-2_Saudi Arabia_2014	nucleocapsid

KJ829365	279-21514	Middle East respiratory syndrome coronavirus	Florida/USA-2_Saudi Arabia_2014	orf1ab

KJ829365	21456-25517	Middle East respiratory syndrome coronavirus	Florida/USA-2_Saudi Arabia_2014	spike

KP223131	27853-28512	Middle East respiratory syndrome coronavirus	Florida/USA-2_Saudi Arabia_2014	membrane

KP223131	28566-29807	Middle East respiratory syndrome coronavirus	Florida/USA-2_Saudi Arabia_2014	nucleocapsid

KP223131	279-21514	Middle East respiratory syndrome coronavirus	Florida/USA-2_Saudi Arabia_2014	orf1ab

KP223131	21456-25517	Middle East respiratory syndrome coronavirus	Florida/USA-2_Saudi Arabia_2014	spike

KF745068	NA	Middle East respiratory syndrome coronavirus	FRA/UAE	membrane

KF745068	NA	Middle East respiratory syndrome coronavirus	FRA/UAE	nucleocapsid

KF745068	NA	Middle East respiratory syndrome coronavirus	FRA/UAE	orf1ab

KF745068	NA	Middle East respiratory syndrome coronavirus	FRA/UAE	spike

KF600628	NA	Middle East respiratory syndrome coronavirus	Hafr-Al-Batin_1_2013	membrane

KF600628	NA	Middle East respiratory syndrome coronavirus	Hafr-Al-Batin_1_2013	nucleocapsid

KF600628	NA	Middle East respiratory syndrome coronavirus	Hafr-Al-Batin_1_2013	orf1ab

KF600628	NA	Middle East respiratory syndrome coronavirus	Hafr-Al-Batin_1_2013	spike

KJ156910	NA	Middle East respiratory syndrome coronavirus	Hafr-Al-Batin_2_2013	membrane

KJ156910	NA	Middle East respiratory syndrome coronavirus	Hafr-Al-Batin_2_2013	nucleocapsid

KJ156910	NA	Middle East respiratory syndrome coronavirus	Hafr-Al-Batin_2_2013	orf1ab

KJ156910	NA	Middle East respiratory syndrome coronavirus	Hafr-Al-Batin_2_2013	spike

KJ156874	NA	Middle East respiratory syndrome coronavirus	Hafr-Al-Batin_6_2013	membrane

KJ156874	NA	Middle East respiratory syndrome coronavirus	Hafr-Al-Batin_6_2013	nucleocapsid

KJ156874	NA	Middle East respiratory syndrome coronavirus	Hafr-Al-Batin_6_2013	orf1ab

KJ156874	NA	Middle East respiratory syndrome coronavirus	Hafr-Al-Batin_6_2013	spike

KJ361500	NA	Middle East respiratory syndrome coronavirus	Hu-France (UAE) - FRA1_1627-2013_BAL_Sanger	membrane

KJ361500	NA	Middle East respiratory syndrome coronavirus	Hu-France (UAE) - FRA1_1627-2013_BAL_Sanger	nucleocapsid

KJ361500	NA	Middle East respiratory syndrome coronavirus	Hu-France (UAE) - FRA1_1627-2013_BAL_Sanger	spike

KJ361502	NA	Middle East respiratory syndrome coronavirus	Hu-France - FRA2_130569-2013_InSpu_Sanger	membrane

KJ361502	NA	Middle East respiratory syndrome coronavirus	Hu-France - FRA2_130569-2013_InSpu_Sanger	nucleocapsid

KJ361502	NA	Middle East respiratory syndrome coronavirus	Hu-France - FRA2_130569-2013_InSpu_Sanger	spike

KJ361501	NA	Middle East respiratory syndrome coronavirus	Hu-France - FRA2_130569-2013_IS_HTS	membrane

KJ361501	NA	Middle East respiratory syndrome coronavirus	Hu-France - FRA2_130569-2013_IS_HTS	nucleocapsid

KJ361501	NA	Middle East respiratory syndrome coronavirus	Hu-France - FRA2_130569-2013_IS_HTS	spike

KJ361503	NA	Middle East respiratory syndrome coronavirus	Hu-France - FRA2_130569-2013_Isolate_Sanger	membrane

KJ361503	NA	Middle East respiratory syndrome coronavirus	Hu-France - FRA2_130569-2013_Isolate_Sanger	nucleocapsid

KJ361503	NA	Middle East respiratory syndrome coronavirus	Hu-France - FRA2_130569-2013_Isolate_Sanger	spike

KT806046	27831-28490	Middle East respiratory syndrome coronavirus	Hu/Hufuf-KSA-11002/2015	membrane

KT806046	28544-29785	Middle East respiratory syndrome coronavirus	Hu/Hufuf-KSA-11002/2015	nucleocapsid

KT806046	257-21492	Middle East respiratory syndrome coronavirus	Hu/Hufuf-KSA-11002/2015	orf1ab

KT806046	21434-25495	Middle East respiratory syndrome coronavirus	Hu/Hufuf-KSA-11002/2015	spike

KT806047	27822-28481	Middle East respiratory syndrome coronavirus	Hu/Hufuf-KSA-9158/2015	membrane

KT806047	28535-29776	Middle East respiratory syndrome coronavirus	Hu/Hufuf-KSA-9158/2015	nucleocapsid

KT806047	257-21492	Middle East respiratory syndrome coronavirus	Hu/Hufuf-KSA-9158/2015	orf1ab

KT806047	21434-25495	Middle East respiratory syndrome coronavirus	Hu/Hufuf-KSA-9158/2015	spike

KU851859	27811-28470	Middle East respiratory syndrome coronavirus	Hu/Jeddah-KSA-3RS2702/2015	membrane

KU851859	28524-29765	Middle East respiratory syndrome coronavirus	Hu/Jeddah-KSA-3RS2702/2015	nucleocapsid

KU851859	257-21492	Middle East respiratory syndrome coronavirus	Hu/Jeddah-KSA-3RS2702/2015	orf1ab

KU851859	21434-25495	Middle East respiratory syndrome coronavirus	Hu/Jeddah-KSA-3RS2702/2015	spike

KT806044	27831-28490	Middle East respiratory syndrome coronavirus	Hu/Jeddah-KSA-C20843/2015	membrane

KT806044	28544-29785	Middle East respiratory syndrome coronavirus	Hu/Jeddah-KSA-C20843/2015	nucleocapsid

KT806044	257-21492	Middle East respiratory syndrome coronavirus	Hu/Jeddah-KSA-C20843/2015	orf1ab

KT806044	21434-25495	Middle East respiratory syndrome coronavirus	Hu/Jeddah-KSA-C20843/2015	spike

KT806055	27831-28490	Middle East respiratory syndrome coronavirus	Hu/Jeddah-KSA-C20860/2015	membrane

KT806055	28544-29785	Middle East respiratory syndrome coronavirus	Hu/Jeddah-KSA-C20860/2015	nucleocapsid

KT806055	257-21492	Middle East respiratory syndrome coronavirus	Hu/Jeddah-KSA-C20860/2015	orf1ab

KT806055	21434-25495	Middle East respiratory syndrome coronavirus	Hu/Jeddah-KSA-C20860/2015	spike

KT806045	27831-28490	Middle East respiratory syndrome coronavirus	Hu/Jeddah-KSA-C21271/2015	membrane

KT806045	28544-29785	Middle East respiratory syndrome coronavirus	Hu/Jeddah-KSA-C21271/2015	nucleocapsid

KT806045	257-21492	Middle East respiratory syndrome coronavirus	Hu/Jeddah-KSA-C21271/2015	orf1ab

KT806045	21434-25495	Middle East respiratory syndrome coronavirus	Hu/Jeddah-KSA-C21271/2015	spike

KT861627	27831-28490	Middle East respiratory syndrome coronavirus	Hu/Jordan-20140010168/2014	membrane

KT861627	28537-29778	Middle East respiratory syndrome coronavirus	Hu/Jordan-20140010168/2014	nucleocapsid

KT861627	257-21492	Middle East respiratory syndrome coronavirus	Hu/Jordan-20140010168/2014	orf1ab

KT861627	21434-25495	Middle East respiratory syndrome coronavirus	Hu/Jordan-20140010168/2014	spike

KT861628	27831-28490	Middle East respiratory syndrome coronavirus	Hu/Jordan-201440011123/2014	membrane

KT861628	28544-29785	Middle East respiratory syndrome coronavirus	Hu/Jordan-201440011123/2014	nucleocapsid

KT861628	257-21492	Middle East respiratory syndrome coronavirus	Hu/Jordan-201440011123/2014	orf1ab

KT861628	21434-25495	Middle East respiratory syndrome coronavirus	Hu/Jordan-201440011123/2014	spike

KT806053	27831-28490	Middle East respiratory syndrome coronavirus	Hu/Kharj-KSA-2598/2015	membrane

KT806053	28544-29785	Middle East respiratory syndrome coronavirus	Hu/Kharj-KSA-2598/2015	nucleocapsid

KT806053	257-21492	Middle East respiratory syndrome coronavirus	Hu/Kharj-KSA-2598/2015	orf1ab

KT806053	21434-25495	Middle East respiratory syndrome coronavirus	Hu/Kharj-KSA-2598/2015	spike

KT806052	27831-28490	Middle East respiratory syndrome coronavirus	Hu/Kharj-KSA-2599/2015	membrane

KT806052	28544-29785	Middle East respiratory syndrome coronavirus	Hu/Kharj-KSA-2599/2015	nucleocapsid

KT806052	257-21492	Middle East respiratory syndrome coronavirus	Hu/Kharj-KSA-2599/2015	orf1ab

KT806052	21434-25495	Middle East respiratory syndrome coronavirus	Hu/Kharj-KSA-2599/2015	spike

KT806048	27831-28490	Middle East respiratory syndrome coronavirus	Hu/Khobar-KSA-6736/2015	membrane

KT806048	28544-29785	Middle East respiratory syndrome coronavirus	Hu/Khobar-KSA-6736/2015	nucleocapsid

KT806048	257-21492	Middle East respiratory syndrome coronavirus	Hu/Khobar-KSA-6736/2015	orf1ab

KT806048	21434-25495	Middle East respiratory syndrome coronavirus	Hu/Khobar-KSA-6736/2015	spike

KT806054	27717-28376	Middle East respiratory syndrome coronavirus	Hu/Najran-KSA-C20915/2015	membrane

KT806054	28430-29671	Middle East respiratory syndrome coronavirus	Hu/Najran-KSA-C20915/2015	nucleocapsid

KT806054	143-21378	Middle East respiratory syndrome coronavirus	Hu/Najran-KSA-C20915/2015	orf1ab

KT806054	21320-25381	Middle East respiratory syndrome coronavirus	Hu/Najran-KSA-C20915/2015	spike

KT156560	27853-28512	Middle East respiratory syndrome coronavirus	Hu/Oman_2285_2013	membrane

KT156560	28566-29807	Middle East respiratory syndrome coronavirus	Hu/Oman_2285_2013	nucleocapsid

KT156560	279-21514	Middle East respiratory syndrome coronavirus	Hu/Oman_2285_2013	orf1ab

KT156560	21456-25517	Middle East respiratory syndrome coronavirus	Hu/Oman_2285_2013	spike

KT156561	27853-28512	Middle East respiratory syndrome coronavirus	Hu/Oman_2874_2013	membrane

KT156561	28566-29807	Middle East respiratory syndrome coronavirus	Hu/Oman_2874_2013	nucleocapsid

KT156561	279-21514	Middle East respiratory syndrome coronavirus	Hu/Oman_2874_2013	orf1ab

KT156561	21456-25517	Middle East respiratory syndrome coronavirus	Hu/Oman_2874_2013	spike

KU851863	27831-28490	Middle East respiratory syndrome coronavirus	Hu/Riyadh-KSA-16077/2015	membrane

KU851863	28544-29785	Middle East respiratory syndrome coronavirus	Hu/Riyadh-KSA-16077/2015	nucleocapsid

KU851863	257-21492	Middle East respiratory syndrome coronavirus	Hu/Riyadh-KSA-16077/2015	orf1ab

KU851863	21434-25495	Middle East respiratory syndrome coronavirus	Hu/Riyadh-KSA-16077/2015	spike

KU851864	27831-28490	Middle East respiratory syndrome coronavirus	Hu/Riyadh-KSA-16098/2015	membrane

KU851864	28544-29785	Middle East respiratory syndrome coronavirus	Hu/Riyadh-KSA-16098/2015	nucleocapsid

KU851864	257-21492	Middle East respiratory syndrome coronavirus	Hu/Riyadh-KSA-16098/2015	orf1ab

KU851864	21434-25495	Middle East respiratory syndrome coronavirus	Hu/Riyadh-KSA-16098/2015	spike

KU851862	27831-28490	Middle East respiratory syndrome coronavirus	Hu/Riyadh-KSA-16117/2015	membrane

KU851862	28544-29785	Middle East respiratory syndrome coronavirus	Hu/Riyadh-KSA-16117/2015	nucleocapsid

KU851862	257-21492	Middle East respiratory syndrome coronavirus	Hu/Riyadh-KSA-16117/2015	orf1ab

KU851862	21434-25495	Middle East respiratory syndrome coronavirus	Hu/Riyadh-KSA-16117/2015	spike

KU851861	27831-28490	Middle East respiratory syndrome coronavirus	Hu/Riyadh-KSA-16120/2015	membrane

KU851861	28544-29785	Middle East respiratory syndrome coronavirus	Hu/Riyadh-KSA-16120/2015	nucleocapsid

KU851861	257-21492	Middle East respiratory syndrome coronavirus	Hu/Riyadh-KSA-16120/2015	orf1ab

KU851861	21434-25495	Middle East respiratory syndrome coronavirus	Hu/Riyadh-KSA-16120/2015	spike

KU851860	27831-28490	Middle East respiratory syndrome coronavirus	Hu/Riyadh-KSA-16121/2015	membrane

KU851860	28544-29785	Middle East respiratory syndrome coronavirus	Hu/Riyadh-KSA-16121/2015	nucleocapsid

KU851860	257-21492	Middle East respiratory syndrome coronavirus	Hu/Riyadh-KSA-16121/2015	orf1ab

KU851860	21434-25495	Middle East respiratory syndrome coronavirus	Hu/Riyadh-KSA-16121/2015	spike

KR011263	NA	Middle East respiratory syndrome coronavirus	Hu/Riyadh-KSA-2345/2015	membrane

KR011263	NA	Middle East respiratory syndrome coronavirus	Hu/Riyadh-KSA-2345/2015	nucleocapsid

KR011263	NA	Middle East respiratory syndrome coronavirus	Hu/Riyadh-KSA-2345/2015	orf1ab

KR011263	NA	Middle East respiratory syndrome coronavirus	Hu/Riyadh-KSA-2345/2015	spike

KR011265	NA	Middle East respiratory syndrome coronavirus	Hu/Riyadh-KSA-2466/2015	membrane

KR011265	NA	Middle East respiratory syndrome coronavirus	Hu/Riyadh-KSA-2466/2015	nucleocapsid

KR011265	NA	Middle East respiratory syndrome coronavirus	Hu/Riyadh-KSA-2466/2015	orf1ab

KR011265	NA	Middle East respiratory syndrome coronavirus	Hu/Riyadh-KSA-2466/2015	spike

KT806051	27831-28490	Middle East respiratory syndrome coronavirus	Hu/Riyadh-KSA-2716/2015	membrane

KT806051	28544-29785	Middle East respiratory syndrome coronavirus	Hu/Riyadh-KSA-2716/2015	nucleocapsid

KT806051	257-21492	Middle East respiratory syndrome coronavirus	Hu/Riyadh-KSA-2716/2015	orf1ab

KT806051	21434-25495	Middle East respiratory syndrome coronavirus	Hu/Riyadh-KSA-2716/2015	spike

KT806049	27831-28490	Middle East respiratory syndrome coronavirus	Hu/Riyadh-KSA-3181/2015	membrane

KT806049	28544-29785	Middle East respiratory syndrome coronavirus	Hu/Riyadh-KSA-3181/2015	nucleocapsid

KT806049	257-21492	Middle East respiratory syndrome coronavirus	Hu/Riyadh-KSA-3181/2015	orf1ab

KT806049	21434-25495	Middle East respiratory syndrome coronavirus	Hu/Riyadh-KSA-3181/2015	spike

KT026453	27844-28503	Middle East respiratory syndrome coronavirus	Hu/Riyadh_KSA_2959_2015	membrane

KT026453	28557-29798	Middle East respiratory syndrome coronavirus	Hu/Riyadh_KSA_2959_2015	nucleocapsid

KT026453	270-21505	Middle East respiratory syndrome coronavirus	Hu/Riyadh_KSA_2959_2015	orf1ab

KT026453	21447-25508	Middle East respiratory syndrome coronavirus	Hu/Riyadh_KSA_2959_2015	spike

KT026454	27853-28512	Middle East respiratory syndrome coronavirus	Hu/Riyadh_KSA_4050_2015	membrane

KT026454	28566-29807	Middle East respiratory syndrome coronavirus	Hu/Riyadh_KSA_4050_2015	nucleocapsid

KT026454	279-21514	Middle East respiratory syndrome coronavirus	Hu/Riyadh_KSA_4050_2015	orf1ab

KT026454	21456-25517	Middle East respiratory syndrome coronavirus	Hu/Riyadh_KSA_4050_2015	spike

KT026456	27850-28509	Middle East respiratory syndrome coronavirus	Hu/Riyadh_KSA_4050_2015	membrane

KT026456	28563-29804	Middle East respiratory syndrome coronavirus	Hu/Riyadh_KSA_4050_2015	nucleocapsid

KT026456	279-21511	Middle East respiratory syndrome coronavirus	Hu/Riyadh_KSA_4050_2015	orf1ab

KT026456	21453-25514	Middle East respiratory syndrome coronavirus	Hu/Riyadh_KSA_4050_2015	spike

KU710264	27831-28490	Middle East respiratory syndrome coronavirus	Hu/Taif/KSA-7032/2014	membrane

KU710264	28544-29785	Middle East respiratory syndrome coronavirus	Hu/Taif/KSA-7032/2014	nucleocapsid

KU710264	257-21492	Middle East respiratory syndrome coronavirus	Hu/Taif/KSA-7032/2014	orf1ab

KU710264	21434-25495	Middle East respiratory syndrome coronavirus	Hu/Taif/KSA-7032/2014	spike

KJ813439	27853-28512	Middle East respiratory syndrome coronavirus	Indiana/USA-1_Saudi Arabia_2014	membrane

KJ813439	28566-29807	Middle East respiratory syndrome coronavirus	Indiana/USA-1_Saudi Arabia_2014	nucleocapsid

KJ813439	279-21514	Middle East respiratory syndrome coronavirus	Indiana/USA-1_Saudi Arabia_2014	orf1ab

KJ813439	21456-25517	Middle East respiratory syndrome coronavirus	Indiana/USA-1_Saudi Arabia_2014	spike

KJ556336	NA	Middle East respiratory syndrome coronavirus	Jeddah_1_2013	membrane

KJ556336	NA	Middle East respiratory syndrome coronavirus	Jeddah_1_2013	nucleocapsid

KJ556336	NA	Middle East respiratory syndrome coronavirus	Jeddah_1_2013	orf1ab

KJ556336	NA	Middle East respiratory syndrome coronavirus	Jeddah_1_2013	spike

KM027260	NA	Middle East respiratory syndrome coronavirus	Jeddah_C10306/KSA/2014-04-20	membrane

KM027260	NA	Middle East respiratory syndrome coronavirus	Jeddah_C10306/KSA/2014-04-20	nucleocapsid

KM027260	21350-25411	Middle East respiratory syndrome coronavirus	Jeddah_C10306/KSA/2014-04-20	spike

KM027255	NA	Middle East respiratory syndrome coronavirus	Jeddah_C7149/KSA/2014-04-05	membrane

KM027255	NA	Middle East respiratory syndrome coronavirus	Jeddah_C7149/KSA/2014-04-05	nucleocapsid

KM027255	21419-25480	Middle East respiratory syndrome coronavirus	Jeddah_C7149/KSA/2014-04-05	spike

KM027256	NA	Middle East respiratory syndrome coronavirus	Jeddah_C7569/KSA/2014-04-03	membrane

KM027256	NA	Middle East respiratory syndrome coronavirus	Jeddah_C7569/KSA/2014-04-03	nucleocapsid

KM027256	21419-25480	Middle East respiratory syndrome coronavirus	Jeddah_C7569/KSA/2014-04-03	spike

KM027257	NA	Middle East respiratory syndrome coronavirus	Jeddah_C7770/KSA/2014-04-07	membrane

KM027257	NA	Middle East respiratory syndrome coronavirus	Jeddah_C7770/KSA/2014-04-07	nucleocapsid

KM027257	21419-25480	Middle East respiratory syndrome coronavirus	Jeddah_C7770/KSA/2014-04-07	spike

KM027258	NA	Middle East respiratory syndrome coronavirus	Jeddah_C8826/KSA/2014-04-12	membrane

KM027258	NA	Middle East respiratory syndrome coronavirus	Jeddah_C8826/KSA/2014-04-12	nucleocapsid

KM027258	21419-25480	Middle East respiratory syndrome coronavirus	Jeddah_C8826/KSA/2014-04-12	spike

KM027259	NA	Middle East respiratory syndrome coronavirus	Jeddah_C9055/KSA/2014-04-14	membrane

KM027259	NA	Middle East respiratory syndrome coronavirus	Jeddah_C9055/KSA/2014-04-14	nucleocapsid

KM027259	21419-25480	Middle East respiratory syndrome coronavirus	Jeddah_C9055/KSA/2014-04-14	spike

KT121580	NA	Middle East respiratory syndrome coronavirus	KFMC-1	orf1ab

KT121580	NA	Middle East respiratory syndrome coronavirus	KFMC-1	membrane

KT121580	NA	Middle East respiratory syndrome coronavirus	KFMC-1	nucleocapsid

KT121580	NA	Middle East respiratory syndrome coronavirus	KFMC-1	spike

KT121578	NA	Middle East respiratory syndrome coronavirus	KFMC-10	orf1ab

KT121578	NA	Middle East respiratory syndrome coronavirus	KFMC-10	membrane

KT121578	NA	Middle East respiratory syndrome coronavirus	KFMC-10	nucleocapsid

KT121578	NA	Middle East respiratory syndrome coronavirus	KFMC-10	spike

KT121577	NA	Middle East respiratory syndrome coronavirus	KFMC-2	orf1ab

KT121577	NA	Middle East respiratory syndrome coronavirus	KFMC-2	membrane

KT121577	NA	Middle East respiratory syndrome coronavirus	KFMC-2	nucleocapsid

KT121577	NA	Middle East respiratory syndrome coronavirus	KFMC-2	spike

KT121572	NA	Middle East respiratory syndrome coronavirus	KFMC-5	orf1ab

KT121572	NA	Middle East respiratory syndrome coronavirus	KFMC-5	membrane

KT121572	NA	Middle East respiratory syndrome coronavirus	KFMC-5	nucleocapsid

KT121572	NA	Middle East respiratory syndrome coronavirus	KFMC-5	spike

KT121576	NA	Middle East respiratory syndrome coronavirus	KFMC-6	orf1ab

KT121576	NA	Middle East respiratory syndrome coronavirus	KFMC-6	membrane

KT121576	NA	Middle East respiratory syndrome coronavirus	KFMC-6	nucleocapsid

KT121576	NA	Middle East respiratory syndrome coronavirus	KFMC-6	spike

KT121581	NA	Middle East respiratory syndrome coronavirus	KFMC-7	orf1ab

KT121581	NA	Middle East respiratory syndrome coronavirus	KFMC-7	membrane

KT121581	NA	Middle East respiratory syndrome coronavirus	KFMC-7	nucleocapsid

KT121581	NA	Middle East respiratory syndrome coronavirus	KFMC-7	spike

KT121579	NA	Middle East respiratory syndrome coronavirus	KFMC-8	orf1ab

KT121579	NA	Middle East respiratory syndrome coronavirus	KFMC-8	membrane

KT121579	NA	Middle East respiratory syndrome coronavirus	KFMC-8	nucleocapsid

KT121579	NA	Middle East respiratory syndrome coronavirus	KFMC-8	spike

KT121574	NA	Middle East respiratory syndrome coronavirus	KFMC-9	orf1ab

KT121574	NA	Middle East respiratory syndrome coronavirus	KFMC-9	membrane

KT121574	NA	Middle East respiratory syndrome coronavirus	KFMC-9	nucleocapsid

KT121574	NA	Middle East respiratory syndrome coronavirus	KFMC-9	spike

KJ650297	NA	Middle East respiratory syndrome coronavirus	KFU-HKU 1	orf1ab

KJ650297	NA	Middle East respiratory syndrome coronavirus	KFU-HKU 1	membrane

KJ650297	NA	Middle East respiratory syndrome coronavirus	KFU-HKU 1	nucleocapsid

KJ650297	NA	Middle East respiratory syndrome coronavirus	KFU-HKU 1	spike

KJ650295	NA	Middle East respiratory syndrome coronavirus	KFU-HKU 13	orf1ab

KJ650295	NA	Middle East respiratory syndrome coronavirus	KFU-HKU 13	membrane

KJ650295	NA	Middle East respiratory syndrome coronavirus	KFU-HKU 13	nucleocapsid

KJ650295	NA	Middle East respiratory syndrome coronavirus	KFU-HKU 13	spike

KJ650296	NA	Middle East respiratory syndrome coronavirus	KFU-HKU 19Dam	orf1ab

KJ650296	NA	Middle East respiratory syndrome coronavirus	KFU-HKU 19Dam	membrane

KJ650296	NA	Middle East respiratory syndrome coronavirus	KFU-HKU 19Dam	nucleocapsid

KJ650296	NA	Middle East respiratory syndrome coronavirus	KFU-HKU 19Dam	spike

KT374052	NA	Middle East respiratory syndrome coronavirus	KOREA/Seoul/014-1-2015	membrane

KT374052	NA	Middle East respiratory syndrome coronavirus	KOREA/Seoul/014-1-2015	nucleocapsid

KT374052	NA	Middle East respiratory syndrome coronavirus	KOREA/Seoul/014-1-2015	orf1ab

KT374052	NA	Middle East respiratory syndrome coronavirus	KOREA/Seoul/014-1-2015	spike

KT374053	NA	Middle East respiratory syndrome coronavirus	KOREA/Seoul/014-2-2015	membrane

KT374053	NA	Middle East respiratory syndrome coronavirus	KOREA/Seoul/014-2-2015	nucleocapsid

KT374053	NA	Middle East respiratory syndrome coronavirus	KOREA/Seoul/014-2-2015	orf1ab

KT374053	NA	Middle East respiratory syndrome coronavirus	KOREA/Seoul/014-2-2015	spike

KT374054	NA	Middle East respiratory syndrome coronavirus	KOREA/Seoul/035-1-2015	membrane

KT374054	NA	Middle East respiratory syndrome coronavirus	KOREA/Seoul/035-1-2015	nucleocapsid

KT374054	NA	Middle East respiratory syndrome coronavirus	KOREA/Seoul/035-1-2015	orf1ab

KT374054	NA	Middle East respiratory syndrome coronavirus	KOREA/Seoul/035-1-2015	spike

KT374050	NA	Middle East respiratory syndrome coronavirus	KOREA/Seoul/163-2-2015	membrane

KT374050	NA	Middle East respiratory syndrome coronavirus	KOREA/Seoul/163-2-2015	nucleocapsid

KT374050	NA	Middle East respiratory syndrome coronavirus	KOREA/Seoul/163-2-2015	orf1ab

KT374050	NA	Middle East respiratory syndrome coronavirus	KOREA/Seoul/163-2-2015	spike

KT374056	NA	Middle East respiratory syndrome coronavirus	KOREA/Seoul/168-1-2015	membrane

KT374056	NA	Middle East respiratory syndrome coronavirus	KOREA/Seoul/168-1-2015	nucleocapsid

KT374056	NA	Middle East respiratory syndrome coronavirus	KOREA/Seoul/168-1-2015	orf1ab

KT374056	NA	Middle East respiratory syndrome coronavirus	KOREA/Seoul/168-1-2015	spike

KT374057	NA	Middle East respiratory syndrome coronavirus	KOREA/Seoul/168-2-2015	membrane

KT374057	NA	Middle East respiratory syndrome coronavirus	KOREA/Seoul/168-2-2015	nucleocapsid

KT374057	NA	Middle East respiratory syndrome coronavirus	KOREA/Seoul/168-2-2015	orf1ab

KT374057	NA	Middle East respiratory syndrome coronavirus	KOREA/Seoul/168-2-2015	spike

KU308549	NA	Middle East respiratory syndrome coronavirus	Korea/Seoul/SNU1-035/2015	membrane

KU308549	NA	Middle East respiratory syndrome coronavirus	Korea/Seoul/SNU1-035/2015	nucleocapsid

KU308549	NA	Middle East respiratory syndrome coronavirus	Korea/Seoul/SNU1-035/2015	orf1ab

KU308549	NA	Middle East respiratory syndrome coronavirus	Korea/Seoul/SNU1-035/2015	spike

KJ713298	NA	Middle East respiratory syndrome coronavirus	KSA-CAMEL-363	membrane

KJ713298	NA	Middle East respiratory syndrome coronavirus	KSA-CAMEL-363	nucleocapsid

KJ713298	NA	Middle East respiratory syndrome coronavirus	KSA-CAMEL-363	orf1ab

KJ713298	NA	Middle East respiratory syndrome coronavirus	KSA-CAMEL-363	spike

KJ713299	NA	Middle East respiratory syndrome coronavirus	KSA-CAMEL-376	membrane

KJ713299	NA	Middle East respiratory syndrome coronavirus	KSA-CAMEL-376	nucleocapsid

KJ713299	NA	Middle East respiratory syndrome coronavirus	KSA-CAMEL-376	orf1ab

KJ713299	NA	Middle East respiratory syndrome coronavirus	KSA-CAMEL-376	spike

KJ713296	NA	Middle East respiratory syndrome coronavirus	KSA-CAMEL-378	membrane

KJ713296	NA	Middle East respiratory syndrome coronavirus	KSA-CAMEL-378	nucleocapsid

KJ713296	NA	Middle East respiratory syndrome coronavirus	KSA-CAMEL-378	orf1ab

KJ713296	NA	Middle East respiratory syndrome coronavirus	KSA-CAMEL-378	spike

KJ713297	NA	Middle East respiratory syndrome coronavirus	KSA-CAMEL-503	membrane

KJ713297	NA	Middle East respiratory syndrome coronavirus	KSA-CAMEL-503	nucleocapsid

KJ713297	NA	Middle East respiratory syndrome coronavirus	KSA-CAMEL-503	orf1ab

KJ713297	NA	Middle East respiratory syndrome coronavirus	KSA-CAMEL-503	spike

KJ713295	NA	Middle East respiratory syndrome coronavirus	KSA-CAMEL-505	membrane

KJ713295	NA	Middle East respiratory syndrome coronavirus	KSA-CAMEL-505	nucleocapsid

KJ713295	NA	Middle East respiratory syndrome coronavirus	KSA-CAMEL-505	orf1ab

KJ713295	NA	Middle East respiratory syndrome coronavirus	KSA-CAMEL-505	spike

KT877351	NA	Middle East respiratory syndrome coronavirus	KSA_1724	membrane

KT877351	NA	Middle East respiratory syndrome coronavirus	KSA_1724	nucleocapsid

KT877351	NA	Middle East respiratory syndrome coronavirus	KSA_1724	orf1ab

KT877351	NA	Middle East respiratory syndrome coronavirus	KSA_1724	spike

KT877350	NA	Middle East respiratory syndrome coronavirus	KSA_1725	membrane

KT877350	NA	Middle East respiratory syndrome coronavirus	KSA_1725	nucleocapsid

KT877350	NA	Middle East respiratory syndrome coronavirus	KSA_1725	orf1ab

KT877350	NA	Middle East respiratory syndrome coronavirus	KSA_1725	spike

KM027261	NA	Middle East respiratory syndrome coronavirus	Makkah_C9355/KSA/Makkah/2014-04-15	membrane

KM027261	NA	Middle East respiratory syndrome coronavirus	Makkah_C9355/KSA/Makkah/2014-04-15	nucleocapsid

KM027261	21419-25480	Middle East respiratory syndrome coronavirus	Makkah_C9355/KSA/Makkah/2014-04-15	spike

KF958702	NA	Middle East respiratory syndrome coronavirus	MERS-CoV-Jeddah-human-1	membrane

KF958702	NA	Middle East respiratory syndrome coronavirus	MERS-CoV-Jeddah-human-1	nucleocapsid

KF958702	NA	Middle East respiratory syndrome coronavirus	MERS-CoV-Jeddah-human-1	orf1ab

KF958702	NA	Middle East respiratory syndrome coronavirus	MERS-CoV-Jeddah-human-1	spike

KT029139	NA	Middle East respiratory syndrome coronavirus	MERS-CoV/KOR/KNIH/002_05_2015	membrane

KT029139	NA	Middle East respiratory syndrome coronavirus	MERS-CoV/KOR/KNIH/002_05_2015	nucleocapsid

KT029139	NA	Middle East respiratory syndrome coronavirus	MERS-CoV/KOR/KNIH/002_05_2015	orf1ab

KT029139	NA	Middle East respiratory syndrome coronavirus	MERS-CoV/KOR/KNIH/002_05_2015	spike

KX034094	27853-28512	Middle East respiratory syndrome coronavirus	MERS-CoV/KOR/Seoul/050-1-2015	membrane

KX034094	28566-29807	Middle East respiratory syndrome coronavirus	MERS-CoV/KOR/Seoul/050-1-2015	nucleocapsid

KX034094	279-21514	Middle East respiratory syndrome coronavirus	MERS-CoV/KOR/Seoul/050-1-2015	orf1ab

KX034094	21456-25517	Middle East respiratory syndrome coronavirus	MERS-CoV/KOR/Seoul/050-1-2015	spike

KX034095	27853-28512	Middle East respiratory syndrome coronavirus	MERS-CoV/KOR/Seoul/066-2015	membrane

KX034095	28566-29807	Middle East respiratory syndrome coronavirus	MERS-CoV/KOR/Seoul/066-2015	nucleocapsid

KX034095	279-21514	Middle East respiratory syndrome coronavirus	MERS-CoV/KOR/Seoul/066-2015	orf1ab

KX034095	21456-25517	Middle East respiratory syndrome coronavirus	MERS-CoV/KOR/Seoul/066-2015	spike

KX034096	27817-28476	Middle East respiratory syndrome coronavirus	MERS-CoV/KOR/Seoul/077-2-2015	membrane

KX034096	28530-29771	Middle East respiratory syndrome coronavirus	MERS-CoV/KOR/Seoul/077-2-2015	nucleocapsid

KX034096	243-21478	Middle East respiratory syndrome coronavirus	MERS-CoV/KOR/Seoul/077-2-2015	orf1ab

KX034096	21420-25481	Middle East respiratory syndrome coronavirus	MERS-CoV/KOR/Seoul/077-2-2015	spike

KX034097	27853-28512	Middle East respiratory syndrome coronavirus	MERS-CoV/KOR/Seoul/080-3-2015	membrane

KX034097	28566-29807	Middle East respiratory syndrome coronavirus	MERS-CoV/KOR/Seoul/080-3-2015	nucleocapsid

KX034097	279-21514	Middle East respiratory syndrome coronavirus	MERS-CoV/KOR/Seoul/080-3-2015	orf1ab

KX034097	21456-25517	Middle East respiratory syndrome coronavirus	MERS-CoV/KOR/Seoul/080-3-2015	spike

KX034098	27853-28512	Middle East respiratory syndrome coronavirus	MERS-CoV/KOR/Seoul/162-1-2015	membrane

KX034098	28566-29807	Middle East respiratory syndrome coronavirus	MERS-CoV/KOR/Seoul/162-1-2015	nucleocapsid

KX034098	279-21514	Middle East respiratory syndrome coronavirus	MERS-CoV/KOR/Seoul/162-1-2015	orf1ab

KX034098	21456-25517	Middle East respiratory syndrome coronavirus	MERS-CoV/KOR/Seoul/162-1-2015	spike

KX034099	27853-28512	Middle East respiratory syndrome coronavirus	MERS-CoV/KOR/Seoul/169-2015	membrane

KX034099	28566-29807	Middle East respiratory syndrome coronavirus	MERS-CoV/KOR/Seoul/169-2015	nucleocapsid

KX034099	279-21514	Middle East respiratory syndrome coronavirus	MERS-CoV/KOR/Seoul/169-2015	orf1ab

KX034099	21456-25517	Middle East respiratory syndrome coronavirus	MERS-CoV/KOR/Seoul/169-2015	spike

KX034100	27853-28512	Middle East respiratory syndrome coronavirus	MERS-CoV/KOR/Seoul/177-3-2015	membrane

KX034100	28566-29807	Middle East respiratory syndrome coronavirus	MERS-CoV/KOR/Seoul/177-3-2015	nucleocapsid

KX034100	279-21514	Middle East respiratory syndrome coronavirus	MERS-CoV/KOR/Seoul/177-3-2015	orf1ab

KX034100	21456-25517	Middle East respiratory syndrome coronavirus	MERS-CoV/KOR/Seoul/177-3-2015	spike

KF192507	NA	Middle East respiratory syndrome coronavirus	Munich	membrane

KF192507	NA	Middle East respiratory syndrome coronavirus	Munich	nucleocapsid

KF192507	NA	Middle East respiratory syndrome coronavirus	Munich	orf1ab

KF192507	NA	Middle East respiratory syndrome coronavirus	Munich	spike

KJ477102	NA	Middle East respiratory syndrome coronavirus	NRCE-HKU205	membrane

KJ477102	NA	Middle East respiratory syndrome coronavirus	NRCE-HKU205	nucleocapsid

KJ477102	NA	Middle East respiratory syndrome coronavirus	NRCE-HKU205	orf1ab

KJ477102	NA	Middle East respiratory syndrome coronavirus	NRCE-HKU205	spike

KJ156934	NA	Middle East respiratory syndrome coronavirus	Riyadh_14_2013	membrane

KJ156934	NA	Middle East respiratory syndrome coronavirus	Riyadh_14_2013	nucleocapsid

KJ156934	NA	Middle East respiratory syndrome coronavirus	Riyadh_14_2013	orf1ab

KJ156934	NA	Middle East respiratory syndrome coronavirus	Riyadh_14_2013	spike

KF600612	NA	Middle East respiratory syndrome coronavirus	Riyadh_1_2012	membrane

KF600612	NA	Middle East respiratory syndrome coronavirus	Riyadh_1_2012	nucleocapsid

KF600612	NA	Middle East respiratory syndrome coronavirus	Riyadh_1_2012	orf1ab

KF600612	NA	Middle East respiratory syndrome coronavirus	Riyadh_1_2012	spike

KM027262	NA	Middle East respiratory syndrome coronavirus	Riyadh_2014KSA_683/KSA/2014	membrane

KM027262	NA	Middle East respiratory syndrome coronavirus	Riyadh_2014KSA_683/KSA/2014	nucleocapsid

KM027262	21348-25409	Middle East respiratory syndrome coronavirus	Riyadh_2014KSA_683/KSA/2014	spike

KF600652	NA	Middle East respiratory syndrome coronavirus	Riyadh_2_2012	membrane

KF600652	NA	Middle East respiratory syndrome coronavirus	Riyadh_2_2012	nucleocapsid

KF600652	NA	Middle East respiratory syndrome coronavirus	Riyadh_2_2012	orf1ab

KF600652	NA	Middle East respiratory syndrome coronavirus	Riyadh_2_2012	spike

KF600613	NA	Middle East respiratory syndrome coronavirus	Riyadh_3_2013	membrane

KF600613	NA	Middle East respiratory syndrome coronavirus	Riyadh_3_2013	nucleocapsid

KF600613	NA	Middle East respiratory syndrome coronavirus	Riyadh_3_2013	orf1ab

KF600613	NA	Middle East respiratory syndrome coronavirus	Riyadh_3_2013	spike

KJ156952	NA	Middle East respiratory syndrome coronavirus	Riyadh_4_2013	membrane

KJ156952	NA	Middle East respiratory syndrome coronavirus	Riyadh_4_2013	nucleocapsid

KJ156952	NA	Middle East respiratory syndrome coronavirus	Riyadh_4_2013	orf1ab

KJ156952	NA	Middle East respiratory syndrome coronavirus	Riyadh_4_2013	spike

KJ156944	NA	Middle East respiratory syndrome coronavirus	Riyadh_5_2013	membrane

KJ156944	NA	Middle East respiratory syndrome coronavirus	Riyadh_5_2013	nucleocapsid

KJ156944	NA	Middle East respiratory syndrome coronavirus	Riyadh_5_2013	orf1ab

KJ156944	NA	Middle East respiratory syndrome coronavirus	Riyadh_5_2013	spike

KJ156869	NA	Middle East respiratory syndrome coronavirus	Riyadh_9_2013	membrane

KJ156869	NA	Middle East respiratory syndrome coronavirus	Riyadh_9_2013	nucleocapsid

KJ156869	NA	Middle East respiratory syndrome coronavirus	Riyadh_9_2013	orf1ab

KJ156869	NA	Middle East respiratory syndrome coronavirus	Riyadh_9_2013	spike

KJ156949	NA	Middle East respiratory syndrome coronavirus	Taif_1_2013	membrane

KJ156949	NA	Middle East respiratory syndrome coronavirus	Taif_1_2013	nucleocapsid

KJ156949	NA	Middle East respiratory syndrome coronavirus	Taif_1_2013	orf1ab

KJ156949	NA	Middle East respiratory syndrome coronavirus	Taif_1_2013	spike

KJ156881	NA	Middle East respiratory syndrome coronavirus	Wadi-Ad-Dawasir_1_2013	membrane

KJ156881	NA	Middle East respiratory syndrome coronavirus	Wadi-Ad-Dawasir_1_2013	nucleocapsid

KJ156881	NA	Middle East respiratory syndrome coronavirus	Wadi-Ad-Dawasir_1_2013	orf1ab

KJ156881	NA	Middle East respiratory syndrome coronavirus	Wadi-Ad-Dawasir_1_2013	spike

JX169866	28379-29065	Murine coronavirus	JHM-WU-Dns2	membrane

JX169866	29080-30447	Murine coronavirus	JHM-WU-Dns2	nucleocapsid

JX169866	190-21731	Murine coronavirus	JHM-WU-Dns2	orf1ab

JX169866	23600-27307	Murine coronavirus	JHM-WU-Dns2	spike

JX169867	28701-29387	Murine coronavirus	JHM.WU	membrane

JX169867	29402-30769	Murine coronavirus	JHM.WU	nucleocapsid

JX169867	194-21738	Murine coronavirus	JHM.WU	orf1ab

JX169867	23922-27629	Murine coronavirus	JHM.WU	spike

KF268337	28947-29633	Murine coronavirus	MHV/BHKR_lab/USA/icA59_17Cl1/2012	membrane

KF268337	29648-31012	Murine coronavirus	MHV/BHKR_lab/USA/icA59_17Cl1/2012	nucleocapsid

KF268337	189-21724	Murine coronavirus	MHV/BHKR_lab/USA/icA59_17Cl1/2012	orf1ab

KF268337	23908-27882	Murine coronavirus	MHV/BHKR_lab/USA/icA59_17Cl1/2012	spike

KF268338	28947-29633	Murine coronavirus	MHV/BHKR_lab/USA/icA59_L94P/2012	membrane

KF268338	29648-31012	Murine coronavirus	MHV/BHKR_lab/USA/icA59_L94P/2012	nucleocapsid

KF268338	189-21724	Murine coronavirus	MHV/BHKR_lab/USA/icA59_L94P/2012	orf1ab

KF268338	23908-27882	Murine coronavirus	MHV/BHKR_lab/USA/icA59_L94P/2012	spike

KF268339	28943-29629	Murine coronavirus	MHV/BHKR_lab/USA/icA59_ns2M/2012	membrane

KF268339	29644-31008	Murine coronavirus	MHV/BHKR_lab/USA/icA59_ns2M/2012	nucleocapsid

KF268339	185-21720	Murine coronavirus	MHV/BHKR_lab/USA/icA59_ns2M/2012	orf1ab

KF268339	23904-27878	Murine coronavirus	MHV/BHKR_lab/USA/icA59_ns2M/2012	spike

KF268336	28947-29633	Murine coronavirus	MHV/BHKR_lab/USA/infA59_H126A/2012	membrane

KF268336	29648-31012	Murine coronavirus	MHV/BHKR_lab/USA/infA59_H126A/2012	nucleocapsid

KF268336	189-21724	Murine coronavirus	MHV/BHKR_lab/USA/infA59_H126A/2012	orf1ab

KF268336	23908-27882	Murine coronavirus	MHV/BHKR_lab/USA/infA59_H126A/2012	spike

FJ647225	NA	Murine coronavirus inf-MHV-A59	inf-MHV-A59	membrane

FJ647225	NA	Murine coronavirus inf-MHV-A59	inf-MHV-A59	nucleocapsid

FJ647225	NA	Murine coronavirus inf-MHV-A59	inf-MHV-A59	spike

FJ647223	NA	Murine coronavirus MHV-1	MHV-1	membrane

FJ647223	NA	Murine coronavirus MHV-1	MHV-1	nucleocapsid

FJ647223	NA	Murine coronavirus MHV-1	MHV-1	spike

FJ647224	NA	Murine coronavirus MHV-3	MHV-3	membrane

FJ647224	NA	Murine coronavirus MHV-3	MHV-3	nucleocapsid

FJ647224	NA	Murine coronavirus MHV-3	MHV-3	spike

FJ647226	NA	Murine coronavirus MHV-JHM.IA	MHV-JHM.IA	membrane

FJ647226	NA	Murine coronavirus MHV-JHM.IA	MHV-JHM.IA	nucleocapsid

FJ647226	NA	Murine coronavirus MHV-JHM.IA	MHV-JHM.IA	spike

FJ647218	NA	Murine coronavirus RA59/R13	RA59/R13	membrane

FJ647218	NA	Murine coronavirus RA59/R13	RA59/R13	nucleocapsid

FJ647218	NA	Murine coronavirus RA59/R13	RA59/R13	spike

FJ647220	NA	Murine coronavirus RA59/SJHM	RA59/SJHM	membrane

FJ647220	NA	Murine coronavirus RA59/SJHM	RA59/SJHM	nucleocapsid

FJ647220	NA	Murine coronavirus RA59/SJHM	RA59/SJHM	spike

FJ647221	NA	Murine coronavirus repA59/RJHM	repA59/RJHM	membrane

FJ647221	NA	Murine coronavirus repA59/RJHM	repA59/RJHM	nucleocapsid

FJ647221	NA	Murine coronavirus repA59/RJHM	repA59/RJHM	spike

FJ647227	NA	Murine coronavirus repJHM/RA59	repJHM/RA59	membrane

FJ647227	NA	Murine coronavirus repJHM/RA59	repJHM/RA59	nucleocapsid

FJ647227	NA	Murine coronavirus repJHM/RA59	repJHM/RA59	spike

FJ647219	NA	Murine coronavirus RJHM/A	RJHM/A	membrane

FJ647219	NA	Murine coronavirus RJHM/A	RJHM/A	nucleocapsid

FJ647219	NA	Murine coronavirus RJHM/A	RJHM/A	spike

FJ647222	NA	Murine coronavirus SA59/RJHM	SA59/RJHM	membrane

FJ647222	NA	Murine coronavirus SA59/RJHM	SA59/RJHM	nucleocapsid

FJ647222	NA	Murine coronavirus SA59/RJHM	SA59/RJHM	spike

MF618252	NA	Murine hepatitis virus	A59	orf1ab

MF618252	NA	Murine hepatitis virus	A59	membrane

MF618252	NA	Murine hepatitis virus	A59	nucleocapsid

MF618252	NA	Murine hepatitis virus	A59	spike

MF618253	NA	Murine hepatitis virus	A59	orf1ab

MF618253	NA	Murine hepatitis virus	A59	membrane

MF618253	NA	Murine hepatitis virus	A59	nucleocapsid

MF618253	NA	Murine hepatitis virus	A59	spike

FJ884686	NA	Murine hepatitis virus strain A59	A59	membrane

FJ884686	NA	Murine hepatitis virus strain A59	A59	nucleocapsid

FJ884686	NA	Murine hepatitis virus strain A59	A59	spike

FJ884687	NA	Murine hepatitis virus strain A59	A59	membrane

FJ884687	NA	Murine hepatitis virus strain A59	A59	nucleocapsid

FJ884687	NA	Murine hepatitis virus strain A59	A59	spike

AC_000192	29142-29828	Murine hepatitis virus strain JHM	UNKNOWN-AC_000192	membrane

AC_000192	29843-31210	Murine hepatitis virus strain JHM	UNKNOWN-AC_000192	nucleocapsid

AC_000192	215-21756	Murine hepatitis virus strain JHM	UNKNOWN-AC_000192	orf1ab

AC_000192	23940-28070	Murine hepatitis virus strain JHM	UNKNOWN-AC_000192	spike

JQ173883	NA	Murine hepatitis virus strain S/3239-17	MHV/S/3239-17	orf1ab

JQ173883	NA	Murine hepatitis virus strain S/3239-17	MHV/S/3239-17	membrane

JQ173883	NA	Murine hepatitis virus strain S/3239-17	MHV/S/3239-17	nucleocapsid

JQ173883	NA	Murine hepatitis virus strain S/3239-17	MHV/S/3239-17	spike

JF792616	NA	Rat coronavirus	681	orf1ab

JF792616	NA	Rat coronavirus	681	membrane

JF792616	NA	Rat coronavirus	681	nucleocapsid

JF792616	NA	Rat coronavirus	681	spike

JF792617	NA	Rat coronavirus	8190	orf1ab

JF792617	NA	Rat coronavirus	8190	membrane

JF792617	NA	Rat coronavirus	8190	nucleocapsid

JF792617	NA	Rat coronavirus	8190	spike

NC_012936	28902-29588	Rat coronavirus Parker	Parker	membrane

NC_012936	29603-30967	Rat coronavirus Parker	Parker	nucleocapsid

NC_012936	183-21700	Rat coronavirus Parker	Parker	orf1ab

NC_012936	23891-27973	Rat coronavirus Parker	Parker	spike

AF208067	28906-29592	Murine hepatitis virus	ML-10	membrane

AF208067	29607-30971	Murine hepatitis virus	ML-10	nucleocapsid

AF208067	23867-27841	Murine hepatitis virus	ML-10	spike

AF208066	28742-29428	Murine hepatitis virus	Penn 97-1	membrane

AF208066	29443-30798	Murine hepatitis virus	Penn 97-1	nucleocapsid

AF208066	23712-27677	Murine hepatitis virus	Penn 97-1	spike

AF201929	NA	Murine hepatitis virus strain 2	MHV-2	membrane

AF201929	NA	Murine hepatitis virus strain 2	MHV-2	nucleocapsid

AF201929	NA	Murine hepatitis virus strain 2	MHV-2	spike

AF207902	28906-29592	Murine hepatitis virus strain ML-11	ML-11	membrane

AF207902	29607-30962	Murine hepatitis virus strain ML-11	ML-11	nucleocapsid

AF207902	23756-27841	Murine hepatitis virus strain ML-11	ML-11	spike

KJ473820	NA	BtPa-BetaCoV/GD2013	BtPa-GD2013	orf1ab

KJ473820	NA	BtPa-BetaCoV/GD2013	BtPa-GD2013	membrane

KJ473820	NA	BtPa-BetaCoV/GD2013	BtPa-GD2013	nucleocapsid

KJ473820	NA	BtPa-BetaCoV/GD2013	BtPa-GD2013	spike

NC_009020	28166-28828	Pipistrellus bat coronavirus HKU5	HKU5-1 LMH03f	membrane

NC_009020	28878-30161	Pipistrellus bat coronavirus HKU5	HKU5-1 LMH03f	nucleocapsid

NC_009020	261-21808	Pipistrellus bat coronavirus HKU5	HKU5-1 LMH03f	orf1ab

NC_009020	21735-25793	Pipistrellus bat coronavirus HKU5	HKU5-1 LMH03f	spike

NC_017083	28760-29452	Rabbit coronavirus HKU14	HKU14-1	membrane

NC_017083	29462-30796	Rabbit coronavirus HKU14	HKU14-1	nucleocapsid

NC_017083	209-21663	Rabbit coronavirus HKU14	HKU14-1	orf1ab

NC_017083	23793-27881	Rabbit coronavirus HKU14	HKU14-1	spike

JN874562	28567-29259	Rabbit coronavirus HKU14	HKU14-10	membrane

JN874562	29269-30600	Rabbit coronavirus HKU14	HKU14-10	nucleocapsid

JN874562	209-21546	Rabbit coronavirus HKU14	HKU14-10	orf1ab

JN874562	23593-27684	Rabbit coronavirus HKU14	HKU14-10	spike

JN874560	28776-29468	Rabbit coronavirus HKU14	HKU14-3	membrane

JN874560	29478-30812	Rabbit coronavirus HKU14	HKU14-3	nucleocapsid

JN874560	NA	Rabbit coronavirus HKU14	HKU14-3	orf1ab

JN874560	23802-27893	Rabbit coronavirus HKU14	HKU14-3	spike

JN874561	28757-29449	Rabbit coronavirus HKU14	HKU14-8	membrane

JN874561	29459-30793	Rabbit coronavirus HKU14	HKU14-8	nucleocapsid

JN874561	NA	Rabbit coronavirus HKU14	HKU14-8	orf1ab

JN874561	23789-27880	Rabbit coronavirus HKU14	HKU14-8	spike

HM211100	25693-26355	Bat coronavirus HKU9-10-1	UNKNOWN-HM211100	membrane

HM211100	26418-27824	Bat coronavirus HKU9-10-1	UNKNOWN-HM211100	nucleocapsid

HM211100	229-20999	Bat coronavirus HKU9-10-1	UNKNOWN-HM211100	orf1ab

HM211100	20959-24768	Bat coronavirus HKU9-10-1	UNKNOWN-HM211100	spike

HM211101	25665-26327	Bat coronavirus HKU9-10-2	UNKNOWN-HM211101	membrane

HM211101	26390-27799	Bat coronavirus HKU9-10-2	UNKNOWN-HM211101	nucleocapsid

HM211101	228-20938	Bat coronavirus HKU9-10-2	UNKNOWN-HM211101	orf1ab

HM211101	20898-24740	Bat coronavirus HKU9-10-2	UNKNOWN-HM211101	spike

HM211098	25693-26355	Bat coronavirus HKU9-5-1	UNKNOWN-HM211098	membrane

HM211098	26418-27824	Bat coronavirus HKU9-5-1	UNKNOWN-HM211098	nucleocapsid

HM211098	229-20999	Bat coronavirus HKU9-5-1	UNKNOWN-HM211098	orf1ab

HM211098	20959-24768	Bat coronavirus HKU9-5-1	UNKNOWN-HM211098	spike

HM211099	25665-26324	Bat coronavirus HKU9-5-2	UNKNOWN-HM211099	membrane

HM211099	26387-27796	Bat coronavirus HKU9-5-2	UNKNOWN-HM211099	nucleocapsid

HM211099	228-20938	Bat coronavirus HKU9-5-2	UNKNOWN-HM211099	orf1ab

HM211099	20898-24740	Bat coronavirus HKU9-5-2	UNKNOWN-HM211099	spike

NC_009021	25689-26357	Rousettus bat coronavirus HKU9	HKU9-1 BF_005I	membrane

NC_009021	26419-27825	Rousettus bat coronavirus HKU9	HKU9-1 BF_005I	nucleocapsid

NC_009021	229-21020	Rousettus bat coronavirus HKU9	HKU9-1 BF_005I	orf1ab

NC_009021	20974-24798	Rousettus bat coronavirus HKU9	HKU9-1 BF_005I	spike

MG762674	NA	Rousettus bat coronavirus HKU9	Rousettus spp/Jinghong/2009	membrane

MG762674	NA	Rousettus bat coronavirus HKU9	Rousettus spp/Jinghong/2009	nucleocapsid

MG762674	NA	Rousettus bat coronavirus HKU9	Rousettus spp/Jinghong/2009	orf1ab

MG762674	NA	Rousettus bat coronavirus HKU9	Rousettus spp/Jinghong/2009	spike

KF294457	NA	SARS-related bat coronavirus	Longquan-140	membrane

KF294457	NA	SARS-related bat coronavirus	Longquan-140	nucleocapsid

KF294457	NA	SARS-related bat coronavirus	Longquan-140	orf1ab

KF294457	NA	SARS-related bat coronavirus	Longquan-140	spike

DQ022305	26339-27004	Bat SARS coronavirus HKU3-1	HKU3-1	membrane

DQ022305	28100-29365	Bat SARS coronavirus HKU3-1	HKU3-1	nucleocapsid

DQ022305	262-21464	Bat SARS coronavirus HKU3-1	HKU3-1	orf1ab

DQ022305	21471-25199	Bat SARS coronavirus HKU3-1	HKU3-1	spike

DQ412042	NA	Bat SARS CoV Rf1/2004	Rf1	membrane

DQ412042	NA	Bat SARS CoV Rf1/2004	Rf1	nucleocapsid

DQ412042	NA	Bat SARS CoV Rf1/2004	Rf1	spike

DQ412043	NA	Bat SARS CoV Rm1/2004	Rm1	membrane

DQ412043	NA	Bat SARS CoV Rm1/2004	Rm1	nucleocapsid

DQ412043	NA	Bat SARS CoV Rm1/2004	Rm1	spike

DQ071615	NA	Bat SARS CoV Rp3/2004	Rp3	membrane

DQ071615	28112-29377	Bat SARS CoV Rp3/2004	Rp3	nucleocapsid

DQ071615	NA	Bat SARS CoV Rp3/2004	Rp3	spike

AY291315	26398-27063	SARS coronavirus Frankfurt 1	Frankfurt 1	membrane

AY291315	28120-29388	SARS coronavirus Frankfurt 1	Frankfurt 1	nucleocapsid

AY291315	NA	SARS coronavirus Frankfurt 1	Frankfurt 1	orf1ab

AY291315	21492-25259	SARS coronavirus Frankfurt 1	Frankfurt 1	spike

KP886808	26352-27017	Bat SARS-like coronavirus YNLF_31C	YNLF_31C	membrane

KP886808	28103-29368	Bat SARS-like coronavirus YNLF_31C	YNLF_31C	nucleocapsid

KP886808	264-21484	Bat SARS-like coronavirus YNLF_31C	YNLF_31C	orf1ab

KP886808	21491-25216	Bat SARS-like coronavirus YNLF_31C	YNLF_31C	spike

KP886809	26352-27017	Bat SARS-like coronavirus YNLF_34C	YNLF_34C	membrane

KP886809	28103-29368	Bat SARS-like coronavirus YNLF_34C	YNLF_34C	nucleocapsid

KP886809	264-21484	Bat SARS-like coronavirus YNLF_34C	YNLF_34C	orf1ab

KP886809	21491-25216	Bat SARS-like coronavirus YNLF_34C	YNLF_34C	spike

KF569996	NA	Rhinolophus affinis coronavirus	LYRa11	membrane

KF569996	NA	Rhinolophus affinis coronavirus	LYRa11	nucleocapsid

KF569996	NA	Rhinolophus affinis coronavirus	LYRa11	spike

JX163924	26358-27023	SARS coronavirus	Tor2/FP1-10851	membrane

JX163924	28080-29348	SARS coronavirus	Tor2/FP1-10851	nucleocapsid

JX163924	225-21445	SARS coronavirus	Tor2/FP1-10851	orf1ab

JX163924	21452-25219	SARS coronavirus	Tor2/FP1-10851	spike

JX163927	26358-27023	SARS coronavirus	Tor2/FP1-10851	membrane

JX163927	28080-29348	SARS coronavirus	Tor2/FP1-10851	nucleocapsid

JX163927	225-21445	SARS coronavirus	Tor2/FP1-10851	orf1ab

JX163927	21452-25219	SARS coronavirus	Tor2/FP1-10851	spike

JX163925	26358-27023	SARS coronavirus	Tor2/FP1-10895	membrane

JX163925	28080-29348	SARS coronavirus	Tor2/FP1-10895	nucleocapsid

JX163925	225-21445	SARS coronavirus	Tor2/FP1-10895	orf1ab

JX163925	21452-25219	SARS coronavirus	Tor2/FP1-10895	spike

JX163928	26358-27023	SARS coronavirus	Tor2/FP1-10895	membrane

JX163928	28080-29348	SARS coronavirus	Tor2/FP1-10895	nucleocapsid

JX163928	225-21445	SARS coronavirus	Tor2/FP1-10895	orf1ab

JX163928	21452-25219	SARS coronavirus	Tor2/FP1-10895	spike

JX163923	26358-27023	SARS coronavirus	Tor2/FP1-10912	membrane

JX163923	28080-29348	SARS coronavirus	Tor2/FP1-10912	nucleocapsid

JX163923	225-21445	SARS coronavirus	Tor2/FP1-10912	orf1ab

JX163923	21452-25219	SARS coronavirus	Tor2/FP1-10912	spike

JX163926	26358-27023	SARS coronavirus	Tor2/FP1-10912	membrane

JX163926	28080-29348	SARS coronavirus	Tor2/FP1-10912	nucleocapsid

JX163926	225-21445	SARS coronavirus	Tor2/FP1-10912	orf1ab

JX163926	21452-25219	SARS coronavirus	Tor2/FP1-10912	spike

JF292922	NA	SARS coronavirus ExoN1	ExoN1 mutant	membrane

JF292922	NA	SARS coronavirus ExoN1	ExoN1 mutant	nucleocapsid

JF292922	NA	SARS coronavirus ExoN1	ExoN1 mutant	orf1ab

JF292922	NA	SARS coronavirus ExoN1	ExoN1 mutant	spike

JX162087	26378-27043	SARS coronavirus ExoN1	ExoN1 mutant	membrane

JX162087	28100-29368	SARS coronavirus ExoN1	ExoN1 mutant	nucleocapsid

JX162087	245-21465	SARS coronavirus ExoN1	ExoN1 mutant	orf1ab

JX162087	21472-25239	SARS coronavirus ExoN1	ExoN1 mutant	spike

KF514407	26378-27043	SARS coronavirus ExoN1	SARS/VeroE6_lab/USA/ExoN1_c5.7P20/2010	membrane

KF514407	28100-29368	SARS coronavirus ExoN1	SARS/VeroE6_lab/USA/ExoN1_c5.7P20/2010	nucleocapsid

KF514407	245-21465	SARS coronavirus ExoN1	SARS/VeroE6_lab/USA/ExoN1_c5.7P20/2010	orf1ab

KF514407	21472-25239	SARS coronavirus ExoN1	SARS/VeroE6_lab/USA/ExoN1_c5.7P20/2010	spike

KF514389	26378-27043	SARS coronavirus ExoN1	SARS/VeroE6_lab/USA/ExoN1_c8P10/2009	membrane

KF514389	28100-29368	SARS coronavirus ExoN1	SARS/VeroE6_lab/USA/ExoN1_c8P10/2009	nucleocapsid

KF514389	245-21465	SARS coronavirus ExoN1	SARS/VeroE6_lab/USA/ExoN1_c8P10/2009	orf1ab

KF514389	21472-25239	SARS coronavirus ExoN1	SARS/VeroE6_lab/USA/ExoN1_c8P10/2009	spike

JQ316196	NA	SARS coronavirus HKU-39849	HKU-39849	membrane

JQ316196	NA	SARS coronavirus HKU-39849	HKU-39849	nucleocapsid

JQ316196	NA	SARS coronavirus HKU-39849	HKU-39849	orf1ab

JQ316196	NA	SARS coronavirus HKU-39849	HKU-39849	spike

JN854286	NA	SARS coronavirus HKU-39849	recSARS-CoV HKU-39849	membrane

JN854286	NA	SARS coronavirus HKU-39849	recSARS-CoV HKU-39849	nucleocapsid

JN854286	NA	SARS coronavirus HKU-39849	recSARS-CoV HKU-39849	orf1ab

JN854286	NA	SARS coronavirus HKU-39849	recSARS-CoV HKU-39849	spike

HQ890543	NA	SARS coronavirus MA15	MA15	membrane

HQ890543	NA	SARS coronavirus MA15	MA15	nucleocapsid

HQ890543	NA	SARS coronavirus MA15	MA15	orf1ab

HQ890543	NA	SARS coronavirus MA15	MA15	spike

JF292909	NA	SARS coronavirus MA15	MA15	membrane

JF292909	NA	SARS coronavirus MA15	MA15	nucleocapsid

JF292909	NA	SARS coronavirus MA15	MA15	orf1ab

JF292909	NA	SARS coronavirus MA15	MA15	spike

JF292915	NA	SARS coronavirus MA15	MA15	membrane

JF292915	NA	SARS coronavirus MA15	MA15	nucleocapsid

JF292915	NA	SARS coronavirus MA15	MA15	orf1ab

JF292915	NA	SARS coronavirus MA15	MA15	spike

HQ890532	NA	SARS coronavirus MA15 ExoN1	MA15 ExoN1 mutant	membrane

HQ890532	NA	SARS coronavirus MA15 ExoN1	MA15 ExoN1 mutant	nucleocapsid

HQ890532	NA	SARS coronavirus MA15 ExoN1	MA15 ExoN1 mutant	orf1ab

HQ890532	NA	SARS coronavirus MA15 ExoN1	MA15 ExoN1 mutant	spike

HQ890534	NA	SARS coronavirus MA15 ExoN1	MA15 ExoN1 mutant	membrane

HQ890534	NA	SARS coronavirus MA15 ExoN1	MA15 ExoN1 mutant	nucleocapsid

HQ890534	NA	SARS coronavirus MA15 ExoN1	MA15 ExoN1 mutant	orf1ab

HQ890534	NA	SARS coronavirus MA15 ExoN1	MA15 ExoN1 mutant	spike

HQ890535	NA	SARS coronavirus MA15 ExoN1	MA15 ExoN1 mutant	membrane

HQ890535	NA	SARS coronavirus MA15 ExoN1	MA15 ExoN1 mutant	nucleocapsid

HQ890535	NA	SARS coronavirus MA15 ExoN1	MA15 ExoN1 mutant	orf1ab

HQ890535	NA	SARS coronavirus MA15 ExoN1	MA15 ExoN1 mutant	spike

HQ890537	NA	SARS coronavirus MA15 ExoN1	MA15 ExoN1 mutant	membrane

HQ890537	NA	SARS coronavirus MA15 ExoN1	MA15 ExoN1 mutant	nucleocapsid

HQ890537	NA	SARS coronavirus MA15 ExoN1	MA15 ExoN1 mutant	orf1ab

HQ890537	NA	SARS coronavirus MA15 ExoN1	MA15 ExoN1 mutant	spike

HQ890538	NA	SARS coronavirus MA15 ExoN1	MA15 ExoN1 mutant	membrane

HQ890538	NA	SARS coronavirus MA15 ExoN1	MA15 ExoN1 mutant	nucleocapsid

HQ890538	NA	SARS coronavirus MA15 ExoN1	MA15 ExoN1 mutant	orf1ab

HQ890538	NA	SARS coronavirus MA15 ExoN1	MA15 ExoN1 mutant	spike

JF292903	NA	SARS coronavirus MA15 ExoN1	MA15 ExoN1 mutant	membrane

JF292903	NA	SARS coronavirus MA15 ExoN1	MA15 ExoN1 mutant	nucleocapsid

JF292903	NA	SARS coronavirus MA15 ExoN1	MA15 ExoN1 mutant	orf1ab

JF292903	NA	SARS coronavirus MA15 ExoN1	MA15 ExoN1 mutant	spike

JF292904	NA	SARS coronavirus MA15 ExoN1	MA15 ExoN1 mutant	membrane

JF292904	NA	SARS coronavirus MA15 ExoN1	MA15 ExoN1 mutant	nucleocapsid

JF292904	NA	SARS coronavirus MA15 ExoN1	MA15 ExoN1 mutant	orf1ab

JF292904	NA	SARS coronavirus MA15 ExoN1	MA15 ExoN1 mutant	spike

JF292905	NA	SARS coronavirus MA15 ExoN1	MA15 ExoN1 mutant	membrane

JF292905	NA	SARS coronavirus MA15 ExoN1	MA15 ExoN1 mutant	nucleocapsid

JF292905	NA	SARS coronavirus MA15 ExoN1	MA15 ExoN1 mutant	orf1ab

JF292905	NA	SARS coronavirus MA15 ExoN1	MA15 ExoN1 mutant	spike

JF292906	NA	SARS coronavirus MA15 ExoN1	MA15 ExoN1 mutant	membrane

JF292906	NA	SARS coronavirus MA15 ExoN1	MA15 ExoN1 mutant	nucleocapsid

JF292906	NA	SARS coronavirus MA15 ExoN1	MA15 ExoN1 mutant	orf1ab

JF292906	NA	SARS coronavirus MA15 ExoN1	MA15 ExoN1 mutant	spike

MK062179	NA	SARS coronavirus Urbani	Urbani	orf1ab

MK062179	NA	SARS coronavirus Urbani	Urbani	membrane

MK062179	NA	SARS coronavirus Urbani	Urbani	nucleocapsid

MK062179	NA	SARS coronavirus Urbani	Urbani	spike

MK062180	NA	SARS coronavirus Urbani	Urbani	orf1ab

MK062180	NA	SARS coronavirus Urbani	Urbani	membrane

MK062180	NA	SARS coronavirus Urbani	Urbani	nucleocapsid

MK062180	NA	SARS coronavirus Urbani	Urbani	spike

MK062181	NA	SARS coronavirus Urbani	Urbani	orf1ab

MK062181	NA	SARS coronavirus Urbani	Urbani	membrane

MK062181	NA	SARS coronavirus Urbani	Urbani	nucleocapsid

MK062181	NA	SARS coronavirus Urbani	Urbani	spike

MK062182	NA	SARS coronavirus Urbani	Urbani	orf1ab

MK062182	NA	SARS coronavirus Urbani	Urbani	membrane

MK062182	NA	SARS coronavirus Urbani	Urbani	nucleocapsid

MK062182	NA	SARS coronavirus Urbani	Urbani	spike

MK062183	NA	SARS coronavirus Urbani	Urbani	orf1ab

MK062183	NA	SARS coronavirus Urbani	Urbani	membrane

MK062183	NA	SARS coronavirus Urbani	Urbani	nucleocapsid

MK062183	NA	SARS coronavirus Urbani	Urbani	spike

MK062184	NA	SARS coronavirus Urbani	Urbani	orf1ab

MK062184	NA	SARS coronavirus Urbani	Urbani	membrane

MK062184	NA	SARS coronavirus Urbani	Urbani	nucleocapsid

MK062184	NA	SARS coronavirus Urbani	Urbani	spike

KF514419	26378-27043	SARS coronavirus wtic-MB	SARS/VeroE6_lab/USA/WTic_c1P10/2009	membrane

KF514419	28100-29368	SARS coronavirus wtic-MB	SARS/VeroE6_lab/USA/WTic_c1P10/2009	nucleocapsid

KF514419	245-21465	SARS coronavirus wtic-MB	SARS/VeroE6_lab/USA/WTic_c1P10/2009	orf1ab

KF514419	21472-25239	SARS coronavirus wtic-MB	SARS/VeroE6_lab/USA/WTic_c1P10/2009	spike

JF292921	NA	SARS coronavirus wtic-MB	WTic	membrane

JF292921	NA	SARS coronavirus wtic-MB	WTic	nucleocapsid

JF292921	NA	SARS coronavirus wtic-MB	WTic	orf1ab

JF292921	NA	SARS coronavirus wtic-MB	WTic	spike

KY352407	26308-26973	Severe acute respiratory syndrome-related coronavirus	BtKY72	membrane

KY352407	27665-28924	Severe acute respiratory syndrome-related coronavirus	BtKY72	nucleocapsid

KY352407	260-21411	Severe acute respiratory syndrome-related coronavirus	BtKY72	orf1ab

KY352407	21418-25191	Severe acute respiratory syndrome-related coronavirus	BtKY72	spike

KJ473822	NA	BtTp-BetaCoV/GX2012	BtTp-GX2012	orf1ab

KJ473822	NA	BtTp-BetaCoV/GX2012	BtTp-GX2012	membrane

KJ473822	NA	BtTp-BetaCoV/GX2012	BtTp-GX2012	nucleocapsid

KJ473822	NA	BtTp-BetaCoV/GX2012	BtTp-GX2012	spike

NC_009019	28000-28659	Tylonycteris bat coronavirus HKU4	HKU4-1 B04f	membrane

NC_009019	28697-29968	Tylonycteris bat coronavirus HKU4	HKU4-1 B04f	nucleocapsid

NC_009019	267-21625	Tylonycteris bat coronavirus HKU4	HKU4-1 B04f	orf1ab

NC_009019	21570-25628	Tylonycteris bat coronavirus HKU4	HKU4-1 B04f	spike

JX993988	NA	Bat coronavirus Cp/Yunnan2011	Cp/Yunnan2011	orf1ab

JX993987	NA	Bat coronavirus Rp/Shaanxi2011	Rp/Shaanxi2011	orf1ab

KU886219	NA	Bovine coronavirus	BCV-AKS-01	orf1ab

EF424619	195-13346	Bovine coronavirus E-AH187	E-AH187	orf1ab

EF424615	211-13362	Bovine coronavirus E-AH65	E-AH65	orf1ab

EF424616	195-13346	Bovine coronavirus E-AH65-TC	E-AH65-TC	orf1ab

EF424620	195-13346	Bovine coronavirus R-AH187	R-AH187	orf1ab

EF424617	195-13346	Bovine coronavirus R-AH65	R-AH65	orf1ab

EF424618	195-13346	Bovine coronavirus R-AH65-TC	R-AH65-TC	orf1ab

EF424624	195-13346	Calf-giraffe coronavirus US/OH3/2006	US/OH3/2006	orf1ab

EF424622	195-13346	Giraffe coronavirus US/OH3-TC/2006	US/OH3-TC/2006	orf1ab

EF424623	195-13346	Giraffe coronavirus US/OH3/2003	US/OH3/2003	orf1ab

EF424621	196-13347	Sable antelope coronavirus US/OH1/2003	US/OH1/2003	orf1ab

FJ425188	NA	Sambar deer coronavirus US/OH-WD388-TC/1994	UNKNOWN-FJ425188	orf1ab

FJ425190	NA	Sambar deer coronavirus US/OH-WD388-TC/1994	UNKNOWN-FJ425190	orf1ab

FJ425189	NA	Sambar deer coronavirus US/OH-WD388/1994	UNKNOWN-FJ425189	orf1ab

FJ425185	NA	Waterbuck coronavirus US/OH-WD358-GnC/1994	UNKNOWN-FJ425185	orf1ab

FJ425184	NA	Waterbuck coronavirus US/OH-WD358-TC/1994	UNKNOWN-FJ425184	orf1ab

FJ425186	NA	Waterbuck coronavirus US/OH-WD358/1994	UNKNOWN-FJ425186	orf1ab

FJ425187	NA	White-tailed deer coronavirus US/OH-WD470/1994	UNKNOWN-FJ425187	orf1ab

MH810163	NA	Yak coronavirus	YAK/HY24/CH/2017	orf1ab

AF220295	NA	Bovine coronavirus	Quebec	orf1ab

KJ361500	NA	Middle East respiratory syndrome coronavirus	Hu-France (UAE) - FRA1_1627-2013_BAL_Sanger	orf1ab

KJ361502	NA	Middle East respiratory syndrome coronavirus	Hu-France - FRA2_130569-2013_InSpu_Sanger	orf1ab

KJ361501	NA	Middle East respiratory syndrome coronavirus	Hu-France - FRA2_130569-2013_IS_HTS	orf1ab

KJ361503	NA	Middle East respiratory syndrome coronavirus	Hu-France - FRA2_130569-2013_Isolate_Sanger	orf1ab

KM027260	NA	Middle East respiratory syndrome coronavirus	Jeddah_C10306/KSA/2014-04-20	orf1ab

KM027255	NA	Middle East respiratory syndrome coronavirus	Jeddah_C7149/KSA/2014-04-05	orf1ab

KM027256	NA	Middle East respiratory syndrome coronavirus	Jeddah_C7569/KSA/2014-04-03	orf1ab

KM027257	NA	Middle East respiratory syndrome coronavirus	Jeddah_C7770/KSA/2014-04-07	orf1ab

KM027258	NA	Middle East respiratory syndrome coronavirus	Jeddah_C8826/KSA/2014-04-12	orf1ab

KM027259	NA	Middle East respiratory syndrome coronavirus	Jeddah_C9055/KSA/2014-04-14	orf1ab

KM027261	NA	Middle East respiratory syndrome coronavirus	Makkah_C9355/KSA/Makkah/2014-04-15	orf1ab

KM027262	NA	Middle East respiratory syndrome coronavirus	Riyadh_2014KSA_683/KSA/2014	orf1ab

FJ647225	NA	Murine coronavirus inf-MHV-A59	inf-MHV-A59	orf1ab

FJ647223	NA	Murine coronavirus MHV-1	MHV-1	orf1ab

FJ647224	NA	Murine coronavirus MHV-3	MHV-3	orf1ab

FJ647226	NA	Murine coronavirus MHV-JHM.IA	MHV-JHM.IA	orf1ab

FJ647218	NA	Murine coronavirus RA59/R13	RA59/R13	orf1ab

FJ647220	NA	Murine coronavirus RA59/SJHM	RA59/SJHM	orf1ab

FJ647221	NA	Murine coronavirus repA59/RJHM	repA59/RJHM	orf1ab

FJ647227	NA	Murine coronavirus repJHM/RA59	repJHM/RA59	orf1ab

FJ647219	NA	Murine coronavirus RJHM/A	RJHM/A	orf1ab

FJ647222	NA	Murine coronavirus SA59/RJHM	SA59/RJHM	orf1ab

FJ884686	NA	Murine hepatitis virus strain A59	A59	orf1ab

FJ884687	NA	Murine hepatitis virus strain A59	A59	orf1ab

AF208067	210-13613	Murine hepatitis virus	ML-10	orf1ab

AF208066	210-13460	Murine hepatitis virus	Penn 97-1	orf1ab

AF201929	NA	Murine hepatitis virus strain 2	MHV-2	orf1ab

AF207902	210-13460	Murine hepatitis virus strain ML-11	ML-11	orf1ab

DQ412042	NA	Bat SARS CoV Rf1/2004	Rf1	orf1ab

DQ412043	NA	Bat SARS CoV Rm1/2004	Rm1	orf1ab

DQ071615	NA	Bat SARS CoV Rp3/2004	Rp3	orf1ab

KF569996	NA	Rhinolophus affinis coronavirus	LYRa11	orf1ab

MN908947	NA	Wuhan seafood market pneumonia virus	SARS-CoV-2	orf1ab

BetaCoV/Wuhan/IVDC-HB-01/2019|EPI_ISL_402119	NA	Wuhan	SARS-CoV-2	orf1ab

BetaCoV/Wuhan/IVDC-HB-04/2020|EPI_ISL_402120	NA	Wuhan	SARS-CoV-2	orf1ab

BetaCoV/Wuhan/IVDC-HB-05/2019|EPI_ISL_402121	NA	Wuhan	SARS-CoV-2	orf1ab

BetaCoV/Wuhan/IPBCAMS-WH-01/2019|EPI_ISL_402123	NA	Wuhan	SARS-CoV-2	orf1ab

BetaCoV/Wuhan/WIV04/2019|EPI_ISL_402124	NA	Wuhan	SARS-CoV-2	orf1ab

BetaCoV/Wuhan-Hu-1/2019|EPI_ISL_402125	NA	Wuhan	SARS-CoV-2	orf1ab

BetaCoV/Wuhan/WIV02/2019|EPI_ISL_402127	NA	Wuhan	SARS-CoV-2	orf1ab

BetaCoV/Wuhan/WIV05/2019|EPI_ISL_402128	NA	Wuhan	SARS-CoV-2	orf1ab

BetaCoV/Wuhan/WIV06/2019|EPI_ISL_402129	NA	Wuhan	SARS-CoV-2	orf1ab

BetaCoV/Wuhan/WIV07/2019|EPI_ISL_402130	NA	Wuhan	SARS-CoV-2	orf1ab

BetaCoV/Wuhan/HBCDC-HB-01/2019|EPI_ISL_402132	NA	Wuhan	SARS-CoV-2	orf1ab

BetaCoV/Wuhan/IPBCAMS-WH-04/2019|EPI_ISL_403929	NA	Wuhan	SARS-CoV-2	orf1ab

BetaCoV/Wuhan/IPBCAMS-WH-03/2019|EPI_ISL_403930	NA	Wuhan	SARS-CoV-2	orf1ab

BetaCoV/Wuhan/IPBCAMS-WH-02/2019|EPI_ISL_403931	NA	Wuhan	SARS-CoV-2	orf1ab

BetaCoV/Nonthaburi/61/2020|EPI_ISL_403962	NA	Wuhan	SARS-CoV-2	orf1ab

BetaCoV/Nonthaburi/74/2020|EPI_ISL_403963	NA	Wuhan	SARS-CoV-2	orf1ab

BetaCoV/Zhejiang/WZ-01/2020|EPI_ISL_404227	NA	Wuhan	SARS-CoV-2	orf1ab

BetaCoV/Zhejiang/WZ-02/2020|EPI_ISL_404228	NA	Wuhan	SARS-CoV-2	orf1ab

MN908947	NA	Wuhan seafood market pneumonia virus	SARS-CoV-2	spike

BetaCoV/Wuhan/IVDC-HB-01/2019|EPI_ISL_402119	NA	Wuhan	SARS-CoV-2	spike

BetaCoV/Wuhan/IVDC-HB-04/2020|EPI_ISL_402120	NA	Wuhan	SARS-CoV-2	spike

BetaCoV/Wuhan/IVDC-HB-05/2019|EPI_ISL_402121	NA	Wuhan	SARS-CoV-2	spike

BetaCoV/Wuhan/IPBCAMS-WH-01/2019|EPI_ISL_402123	NA	Wuhan	SARS-CoV-2	spike

BetaCoV/Wuhan/WIV04/2019|EPI_ISL_402124	NA	Wuhan	SARS-CoV-2	spike

BetaCoV/Wuhan-Hu-1/2019|EPI_ISL_402125	NA	Wuhan	SARS-CoV-2	spike

BetaCoV/Wuhan/WIV02/2019|EPI_ISL_402127	NA	Wuhan	SARS-CoV-2	spike

BetaCoV/Wuhan/WIV05/2019|EPI_ISL_402128	NA	Wuhan	SARS-CoV-2	spike

BetaCoV/Wuhan/WIV06/2019|EPI_ISL_402129	NA	Wuhan	SARS-CoV-2	spike

BetaCoV/Wuhan/WIV07/2019|EPI_ISL_402130	NA	Wuhan	SARS-CoV-2	spike

BetaCoV/Wuhan/HBCDC-HB-01/2019|EPI_ISL_402132	NA	Wuhan	SARS-CoV-2	spike

BetaCoV/Wuhan/IPBCAMS-WH-04/2019|EPI_ISL_403929	NA	Wuhan	SARS-CoV-2	spike

BetaCoV/Wuhan/IPBCAMS-WH-03/2019|EPI_ISL_403930	NA	Wuhan	SARS-CoV-2	spike

BetaCoV/Wuhan/IPBCAMS-WH-02/2019|EPI_ISL_403931	NA	Wuhan	SARS-CoV-2	spike

BetaCoV/Nonthaburi/61/2020|EPI_ISL_403962	NA	Wuhan	SARS-CoV-2	spike

BetaCoV/Nonthaburi/74/2020|EPI_ISL_403963	NA	Wuhan	SARS-CoV-2	spike

BetaCoV/Zhejiang/WZ-01/2020|EPI_ISL_404227	NA	Wuhan	SARS-CoV-2	spike

BetaCoV/Zhejiang/WZ-02/2020|EPI_ISL_404228	NA	Wuhan	SARS-CoV-2	spike

MN908947	NA	Wuhan seafood market pneumonia virus	SARS-CoV-2	membrane

BetaCoV/Wuhan/IVDC-HB-01/2019|EPI_ISL_402119	NA	Wuhan	SARS-CoV-2	membrane

BetaCoV/Wuhan/IVDC-HB-04/2020|EPI_ISL_402120	NA	Wuhan	SARS-CoV-2	membrane

BetaCoV/Wuhan/IVDC-HB-05/2019|EPI_ISL_402121	NA	Wuhan	SARS-CoV-2	membrane

BetaCoV/Wuhan/IPBCAMS-WH-01/2019|EPI_ISL_402123	NA	Wuhan	SARS-CoV-2	membrane

BetaCoV/Wuhan/WIV04/2019|EPI_ISL_402124	NA	Wuhan	SARS-CoV-2	membrane

BetaCoV/Wuhan-Hu-1/2019|EPI_ISL_402125	NA	Wuhan	SARS-CoV-2	membrane

BetaCoV/Wuhan/WIV02/2019|EPI_ISL_402127	NA	Wuhan	SARS-CoV-2	membrane

BetaCoV/Wuhan/WIV05/2019|EPI_ISL_402128	NA	Wuhan	SARS-CoV-2	membrane

BetaCoV/Wuhan/WIV06/2019|EPI_ISL_402129	NA	Wuhan	SARS-CoV-2	membrane

BetaCoV/Wuhan/WIV07/2019|EPI_ISL_402130	NA	Wuhan	SARS-CoV-2	membrane

BetaCoV/Wuhan/HBCDC-HB-01/2019|EPI_ISL_402132	NA	Wuhan	SARS-CoV-2	membrane

BetaCoV/Wuhan/IPBCAMS-WH-04/2019|EPI_ISL_403929	NA	Wuhan	SARS-CoV-2	membrane

BetaCoV/Wuhan/IPBCAMS-WH-03/2019|EPI_ISL_403930	NA	Wuhan	SARS-CoV-2	membrane

BetaCoV/Wuhan/IPBCAMS-WH-02/2019|EPI_ISL_403931	NA	Wuhan	SARS-CoV-2	membrane

BetaCoV/Nonthaburi/61/2020|EPI_ISL_403962	NA	Wuhan	SARS-CoV-2	membrane

BetaCoV/Nonthaburi/74/2020|EPI_ISL_403963	NA	Wuhan	SARS-CoV-2	membrane

BetaCoV/Zhejiang/WZ-01/2020|EPI_ISL_404227	NA	Wuhan	SARS-CoV-2	membrane

BetaCoV/Zhejiang/WZ-02/2020|EPI_ISL_404228	NA	Wuhan	SARS-CoV-2	membrane

MN908947	NA	Wuhan seafood market pneumonia virus	SARS-CoV-2	nucleocapsid

BetaCoV/Wuhan/IVDC-HB-01/2019|EPI_ISL_402119	NA	Wuhan	SARS-CoV-2	nucleocapsid

BetaCoV/Wuhan/IVDC-HB-04/2020|EPI_ISL_402120	NA	Wuhan	SARS-CoV-2	nucleocapsid

BetaCoV/Wuhan/IVDC-HB-05/2019|EPI_ISL_402121	NA	Wuhan	SARS-CoV-2	nucleocapsid

BetaCoV/Wuhan/IPBCAMS-WH-01/2019|EPI_ISL_402123	NA	Wuhan	SARS-CoV-2	nucleocapsid

BetaCoV/Wuhan/WIV04/2019|EPI_ISL_402124	NA	Wuhan	SARS-CoV-2	nucleocapsid

BetaCoV/Wuhan-Hu-1/2019|EPI_ISL_402125	NA	Wuhan	SARS-CoV-2	nucleocapsid

BetaCoV/Wuhan/WIV02/2019|EPI_ISL_402127	NA	Wuhan	SARS-CoV-2	nucleocapsid

BetaCoV/Wuhan/WIV05/2019|EPI_ISL_402128	NA	Wuhan	SARS-CoV-2	nucleocapsid

BetaCoV/Wuhan/WIV06/2019|EPI_ISL_402129	NA	Wuhan	SARS-CoV-2	nucleocapsid

BetaCoV/Wuhan/WIV07/2019|EPI_ISL_402130	NA	Wuhan	SARS-CoV-2	nucleocapsid

BetaCoV/Wuhan/HBCDC-HB-01/2019|EPI_ISL_402132	NA	Wuhan	SARS-CoV-2	nucleocapsid

BetaCoV/Wuhan/IPBCAMS-WH-04/2019|EPI_ISL_403929	NA	Wuhan	SARS-CoV-2	nucleocapsid

BetaCoV/Wuhan/IPBCAMS-WH-03/2019|EPI_ISL_403930	NA	Wuhan	SARS-CoV-2	nucleocapsid

BetaCoV/Wuhan/IPBCAMS-WH-02/2019|EPI_ISL_403931	NA	Wuhan	SARS-CoV-2	nucleocapsid

BetaCoV/Nonthaburi/61/2020|EPI_ISL_403962	NA	Wuhan	SARS-CoV-2	nucleocapsid

BetaCoV/Nonthaburi/74/2020|EPI_ISL_403963	NA	Wuhan	SARS-CoV-2	nucleocapsid

BetaCoV/Zhejiang/WZ-01/2020|EPI_ISL_404227	NA	Wuhan	SARS-CoV-2	nucleocapsid

BetaCoV/Zhejiang/WZ-02/2020|EPI_ISL_404228	NA	Wuhan	SARS-CoV-2	nucleocapsid

GU190215	NA	BM48-31/BGR/2008	BM48-31	orf1ab

GU190215	NA	BM48-31/BGR/2008	BM48-31	spike

GU190215	NA	BM48-31/BGR/2008	BM48-31	membrane

GU190215	NA	BM48-31/BGR/2008	BM48-31	nucleocapsid

MG772933	NA	bat-SL-CoVZC45	CoVZC45	orf1ab

MG772933	NA	bat-SL-CoVZC45	CoVZC45	spike

MG772933	NA	bat-SL-CoVZC45	CoVZC45	membrane

MG772933	NA	bat-SL-CoVZC45	CoVZC45	nucleocapsid

MG772934	NA	bat-SL-CoVZXC21	CoVZXC21	orf1ab

MG772934	NA	bat-SL-CoVZXC21	CoVZXC21	spike

MG772934	NA	bat-SL-CoVZXC21	CoVZXC21	membrane

MG772934	NA	bat-SL-CoVZXC21	CoVZXC21	nucleocapsid

EPI_ISL_402131	NA	bat/Yunnan/RaTG13/2013	bat/RaTG13/2013	orf1ab

EPI_ISL_402131	NA	bat/Yunnan/RaTG13/2013	bat/RaTG13/2013	spike

EPI_ISL_402131	NA	bat/Yunnan/RaTG13/2013	bat/RaTG13/2013	membrane

EPI_ISL_402131	NA	bat/Yunnan/RaTG13/2013	bat/RaTG13/2013	nucleocapsid

EPI_ISL_410539	NA	pangolin/Guangxi/P1E/2017	pangolin/P1E/2017	orf1ab

EPI_ISL_410539	NA	pangolin/Guangxi/P1E/2017	pangolin/P1E/2017	spike

EPI_ISL_410539	NA	pangolin/Guangxi/P1E/2017	pangolin/P1E/2017	membrane

EPI_ISL_410539	NA	pangolin/Guangxi/P1E/2017	pangolin/P1E/2017	nucleocapsid

Table S2. List of the authors of the sequences in this study shared by GISAID Initiative

| **Accession ID** | **Virus name** | **Location** | **Collection date** | **Originating lab** | **Submitting lab** | **Authors** |
| --- | --- | --- | --- | --- | --- | --- |
| [EPI_ISL_410539](https://platform.gisaid.org/epi3/start/CoV2020) | BetaCoV/pangolin/Guangxi/P1E/2017 | Asia / China / Guangxi | 2017 | Beijing Institute of Microbiology and Epidemiology | Beijing Institute of Microbiology and Epidemiology | Wu-Chun Cao; Tommy Tsan-Yuk Lam; Na Jia; Ya-Wei Zhang; Jia-Fu Jiang; Bao-Gui Jiang |
| EPI_ISL_402119 | BetaCoV/Wuhan/IVDC-HB-01/2019 | Asia / China / Hubei / Wuhan | 12/30/2019 | National Institute for Viral Disease Control and Prevention, China CDC | National Institute for Viral Disease Control and Prevention, China CDC | Wenjie Tan，Xiang Zhao，Wenling Wang，Xuejun Ma，Yongzhong Jiang，Roujian Lu, Ji Wang, Weimin Zhou，Peihua Niu，Peipei Liu，Faxian Zhan，Weifeng Shi，Baoying Huang，Jun Liu，Li Zhao，Yao Meng，Xiaozhou He，Fei Ye，Na Zhu，Yang Li，Jing Chen，Wenbo Xu，George F. Gao，Guizhen Wu |
| EPI_ISL_402120 | BetaCoV/Wuhan/IVDC-HB-04/2020 | Asia / China / Hubei / Wuhan | 1/1/2020 | National Institute for Viral Disease Control and Prevention, China CDC | National Institute for Viral Disease Control and Prevention, China CDC | Wenjie Tan，Xiang Zhao，Wenling Wang，Xuejun Ma，Yongzhong Jiang，Roujian Lu，Ji Wang，Weimin Zhou，Peihua Niu，Peipei Liu，Faxian Zhan，Weifeng Shi，Baoying Huang，Jun Liu，Li Zhao，Yao Meng，Xiaozhou He，Fei Ye，Na Zhu，Yang Li，Jing Chen，Wenbo Xu，George F. Gao，Guizhen Wu |
| EPI_ISL_402121 | BetaCoV/Wuhan/IVDC-HB-05/2019 | Asia / China / Hubei / Wuhan | 12/30/2019 | National Institute for Viral Disease Control and Prevention, China CDC | National Institute for Viral Disease Control and Prevention, China CDC | Wenjie Tan，Xuejun Ma，Xiang Zhao，Wenling Wang，Yongzhong Jiang，Roujian Lu，Ji Wang，Peihua Niu, Weimin Zhou, Faxian Zhan，Weifeng Shi，Baoying Huang，Jun Liu，Li Zhao，Yao Meng，Fei Ye，Na Zhu, Xiaozhou He，Peipei Liu, Yang Li，Jing Chen，Wenbo Xu，George F. Gao，Guizhen Wu |
| EPI_ISL_402123 | BetaCoV/Wuhan/IPBCAMS-WH-01/2019 | Asia / China / Hubei / Wuhan | 12/24/2019 | Institute of Pathogen Biology, Chinese Academy of Medical Sciences & Peking Union Medical College | Institute of Pathogen Biology, Chinese Academy of Medical Sciences & Peking Union Medical College | Lili Ren, Jianwei Wang, Qi Jin, Zichun Xiang, Zhiqiang Wu, Chao Wu, Yiwei Liu |
| EPI_ISL_402124 | BetaCoV/Wuhan/WIV04/2019 | Asia / China / Hubei / Wuhan | 12/30/2019 | Wuhan Jinyintan Hospital | Wuhan Institute of Virology, Chinese Academy of Sciences | Peng Zhou, Xing-Lou Yang, Ding-Yu Zhang, Lei Zhang, Yan Zhu, Hao-Rui Si, Zhengli Shi |
| EPI_ISL_402125 | BetaCoV/Wuhan-Hu-1/2019 | Asia / China | 12/31/2019 | unknown | National Institute for Communicable Disease Control and Prevention (ICDC) Chinese Center for Disease Control and Prevention (China CDC) | Zhang,Y.-Z., Wu,F., Chen,Y.-M., Pei,Y.-Y., Xu,L., Wang,W., Zhao,S., Yu,B., Hu,Y., Tao,Z.-W., Song,Z.-G., Tian,J.-H., Zhang,Y.-L., Liu,Y., Zheng,J.-J., Dai,F.-H., Wang,Q.-M., She,J.-L. and Zhu,T.-Y. |
| EPI_ISL_402127 | BetaCoV/Wuhan/WIV02/2019 | Asia / China / Hubei / Wuhan | 12/30/2019 | Wuhan Jinyintan Hospital | Wuhan Institute of Virology, Chinese Academy of Sciences | Peng Zhou, Xing-Lou Yang, Ding-Yu Zhang, Lei Zhang, Yan Zhu, Hao-Rui Si, Zhengli Shi |
| EPI_ISL_402128 | BetaCoV/Wuhan/WIV05/2019 | Asia / China / Hubei / Wuhan | 12/30/2019 | Wuhan Jinyintan Hospital | Wuhan Institute of Virology, Chinese Academy of Sciences | Peng Zhou, Xing-Lou Yang, Ding-Yu Zhang, Lei Zhang, Yan Zhu, Hao-Rui Si, Zhengli Shi |
| EPI_ISL_402129 | BetaCoV/Wuhan/WIV06/2019 | Asia / China / Hubei / Wuhan | 12/30/2019 | Wuhan Jinyintan Hospital | Wuhan Institute of Virology, Chinese Academy of Sciences | Peng Zhou, Xing-Lou Yang, Ding-Yu Zhang, Lei Zhang, Yan Zhu, Hao-Rui Si, Zhengli Shi |
| EPI_ISL_402130 | BetaCoV/Wuhan/WIV07/2019 | Asia / China / Hubei / Wuhan | 12/30/2019 | Wuhan Jinyintan Hospital | Wuhan Institute of Virology, Chinese Academy of Sciences | Peng Zhou, Xing-Lou Yang, Ding-Yu Zhang, Lei Zhang, Yan Zhu, Hao-Rui Si, Zhengli Shi |
| EPI_ISL_402132 | BetaCoV/Wuhan/HBCDC-HB-01/2019 | Asia / China / Hubei / Wuhan | 12/30/2019 | Wuhan Jinyintan Hospital | Hubei Provincial Center for Disease Control and Prevention | Bin Fang, Xiang Li, Xiao Yu, Linlin Liu, Bo Yang, Faxian Zhan, Guojun Ye, Xixiang Huo, Junqiang Xu, Bo Yu, Kun Cai, Jing Li, Yongzhong Jiang. |
| EPI_ISL_403929 | BetaCoV/Wuhan/IPBCAMS-WH-04/2019 | Asia / China / Hubei / Wuhan | 12/30/2019 | Institute of Pathogen Biology, Chinese Academy of Medical Sciences & Peking Union Medical College | Institute of Pathogen Biology, Chinese Academy of Medical Sciences & Peking Union Medical College | Lili Ren, Jianwei Wang, Qi Jin, Zichun Xiang, Zhiqiang Wu, Chao Wu, Yiwei Liu |
| EPI_ISL_403930 | BetaCoV/Wuhan/IPBCAMS-WH-03/2019 | Asia / China / Hubei / Wuhan | 12/30/2019 | Institute of Pathogen Biology, Chinese Academy of Medical Sciences & Peking Union Medical College | Institute of Pathogen Biology, Chinese Academy of Medical Sciences & Peking Union Medical College | Lili Ren, Jianwei Wang, Qi Jin, Zichun Xiang, Zhiqiang Wu, Chao Wu, Yiwei Liu |
| EPI_ISL_403931 | BetaCoV/Wuhan/IPBCAMS-WH-02/2019 | Asia / China / Hubei / Wuhan | 12/30/2019 | Institute of Pathogen Biology, Chinese Academy of Medical Sciences & Peking Union Medical College | Institute of Pathogen Biology, Chinese Academy of Medical Sciences & Peking Union Medical College | Lili Ren, Jianwei Wang, Qi Jin, Zichun Xiang, Zhiqiang Wu, Chao Wu, Yiwei Liu |
| EPI_ISL_403962 | BetaCoV/Nonthaburi/61/2020 | Asia / Thailand / Nonthaburi | 1/8/2020 | Bamrasnaradura Hospital | 1. Department of Medical Sciences, Ministry of Public Health, Thailand 2. Thai Red Cross Emerging Infectious Diseases - Health Science Centre 3. Department of Disease Control, Ministry of Public Health, Thailand | Pilailuk,Okada; Siripaporn,Phuygun; Thanutsapa,Thanadachakul; Supaporn,Wacharapluesadee; Sittiporn,Parnmen; Warawan,Wongboot; Sunthareeya,Waicharoen; Rome,Buathong; Malinee,Chittaganpitch; Nanthawan,Mekha |
| EPI_ISL_403963 | BetaCoV/Nonthaburi/74/2020 | Asia / Thailand / Nonthaburi | 1/13/2020 | Bamrasnaradura Hospital | 1. Department of Medical Sciences, Ministry of Public Health, Thailand 2. Thai Red Cross Emerging Infectious Diseases - Health Science Centre 3. Department of Disease Control, Ministry of Public Health, Thailand | Pilailuk,Okada; Siripaporn,Phuygun; Thanutsapa,Thanadachakul; Supaporn,Wacharapluesadee; Sittiporn,Parnmen; Warawan,Wongboot; Sunthareeya,Waicharoen; Rome,Buathong; Malinee,Chittaganpitch; Nanthawan,Mekha |
| EPI_ISL_404227 | BetaCoV/Zhejiang/WZ-01/2020 | Asia / China / Zhejiang | 1/16/2020 | Zhejiang Provincial Center for Disease Control and Prevention | Department of Microbiology, Zhejiang Provincial Center for Disease Control and Prevention | Yin Chen, Yanjun Zhang, Haiyan Mao, Junhang Pan, Xiuyu Lou, Yiyu Lu, Juying Yan, Hanping Zhu, Jian Gao, Yan Feng, Yi Sun, Hao Yan, Zhen Li, Yisheng Sun, Liming Gong, Qiong Ge, Wen Shi, Xinying Wang, Wenwu Yao, Zhangnv Yang, Fang Xu, Chen Chen, Enfu Chen, Zhen Wang, Zhiping Chen, Jianmin Jiang, Chonggao Hu |
| EPI_ISL_404228 | BetaCoV/Zhejiang/WZ-02/2020 | Asia / China / Zhejiang | 1/17/2020 | Zhejiang Provincial Center for Disease Control and Prevention | Department of Microbiology, Zhejiang Provincial Center for Disease Control and Prevention | Yanjun Zhang, Yin Chen, Haiyan Mao, Junhang Pan, Xiuyu Lou, Yiyu Lu, Juying Yan, Hanping Zhu, Jian Gao, Yan Feng, Yi Sun, Hao Yan, Zhen Li, Yisheng Sun, Liming Gong, Qiong Ge, Wen Shi, Xinying Wang, Wenwu Yao, Zhangnv Yang, Fang Xu, Chen Chen, Enfu Chen, Zhen Wang, Zhiping Chen, Jianmin Jiang, Chonggao Hu |
